# Molecular Signatures of Normal Pressure Hydrocephalus: A Large-scale Proteomic Analysis of Cerebrospinal Fluid

**DOI:** 10.1101/2024.03.01.583014

**Authors:** Aida Kamalian, Siavash Shirzadeh Barough, Sara G. Ho, Marilyn Albert, Mark G. Luciano, Sevil Yasar, Abhay Moghekar

## Abstract

Given the persistent challenge of differentiating idiopathic Normal Pressure Hydrocephalus (iNPH) from similar clinical entities, we conducted an in-depth proteomic study of cerebrospinal fluid (CSF) in 28 shunt-responsive iNPH patients, 38 Mild Cognitive Impairment (MCI) due to Alzheimer’s disease, and 49 healthy controls. Utilizing the Olink Explore 3072 panel, we identified distinct proteomic profiles in iNPH that highlight significant downregulation of synaptic markers and cell-cell adhesion proteins. Alongside vimentin and inflammatory markers upregulation, these results suggest ependymal layer and transependymal flow dysfunction. Moreover, downregulation of multiple proteins associated with congenital hydrocephalus (e.g., L1CAM, PCDH9, ISLR2, ADAMTSL2, and B4GAT1) points to a possible shared molecular foundation between congenital hydrocephalus and iNPH. Through orthogonal partial least squares discriminant analysis (OPLS-DA), a panel comprising 13 proteins has been identified as potential diagnostic biomarkers of iNPH, pending external validation. These findings offer novel insights into the pathophysiology of iNPH, with implications for improved diagnosis.

## 1. Introduction

Idiopathic Normal Pressure Hydrocephalus (iNPH) is one of the few treatable causes of dementia that is characterized by the triad of gait disturbances, dementia, and urinary incontinence^1^. Previous epidemiologic studies have reported a prevalence of 1.5% among people aged 70-79 and 5.9% in population aged 80 and older^2,3^. Despite potential reversal of symptoms via shunt surgery, iNPH remains underdiagnosed and often misdiagnosed as Alzheimer’s disease (AD) or a Parkinsonian syndrome, highlighting the need for development of reliable, specific diagnostic biomarkers^2–5^.

Recent advancements in diagnostic techniques, particularly in neuroimaging and cerebrospinal fluid dynamics, have enhanced the accuracy of iNPH diagnosis^6–10^. However, a definitive diagnosis remains elusive, relying heavily on clinical judgment and the exclusion of other conditions^10^.

Previous studies have mainly focused on the potential of classical AD cerebrospinal fluid (CSF) biomarkers in differential diagnosis of iNPH from similar clinical entities and long-term response after shunt surgery^11,12^. Lower levels of neurofilament light (NfL), amyloidβ (Aβ ), amyloidβ _1-40_ (Aβ_1–40_), total tau (t-tau), and phosphorylated tau 181 (p-Tau181) have been associated with long-term improvement in gait outcomes after shunt surgery^11–13^. Moreover, high concentrations of CSF Aβ1-42 with a high Aβ_1-42_/Aβ_1-40_ ratio, low p-tau181, and low t-tau levels is a typical CSF profile of iNPH, while lower Aβ_1-42_/Aβ_1-40_, especially when associated with high p-tau levels, may indicate coexistent AD pathology^12,14^.

Weiner et al. performed a deep proteomics analysis of CSF in responsive and unresponsive iNPH patients and identified potential prognostic biomarkers for shunt responsiveness in iNPH, notably FABP3, ANXA4, B3GAT2, ITGB1, YWHAG^13^. However, they did not investigate iNPH-specific proteomics compared to healthy controls or other differential diagnoses^13^. In another investigation, Torreta et al. employed mass spectrometry to examine the sphingolipid profile and proteomics of CSF in individuals with iNPH, contrasting these with AD patients and healthy subjects^10^. This research identified intriguing potential biomarkers for iNPH, including acute-phase reactants, fragments of complement components, glycosylation variations, and reduced synaptic markers^10^. The study primarily emphasized sphingolipid patterns but did not provide extensive details on the quantity and specificities of the proteins measured^10^. Additionally, the study’s limited sample size and the notably older age of the iNPH cohort may affect the applicability of its results^10^.

The present study aims to fill the gaps in current research by conducting a large-scale, comprehensive proteomics analysis of CSF in patients with definite iNPH who have improved significantly after VP shunt surgery, Mild Cognitive Impairment (MCI) due to AD, and healthy controls (HC). Utilizing the Olink Explore 3072 panel, which screens for approximately 3000 proteins, this study seeks to identify distinct iNPH specific proteomic profiles and conceptualize a protein signature panel that could potentially differentiate iNPH from MCI due to AD and normal aging processes, while simultaneously providing clues into the pathophysiology of this poorly understood disease.

## 2. Materials and Methods

### 2.1. Study Cohorts

#### Participants

We evaluated individuals diagnosed with probable iNPH based on iNPH guidelines that presented with symptoms of gait disturbances, cognitive impairments, and/or urinary incontinence^15^. These patients were referred to the Johns Hopkins Center for Cerebral Fluid Disorders for evaluation and underwent a diagnostic CSF drainage trial in the form of Large Volume Lumbar Puncture (LVLP) with pre- and post CSF drainage gait assessments (Speed, balance, and endurance tests) to ascertain their suitability for shunt surgery. Following a comprehensive evaluation by a multidisciplinary team comprising neurology and neurosurgery specialists, eligible patients underwent ventriculoperitoneal (VP) shunt surgery (Table 1). The improvement after the shunt surgery was assessed using the Timed-Up-Go 3-meter (TUG 3m) test approximately 12 months post-surgery. The inclusion criteria for iNPH patients were: (1) Evan’s index greater than 0.3 (2) No secondary etiology such as trauma, stroke, hemorrhage etc. (3) Gait dysfunction with either cognitive impairment and/or urinary dysfunction (4) Improvement in the TUG 3m test results 12 months postoperatively (Median and mean interval of 12 and 14 months, respectively; see Table 2) of at least 30%, and (5) Availability of CSF samples from LVLP prior to the shunt surgery. Patients with concurrent neurological conditions, such as Parkinson’s disease or vascular dementia, were excluded from the study.

**Table 1.** Demographic table of normal pressure hydrocephalus (iNPH), healthy controls (HC), and mild cognitive impairment (MCI) participants detailing their cerebrospinal fluid (CSF) biomarkers such as neurofilament light concentrations (NfL), amyloid-β_1-42_ to amyloid-β_1-40_ ratio (Aβ_1-42_/ Aβ_1-40_), and phosphorylated tau 181 (p-tau181). MoCA: Montreal Cognitive Assessment.

|  | HC (N = 49) | iNPH (N = 28) | MCI (N = 38) | P-value |
| --- | --- | --- | --- | --- |
| **Sex** |  |  |  |  |
| F | 25 (51%) | 11 (39%) | 21 (55%) | 0.42 |
| M | 24 (49%) | 17 (61%) | 17 (45%) |  |
| **Age (Years)** |  |  |  |  |
| Mean ± SD | 69 ± 6.3 | 71 ± 5.0 | 75 ± 9.6 | <0.001 |
| **MoCA** |  |  |  |  |
| Mean (SD) | 26 ± 2.6 | 22 ± 5.9 | 21 ± 4.9 | 0.049 |
| **Evan's Index (EI)** |  |  |  |  |
| Mean ± SD | 0.28 ± 0.029 | 0.37 ± 0.024 | - | <0.001 |
| Missing | 13 (26.5%) | 0 (0%) | - |  |
| **Transependymal Flow Index** |  |  |  |  |
| Mean ± SD | 0 ± 0 | 0.82 ± 0.39 | - | <0.001 |
| Missing | 13 (26.5%) | 0 (0%) | - |  |
| **$A\beta_{1-42}/A\beta_{1-40}$** |  |  |  |  |
| Mean ± SD | 0.095 ± 0.013 | 0.12 ± 0.024 | 0.058 ± 0.020 | <0.001 |
| **P-tau181 (pg/mL)** |  |  |  |  |
| Mean ± SD | 34 ± 9.8 | 23 ± 9.0 | 86 ± 47 | <0.001 |
| **NfL (pg/mL)** |  |  |  |  |
| Mean ± SD | 2200 ± 830 | 1100 ± 460 | 3400 ± 2100 | <0.001 |

**Table 2.**
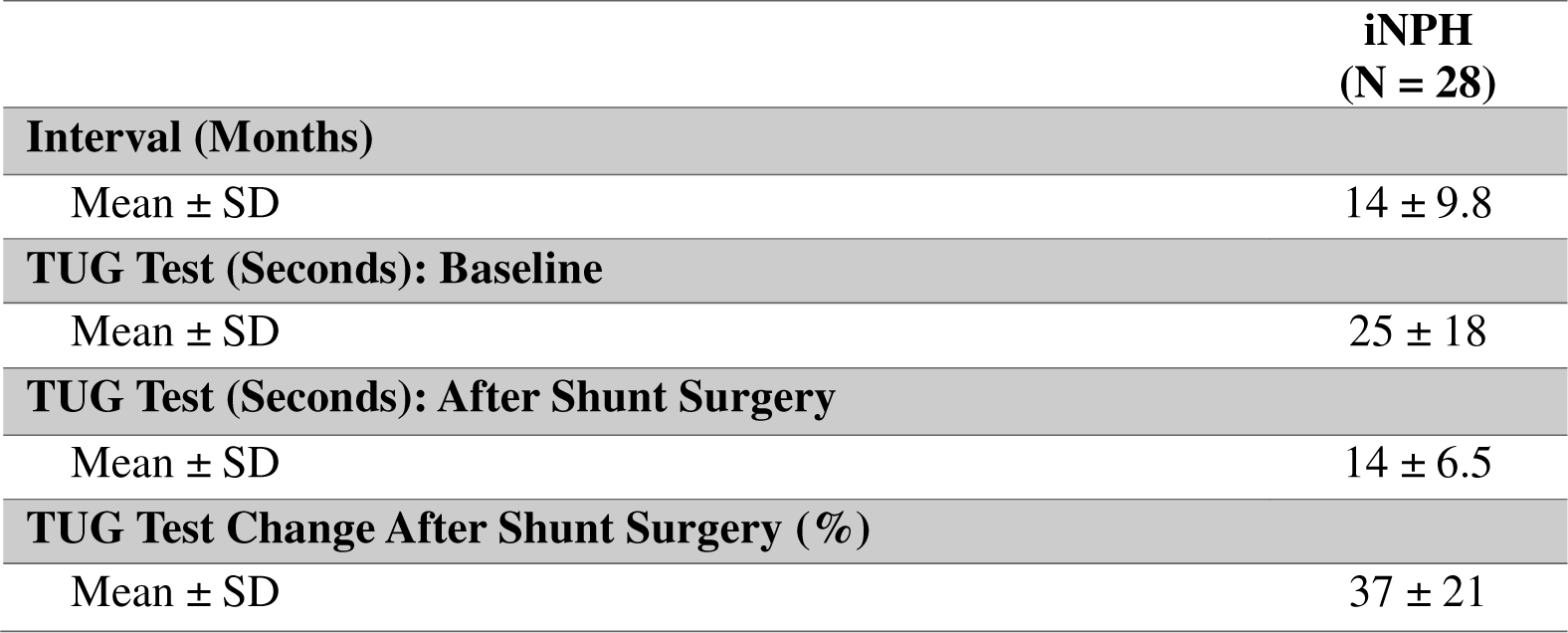
The interval between pre- and postoperative gait testing and Timed-Up-Go 3-meter (TUG 3m) gait test values of normal pressure hydrocephalus (iNPH) participants included in the study.

Additionally, our study included participants from both the Johns Hopkins Alzheimer’s Disease Research Center (ADRC) and Center for Cerebral Fluid Disorders within the Department of Neurology, Johns Hopkins University. These individuals were either cognitively intact or fulfilled the criteria for MCI. The clinical categorization of the MCI and the HC cohorts adhered to the guidelines set forth by the National Institute on Aging/Alzheimer’s Association workgroups^16^.

#### Consent statement

All subjects gave their informed consent for inclusion before they participated in the study. This study was conducted in accordance with the Declaration of Helsinki, and the protocol was approved by the institutional review board of Johns Hopkins University.

### 2.2. CSF Collection and CSF Biomarker Assays

CSF samples were collected from 28 iNPH, 49 HC, and 38 MCI participants. CSF of iNPH patients was collected during a LVLP procedure during which 40 ml of CSF was drained and they were not asked to fast before their procedure as is standard of care in the clinic. HC and MCI subjects underwent a lumbar puncture in the fasted state during which 20 ml of CSF was collected. All CSF samples were collected in a 50 mL polypropylene tube and transported on ice to the lab, where they underwent centrifugation at 2500× g for 15 min. Samples were then aliquoted in 0.5 mL aliquots in polypropylene cryovials and frozen at −80 C within 2 h of collection. CSF Aβ1–42, Aβ1–40, and p-tau181 were measured using the Lumipulse G1200 assay (Fujirebio, Malvern, PA, USA). The intra-assay coefficients of variation for this assay were 3.4% for Aβ1–42, 2.7% for Aβ1–40, and 1.8% for p-tau181. The ratio of CSF Aβ1–42/Aβ1–40 and p-tau181 were used in the current analyses. The participants with an Aβ1–42/Aβ1–40 ratio below 0.068 and p-tau181 levels above 50.6 pg/mL were identified as MCI participants with positive Alzheimer’s disease (AD) biomarkers. All MCI participants selected for this study were required to be AD-biomarker positive based on their CSF values, whereas all HC participants were required to be AD-biomarker negative based on the cut-offs established by Greenberg et al^17^.

### 2.3. Magnetic Resonance Imaging (MRI)-Based Quantification of Intracranial Volume (ICV), Transependymal Flow (TEF), and Evans’ Index (EI) Calculations

All iNPH patients underwent standard of care Magnetic Resonance Imaging (MRI) that included 3D-MPRAGE T1, T2, Fluid attenuated inversion recovery (FLAIR), and Diffusion-weighted imaging (DWI) sequences before referral to our center except for 2 participants that had undergone imaging studies at another medical institute. The healthy controls included in this study were recruited from a separate ADRC study and had undergone only T2 and FLAIR MR imaging.

In the axial images, aligned with the anterior commissure-posterior commissure plane and using T2-weighted sequences, we assessed the maximal width of the frontal horns. The EI was determined for iNPH patients and HC by dividing the maximal lateral width of the frontal horns of the lateral ventricles by the maximal internal diameter of the skull at the corresponding axial level. The presence or absence of transependymal flow (TEF) was reported by a neuroradiologist based on T2-weighted sequences for iNPH patients and HC.

In this study, brain mask and volumetric data were extracted for each iNPH patient employing FreeSurfer’s recon-all function. Given FreeSurfer’s requirement for a slice thickness of less than 1 mm, 23 out of the 26 iNPH patients who had available MPRAGE T1 scans underwent analysis with their original scans. SynthSR, an artificial intelligence (AI) super-resolution tool, was then employed to enhance the resolution of the 3 remaining scans, enabling compatibility with FreeSurfer’s analysis^18^. Prior to analysis, all scans underwent preprocessing and harmonization procedures to standardize contrast, resolution, and orientation, thus ensuring consistent results^19^. Subsequently, outputs from FreeSurfer were visualized and checked for any inconsistencies to ensure the integrity and quality of the results. Finally, total intracranial volume for each patient was estimated using FreeSurfer’s native functions (Figure s1)^20^.

### 2.4. Protein Measures Using Olink Proximity Extension Assay (PEA), Quality Control (QC) and Data Pre-Processing

The Olink Explore panel, a highly sensitive and specific technique, measures the expression of 3072 proteins in 10 µL CSF (Olink, Uppsala, Sweden). CSF protein measurements were conducted using PEA technology, following the manufacturer’s protocol^21^.

The CSF samples were measured on 2 Olink Explore panels with 17 bridging samples between the two plates. We then bridged the data from two plates using bridging normalization and verified the process using principal component analysis (PCA) plots before and after the normalization step for each of the 8 Olink Explore 3072 panels.

Pre-processing handling of data included plate-based normalization and QC checks based on appropriate Olink protocols^22^. Outlier deletion was performed subsequently by detecting and deleting datapoints that were above or below 5 SD of mean normalized protein expression (NPX) of each assay (protein). All datapoints with QC or assay warning were also deleted. PCA plots per panel were used to detect and delete outlier samples^22^.

### 2.5 Data Analysis and Statistical Methods

#### Differential expression analysis

Statistical analyses were performed with R software (R-project.org) using the publicly available OlinkAnalyze package^23,24^. Groups were compared using Welch’s two-sample, two-sided t-test analysis and Benjamini–Hochberg (BH) post hoc analysis. Differentially expressed proteins were defined as assays with false discovery rates (FDR) below 0.05 (FDR-adjusted p-value < 0.05).

#### Group comparisons

Additionally, after verifying normality (Using Shapiro-Wilk test) and variance comparability in each assay, we applied ANOVA and Kruskal-Wallis (KW) tests, as appropriate, for comparative analysis among iNPH, HC, and MCI patient groups. These analyses were performed using RStudio, considering a p-value of ≤0.05 as statistically significant. Posthoc ANOVA and non-parametric analysis was performed using Tukey and BH p-value adjustment, respectively, per assay (by OlinkID) at confidence level 0.95.

#### Orthogonal Partial Least Squares Regression Discriminant Analysis (OPLS-DA)

In our study, comprehensive multivariate analyses were executed utilizing SIMCA software (version 18, Sartorius). A preliminary PCA incorporated all proteins and participants, primarily to inspect the dataset’s structure and to validate the discernibility of sample groups based on their respective conditions^25^. This exploratory step led to the identification of two outliers, as determined by metrics such as distance to model X and Hotelling’s T2 criterion^26^. However, they were retained in subsequent analyses, given the absence of substantial deviations in individual protein levels. Subsequently, we engaged in Orthogonal Partial Least Squares Regression Discriminant Analysis (OPLS-DA) with a missingness threshold of 90%^26^. This approach is particularly suitable for large-scale omics datasets, as it models variances in predictor variables (proteins) against the dependent variable (group classifications)^26^. During this analysis, data preprocessing included mean centering and unit variance scaling, ensuring that both low- and high-abundance proteins exerted comparable influence on the model.

The validity of the OPLS-DA model was ascertained through a cross-validated analysis of variance (CV-ANOVA)^26^. The model’s effectiveness was evaluated based on R^2^ (goodness of fit) and Q^2^ (goodness of prediction) metrics. To determine the proteins with the highest discriminative power, we computed Variable Influence in the Projection (VIPpred) statistics.

These metrics elucidate the relative contribution of each protein to the model’s predictive accuracy. Additionally, we normalized the loadings of each protein within the model as correlation coefficients (p(corr)[1]), thus standardizing their range from –1.0 to 1.0. Proteins demonstrating a VIPpred value of ≥2.0 combined with a p(corr)[1] of ≥0.5 or ≤ -0.5 were marked as possessing the highest discriminative potential.

### 2.6. Protein Pathway Analysis

#### Overrepresentation analysis (ORA)

After performing differential abundance analysis on our proteomics dataset, we conducted overrepresentation analysis (ORA) to identify differentially expressed pathways. This bioinformatics analysis helped us identify pathways that were enriched with significant changes in protein abundance. The goal of this analysis was to gain a better understanding of the functional implications of the differentially expressed proteins and to identify potential biological processes involved in the observed phenotypes.

We performed overrepresentation analysis on differentially expressed proteins (with differential abundance of FDR < 0.05) using Gene Ontology (GO) biological process and the Wikipathway database^27–29^. ORA was performed using the WEB-based GEne SeT AnaLysis Toolkit (WebGestalt), which is a functional enrichment analysis web tool implementing several biological functional category databases such as KEGG, Reactome, WikiPathway, and PANTHER. The default settings of WebGestalt were used with the FDR cut-off of 0.05^28,29^.

#### Gene Set Enrichment Analysis (GSEA)

Gene Set Enrichment Analysis (GSEA) is a computational method that determines whether the expression of a set of genes is significantly different between two phenotypes^30^. GSEA was performed using GSEA 4.3.2 software^31,32^. The GSEA calculates the signal-to-noise ratio for all proteins and orders gene sets by normalized enrichment scores (NES). We performed the GSEA with the default settings of the software, which included 1000 permutations, phenotype permutation type, exclusion of gene sets larger than 500 and smaller than 15 and using weighted enrichment statistics. FDR cut-off of 0.25 was adopted in this analysis as recommended by GSEA developers to avoid overlooking potentially significant pathways in the context of relatively small number of gene sets being analyzed^31,32^.

In addition to ORA, we also performed GSEA in our proteomics study of CSF from iNPH and MCI patients. We chose to use both methods because they provide complementary information and can help to validate each other’s results^30^.

ORA allowed us to identify pathways that are overrepresented in our list of differentially expressed proteins, whereas GSEA enabled us to assess the enrichment of pre-defined gene sets in the list of differentially expressed proteins ranked by normalized enrichment scores, which can provide a more comprehensive understanding of the biological processes affected by the disease^30^.

Using both methods also helped to overcome the limitations of each method, such as the dependence on pre-defined gene sets in ORA and the need for a large reference gene set in GSEA. Overall, by using both ORA and GSEA, we were able to identify novel and previously known pathways that are dysregulated in iNPH and gain a better understanding of the underlying mechanisms contributing to the pathogenesis of this disease.

## 3. Results

### 3.1. Proximity Extension Analysis (PEA) Experiment Results

We used PEA technology with the Olink Explore panel to determine the differential expression of approximately 3072 proteins in 28 iNPH patients that improved significantly after shunting, 38 AD-biomarker positive MCI patients and 49 HC (Table 1). The intra-assay and inter-assay coefficient of variation (CV) of the panels did not exceed 15%. The proteins that did not pass the Olink batch release quality control criteria were excluded from the study (Refer to supplementary material).

### 3.2. Comparative Proteomic Analysis of CSF in iNPH, HC, and MCI

In our study, proteomic analysis of CSF was conducted, resulting in the assay of 2,888 proteins. Comparative analysis revealed 583 proteins exhibiting differential expression in iNPH patients, as determined by FDR-adjusted p-values <0.05, detailed in Tables s1 and s2. Notably, of these 583 proteins, 416 were identified as downregulated, as depicted in Figure 1c.

**Figure 1.**
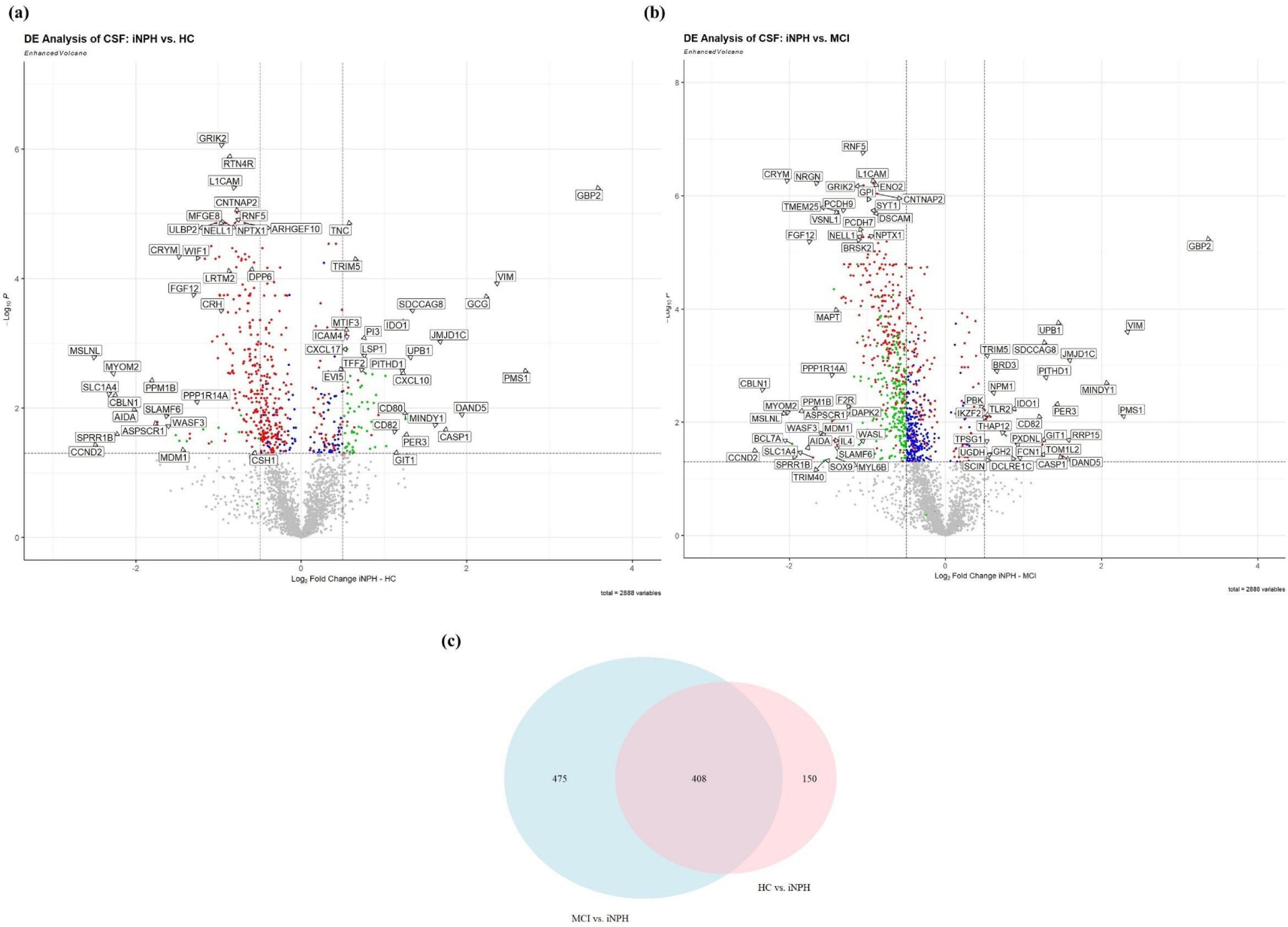
Comparative proteomics profile between normal pressure hydrocephalus (iNPH) and healthy control (HC), and mild cognitive impairment (MCI) participants. (a) Volcano plot of differentially expressed proteins (Adjusted p-value <0.05) in iNPH compared to HC and (b) iNPH compared to MCI. The overlapping significant proteins between the two comparisons are illustrated as red in both volcano plots, the green dots represent significant differentially expressed proteins that had log_2_ Fold Change of over 0.5 and the blue dots are significant differentially expressed proteins with log_2_ Fold Change below 0.5. (c) Venn diagram of the number of significant differentially expressed proteins in each comparison.

Comparative analysis between iNPH and MCI subjects identified 930 proteins with differential expression, predominantly downregulated in iNPH (823 proteins), as shown in Figures 1a and 1b. Figure 1c illustrates the significant overlap in differentially expressed proteins across both comparative analyses.

ANOVA and Kruskal-Wallis tests identified 1,174 proteins with significant variance across HC, MCI, and iNPH groups (Refer to Tables s3 and s4, and Figures s2 and s3).

### 3.3. Multivariate OPLS-DA Analysis Outcomes

The dataset was subjected to OPLS-DA, yielding a robust model (F(8.25) = 18.45, p = 8.14e-32) with two principal and two orthogonal components, demonstrating high model fit (R^2^Y = 87%) and moderate predictive capacity (Q^2^ = 55%). The model’s score plot (Figure 2a) delineates clear group separations. Figure 2b highlights proteins significantly differentiating iNPH patients from HC and MCIs. Of 2888 proteins analyzed, 13 were identified as significant discriminators among the three groups, with the threshold of variable importance in projection (VIPpred) ≥2.0 and p(corr) ≥0.5 or ≤-0.05. These discriminators are detailed in Figure 2b, while comprehensive protein data is presented in Table s7. We then created three separate OPLS-DA models consisting of the traditional AD biomarkers (Aβ_1-42_/Aβ_1-40_, p-tau181, and NfL; Figure 2c), the 13 key discriminator protein panel (Figure 2d), and the 13-protein panel plus the AD biomarker set together (Figure 2e). The comparison of these three models demonstrated that the discriminative power of a model consisting of both traditional AD markers and the 13 protein panel (Figure 2e) has an especially enhanced goodness of fit and predictive power (R^2^Y = 66% , Q^2^= 59%) compared to either one of these panels alone (R^2^Y = 47% , Q^2^= 38% and R^2^Y = 37% , Q^2^= 32% for 13-protein model and AD biomarker model, respectively).

**Figure 2.**
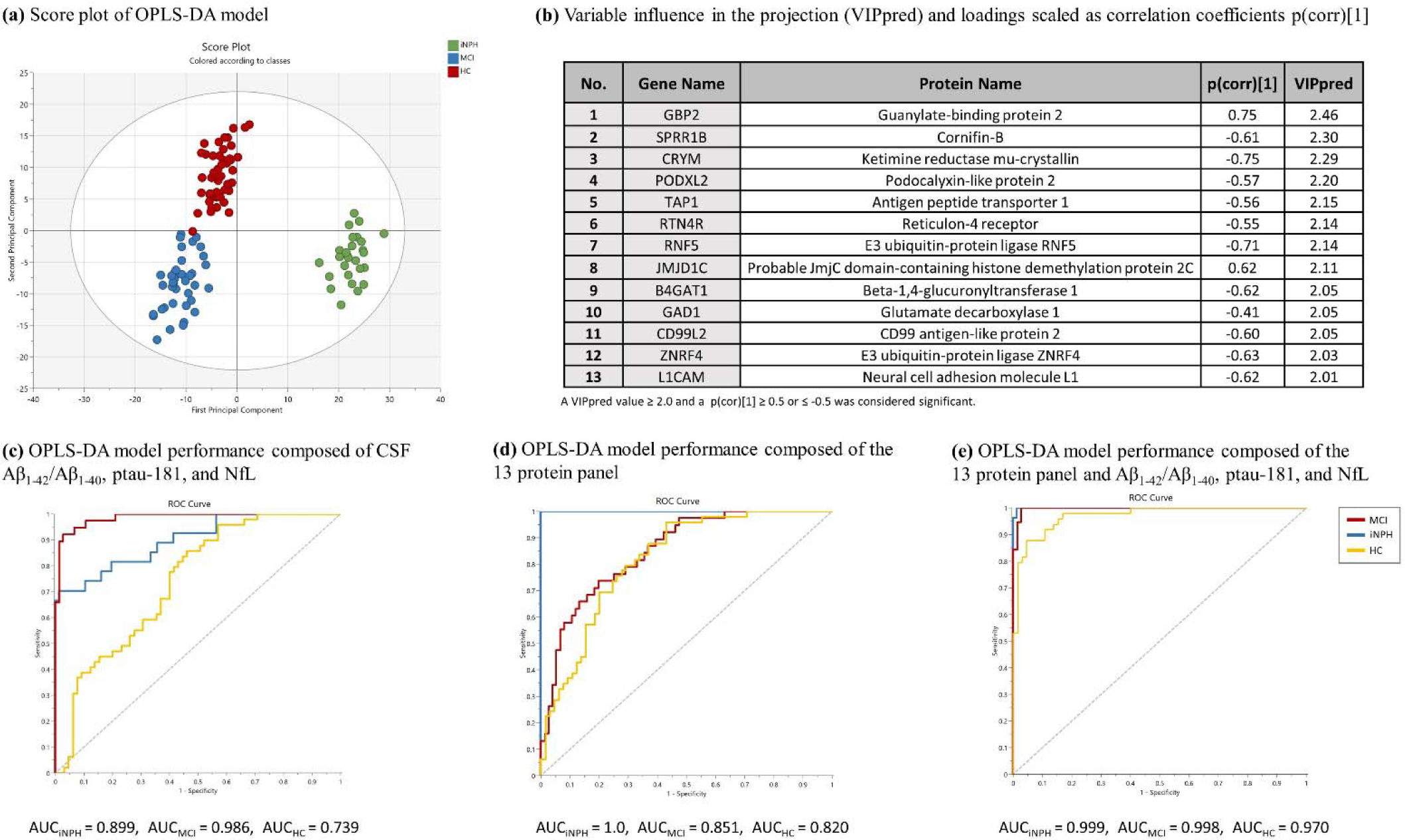
Score plot of the orthogonal partial least squares regressions discriminant analysis (OPLS-DA) model delineating the separation of normal pressure hydrocephalus (iNPH), mild cognitive impairment (MCI), and healthy control (HC) groups based on protein expressions (NPX) in cerebrospinal fluid (CSF), (b) Visualization of 13 proteins with highest distinctive power selected based on OPLS-DA model that significantly differentiate iNPH from HC and MCI with their variable influence in the projection (VIPpred) and loadings scaled as correlation coefficient (p(corr)[1]). (c) Receiver operating characteristic (ROC) curve of an OPLS-DA model that uses cerebrospinal fluid (CSF) levels of Aβ_1-42_/ Aβ_1-40_, p-tau181, and neurofilament light (NfL) to discriminate between iNPH, MCI, and HC. (d) ROC curve of an OPLS-DA model that uses CSF levels of the 13 most distinctive proteins identified in previous OPLS-DA analysis to discriminate between three conditions, and (e) ROC curve of an OPLS-DA model that uses CSF levels of the 13 most distinctive proteins and biomarkers detailed in Figure 2(c) to discriminate between iNPH, MCI, and HC.

### 3.4. Overrepresentation Analysis (ORA) Outcomes

ORA, utilizing the GO database, revealed significant enrichment in biological processes such as the “synaptic vesicle cycle”, “neuron migration”, “central nervous system neuron differentiation”, and “response to chemokine”, as illustrated in Figure s4. Additional processes like “forebrain development” and “cell-cell adhesion” were also significantly enriched. Notably, the enrichment patterns in iNPH vs. HC comparisons were largely mirrored, except for the GO terms “cognition” and “response to chemokine,” which were not enriched in the latter comparison (Refer to Figure s4).

### 3.5. Gene Set Enrichment Analysis (GSEA) Outcomes

GSEA indicated an upregulation of “chemokine receptor binding” and “lymphocyte chemotaxis”, and a downregulation of synaptic membrane-related proteins and cellular components in addition to cell-cell adhesion in iNPH (Figure 3 and Table s5). “Cell-cell junction assembly” and “tight junction” are among the significantly downregulated GO terms in iNPH.

**Figure 3.**
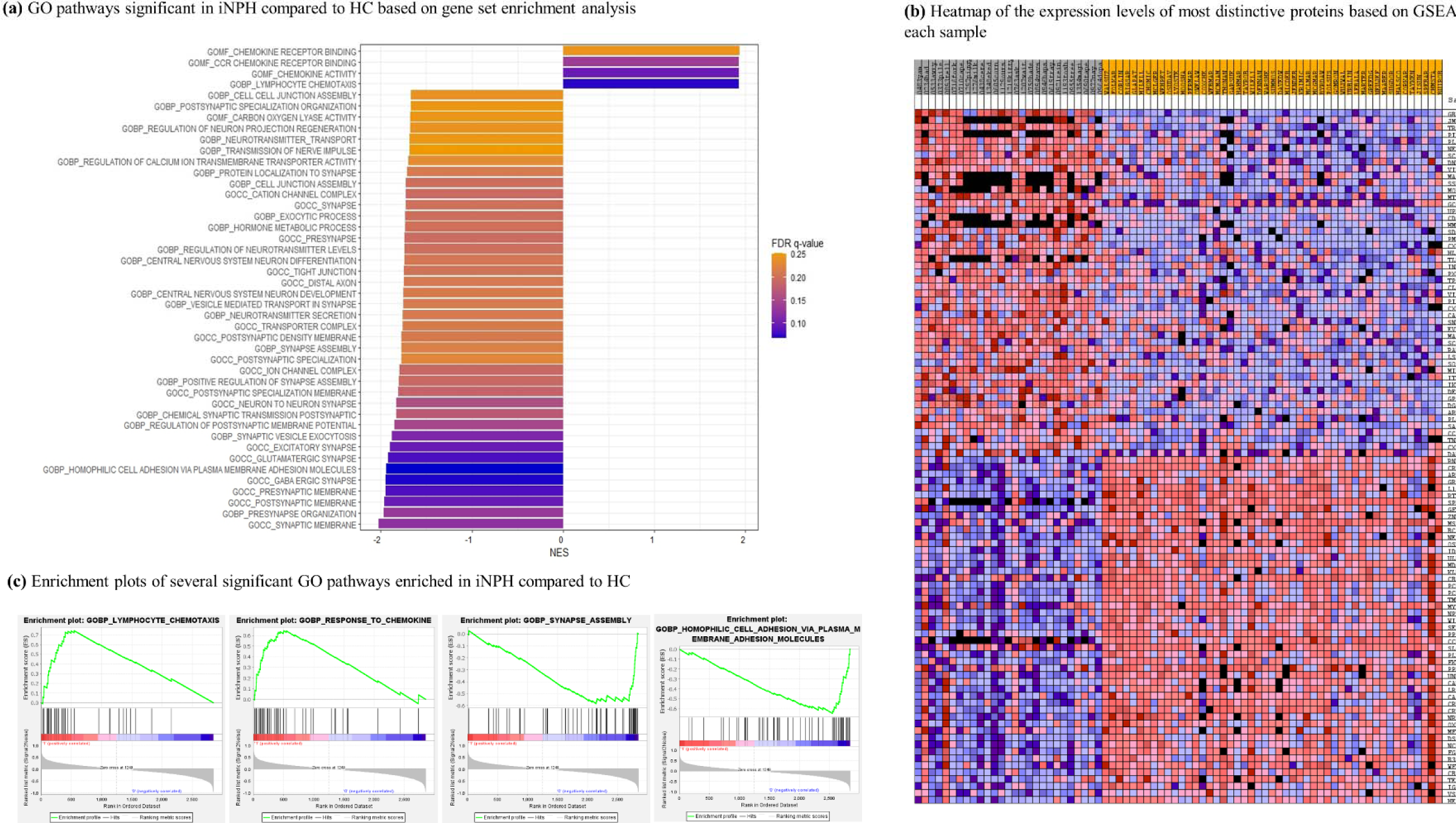
The results of gene set enrichment analysis (GSEA) demonstrating the (a) significantly enriched gene ontological (GO) terms in normal pressure hydrocephalus (iNPH) compared to healthy control (HC) group and their normalized enrichment scores (NES), (b) a heatmap of the most distinctive proteins based on GSEA and their expression levels in each sample of iNPH (grey samples) and HC (orange samples) is demonstrated. (c) Enrichment plots of two most significantly downregulated and two most significantly upregulated GO biological processes in iNPH is demonstrated.

Similar synaptic membrane-related set of proteins were found to be significantly downregulated in iNPH when compared to MCI, as summarized in Table s6. However, no upregulation of inflammatory and chemokine-related pathways was found to be significant in this comparison.

## 4. Discussion

### 4.1. Overview of Proteomic Alterations in iNPH

Employing PEA with the Olink Explore 3072 panel, our study embarked on a comprehensive examination of approximately 3000 proteins across 28 shunt-responsive iNPH patients with post-shunt improvement, 38 AD-biomarker positive MCI patients, and 49 HC. Out of 2,888 proteins assayed, 583 exhibited differential expression in iNPH, with a significant downregulation of 416 proteins (Figure 1). This downregulation is notably profound when juxtaposed with MCI subjects, revealing 823 proteins predominantly downregulated in iNPH. This extensive alteration suggests a unique CSF proteomics profile of iNPH.

The unaffected concentrations of established biomarkers for conditions often misdiagnosed as iNPH, such as MCI due to AD (indicated by CSF levels of Aβ_1-42_/Aβ_1-40_, p-tau181, secernin-1 [SCRN1], and microtubule-associated protein tau [MAPT]) and Parkinsonian syndrome (marked by α-synuclein [SNCA] and DOPA decarboxylase [DDC]), underscore the phenotypic consistency of the iNPH participants in our study^33,34^.

### 4.2. Potential Mechanisms Underlying Extensive Protein Downregulation

The skewed volcano plots, indicating a significant number of downregulated proteins in iNPH CSF, prompt the exploration of underlying mechanisms. The dilution effect due to ventriculomegaly is a plausible explanation, considering the increased ventricular space in iNPH^35^. However, our comparative analysis of total CSF protein concentrations and constitutively secreted protein concentrations challenges this hypothesis. Transthyretin (TTR), which is one of the most abundant proteins in CSF and is exclusively expressed by choroid plexus into CSF, and autotaxin (ENPP2), also constitutively secreted into the CSF were not significantly different between iNPH and HC (Figure s3 diagrams 320 and 892, respectively)^36–38^. Moreover, total CSF protein concentrations measured as part of clinical laboratory testing on LVLP CSF samples of our iNPH participants did not support the dilution effect hypothesis (Mean total CSF protein = 51.2 ± 22.5 mg/dL with normal range of 15-45 mg/dL). Thus, we found no basis for the hypothesis that this might be due to dilution which is also in line with previous studies^39^.

The potential dysfunction of the glymphatic system in iNPH offers an intriguing perspective on the observed protein downregulation in our study. This system, crucial for eliminating peptides and neurotoxic waste from the perineural extracellular efflux and cerebral interstitial fluid (ISF) to the CSF, is implicated in various neurodegenerative diseases^40–42^. Impairment in this system could lead to inadequate drainage of neuronal markers from ISF, contributing to their decreased presence in CSF and reflecting broader neurodegenerative changes within the central nervous system (CNS) (Figure 4). This hypothesis is supported by imaging studies indicating reduced glymphatic activity in iNPH and aligns with observed synaptic and neuronal marker downregulation^6,40,43,44^.

**Figure 4.**
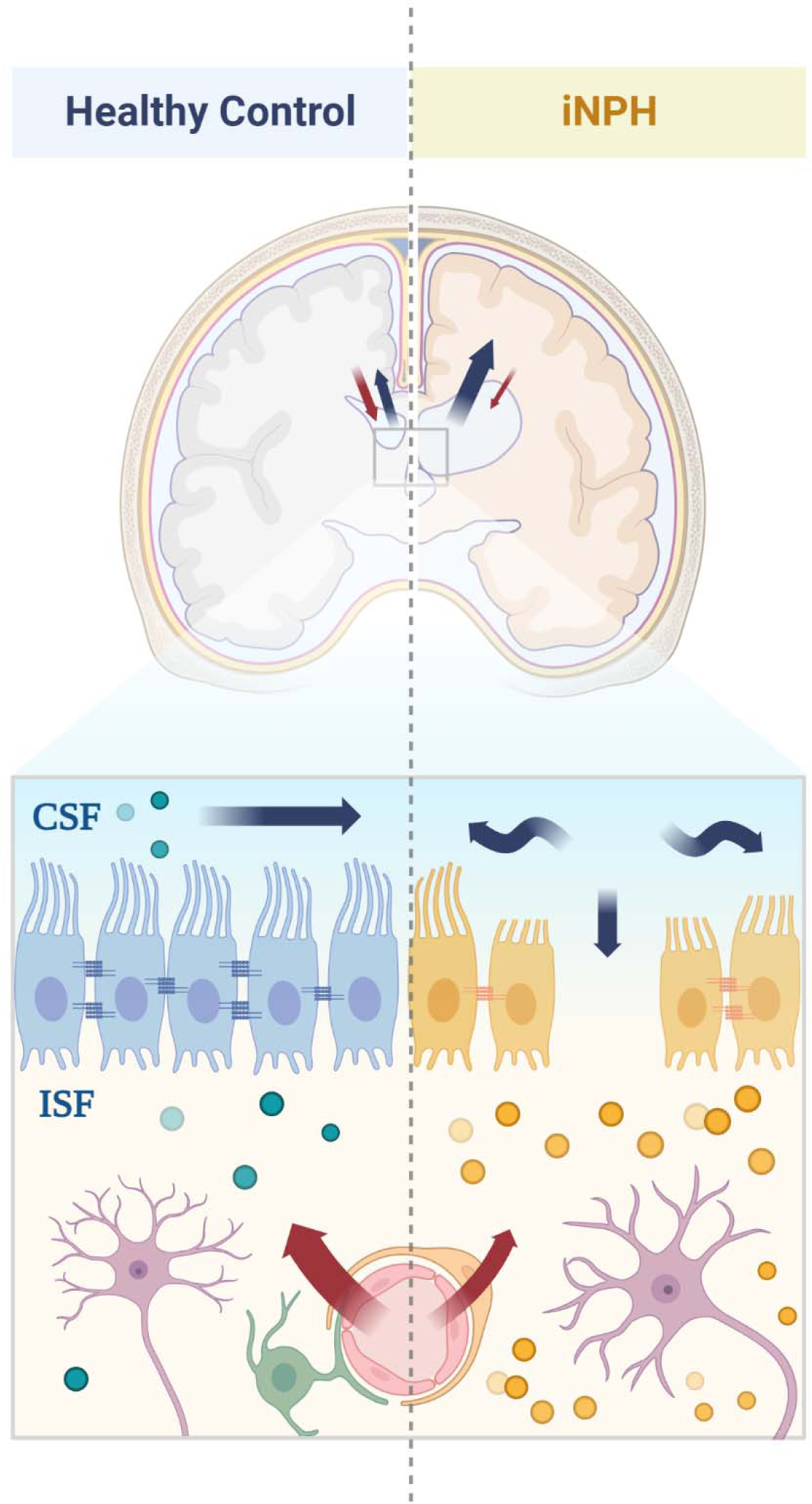
Depiction of the hypothesized glymphatic system dysfunction and transependymal flow disruption affecting the clearance of Interstitial fluid (ISF) into cerebrospinal fluid (CSF) in normal pressure hydrocephalus (iNPH), illustrating its potential role in the observed CSF proteomic changes.

Moreover, the TEF index, as shown in demographic Table 1, indicates disrupted ISF to CSF flow in iNPH patients, with a TEF index of 1, compared to 0 in healthy controls. This index reflects the abnormal movement of CSF across the ependymal layer from the ventricular compartment into the interstitial spaces of the brain parenchyma, suggesting compromised fluid dynamics and ependymal integrity in iNPH. Such disruptions could contribute to the accumulation of metabolic waste in the brain’s interstitial space, impacting neuronal health and potentially leading to the observed downregulation of synaptic and neuronal markers.

### 4.3. The Role of Synaptic and Neuronal Markers

The downregulated synaptic and neuronal markers identified, including neuronal pentraxins and their receptor (NPTX1, NPTX2, NPTXR), neurogranin (NRGN), synaptotagmin-1 (SYT1), and neuroplastin (NPTN), Dyslexia-associated protein (KIAA0319), and synaptosomal-associated protein 25 (SNAP25) among others, are integral to synaptic function and neuronal communication. The low concentrations of these proteins in CSF of iNPH, if not affected by the disrupted ISF-CSF flow and glymphatic dysfunction, could indicate a disruption in synaptic integrity and neuronal signaling, contributing to the clinical manifestations of the condition. Similar observations in previous studies further substantiates the potential role of glymphatic dysfunction and/or synaptic dysfunction in iNPH pathogenesis^10,45^.

### 4.4. Cell-Cell Adhesion and Neuroepithelial Integrity

Our study emphasizes a significant downregulation in cell-cell adhesion processes in iNPH. The dysregulation of neuron-specific adhesion molecules such as CNTNAP2, CELSR2, and various other cadherins (e.g., PCDH7, PCDH-9, PCDH-17, CDH2, CDH6, and CDH15) which are critical for axonogenesis and synaptic maturation, suggests a disruption in neuroepithelial integrity and possibly neuronal connectivity^46,47^. This is further supported by the literature highlighting the role of various tight and adherens junctions, especially N-cadherin (CDH2) in maintaining the integrity of the neuroepithelium and glial progenitor cells and hydrocephalus development^47,48^. The disruption of ependymal planar polarity and CSF circulation, as evidenced by the downregulated polycystin 1 (PKD1) and CELSR2, is also consistent with established hypotheses of hydrocephalus development^49–52^.

Moreover, vimentin’s upregulation in iNPH CSF, as the second most upregulated protein, is noteworthy (Figure 1a). As an intermediate filament prevalent in ependymal cells, its increased expression is a marker of ongoing ependymal injury and a potential role in the pathogenesis of iNPH. The consistency of our findings with previous literature, which links ependymal injury and congenital hydrocephalus to Vimentin upregulation, indicates a common pathway that might be crucial for understanding iNPH development^53^.

Together, these findings suggest that iNPH may arise from a cascade of intertwined cellular events, encompassing altered cell adhesion, ependymal cell pathology and denudation, and impaired ISF-CSF flow and glymphatic clearance, each contributing uniquely to the disease’s progression and the diverse symptomatology observed in patients.

### 4.5. Inflammatory Pathways and Chemokine Upregulation

The differential expression and pathway analyses from our study point to an upregulation of inflammatory pathways in iNPH (e.g., CXCL1, CXCL9, CXCL10, CCL23), particularly those related to lymphocytic and monocytic chemotactic pathways and adaptive immunity (e.g., CCL2, CXCL11, CCL4, CCL8, CCL21). The upregulation of chemokines, especially Monocyte chemoattractant protein-1 (CCL2), aligns with previous studies, suggesting an inflammatory component in iNPH pathogenesis ^1,9,54–56^. The role of this observed inflammation, whether as a cause or consequence of choroid plexus hypersecretion or BBB dysfunction, remains to be elucidated ^9^.

### 4.6. Possibility of Shared Molecular Pathways in Congenital Hydrocephalus and iNPH

Moreover, our study identifies the downregulation of specific genes, such as L1CAM, PCDH9, ISLR2, ADAMTSL2, and B4GAT1, which have been previously well characterized as playing an important role in congenital hydrocephalus and aqueductal stenosis ^57–64^. The downregulation of L1CAM, a gene extensively studied and phenotyped for its role in X-linked congenital hydrocephalus, is particularly noteworthy^57,61^. In addition, previous studies have explored genetic markers of adult-onset iNPH and have discovered mutations of CWH43 gene (not measured in Explore 3072 panel) in iNPH patients that increases L1CAM proteolysis^65,66^. This finding, alongside the uniformly downregulated expression of PCDH9, ISLR2, ADAMTSL2, and B4GAT1 in our iNPH patients, reinforces the potential parallel mechanisms in the pathophysiology of congenital hydrocephalus and iNPH and raises the possibility that iNPH may be a forme-fruste of congenital hydrocephalus that manifests in adulthood at least in a subset of patients. This is supported by prior findings that patients with iNPH have a larger head circumference than normal controls of the same height^67^.

These proteins are integral to various cellular processes, including but not limited to neuronal-cell adhesion, neurite migration, glycosylation, TGF-β bioavailability, and axonogenesis, which are crucial in maintaining normal brain development and fluid dynamics^46,57,63^. Their downregulation in our study not only aligns with the observed disruptions in cell junctions, axonogenesis, and ependymal integrity, as discussed earlier, but also suggests a broader connection to congenital hydrocephalus.

Prior research has shown that patients with iNPH exhibit significantly increased ICV in comparison to healthy individuals, suggesting that iNPH originates in early infancy, consequent to diminished CSF uptake before cranial sutures close^68^. Consequently, we assessed the ICV of patients with iNPH in our study and aligned these findings with previously established normative data^69^. Our results revealed a considerable enlargement in ICV among the iNPH patients (Mean ICV Percentile = 92 ± 9.6, see Table s8), pointing to the early onset of the disease pathology. This connection further underscores a possible shared genetic and molecular foundation between congenital hydrocephalus and iNPH, providing further insight into the complex network of factors contributing to the development and progression of these conditions.

### 4.7. Multivariate OPLS-DA Analysis and Key Discriminator Proteins

Our robust multivariate OPLS-DA analysis, demonstrating high model fit and moderate predictive capacity in differentiating between iNPH, MCI and HC, underscores the potential of identified proteins as CSF biomarkers for iNPH. Thirteen proteins were distinguished as significant discriminators among the groups, identified with high variable importance in projection (VIPpred) and correlation coefficient (p(corr)[1]), offer a panel of proteins that can be utilized in designing possible targets for a diagnostic panel pending external validation in a larger dataset.

### 4.8. Limitations and Novelty of the Study

Our conclusions are primarily limited by the small number of iNPH patients included, which may affect the robustness of our analytical findings. Furthermore, the 13-protein panel identified for potential iNPH diagnosis can only be deemed hypothesis generating for now and necessitates further validation in more extensive and diverse patient cohorts to establish its diagnostic accuracy. The age difference between the MCI and iNPH groups presents another challenge, potentially leading to age-related differences in proteomic profiles. Additionally, our analysis, while extensive, is confined to the proteins within the Olink Explore 3072 panel and does not encompass the entire spectrum of CSF proteins. We also were not able to assess alterations in many of the recently identified genes related to iNPH (e.g., CWH43, SFMBT1) in our patients given they were not measured in our proteomics panel^70^.

Despite these limitations, this study is pioneering in its comprehensive proteomic analysis of CSF in iNPH, using cutting-edge PEA technology. It stands as the first to undertake such a detailed comparison between iNPH, HC, and MCI, which is often considered in differential diagnosis. Moreover, our study offers unique insights into iNPH pathophysiology and potential connections with congenital hydrocephalus.

## 5. Conclusion

In conclusion, our comprehensive CSF proteomics study of iNPH compared to MCI and HC illuminated several distinctive patterns of proteomics changes in iNPH including the downregulation of synaptic markers and downregulation of proteins vital for cell-cell adhesion (e.g., L1CAM, DSCAM, and various cadherins) and ependymal planar polarity coupled with vimentin upregulation suggested underlying ependymal layer denudation and dysfunction. Furthermore, the potential parallels drawn with congenital hydrocephalus (e.g., L1CAM) provide a new perspective on possible shared molecular pathways.

The discovery of a 13-protein panel of the most distinctive proteins offers potential for future diagnostic applications, pending further validation. Hence, our findings pave the way for new avenues in iNPH research, opening possibilities for improved diagnosis and a deeper comprehension of this multifaceted neurological disorder.

## 6. Acknowledgement

This work was supported by the National Institutes of Aging (Grant Number: U19–AG033655 and P30–AG066507).

## 7. Informed Consent Statement

Informed consent was obtained from all subjects involved in the study.

