## Supplementary Figures for "Molecular Signatures of Normal Pressure Hydrocephalus: A Large-scale Proteomic Analysis of Cerebrospinal Fluid"

### Supplementary Methods

The proteins that did not pass the Olink batch release quality control criteria were excluded from the study included: KNG1, BMP6, EPHX2, PGLYRP1, EDEM2, CALY, ARL13B, ARNTL, BCL2L11, BID, LTA, GZMB and MGLL, EP300, FGF3, FUOM, KNG1, ADIPOQ, CDHR1, CLSTN1, PSG1, CGA, EFNA1, HTR1B, KCNH2, STXBP1 and YAP1, FLI1, MPI, EBI3_IL27, ANGPTL7, CPLX2, TAGLN3, GABARAPL1, NFKB2, CTAG1A_CTAG1B, OGT, MTHFSD, IFIT1, TNPO1, MAGEA3, SH3GL3, and RAPGEF2.

**Figure s1. Workflow of Intracranial volume (ICV) estimation using FreeSurfer for idiopathic Normal Pressure Hydrocephalus (iNPH) patients.** Patients that had scans with a slice thickness of less than 1 mm (23 out of the 26 iNPH patients) underwent analysis with their original scans (b) . SynthSR, an artificial intelligence (AI) super-resolution tool, was employed to enhance the resolution of the 3 remaining scans, enabling compatibility with FreeSurfer's analysis (a).

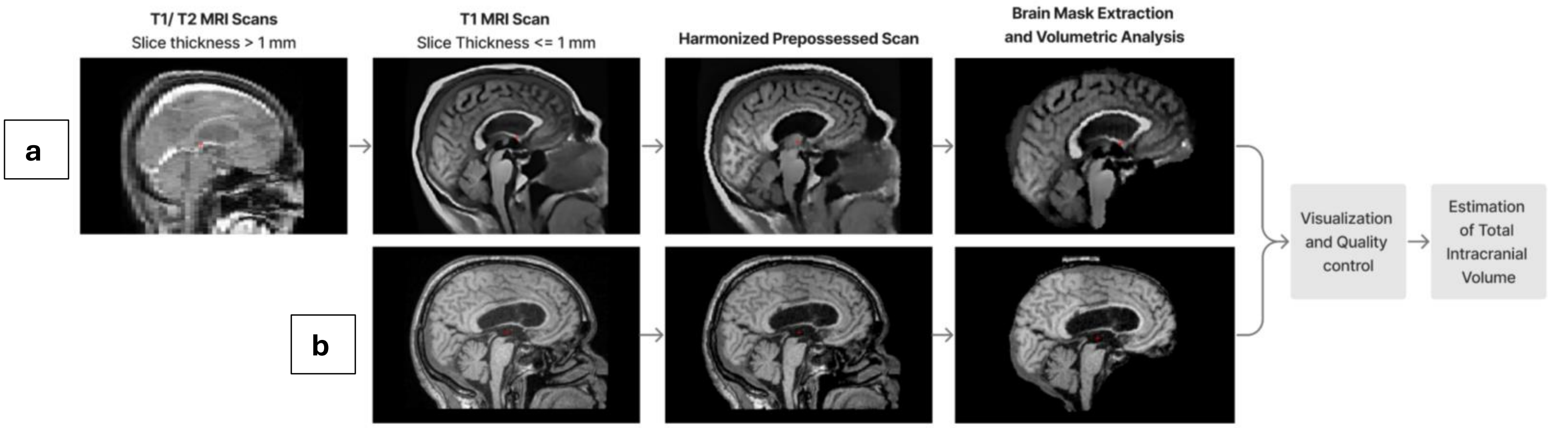


**Figure s2. Boxplots of Analysis of Variance (ANOVA) of Normalized Protein Expression (NPX) Across Different Conditions.** This visual representation aids in the comparative analysis of protein expression levels (NPX) between conditions such as idiopathic Normal Pressure Hydrocephalus (iNPH), Mild Cognitive Impairment (MCI), and Healthy Control (HC). Each boxplot corresponds to a unique protein demonstrated by their gene symbols, identified through Analysis of Variance (ANOVA) testing across conditions. The significance levels, adjusted using the Benjamini-Hochberg (BH) method, are indicated by asterisks on the plots: $p < 0.0001$ = ‘****’, $p < 0.001$= ‘***’, $p < 0.01$ = ‘**’, $p < 0.05$ = ‘*’, and ‘ns’ for non-significant differences.

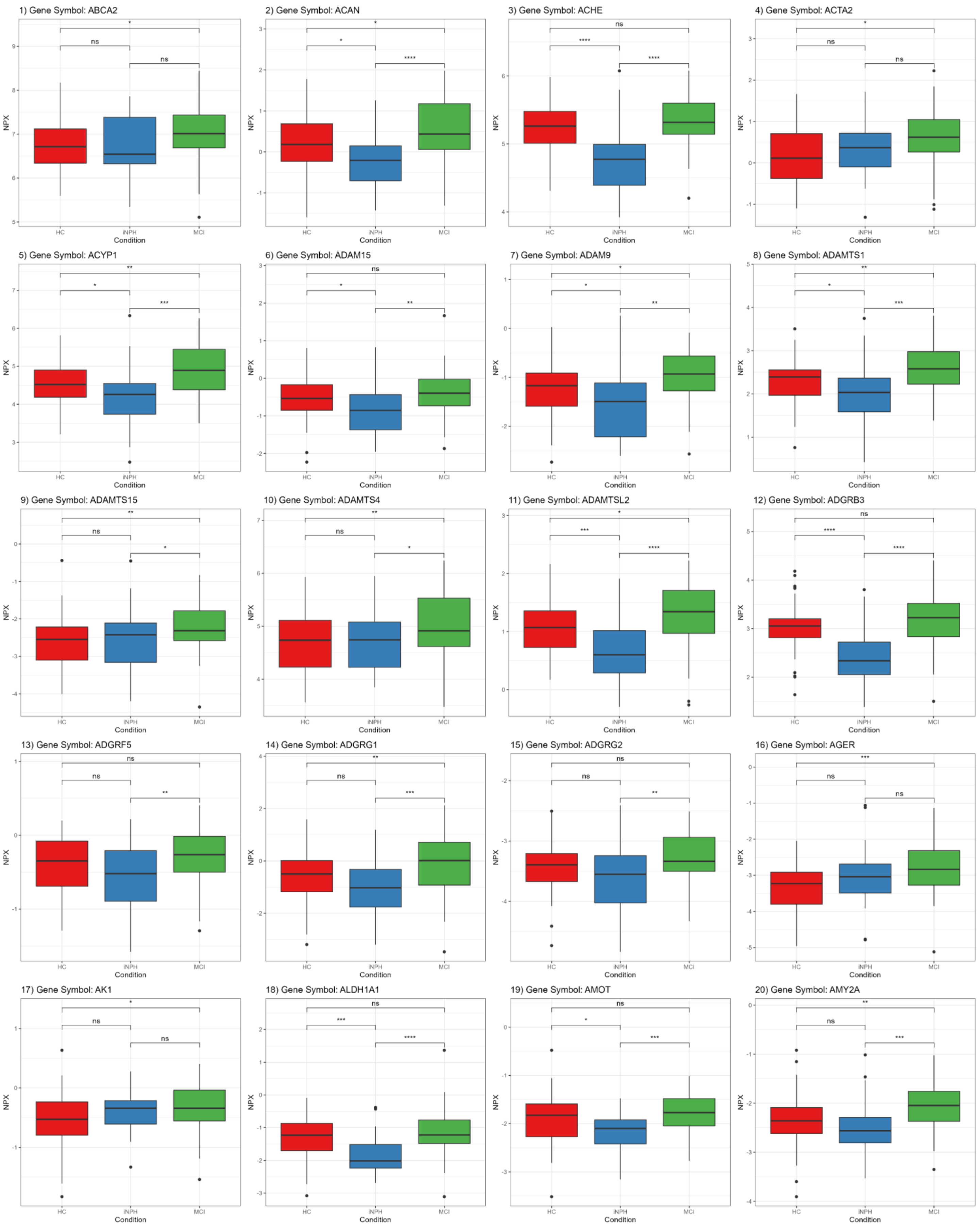
1) Gene Symbol: ABCA2
2) Gene Symbol: ACAN
3) Gene Symbol: ACHE
4) Gene Symbol: ACTA2
5) Gene Symbol: ACYP1
6) Gene Symbol: ADAM15
7) Gene Symbol: ADAM9
8) Gene Symbol: ADAMTS1
9) Gene Symbol: ADAMTS15
10) Gene Symbol: ADAMTS4
11) Gene Symbol: ADAMTSL2
12) Gene Symbol: ADGRB3
13) Gene Symbol: ADGRF5
14) Gene Symbol: ADGRG1
15) Gene Symbol: ADGRG2
16) Gene Symbol: AGER
17) Gene Symbol: AK1
18) Gene Symbol: ALDH1A1
19) Gene Symbol: AMOT
20) Gene Symbol: AMY2A
NPX
Condition
HC
iNPH
MCI

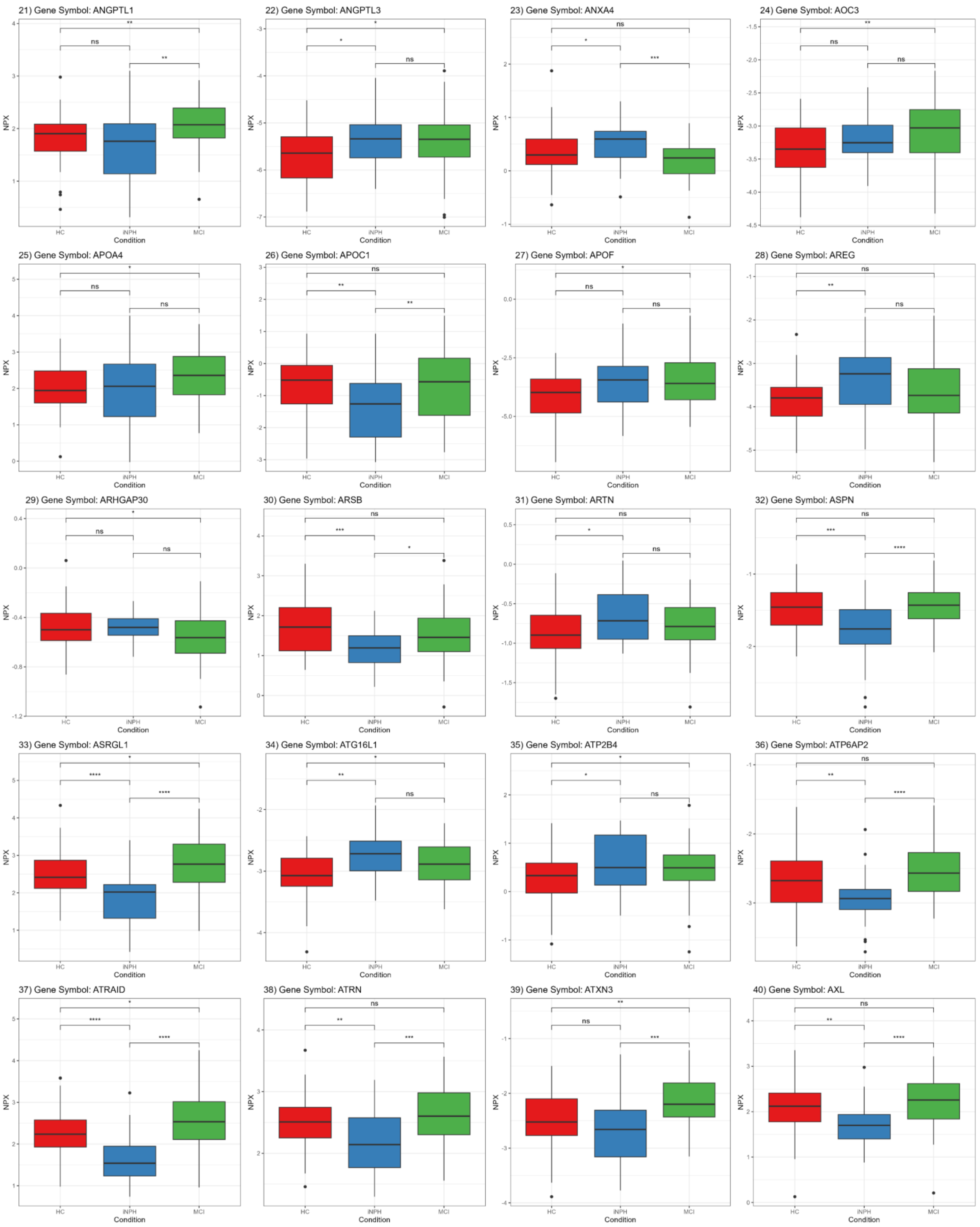
21) Gene Symbol: ANGPTL1
22) Gene Symbol: ANGPTL3
23) Gene Symbol: ANXA4
24) Gene Symbol: AOC3
25) Gene Symbol: APOA4
26) Gene Symbol: APOC1
27) Gene Symbol: APOF
28) Gene Symbol: AREG
29) Gene Symbol: ARHGAP30
30) Gene Symbol: ARSB
31) Gene Symbol: ARTN
32) Gene Symbol: ASPN
33) Gene Symbol: ASRGL1
34) Gene Symbol: ATG16L1
35) Gene Symbol: ATP2B4
36) Gene Symbol: ATP6AP2
37) Gene Symbol: ATRAID
38) Gene Symbol: ATRN
39) Gene Symbol: ATXN3
40) Gene Symbol: AXL
NPX
Condition
HC
iNPH
MCI

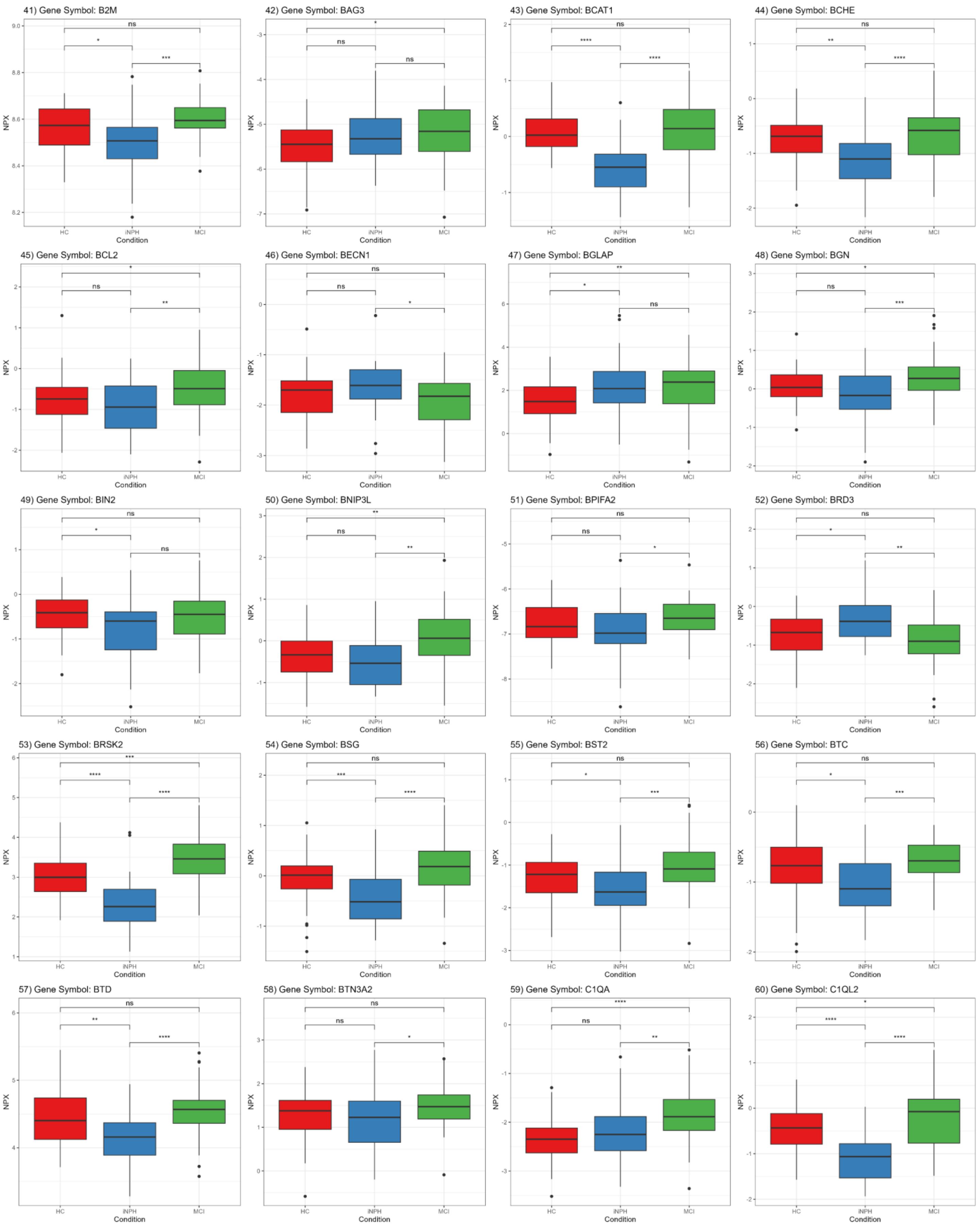
41) Gene Symbol: B2M
42) Gene Symbol: BAG3
43) Gene Symbol: BCAT1
44) Gene Symbol: BCHE
45) Gene Symbol: BCL2
46) Gene Symbol: BECN1
47) Gene Symbol: BGLAP
48) Gene Symbol: BGN
49) Gene Symbol: BIN2
50) Gene Symbol: BNIP3L
51) Gene Symbol: BPIFA2
52) Gene Symbol: BRD3
53) Gene Symbol: BRSK2
54) Gene Symbol: BSG
55) Gene Symbol: BST2
56) Gene Symbol: BTC
57) Gene Symbol: BTD
58) Gene Symbol: BTN3A2
59) Gene Symbol: C1QA
60) Gene Symbol: C1QL2
NPX
Condition
HC
iNPH
MCI

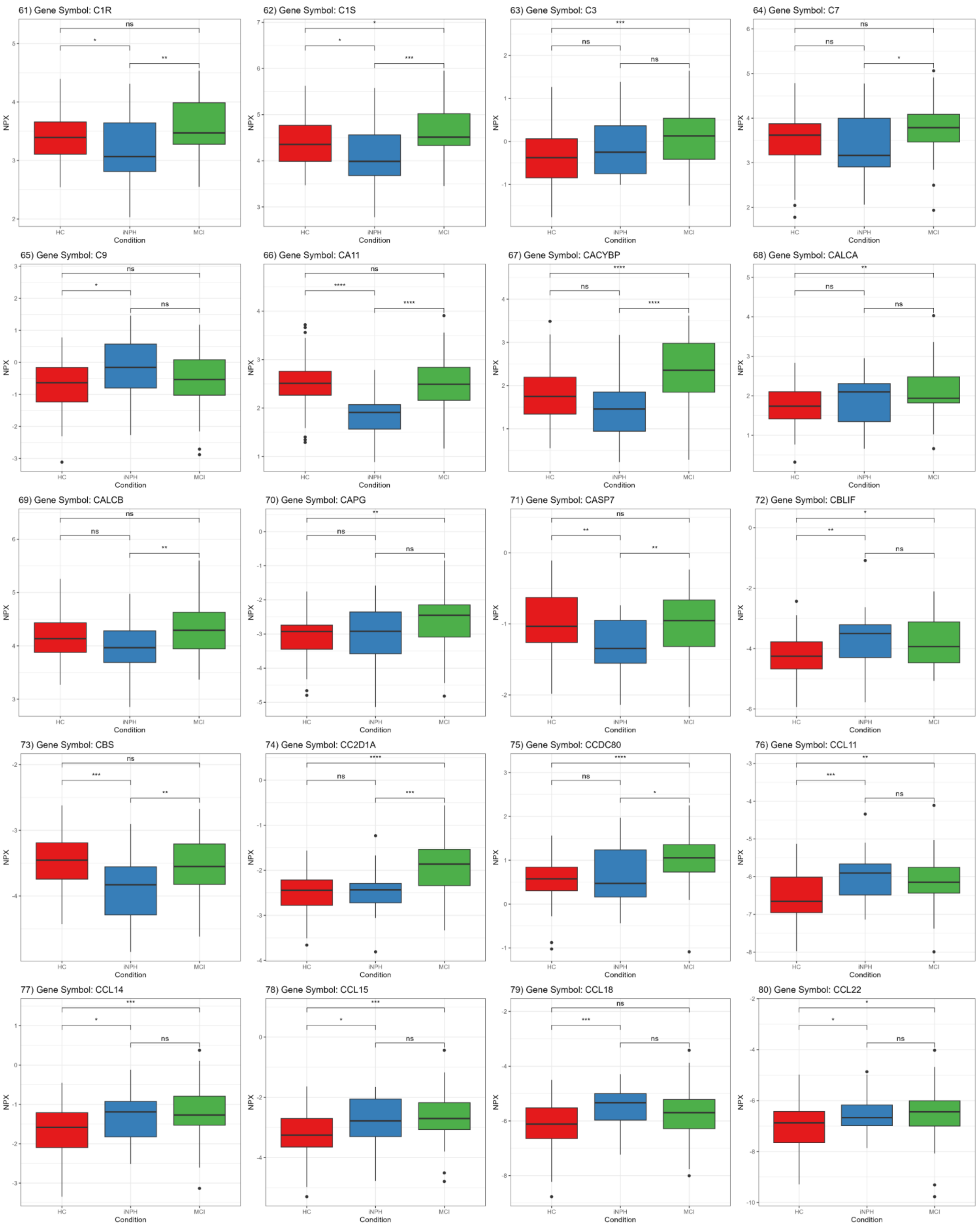
61) Gene Symbol: C1R
ns
*
**
62) Gene Symbol: C1S
*
*
***
63) Gene Symbol: C3
***
ns
ns
64) Gene Symbol: C7
ns
ns
*
65) Gene Symbol: C9
ns
*
ns
66) Gene Symbol: CA11
ns
****
****
67) Gene Symbol: CACYBP
****
ns
****
68) Gene Symbol: CALCA
**
ns
ns
69) Gene Symbol: CALCB
ns
ns
**
70) Gene Symbol: CAPG
**
ns
ns
71) Gene Symbol: CASP7
ns
**
**
72) Gene Symbol: CBLIF
*
**
ns
73) Gene Symbol: CBS
ns
***
**
74) Gene Symbol: CC2D1A
****
ns
***
75) Gene Symbol: CCDC80
****
ns
*
76) Gene Symbol: CCL11
**
***
ns
77) Gene Symbol: CCL14
***
*
ns
78) Gene Symbol: CCL15
***
*
ns
79) Gene Symbol: CCL18
ns
***
ns
80) Gene Symbol: CCL22
*
*
ns
NPX
HC
iNPH
MCI
Condition

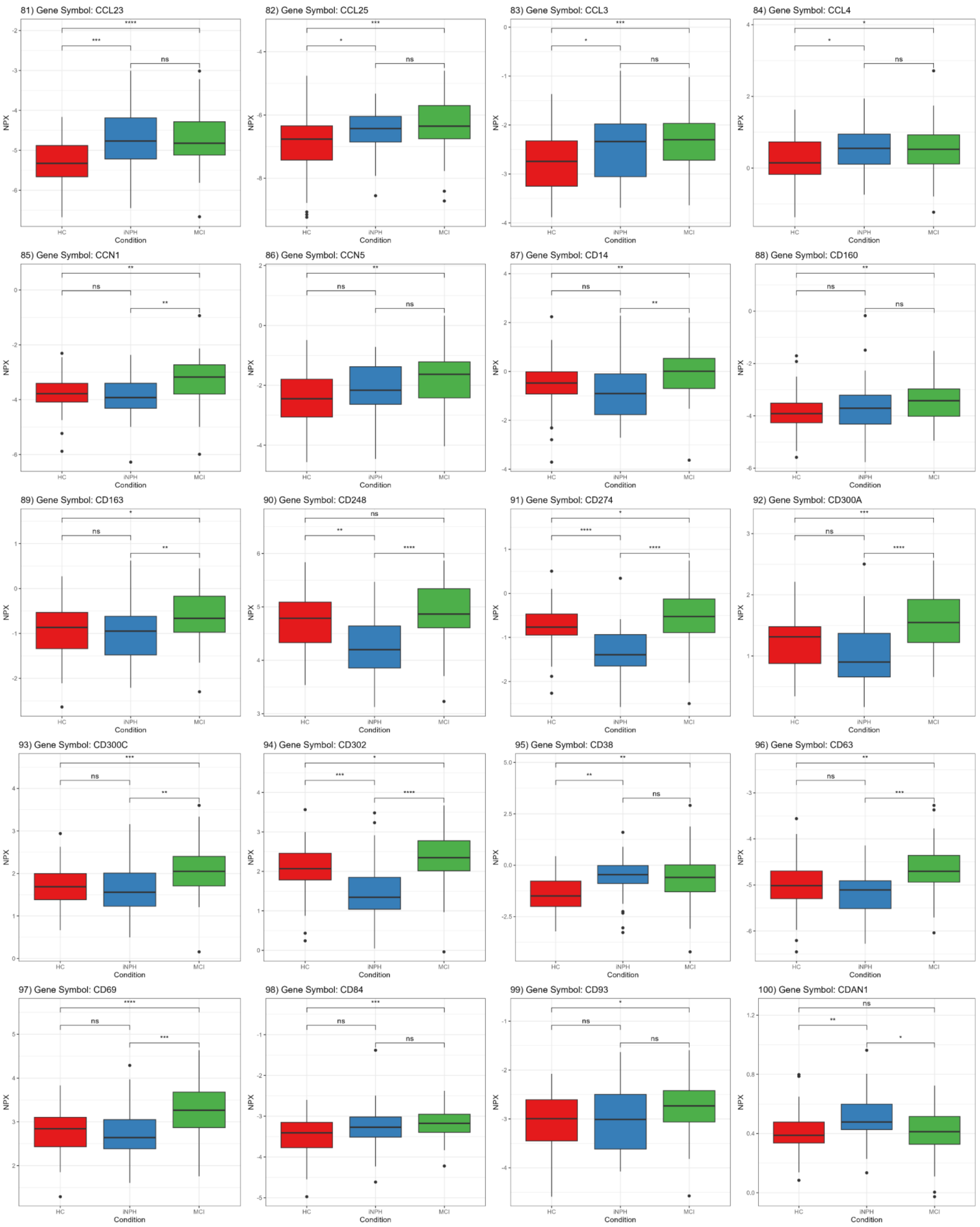
81) Gene Symbol: CCL23
82) Gene Symbol: CCL25
83) Gene Symbol: CCL3
84) Gene Symbol: CCL4
85) Gene Symbol: CCN1
86) Gene Symbol: CCN5
87) Gene Symbol: CD14
88) Gene Symbol: CD160
89) Gene Symbol: CD163
90) Gene Symbol: CD248
91) Gene Symbol: CD274
92) Gene Symbol: CD300A
93) Gene Symbol: CD300C
94) Gene Symbol: CD302
95) Gene Symbol: CD38
96) Gene Symbol: CD63
97) Gene Symbol: CD69
98) Gene Symbol: CD84
99) Gene Symbol: CD93
100) Gene Symbol: CDAN1
NPX
Condition
HC
iNPH
MCI

101) Gene Symbol: CDCP1

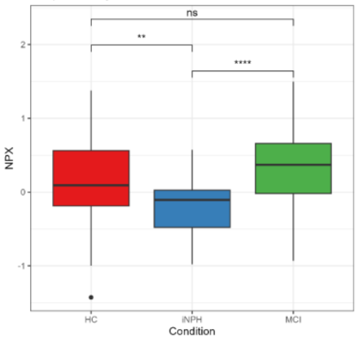


102) Gene Symbol: CDH5

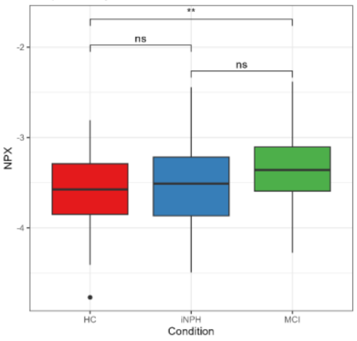


103) Gene Symbol: CDNF

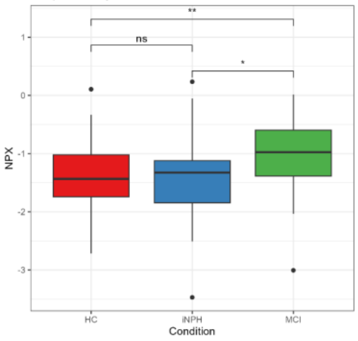


104) Gene Symbol: CDON

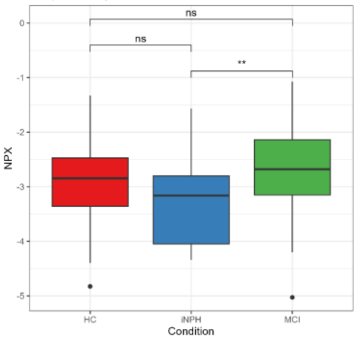


105) Gene Symbol: CEACAM16

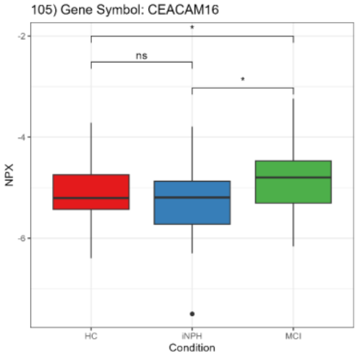


106) Gene Symbol: CEND1

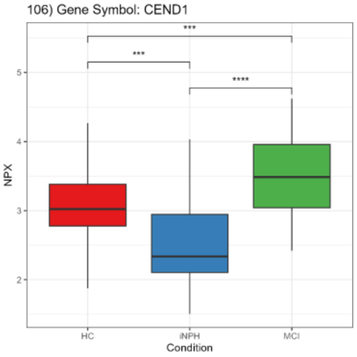


107) Gene Symbol: CEP170

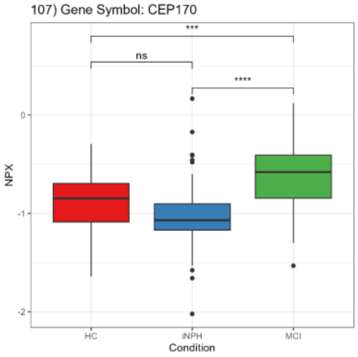


108) Gene Symbol: CFH

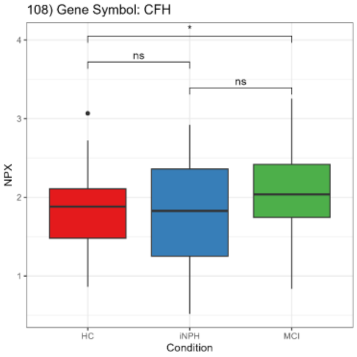


109) Gene Symbol: CFHR2

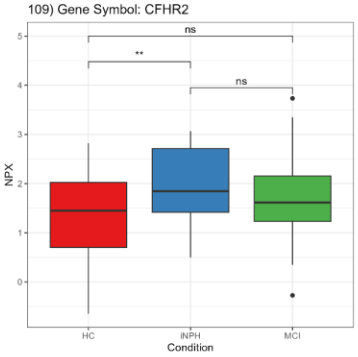


110) Gene Symbol: CFHR5

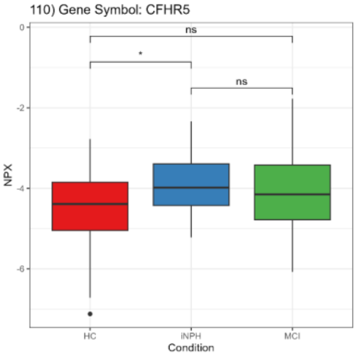


111) Gene Symbol: CHCHD10

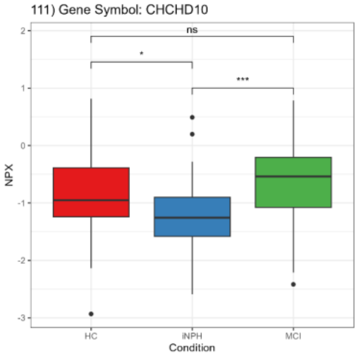


112) Gene Symbol: CHRDL2

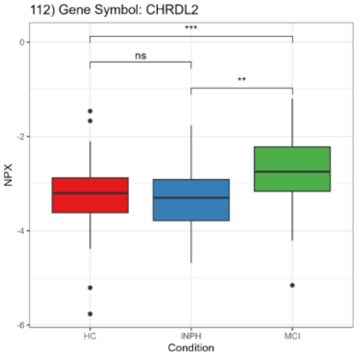


113) Gene Symbol: CKAP4

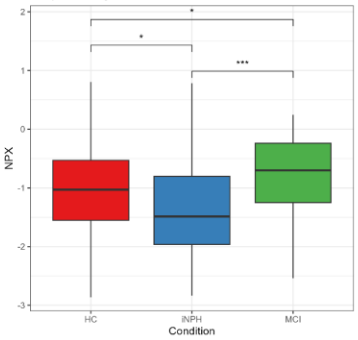


114) Gene Symbol: CLEC3B

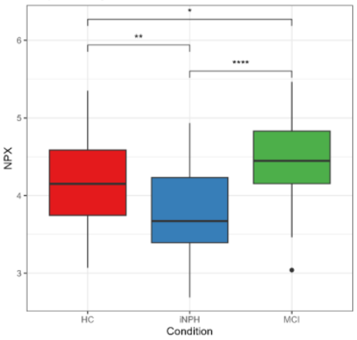


115) Gene Symbol: CLEC4M

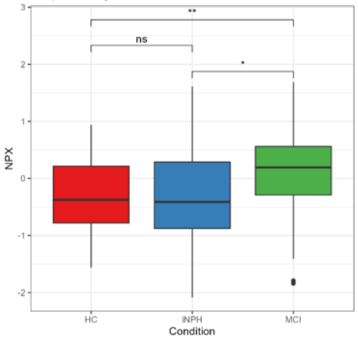


116) Gene Symbol: CLEC5A

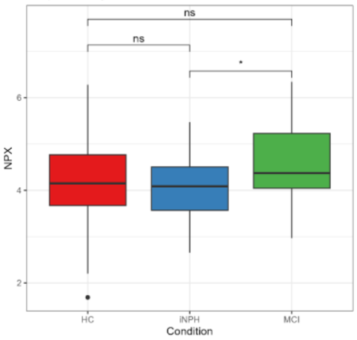


117) Gene Symbol: CLEC7A

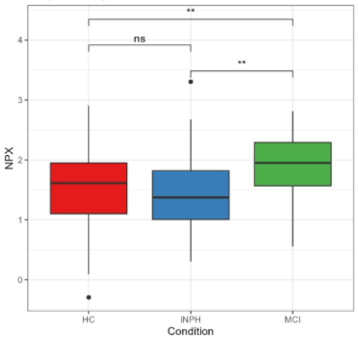


118) Gene Symbol: CLIP2

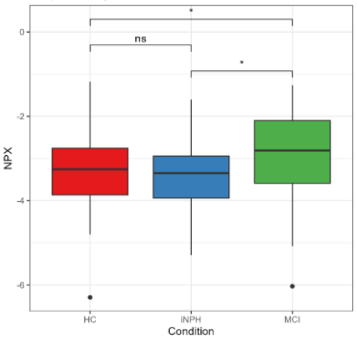


119) Gene Symbol: CLPS

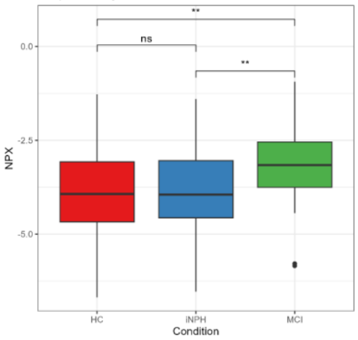


120) Gene Symbol: CLSPN

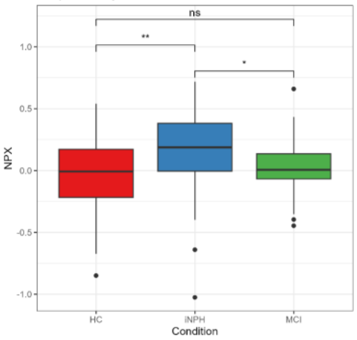

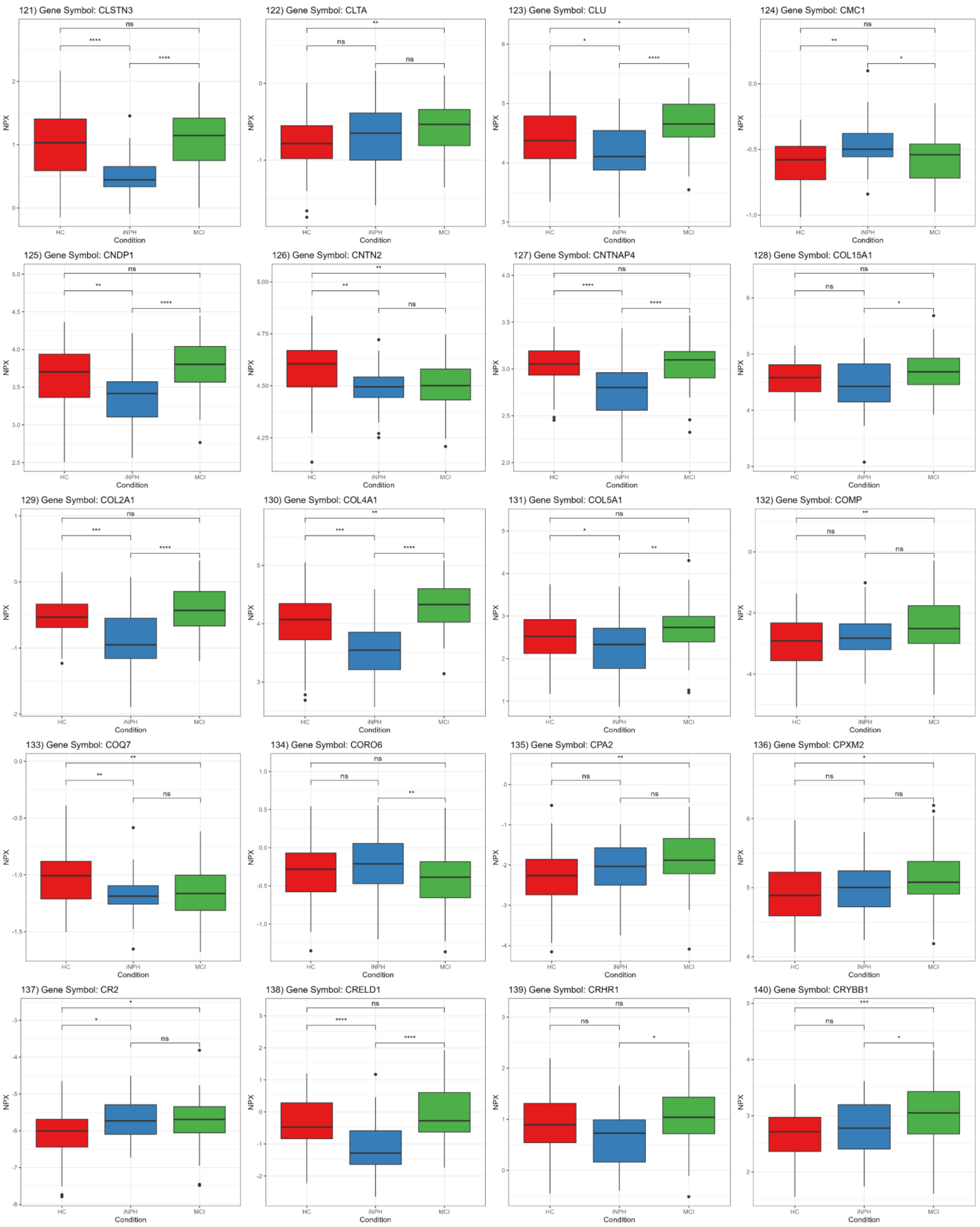
121) Gene Symbol: CLSTN3
122) Gene Symbol: CLTA
123) Gene Symbol: CLU
124) Gene Symbol: CMC1
125) Gene Symbol: CNDP1
126) Gene Symbol: CNTN2
127) Gene Symbol: CNTNAP4
128) Gene Symbol: COL15A1
129) Gene Symbol: COL2A1
130) Gene Symbol: COL4A1
131) Gene Symbol: COL5A1
132) Gene Symbol: COMP
133) Gene Symbol: COQ7
134) Gene Symbol: CORO6
135) Gene Symbol: CPA2
136) Gene Symbol: CPXM2
137) Gene Symbol: CR2
138) Gene Symbol: CRELD1
139) Gene Symbol: CRHR1
140) Gene Symbol: CRYBB1
NPX
Condition
HC
iNPH
MCI

141) Gene Symbol: CRYGD

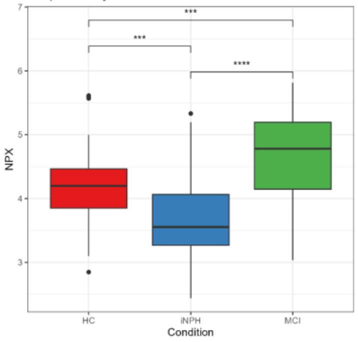


142) Gene Symbol: CSF1R

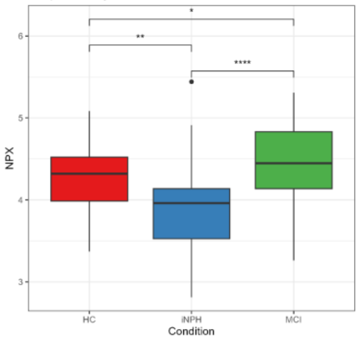


143) Gene Symbol: CSF2RA

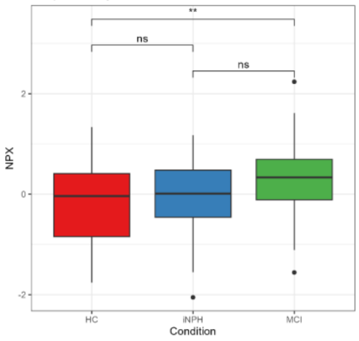


144) Gene Symbol: CSPG4

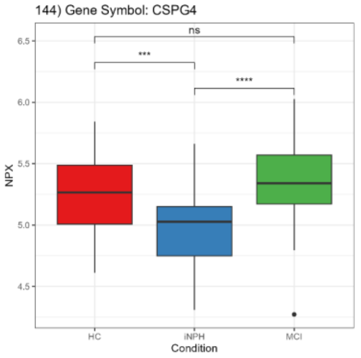


145) Gene Symbol: CSPG5

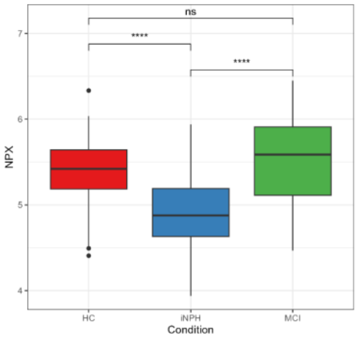


146) Gene Symbol: CST1

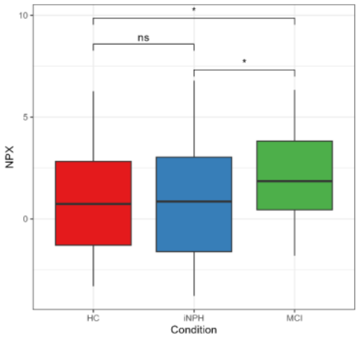


147) Gene Symbol: CST6

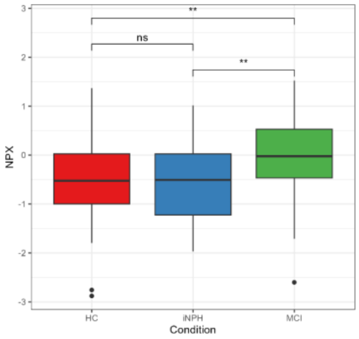


148) Gene Symbol: CSTB

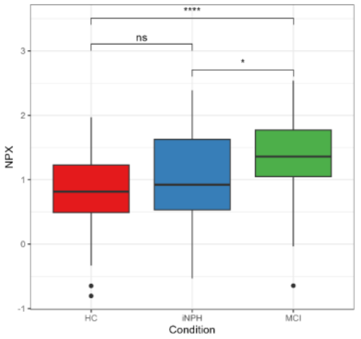


149) Gene Symbol: CTBS

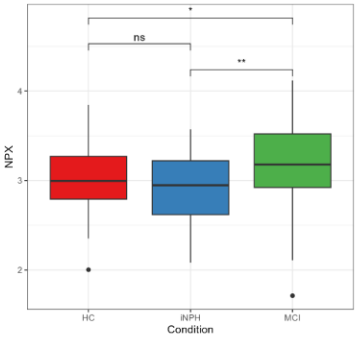


150) Gene Symbol: CTSS

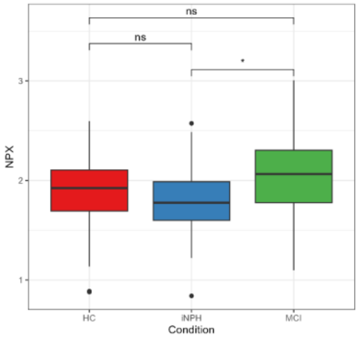


151) Gene Symbol: CWC15

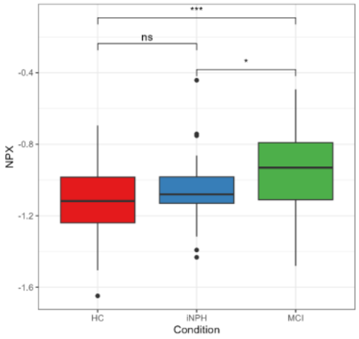


152) Gene Symbol: CX3CL1

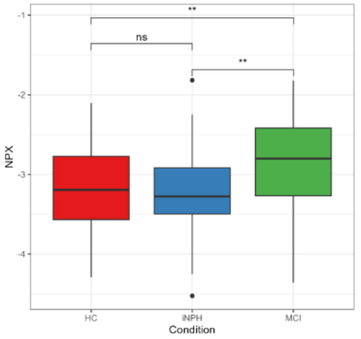


153) Gene Symbol: CXADR

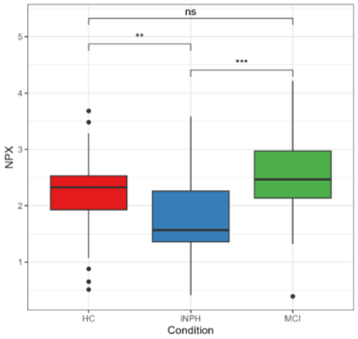


154) Gene Symbol: CXCL10

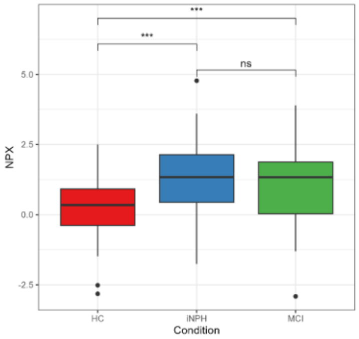


155) Gene Symbol: CXCL17

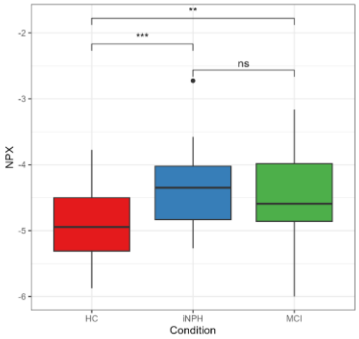


156) Gene Symbol: DDT

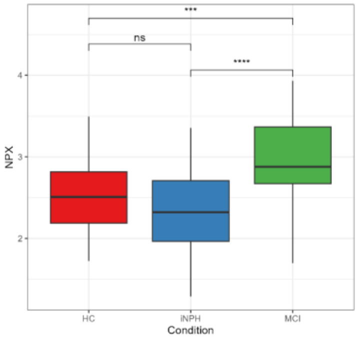


157) Gene Symbol: DIPK2B

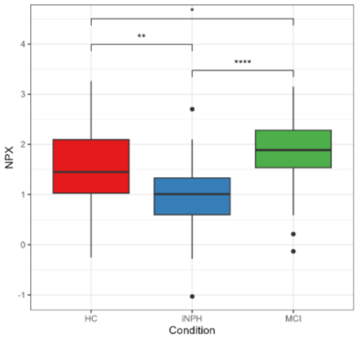


158) Gene Symbol: DKK4

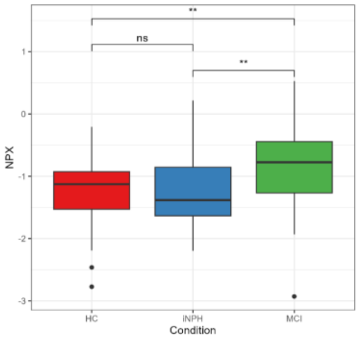


159) Gene Symbol: DKKL1

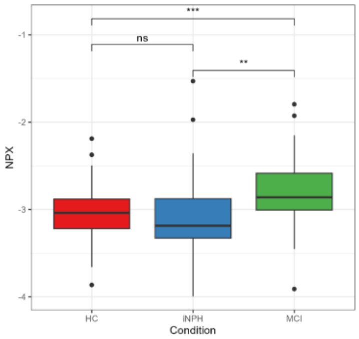


160) Gene Symbol: DNAJA2

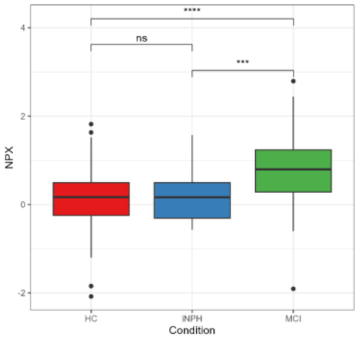

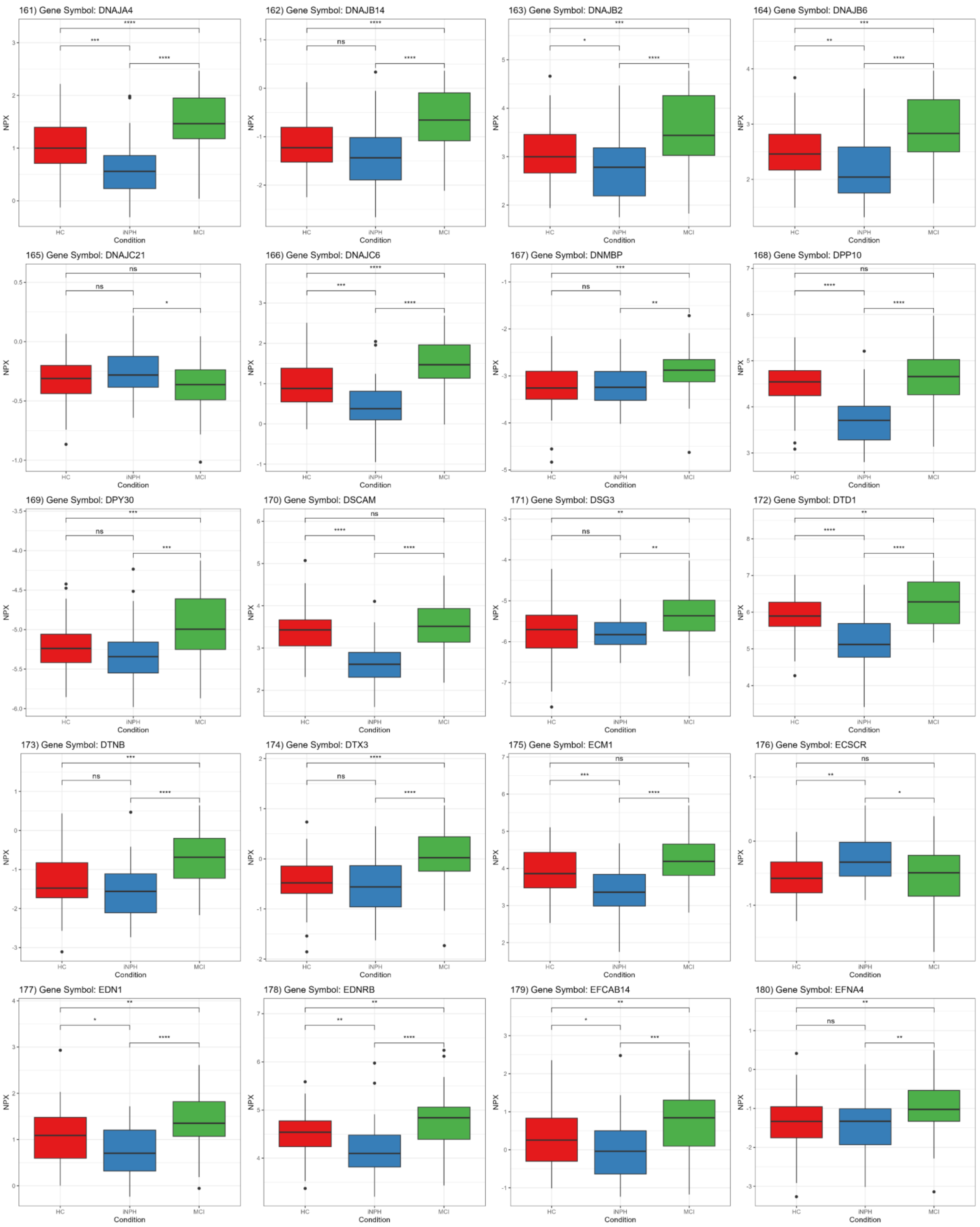
161) Gene Symbol: DNAJA4
162) Gene Symbol: DNAJB14
163) Gene Symbol: DNAJB2
164) Gene Symbol: DNAJB6
165) Gene Symbol: DNAJC21
166) Gene Symbol: DNAJC6
167) Gene Symbol: DNMBP
168) Gene Symbol: DPP10
169) Gene Symbol: DPY30
170) Gene Symbol: DSCAM
171) Gene Symbol: DSG3
172) Gene Symbol: DTD1
173) Gene Symbol: DTNB
174) Gene Symbol: DTX3
175) Gene Symbol: ECM1
176) Gene Symbol: ECSCR
177) Gene Symbol: EDN1
178) Gene Symbol: EDNRB
179) Gene Symbol: EFCAB14
180) Gene Symbol: EFNA4
NPX
Condition
HC
iNPH
MCI

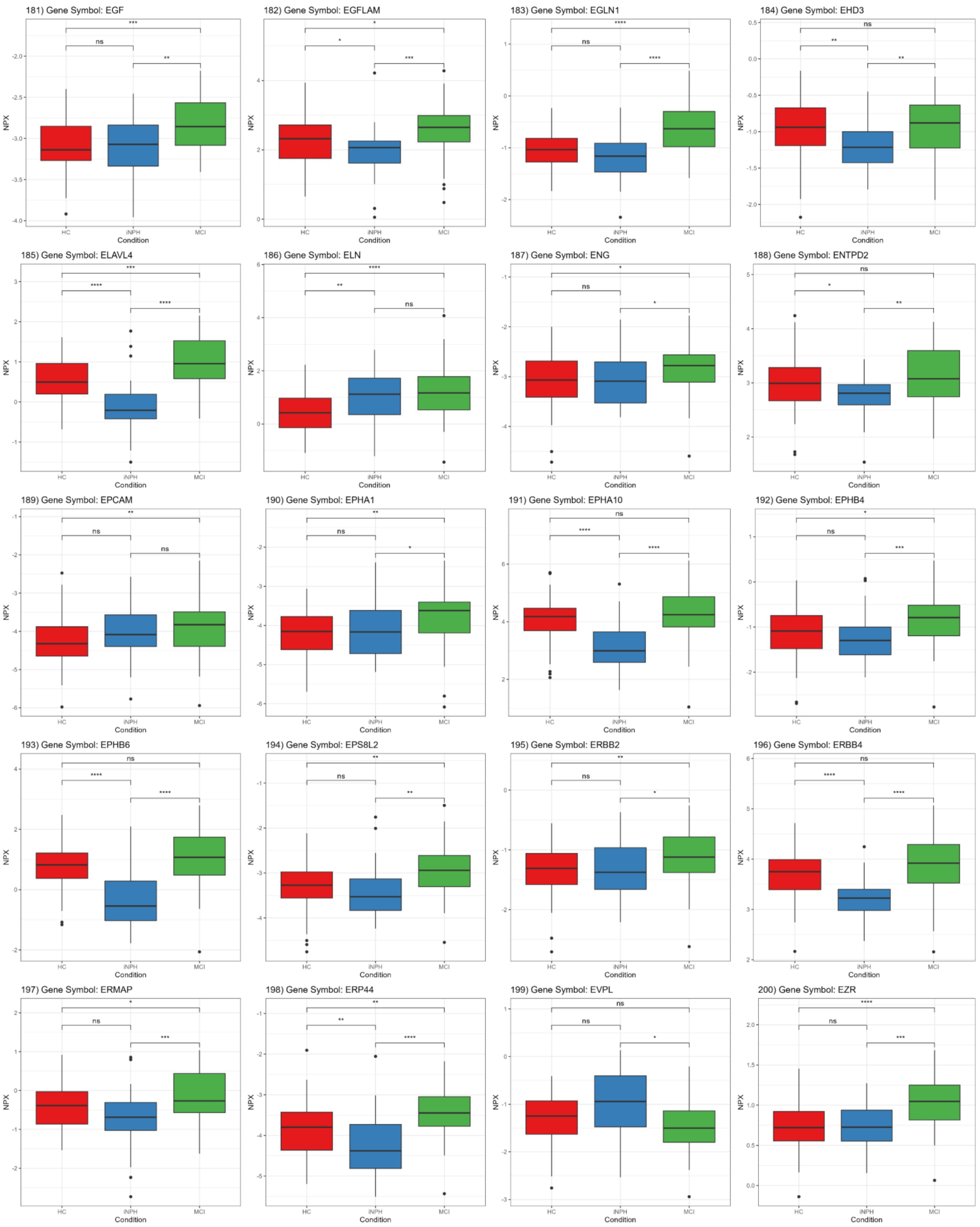
181) Gene Symbol: EGF
182) Gene Symbol: EGFLAM
183) Gene Symbol: EGLN1
184) Gene Symbol: EHD3
185) Gene Symbol: ELAVL4
186) Gene Symbol: ELN
187) Gene Symbol: ENG
188) Gene Symbol: ENTPD2
189) Gene Symbol: EPCAM
190) Gene Symbol: EPHA1
191) Gene Symbol: EPHA10
192) Gene Symbol: EPHB4
193) Gene Symbol: EPHB6
194) Gene Symbol: EPS8L2
195) Gene Symbol: ERBB2
196) Gene Symbol: ERBB4
197) Gene Symbol: ERMAP
198) Gene Symbol: ERP44
199) Gene Symbol: EVPL
200) Gene Symbol: EZR
NPX
Condition
HC
iNPH
MCI

201) Gene Symbol: FABP3

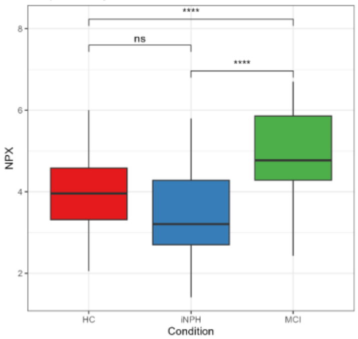


202) Gene Symbol: FAM171B

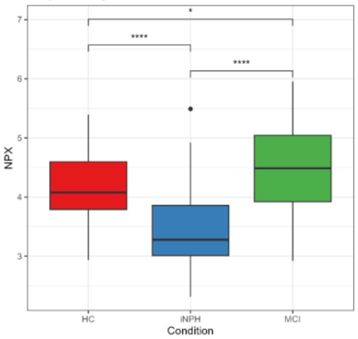


203) Gene Symbol: FAM20A

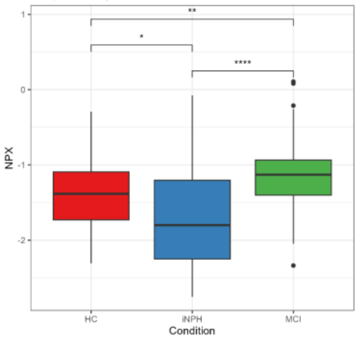


204) Gene Symbol: FAS

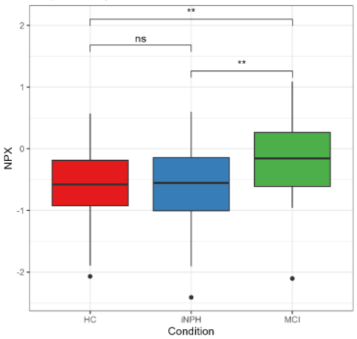


205) Gene Symbol: FASLG

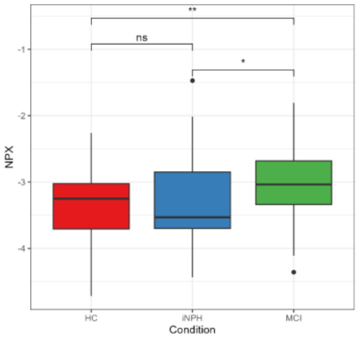


206) Gene Symbol: FCAR

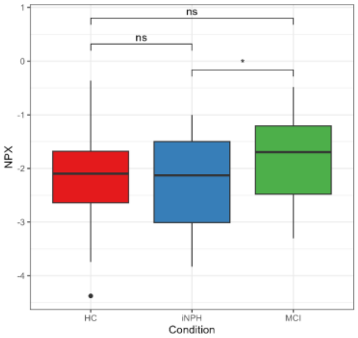


207) Gene Symbol: FGF12

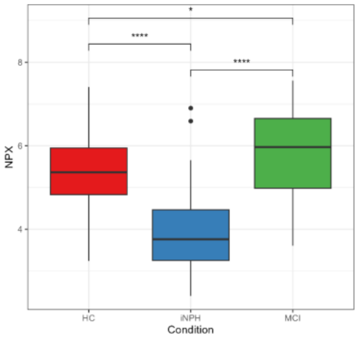


208) Gene Symbol: FGFBP3

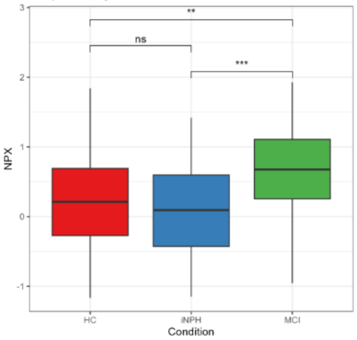


209) Gene Symbol: FGFR4

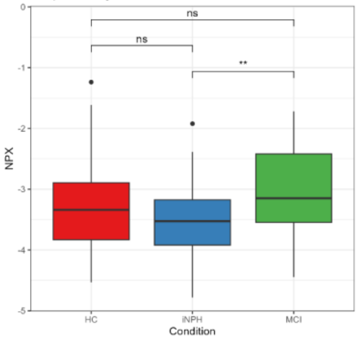


210) Gene Symbol: FGR

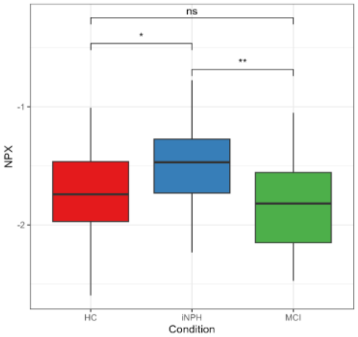


211) Gene Symbol: FIS1

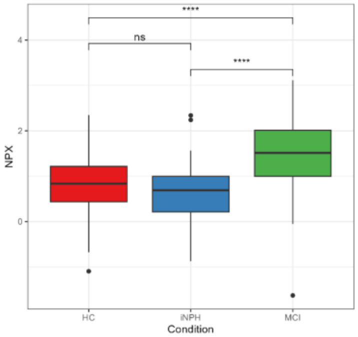


212) Gene Symbol: FKBP4

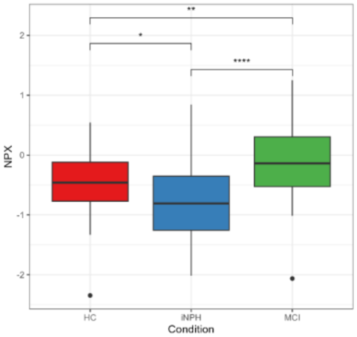


213) Gene Symbol: FKBP5

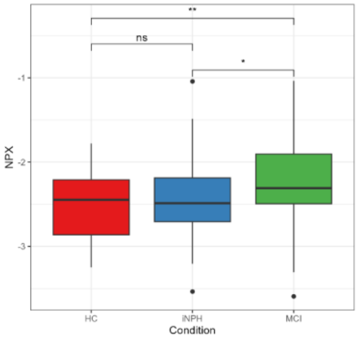


214) Gene Symbol: FLT1

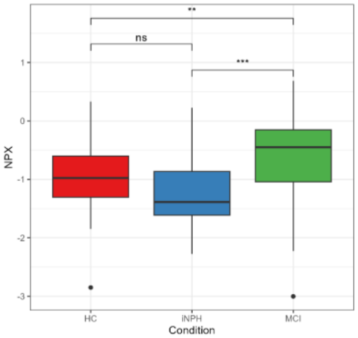


215) Gene Symbol: FLT3

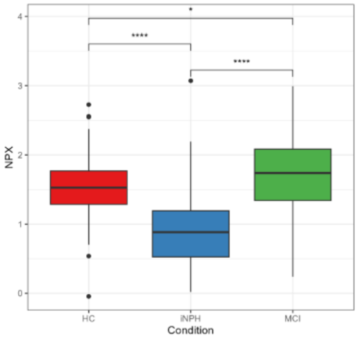


216) Gene Symbol: FLT3LG

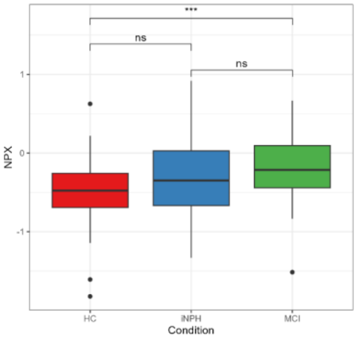


217) Gene Symbol: FN1

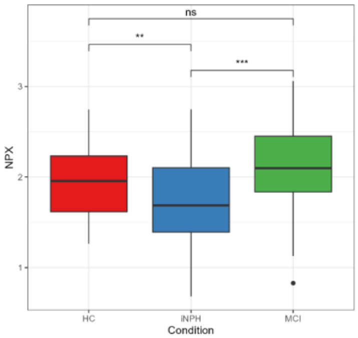


218) Gene Symbol: FOSB

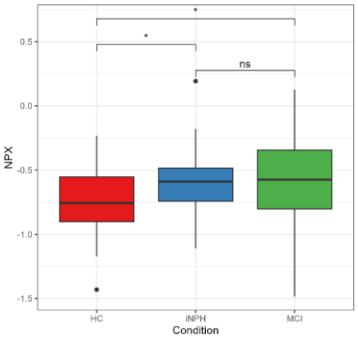


219) Gene Symbol: FOXJ3

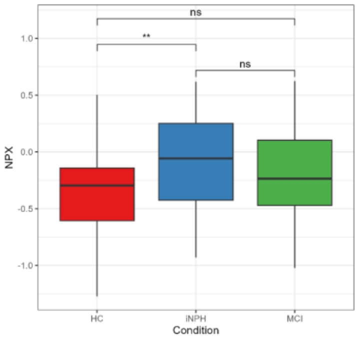


220) Gene Symbol: FUT1

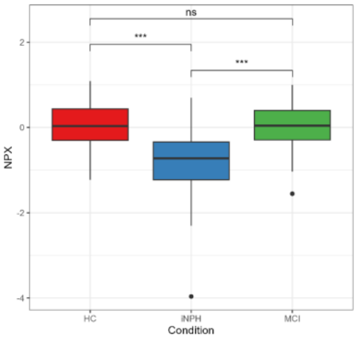

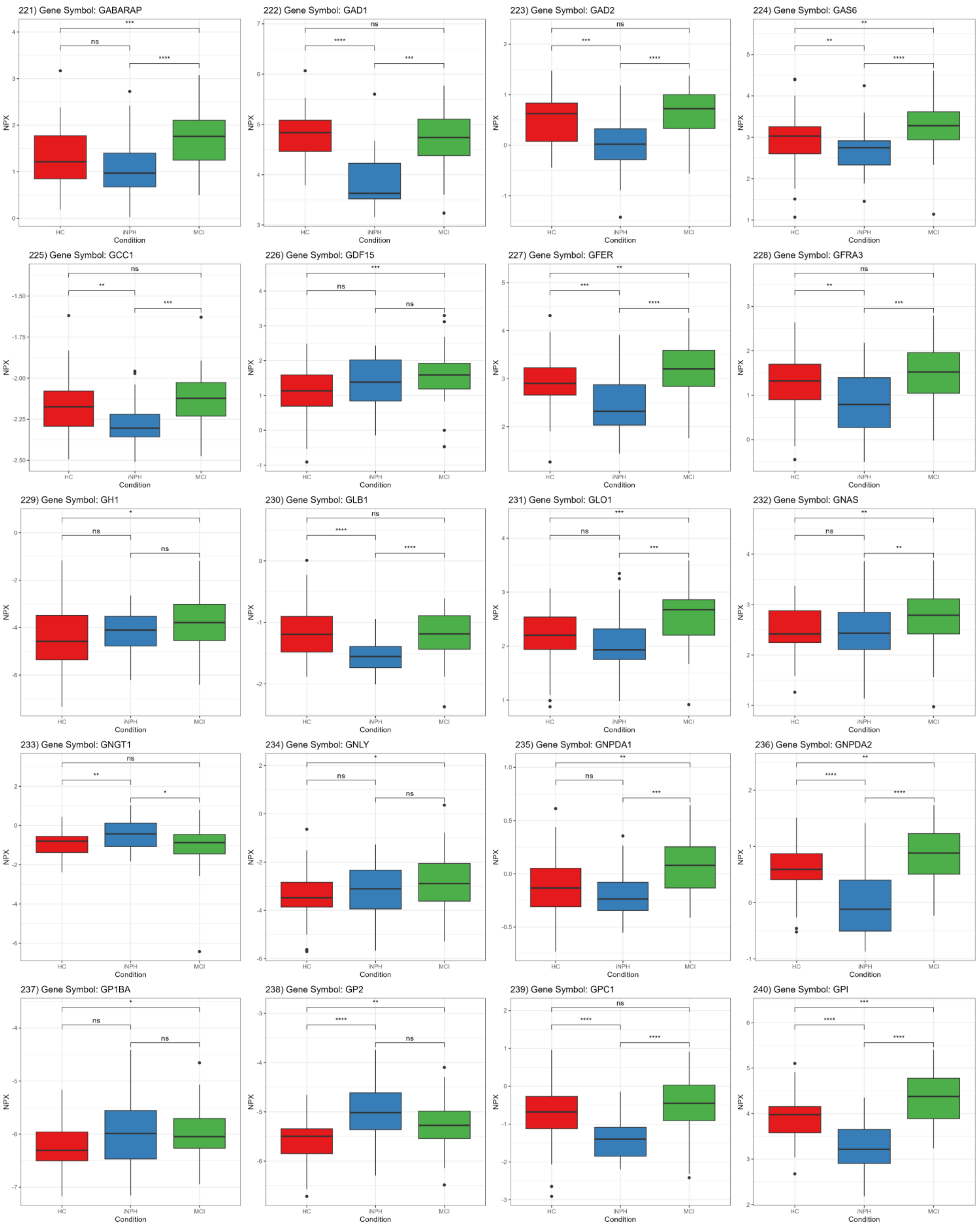
221) Gene Symbol: GABARAP
222) Gene Symbol: GAD1
223) Gene Symbol: GAD2
224) Gene Symbol: GAS6
225) Gene Symbol: GCC1
226) Gene Symbol: GDF15
227) Gene Symbol: GFER
228) Gene Symbol: GFRA3
229) Gene Symbol: GH1
230) Gene Symbol: GLB1
231) Gene Symbol: GLO1
232) Gene Symbol: GNAS
233) Gene Symbol: GNGT1
234) Gene Symbol: GNLY
235) Gene Symbol: GNPDA1
236) Gene Symbol: GNPDA2
237) Gene Symbol: GP1BA
238) Gene Symbol: GP2
239) Gene Symbol: GPC1
240) Gene Symbol: GPI
NPX
Condition
HC
iNPH
MCI

241) Gene Symbol: GRIK2

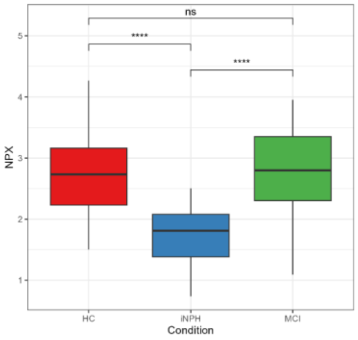


242) Gene Symbol: GSN

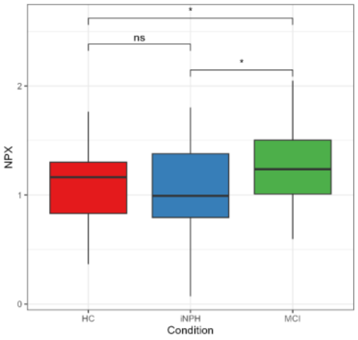


243) Gene Symbol: GSTA3

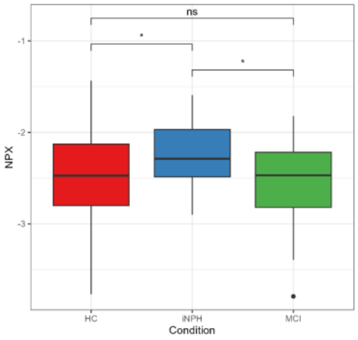


244) Gene Symbol: GUCA2A

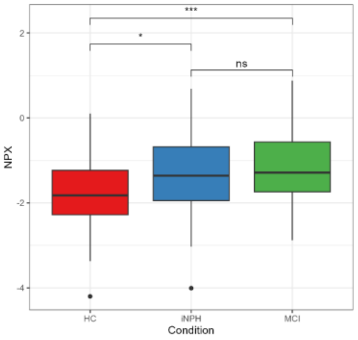


245) Gene Symbol: GUK1

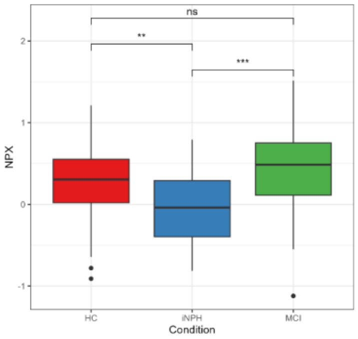


246) Gene Symbol: HAVCR1

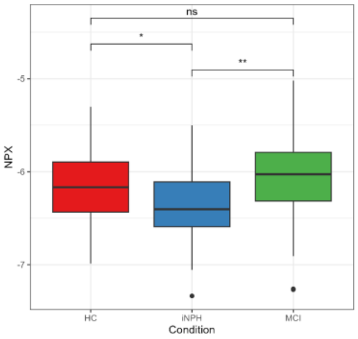


247) Gene Symbol: HDDC2

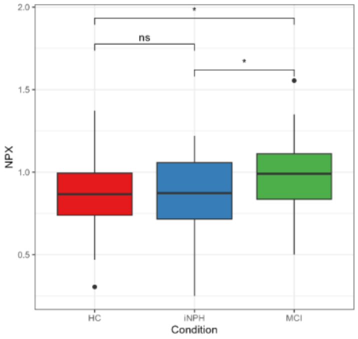


248) Gene Symbol: HEBP1

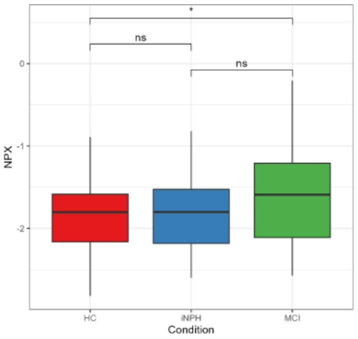


249) Gene Symbol: HEG1

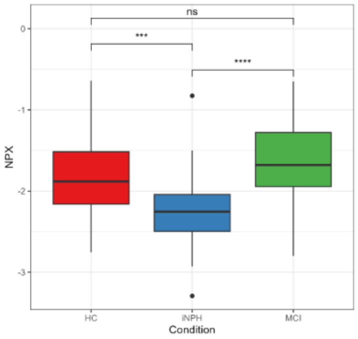


250) Gene Symbol: HEPH

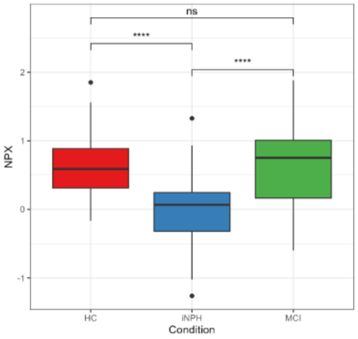


251) Gene Symbol: HGFAC

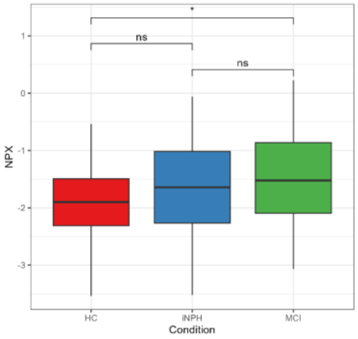


252) Gene Symbol: HGS

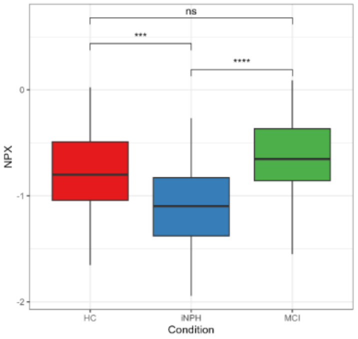


253) Gene Symbol: HIP1R

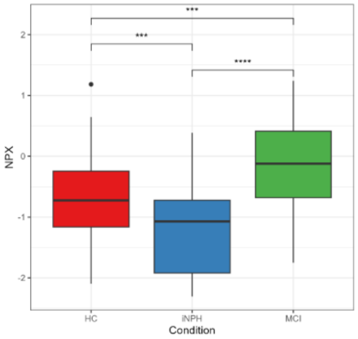


254) Gene Symbol: HMOX2

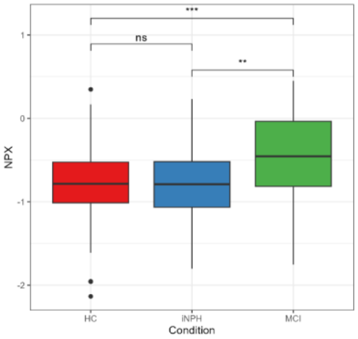


255) Gene Symbol: HS6ST1

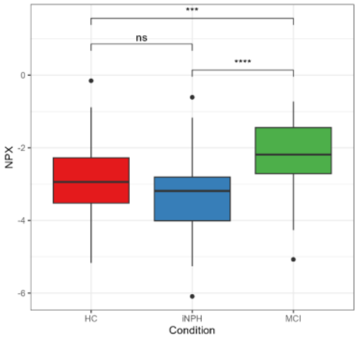


256) Gene Symbol: HSPB1

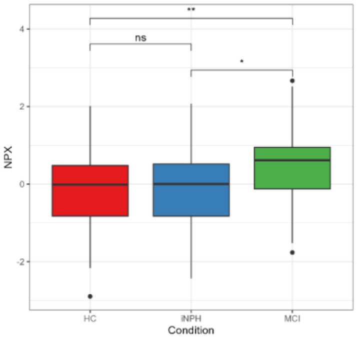


257) Gene Symbol: HSPB6

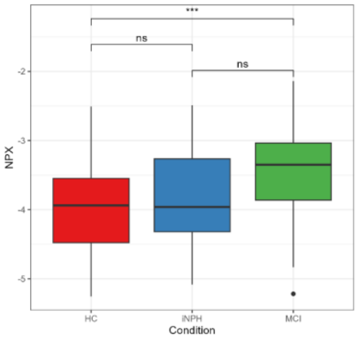


258) Gene Symbol: HTRA2

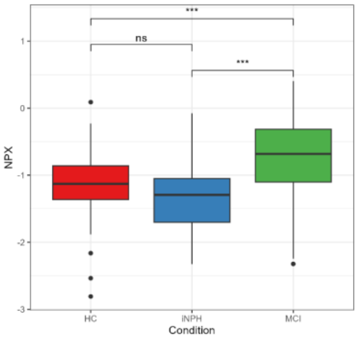


259) Gene Symbol: HYOU1

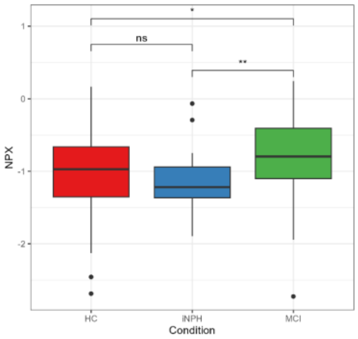


260) Gene Symbol: ICAM1

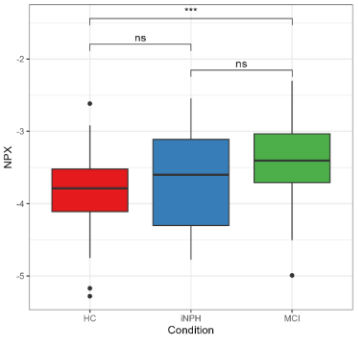

261) Gene Symbol: ICAM2

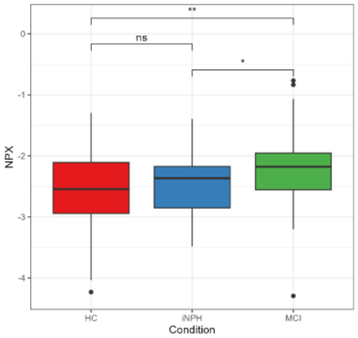


262) Gene Symbol: ICAM5

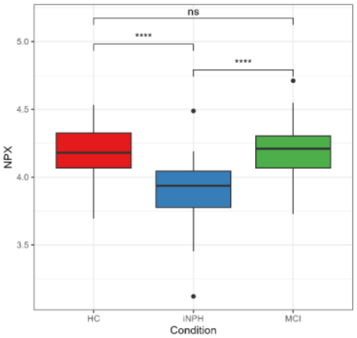


263) Gene Symbol: IFNAR1

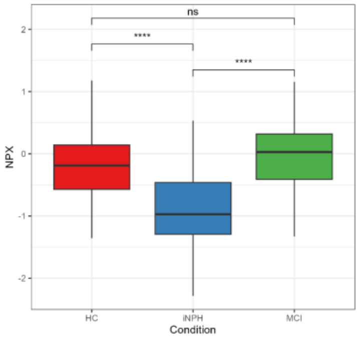


264) Gene Symbol: IFNGR2

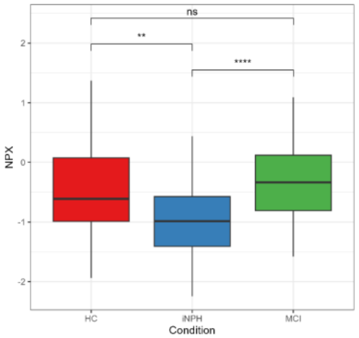


265) Gene Symbol: IFNLR1

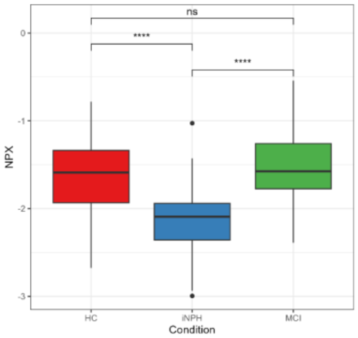


266) Gene Symbol: IGDCC3

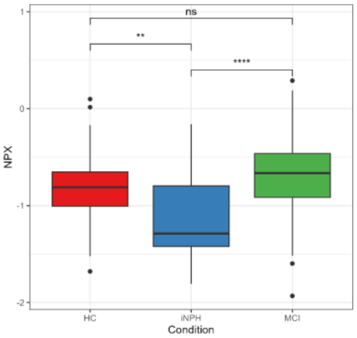


267) Gene Symbol: IGDCC4

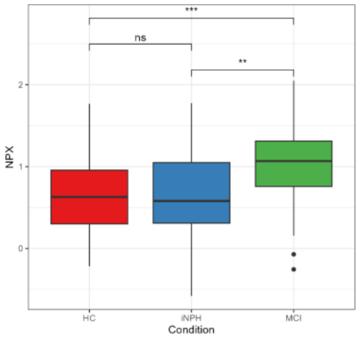


268) Gene Symbol: IGF1R

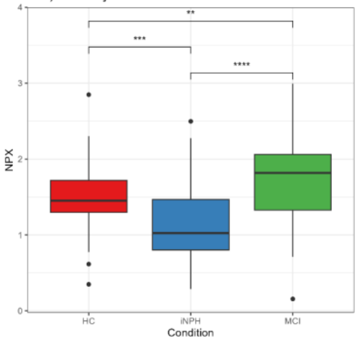


269) Gene Symbol: IGFBP1

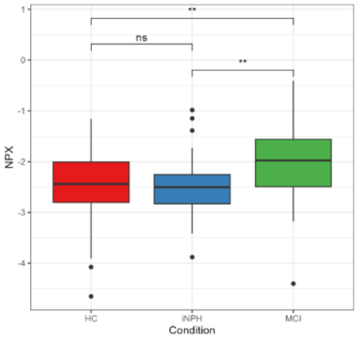


270) Gene Symbol: IGFL4

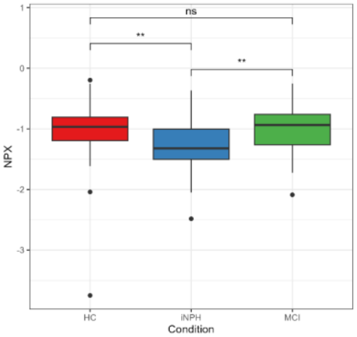


271) Gene Symbol: IGLON5

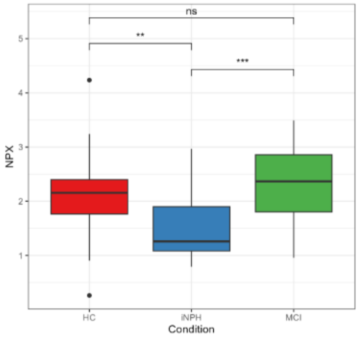


272) Gene Symbol: IGSF3

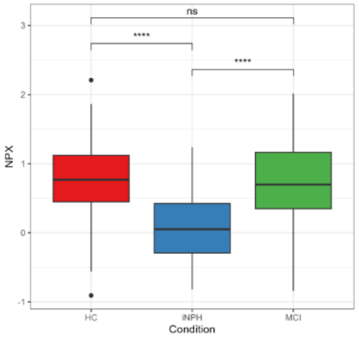


273) Gene Symbol: IL13RA1

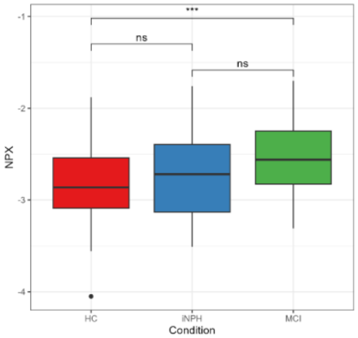


274) Gene Symbol: IL15

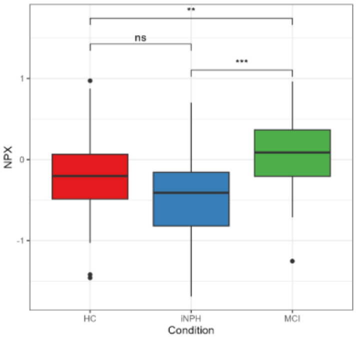


275) Gene Symbol: IL17RA

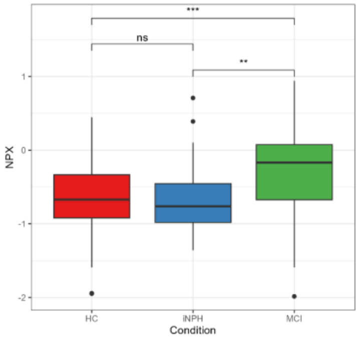


276) Gene Symbol: IL17RB

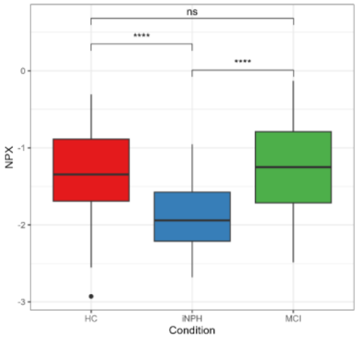


277) Gene Symbol: IL18BP

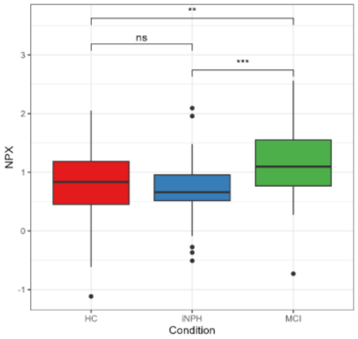


278) Gene Symbol: IL1R1

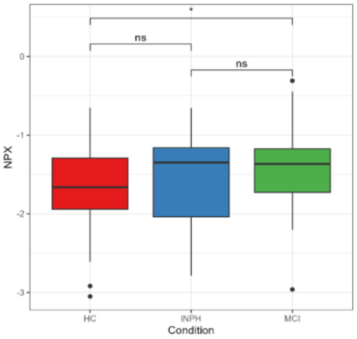


279) Gene Symbol: IL1RL2

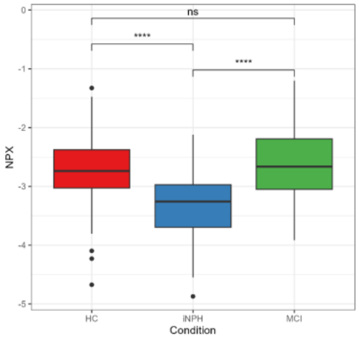


280) Gene Symbol: IL22

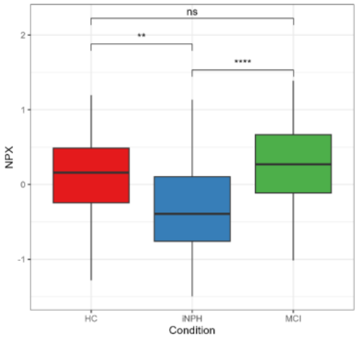

281) Gene Symbol: IL24

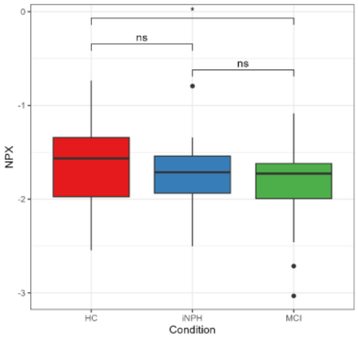


282) Gene Symbol: IL2RA

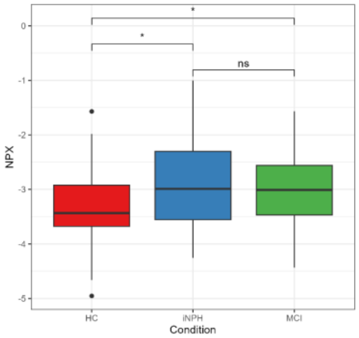


283) Gene Symbol: IL7R

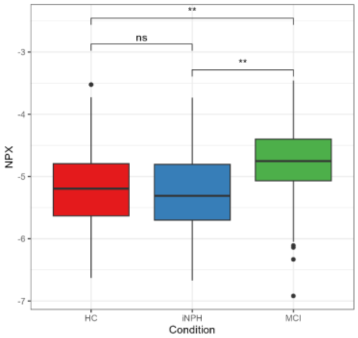


284) Gene Symbol: IMMT

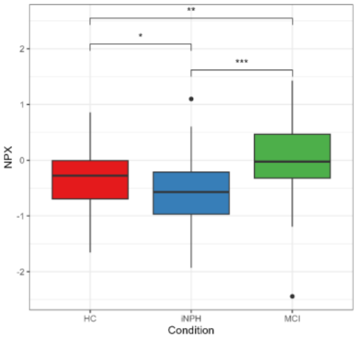


285) Gene Symbol: IPCEF1

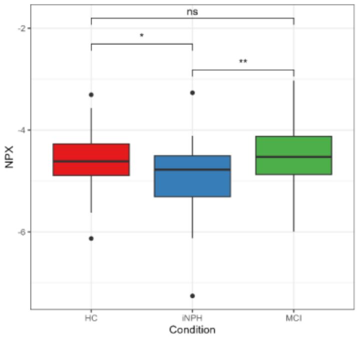


286) Gene Symbol: IRAK1

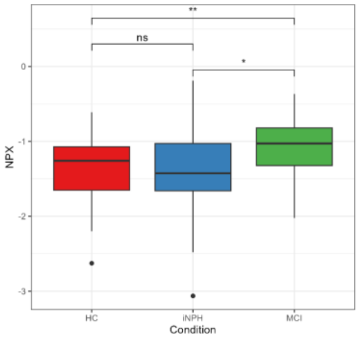


287) Gene Symbol: ISM1

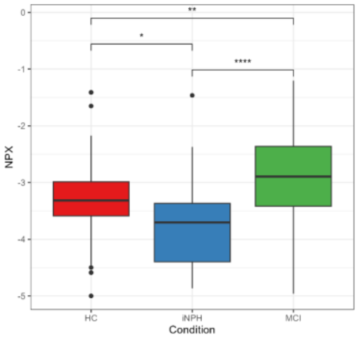


288) Gene Symbol: ISM2

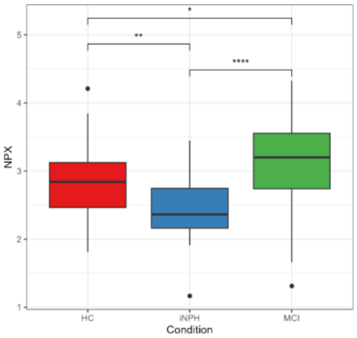


289) Gene Symbol: ITGA11

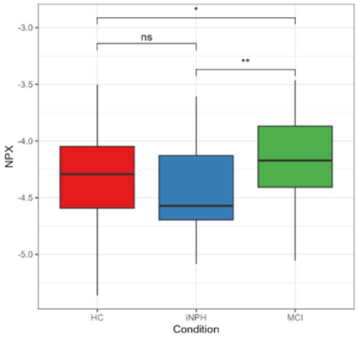


290) Gene Symbol: ITGAM

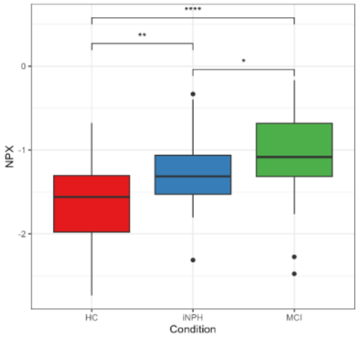


291) Gene Symbol: ITGB2

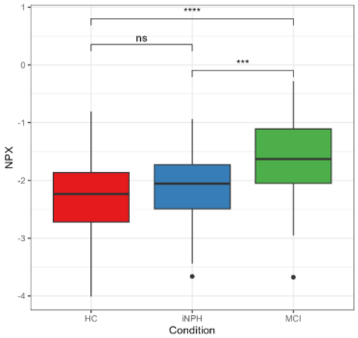


292) Gene Symbol: ITGBL1

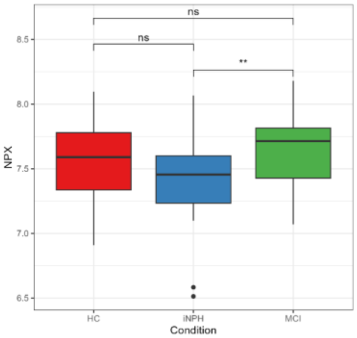


293) Gene Symbol: ITIH1

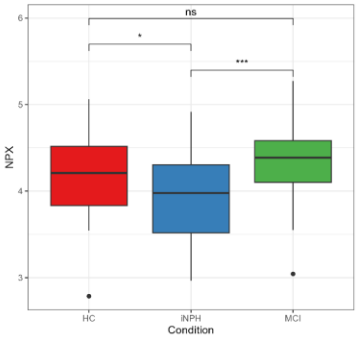


294) Gene Symbol: ITM2A

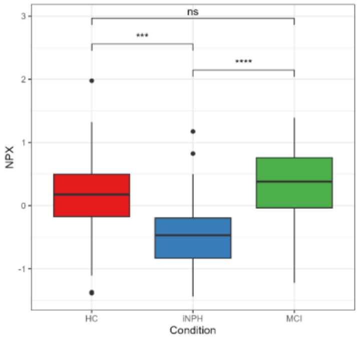


295) Gene Symbol: JAM3

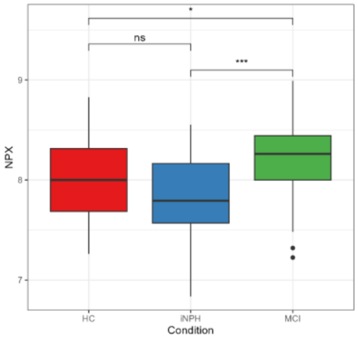


296) Gene Symbol: KCNC4

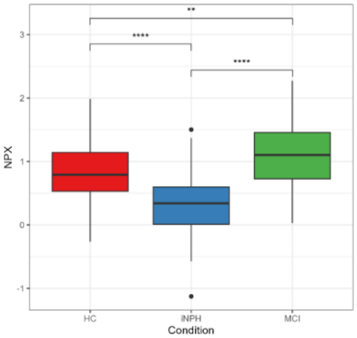


297) Gene Symbol: KHK

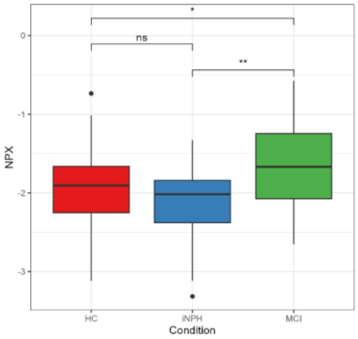


298) Gene Symbol: KIRREL2

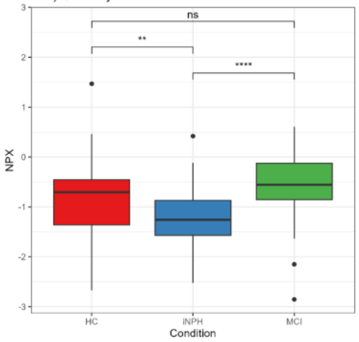


299) Gene Symbol: KLK10

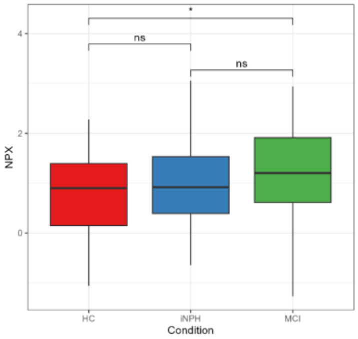


300) Gene Symbol: KLK7

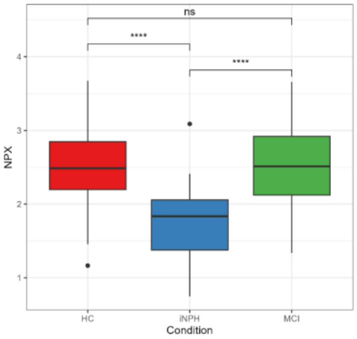

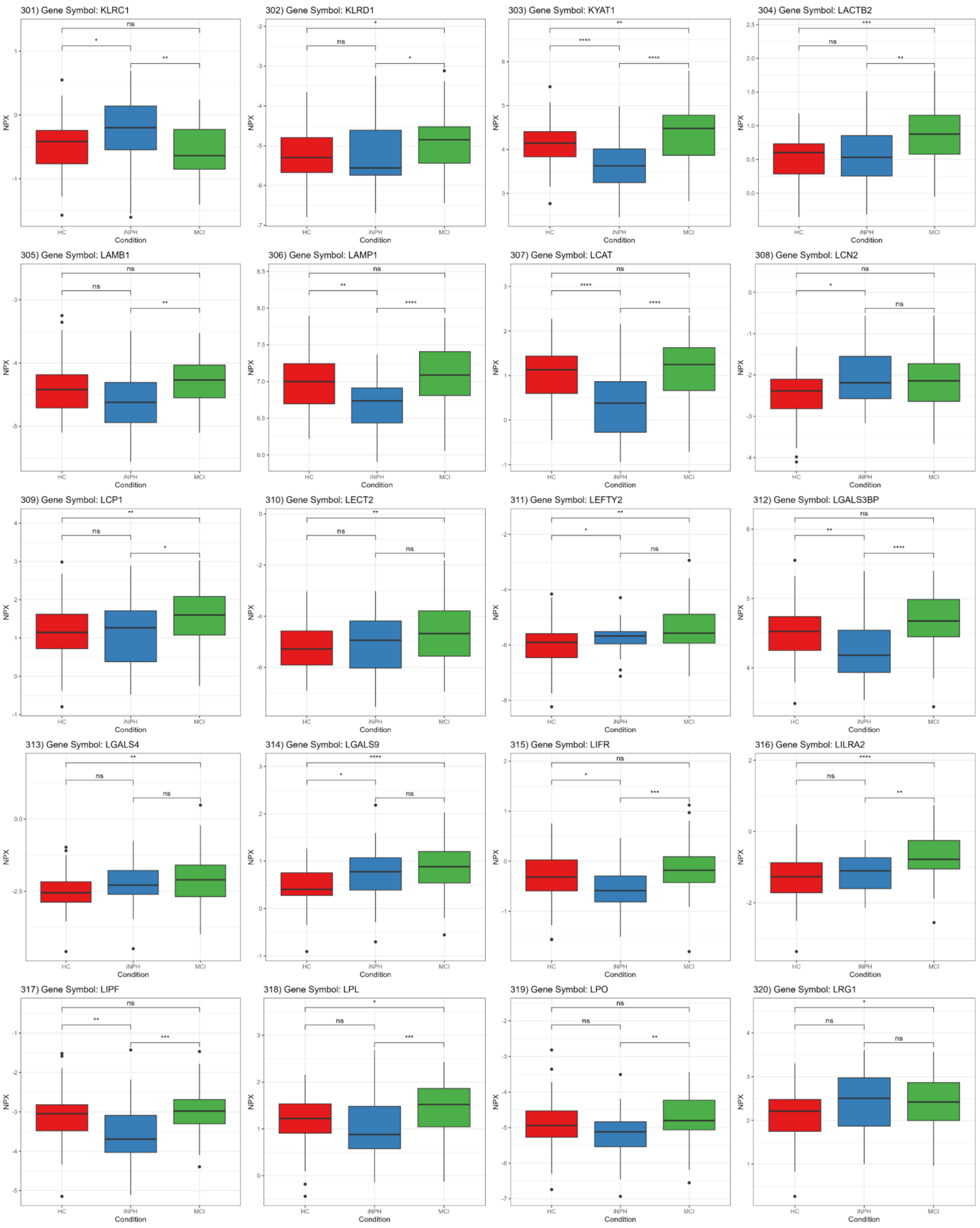
301) Gene Symbol: KLRC1
302) Gene Symbol: KLRD1
303) Gene Symbol: KYAT1
304) Gene Symbol: LACTB2
305) Gene Symbol: LAMB1
306) Gene Symbol: LAMP1
307) Gene Symbol: LCAT
308) Gene Symbol: LCN2
309) Gene Symbol: LCP1
310) Gene Symbol: LECT2
311) Gene Symbol: LEFTY2
312) Gene Symbol: LGALS3BP
313) Gene Symbol: LGALS4
314) Gene Symbol: LGALS9
315) Gene Symbol: LIFR
316) Gene Symbol: LILRA2
317) Gene Symbol: LIPF
318) Gene Symbol: LPL
319) Gene Symbol: LPO
320) Gene Symbol: LRG1
NPX
Condition
HC
iNPH
MCI

321) Gene Symbol: LRIG3

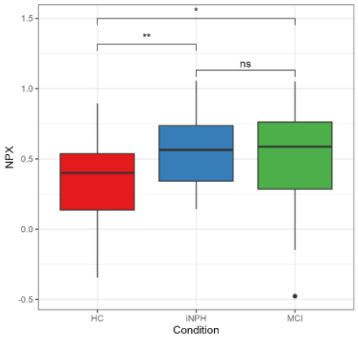


322) Gene Symbol: LRP1

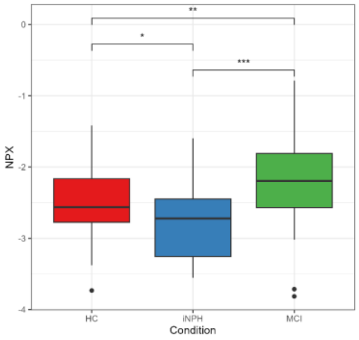


323) Gene Symbol: LRRC37A2

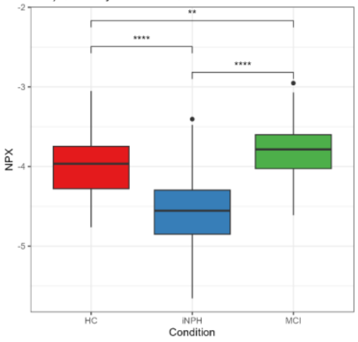


324) Gene Symbol: LRTM2

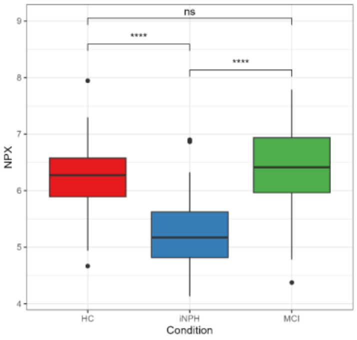


325) Gene Symbol: LTBP3

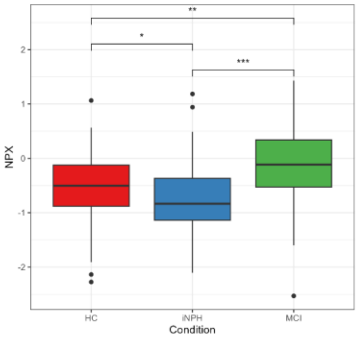


326) Gene Symbol: LUZP2

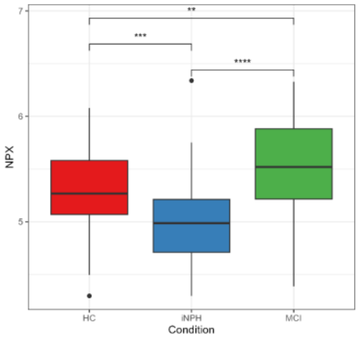


327) Gene Symbol: LY9

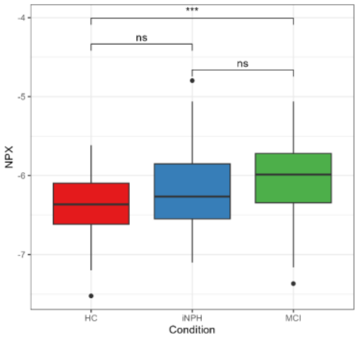


328) Gene Symbol: LYPD8

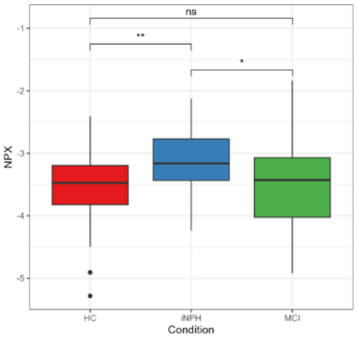


329) Gene Symbol: LYVE1

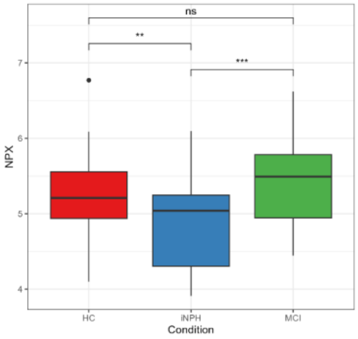


330) Gene Symbol: MAMDC2

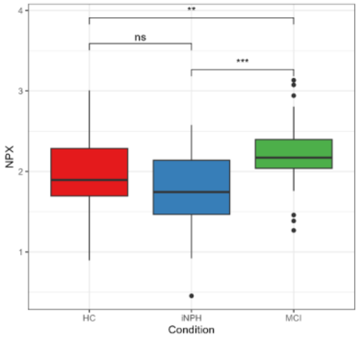


331) Gene Symbol: MAN1A2

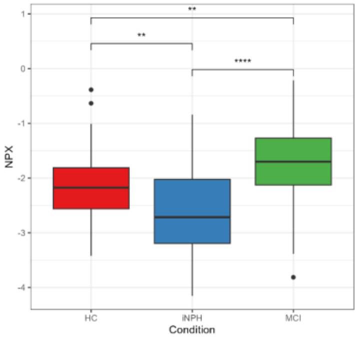


332) Gene Symbol: MAN2B2

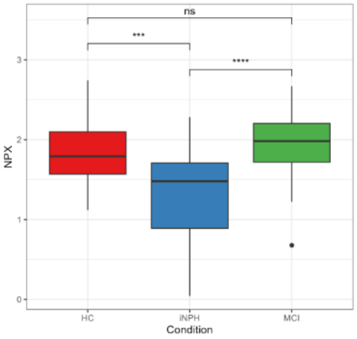


333) Gene Symbol: MANEAL

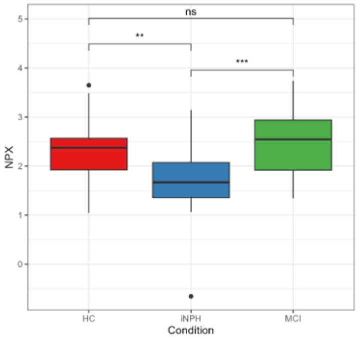


334) Gene Symbol: MANF

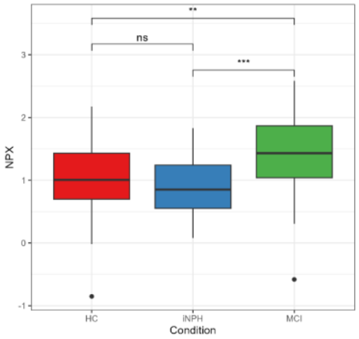


335) Gene Symbol: MAP1LC3A

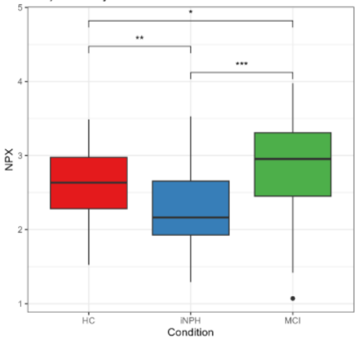


336) Gene Symbol: MAP1LC3B2

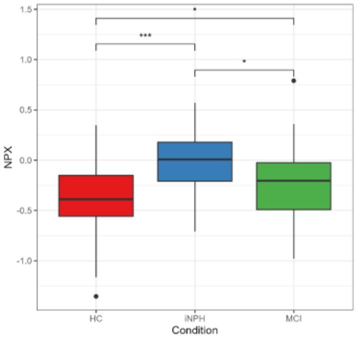


337) Gene Symbol: MAP2

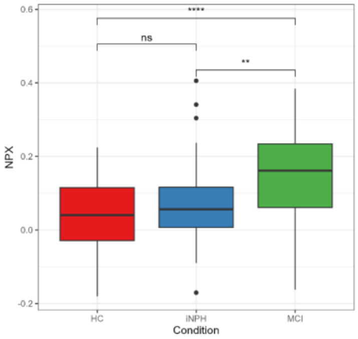


338) Gene Symbol: MAP4K5

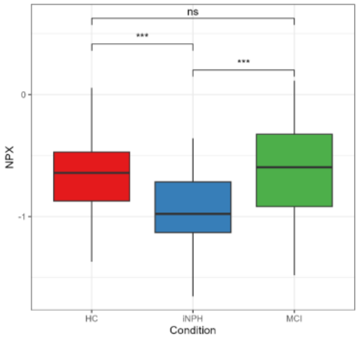


339) Gene Symbol: MAPT

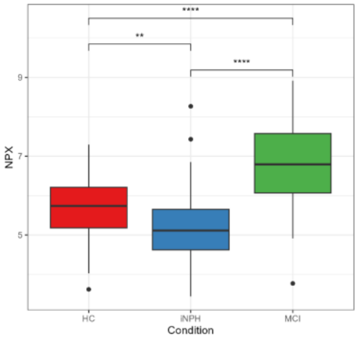


340) Gene Symbol: MARS1

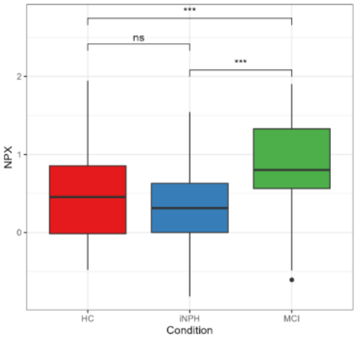

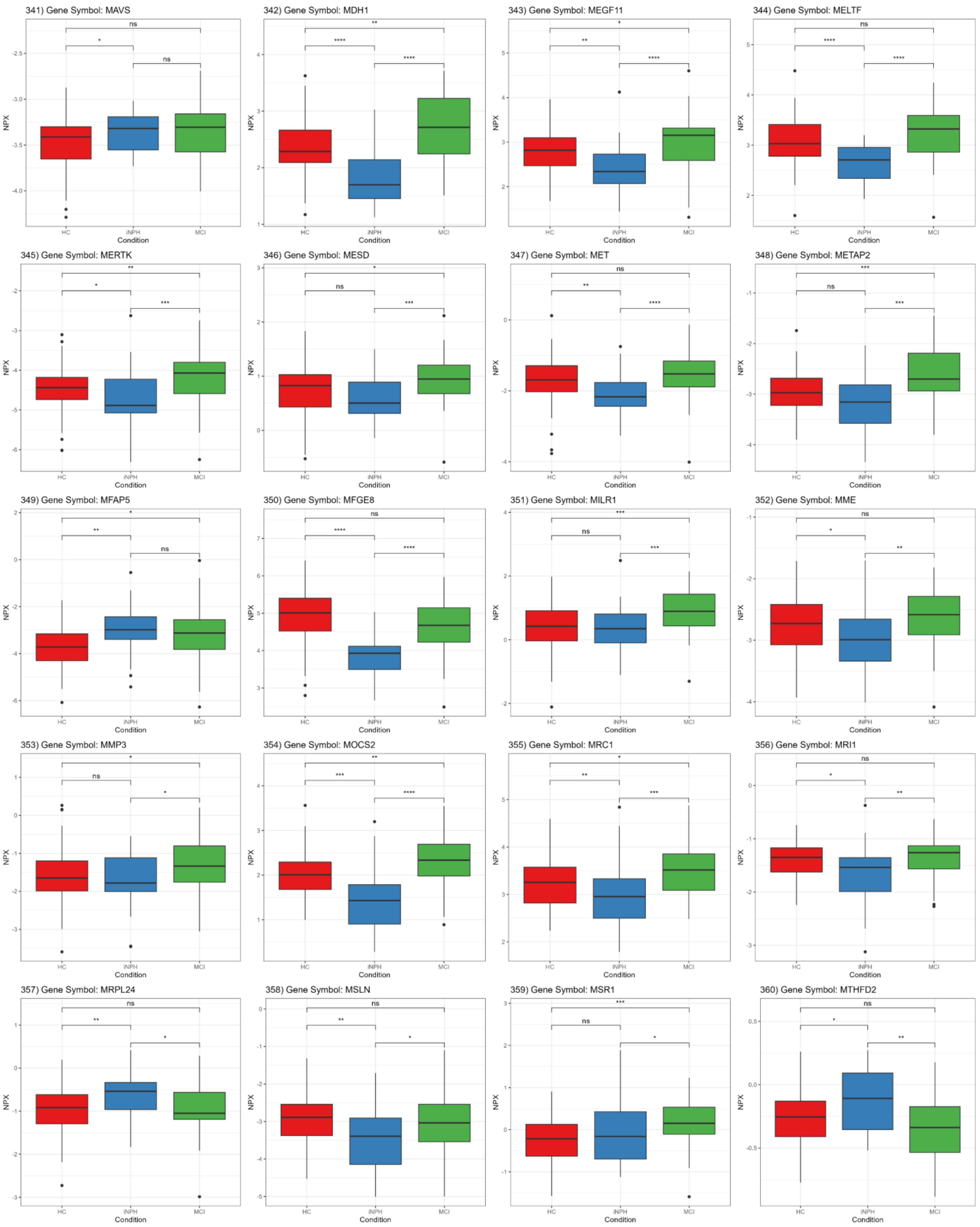
341) Gene Symbol: MAVS
342) Gene Symbol: MDH1
343) Gene Symbol: MEGF11
344) Gene Symbol: MELTF
345) Gene Symbol: MERTK
346) Gene Symbol: MESD
347) Gene Symbol: MET
348) Gene Symbol: METAP2
349) Gene Symbol: MFAP5
350) Gene Symbol: MFGE8
351) Gene Symbol: MILR1
352) Gene Symbol: MME
353) Gene Symbol: MMP3
354) Gene Symbol: MOCS2
355) Gene Symbol: MRC1
356) Gene Symbol: MRI1
357) Gene Symbol: MRPL24
358) Gene Symbol: MSLN
359) Gene Symbol: MSR1
360) Gene Symbol: MTHFD2
NPX
Condition
HC
iNPH
MCI

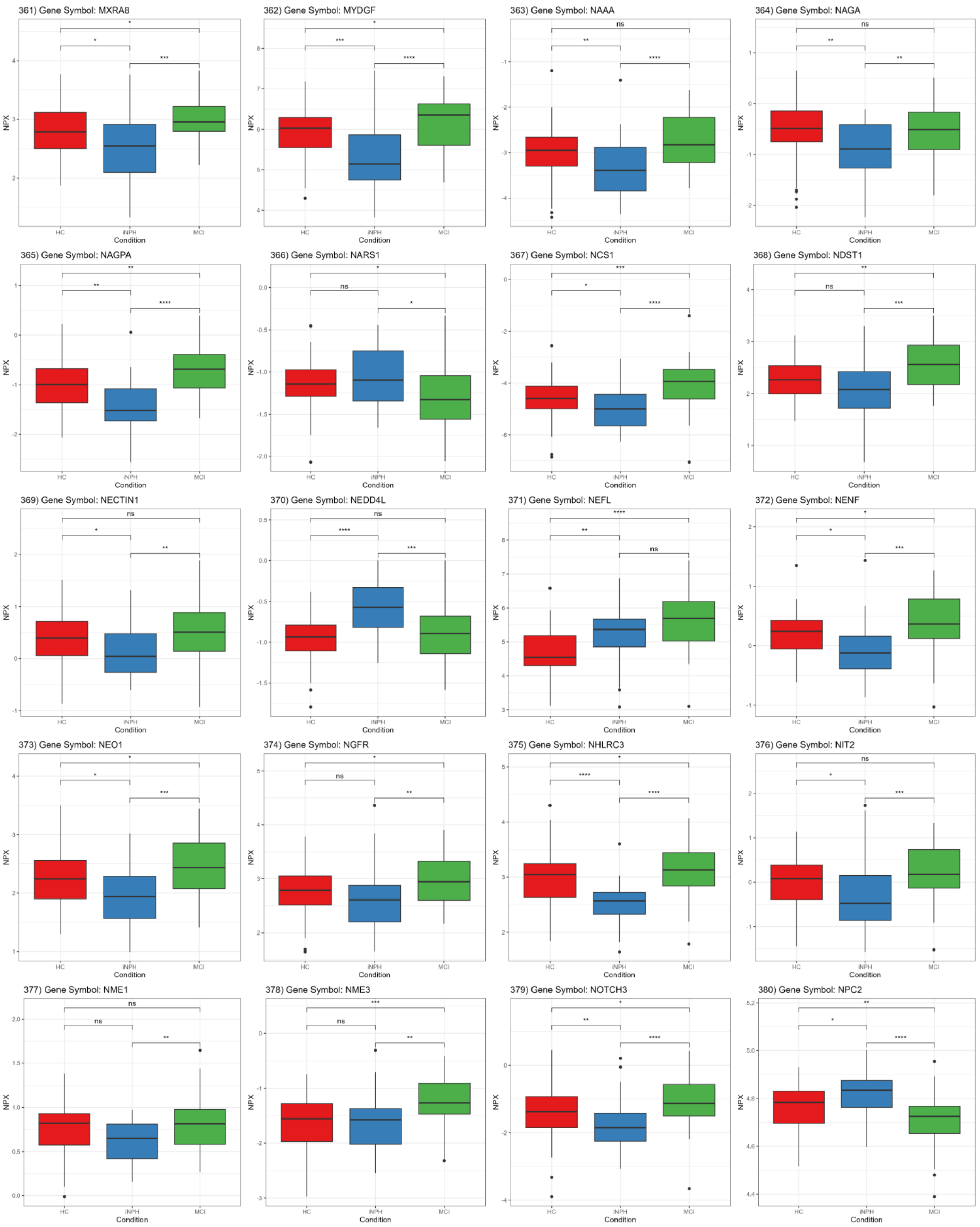
361) Gene Symbol: MXRA8
362) Gene Symbol: MYDGF
363) Gene Symbol: NAAA
364) Gene Symbol: NAGA
365) Gene Symbol: NAGPA
366) Gene Symbol: NARS1
367) Gene Symbol: NCS1
368) Gene Symbol: NDST1
369) Gene Symbol: NECTIN1
370) Gene Symbol: NEDD4L
371) Gene Symbol: NEFL
372) Gene Symbol: NENF
373) Gene Symbol: NEO1
374) Gene Symbol: NGFR
375) Gene Symbol: NHLRC3
376) Gene Symbol: NIT2
377) Gene Symbol: NME1
378) Gene Symbol: NME3
379) Gene Symbol: NOTCH3
380) Gene Symbol: NPC2
NPX
Condition
HC
iNPH
MCI

381) Gene Symbol: NPTN

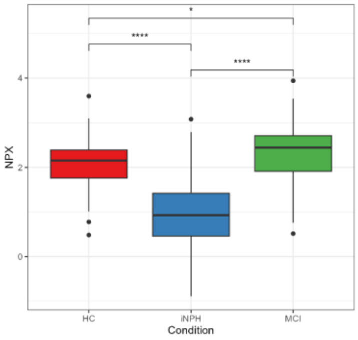


382) Gene Symbol: NPTXR

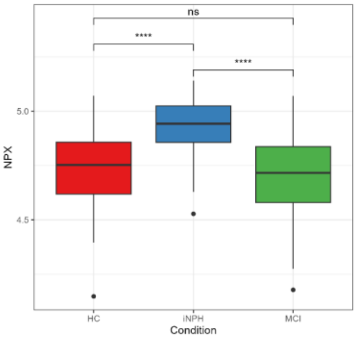


383) Gene Symbol: NRCAM

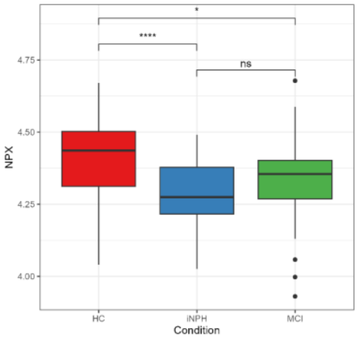


384) Gene Symbol: NRGN

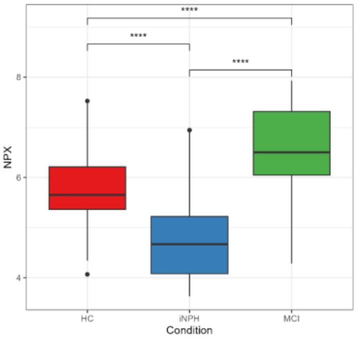


385) Gene Symbol: NT5C

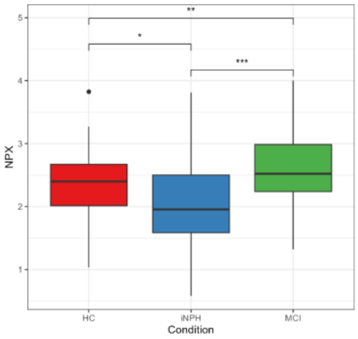


386) Gene Symbol: NUDC

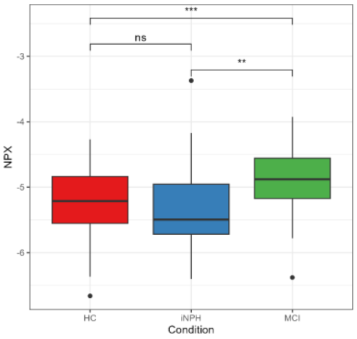


387) Gene Symbol: OCLN

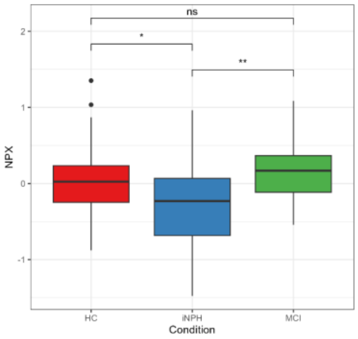


388) Gene Symbol: OSMR

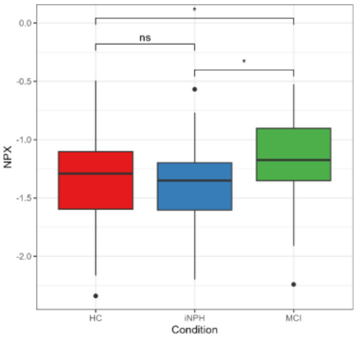


389) Gene Symbol: OTUD7B

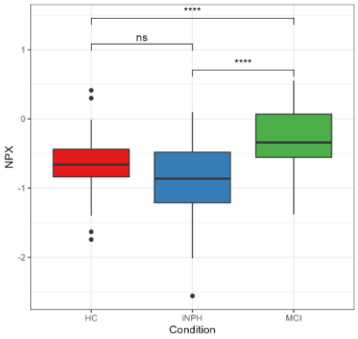


390) Gene Symbol: PAFAH1B3

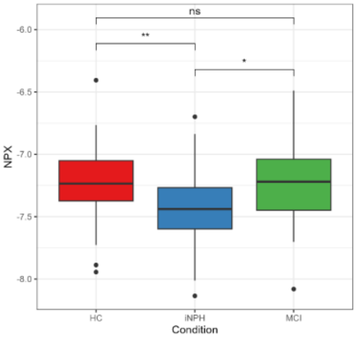


391) Gene Symbol: PALM2

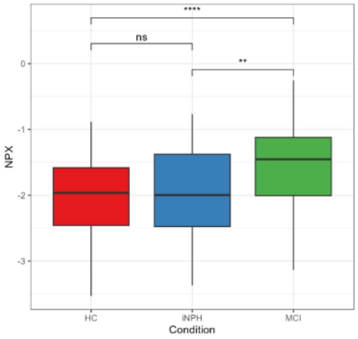


392) Gene Symbol: PARK7

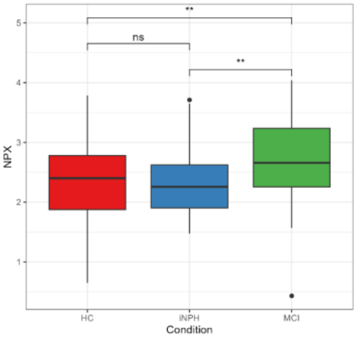


393) Gene Symbol: PAXX

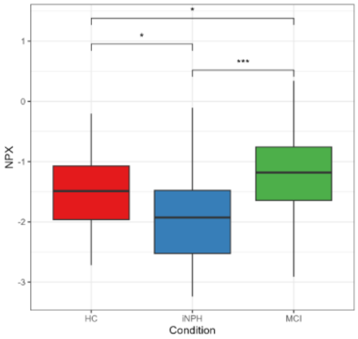


394) Gene Symbol: PBK

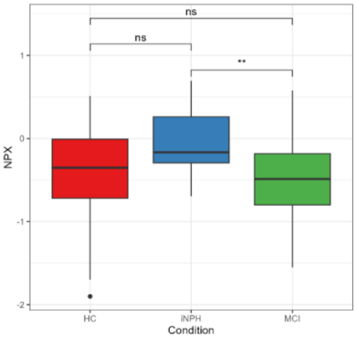


395) Gene Symbol: PCBD1

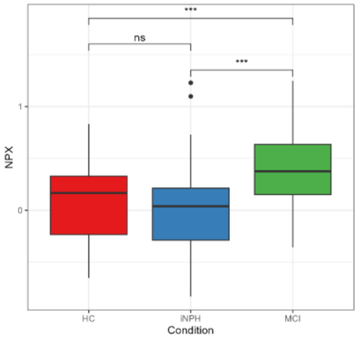


396) Gene Symbol: PCDH12

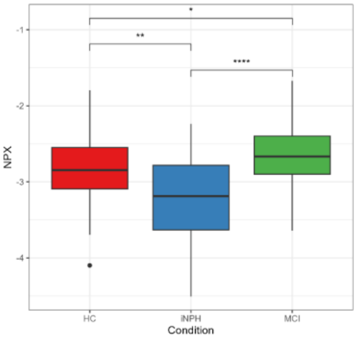


397) Gene Symbol: PCDH7

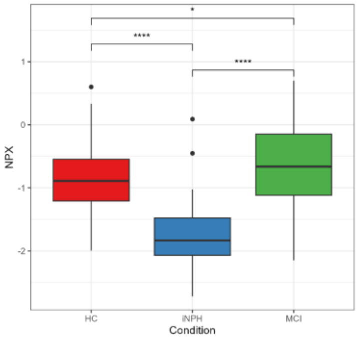


398) Gene Symbol: PCDH9

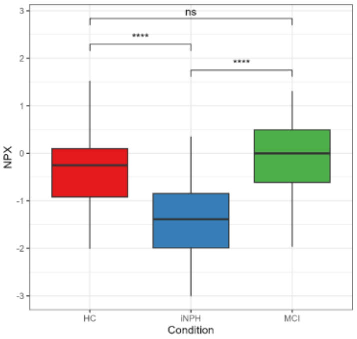


399) Gene Symbol: PCDHB15

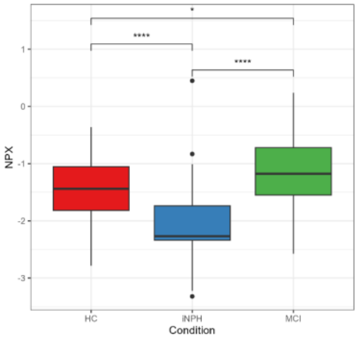


400) Gene Symbol: PCSK9

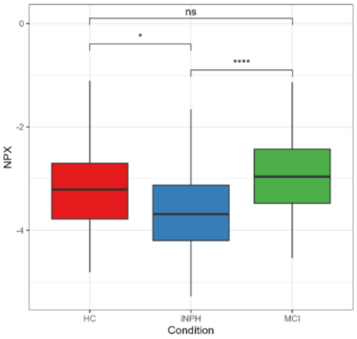

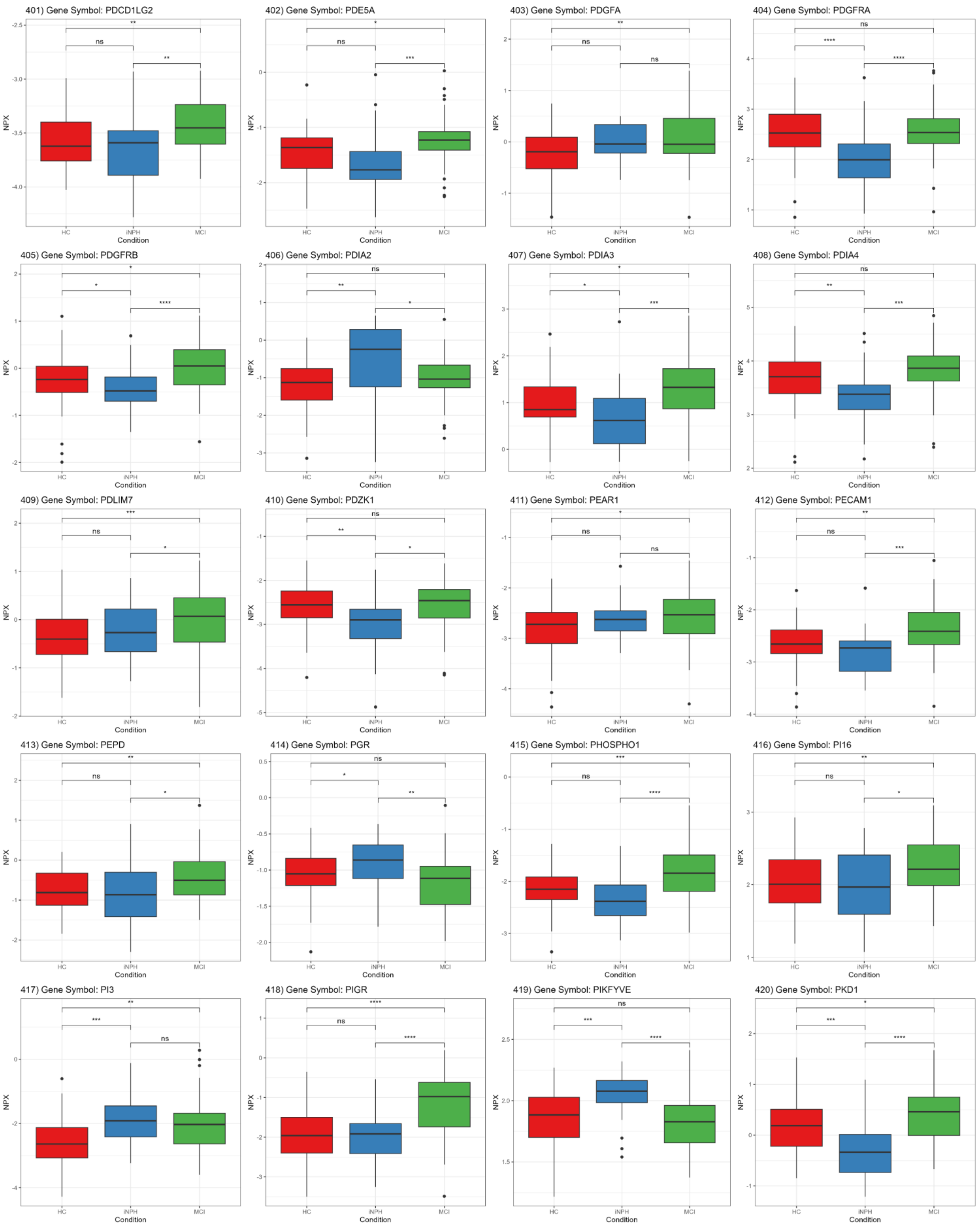
401) Gene Symbol: PDCD1LG2
402) Gene Symbol: PDE5A
403) Gene Symbol: PDGFA
404) Gene Symbol: PDGFRA
405) Gene Symbol: PDGFRB
406) Gene Symbol: PDIA2
407) Gene Symbol: PDIA3
408) Gene Symbol: PDIA4
409) Gene Symbol: PDLIM7
410) Gene Symbol: PDZK1
411) Gene Symbol: PEAR1
412) Gene Symbol: PECAM1
413) Gene Symbol: PEPD
414) Gene Symbol: PGR
415) Gene Symbol: PHOSPHO1
416) Gene Symbol: PI16
417) Gene Symbol: PI3
418) Gene Symbol: PIGR
419) Gene Symbol: PIKFYVE
420) Gene Symbol: PKD1
NPX
Condition
HC
iNPH
MCI

421) Gene Symbol: PLA2G15

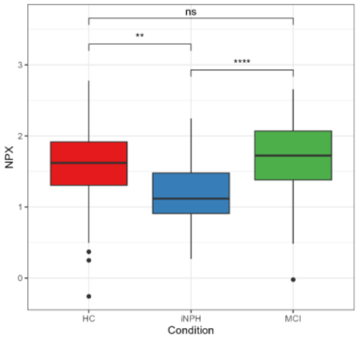


422) Gene Symbol: PLA2G7

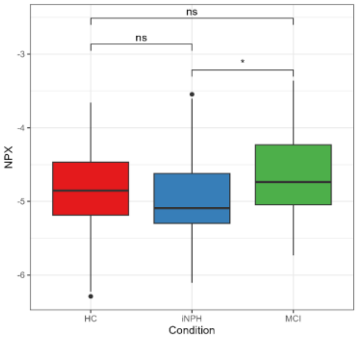


423) Gene Symbol: PLB1

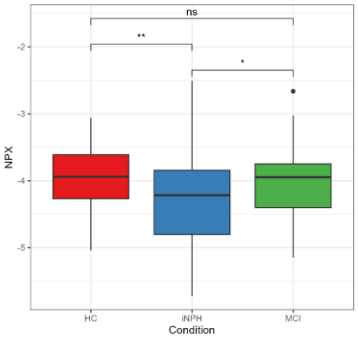


424) Gene Symbol: PLCB1

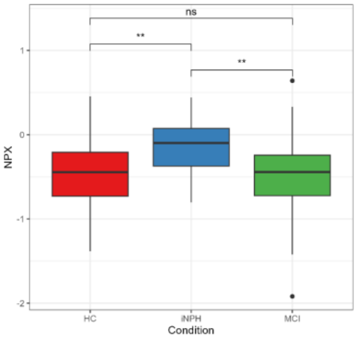


425) Gene Symbol: PLIN3

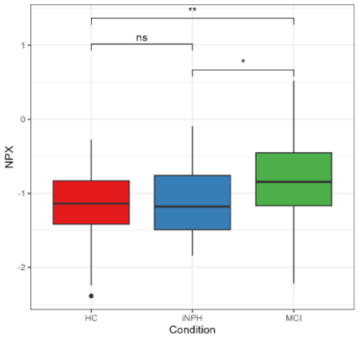


426) Gene Symbol: PLSCR3

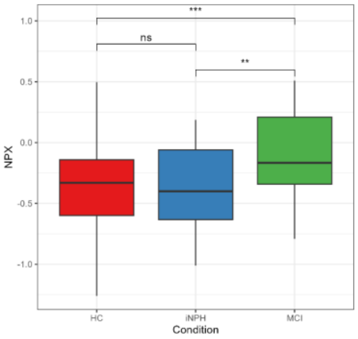


427) Gene Symbol: PLXDC1

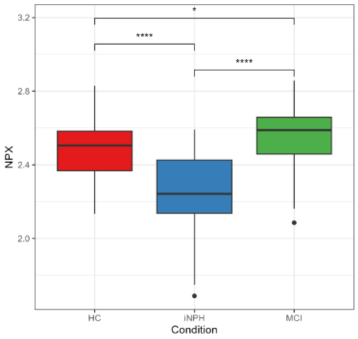


428) Gene Symbol: PLXNA4

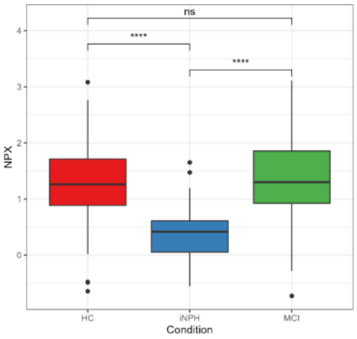


429) Gene Symbol: PMCH

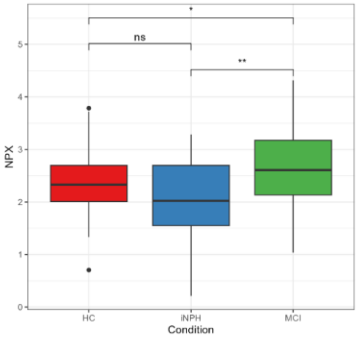


430) Gene Symbol: PODXL

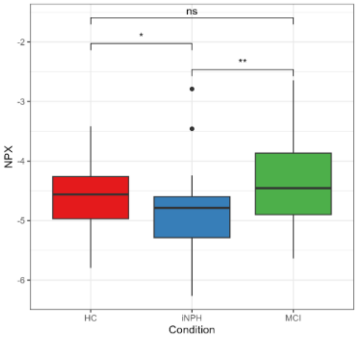


431) Gene Symbol: POMC

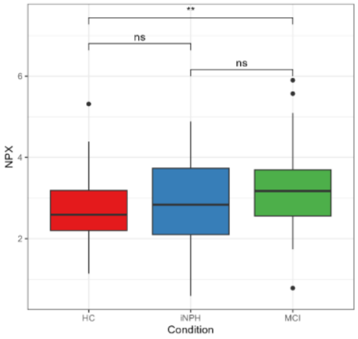


432) Gene Symbol: PON2

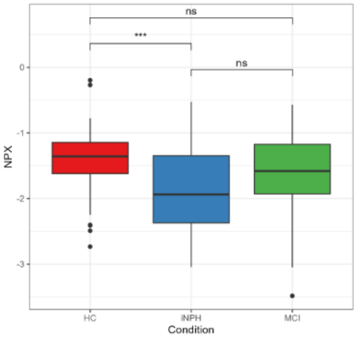


433) Gene Symbol: PPL

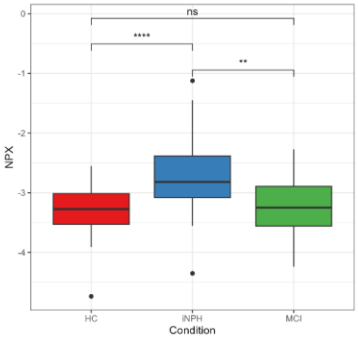


434) Gene Symbol: PPM1F

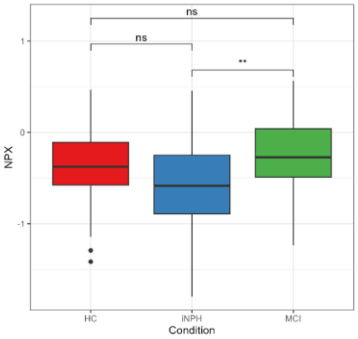


435) Gene Symbol: PPP1R2

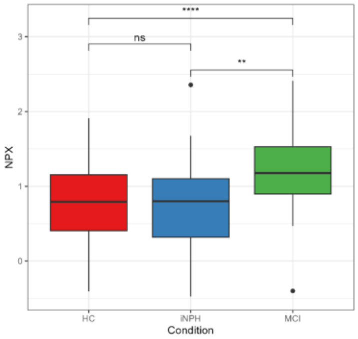


436) Gene Symbol: PQBP1

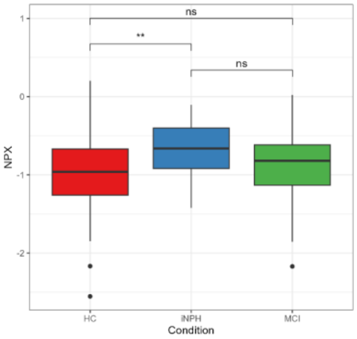


437) Gene Symbol: PRDX5

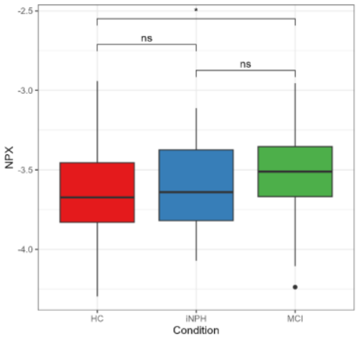


438) Gene Symbol: PRKAR1A

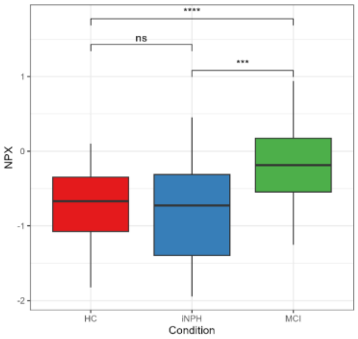


439) Gene Symbol: PRKAR2A

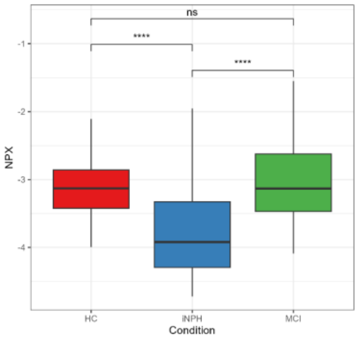


440) Gene Symbol: PROCR

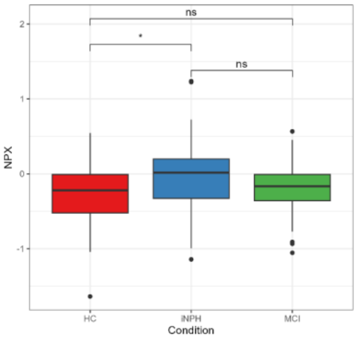

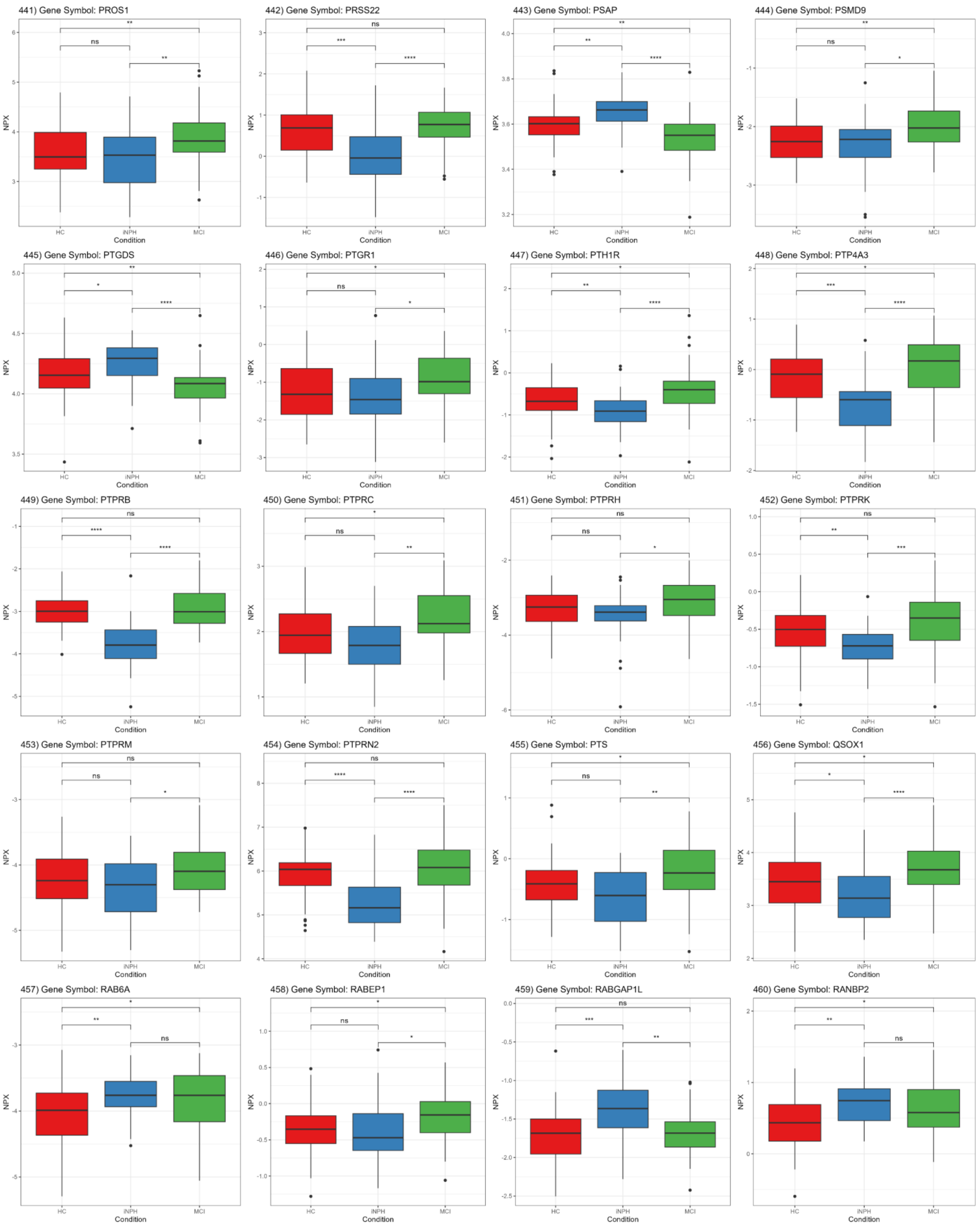
441) Gene Symbol: PROS1
442) Gene Symbol: PRSS22
443) Gene Symbol: PSAP
444) Gene Symbol: PSMD9
445) Gene Symbol: PTGDS
446) Gene Symbol: PTGR1
447) Gene Symbol: PTH1R
448) Gene Symbol: PTP4A3
449) Gene Symbol: PTPRB
450) Gene Symbol: PTPRC
451) Gene Symbol: PTPRH
452) Gene Symbol: PTPRK
453) Gene Symbol: PTPRM
454) Gene Symbol: PTPRN2
455) Gene Symbol: PTS
456) Gene Symbol: QSOX1
457) Gene Symbol: RAB6A
458) Gene Symbol: RABEP1
459) Gene Symbol: RABGAP1L
460) Gene Symbol: RANBP2
NPX
Condition
HC
iNPH
MCI

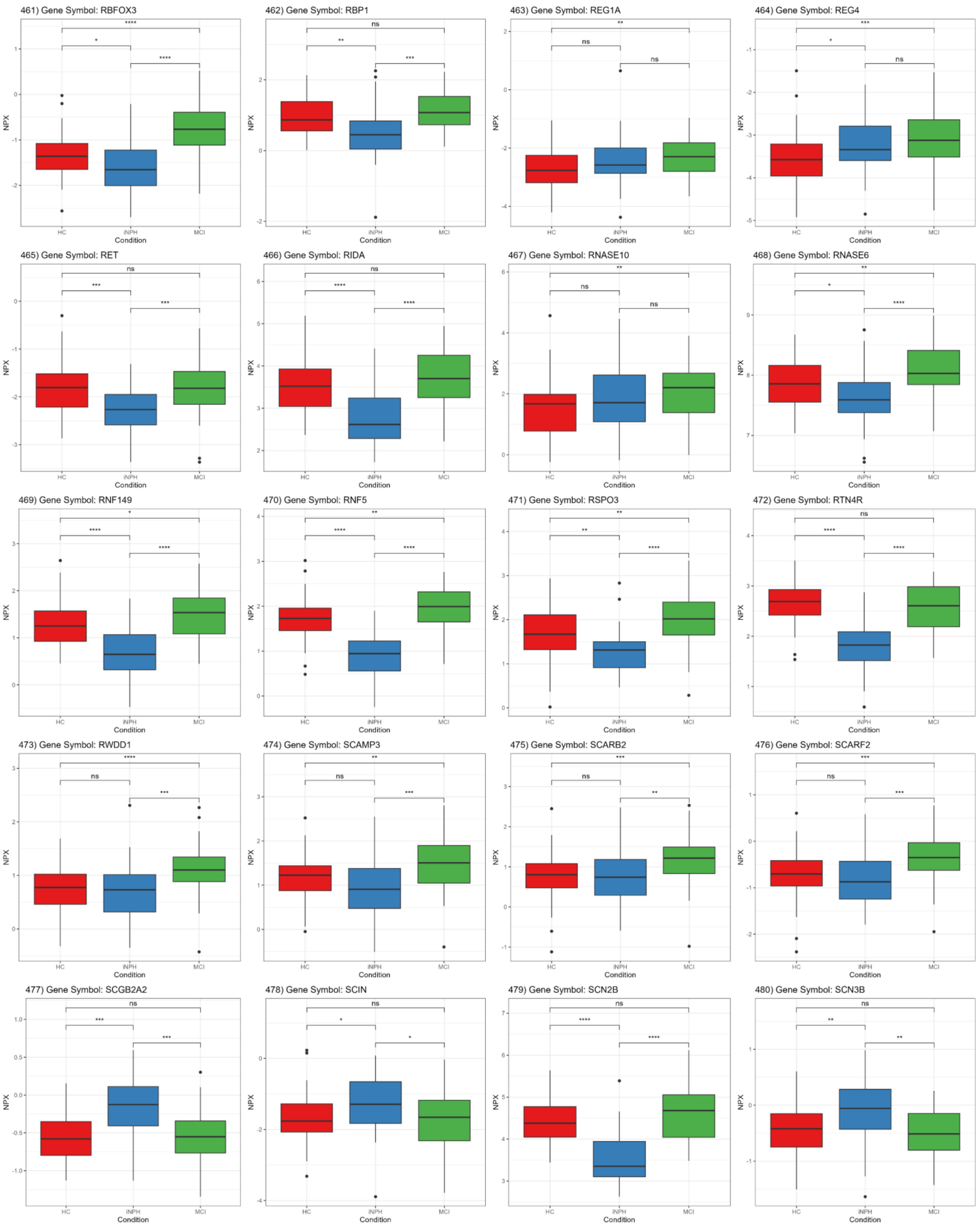
461) Gene Symbol: RBFOX3
462) Gene Symbol: RBP1
463) Gene Symbol: REG1A
464) Gene Symbol: REG4
465) Gene Symbol: RET
466) Gene Symbol: RIDA
467) Gene Symbol: RNASE10
468) Gene Symbol: RNASE6
469) Gene Symbol: RNF149
470) Gene Symbol: RNF5
471) Gene Symbol: RSPO3
472) Gene Symbol: RTN4R
473) Gene Symbol: RWDD1
474) Gene Symbol: SCAMP3
475) Gene Symbol: SCARB2
476) Gene Symbol: SCARF2
477) Gene Symbol: SCGB2A2
478) Gene Symbol: SCIN
479) Gene Symbol: SCN2B
480) Gene Symbol: SCN3B
NPX
Condition
HC
iNPH
MCI

481) Gene Symbol: SCN4B

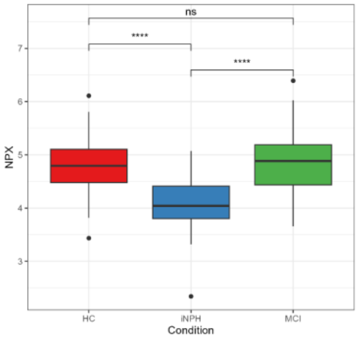


482) Gene Symbol: SCRN1

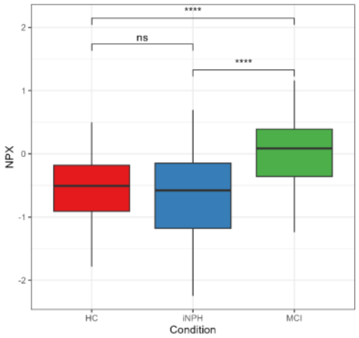


483) Gene Symbol: SDC1

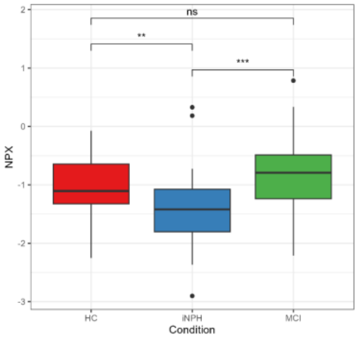


484) Gene Symbol: SDC4

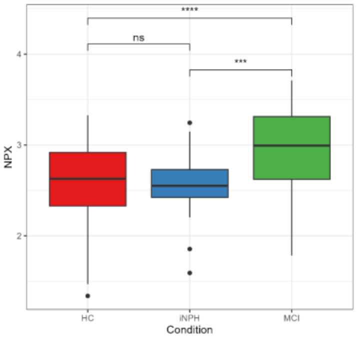


485) Gene Symbol: SDHB

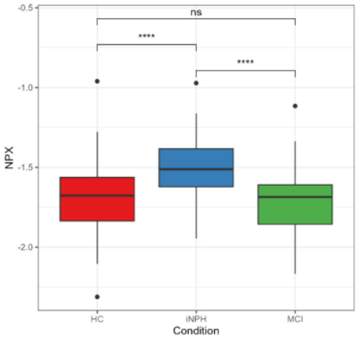


486) Gene Symbol: SDK2

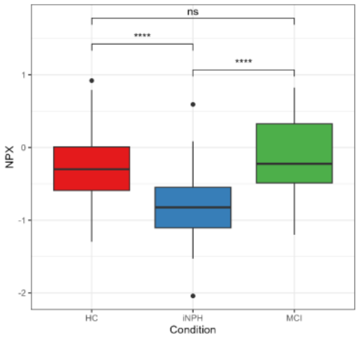


487) Gene Symbol: SEL1L

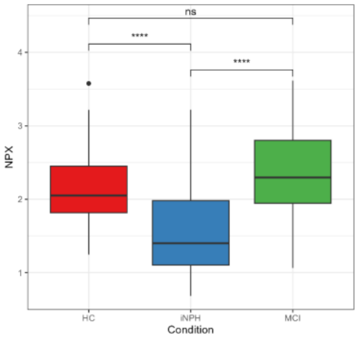


488) Gene Symbol: SELENOP

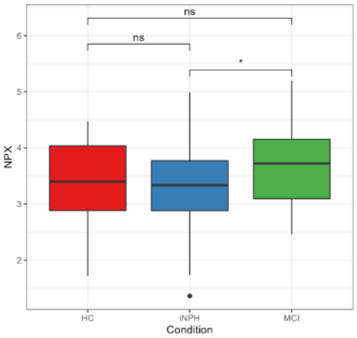


489) Gene Symbol: SEMA3G

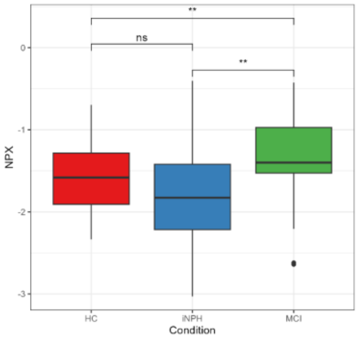


490) Gene Symbol: SEPTIN8

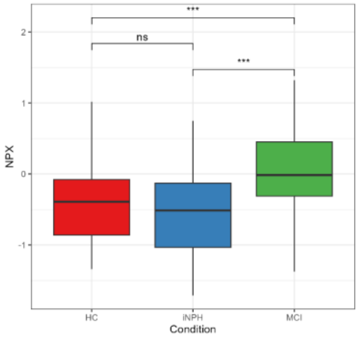


491) Gene Symbol: SEPTIN9

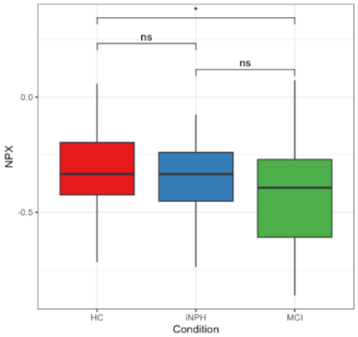


492) Gene Symbol: SERPINA9

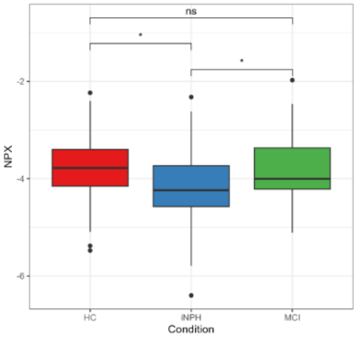


493) Gene Symbol: SERPINB1

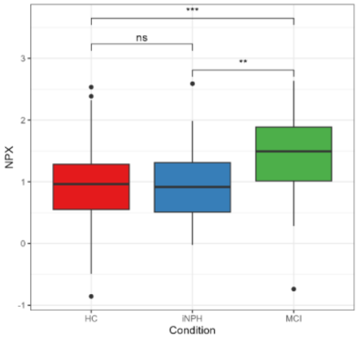


494) Gene Symbol: SERPINB8

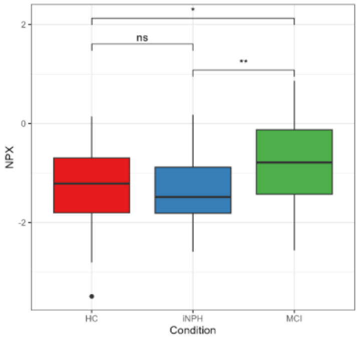


495) Gene Symbol: SERPINB9

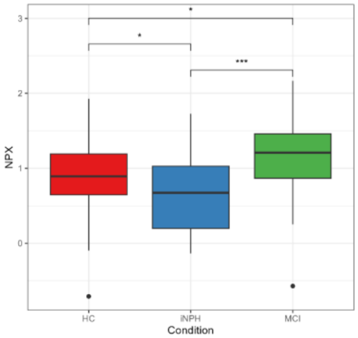


496) Gene Symbol: SERPIND1

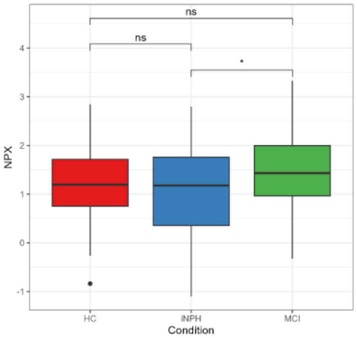


497) Gene Symbol: SERPINF1

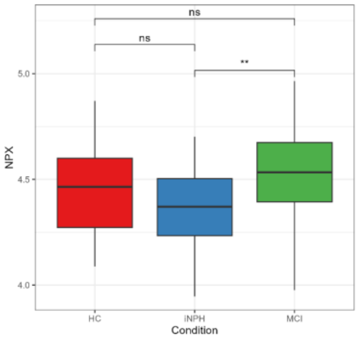


498) Gene Symbol: SERPING1

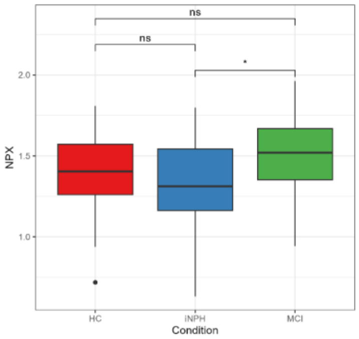


499) Gene Symbol: SERPINI1

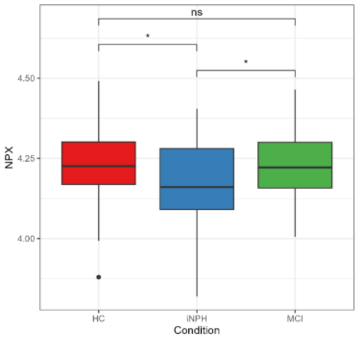


500) Gene Symbol: SESTD1

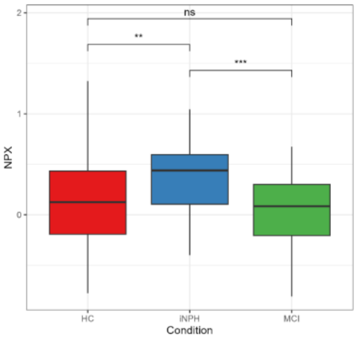

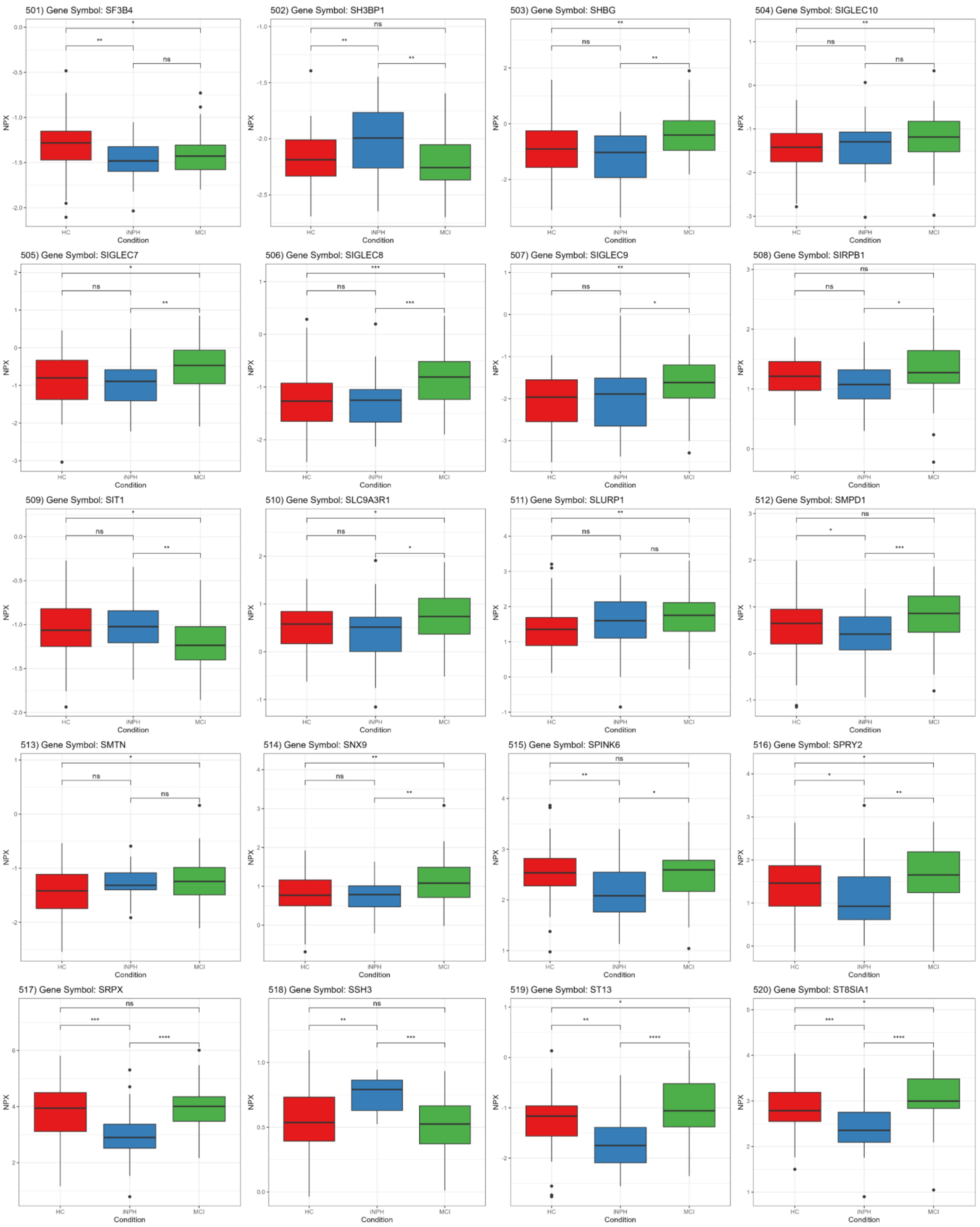
501) Gene Symbol: SF3B4
502) Gene Symbol: SH3BP1
503) Gene Symbol: SHBG
504) Gene Symbol: SIGLEC10
505) Gene Symbol: SIGLEC7
506) Gene Symbol: SIGLEC8
507) Gene Symbol: SIGLEC9
508) Gene Symbol: SIRPB1
509) Gene Symbol: SIT1
510) Gene Symbol: SLC9A3R1
511) Gene Symbol: SLURP1
512) Gene Symbol: SMPD1
513) Gene Symbol: SMTN
514) Gene Symbol: SNX9
515) Gene Symbol: SPINK6
516) Gene Symbol: SPRY2
517) Gene Symbol: SRPX
518) Gene Symbol: SSH3
519) Gene Symbol: ST13
520) Gene Symbol: ST8SIA1
NPX
Condition
HC
iNPH
MCI

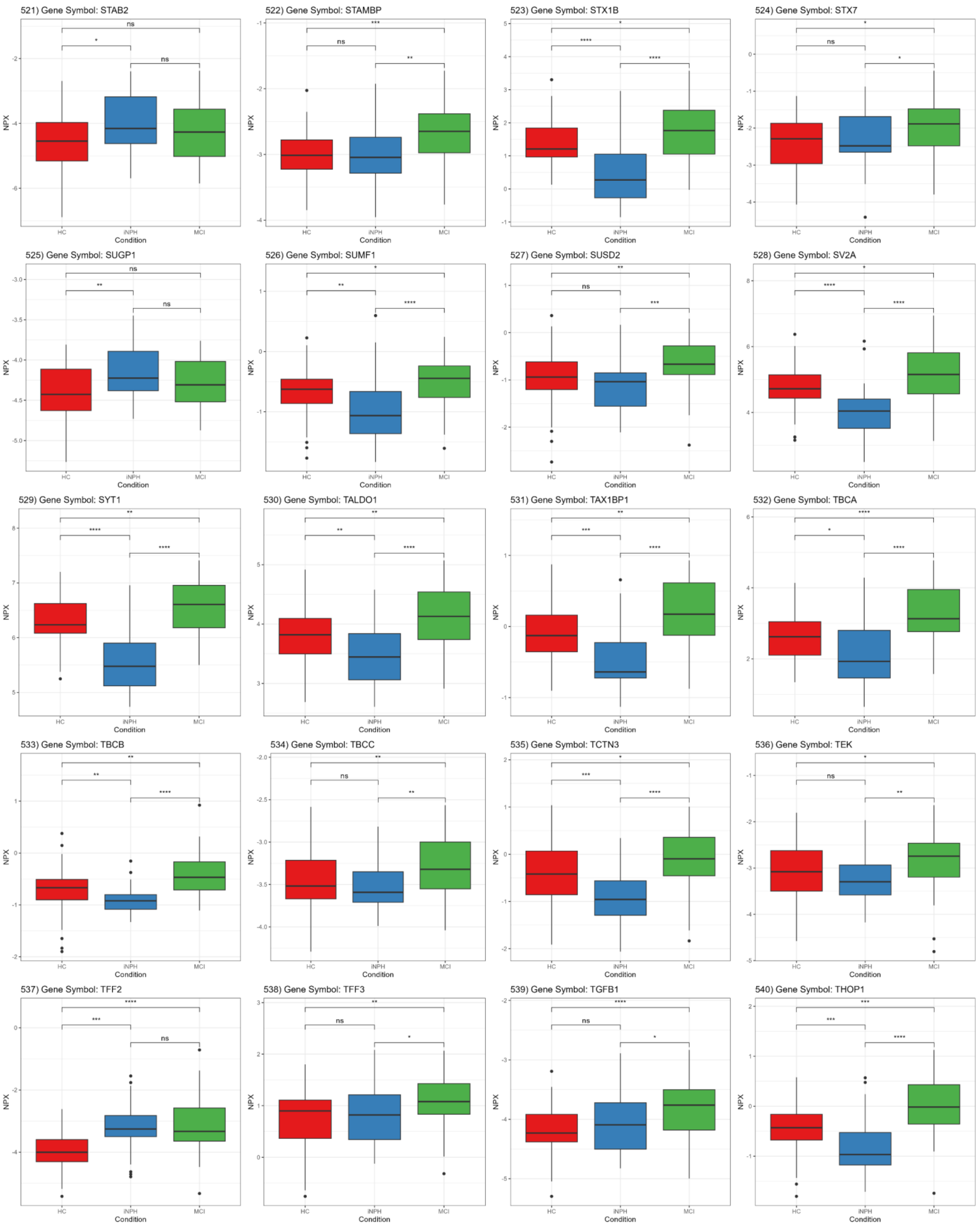
521) Gene Symbol: STAB2
522) Gene Symbol: STAMBP
523) Gene Symbol: STX1B
524) Gene Symbol: STX7
525) Gene Symbol: SUGP1
526) Gene Symbol: SUMF1
527) Gene Symbol: SUSD2
528) Gene Symbol: SV2A
529) Gene Symbol: SYT1
530) Gene Symbol: TALDO1
531) Gene Symbol: TAX1BP1
532) Gene Symbol: TBCA
533) Gene Symbol: TBCB
534) Gene Symbol: TBCC
535) Gene Symbol: TCTN3
536) Gene Symbol: TEK
537) Gene Symbol: TFF2
538) Gene Symbol: TFF3
539) Gene Symbol: TGFB1
540) Gene Symbol: THOP1
NPX
HC
iNPH
MCI
Condition

541) Gene Symbol: TLR2

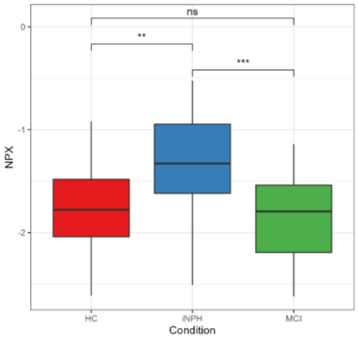


542) Gene Symbol: TMEM25

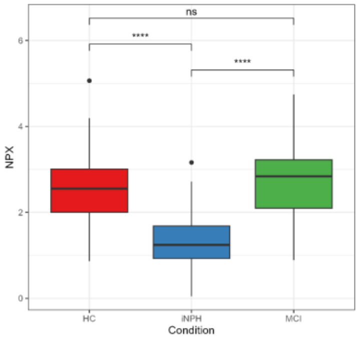


543) Gene Symbol: TNFRSF10A

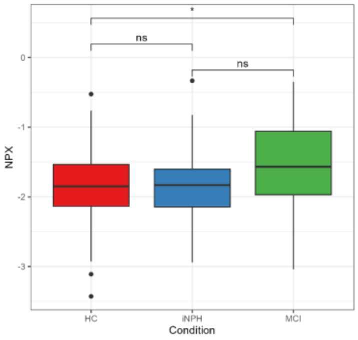


544) Gene Symbol: TNFRSF10C

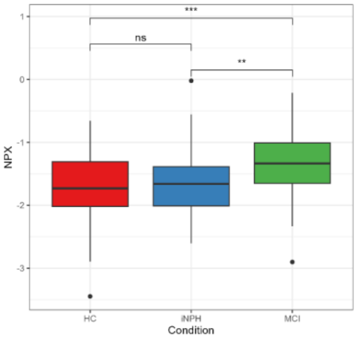


545) Gene Symbol: TNFRSF11A

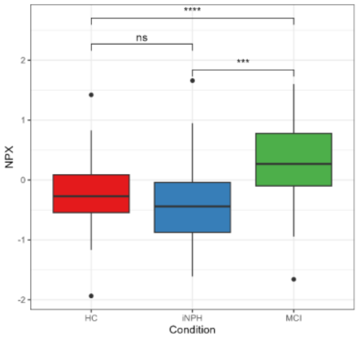


546) Gene Symbol: TNFRSF11B

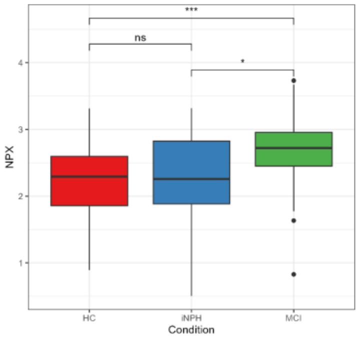


547) Gene Symbol: TNFRSF14

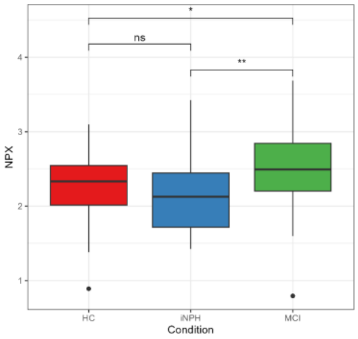


548) Gene Symbol: TNFRSF1B

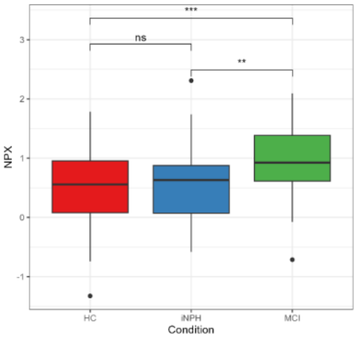


549) Gene Symbol: TNFSF10

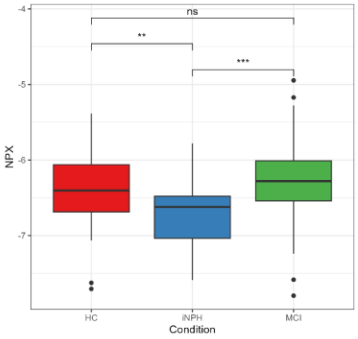


550) Gene Symbol: TNFSF12

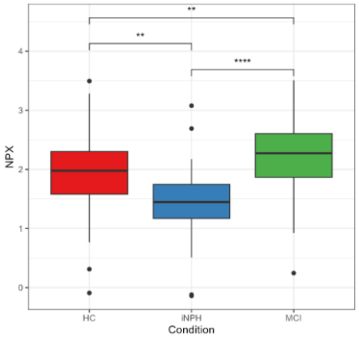


551) Gene Symbol: TNFSF13B

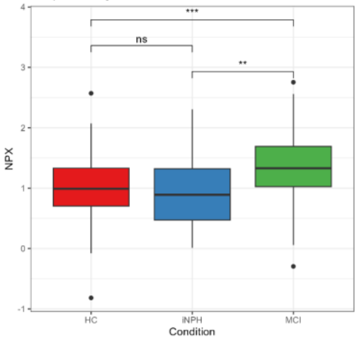


552) Gene Symbol: TNFSF8

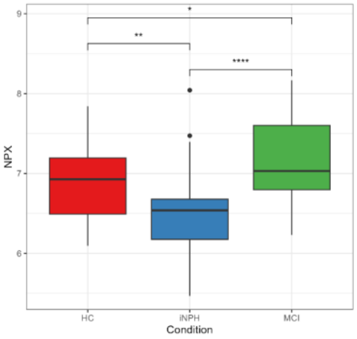


553) Gene Symbol: TNFSF9

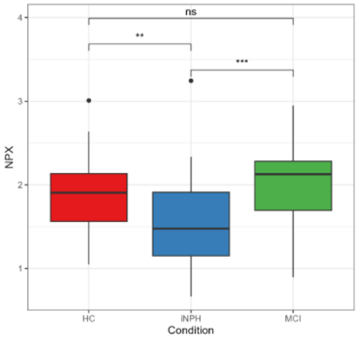


554) Gene Symbol: TNR

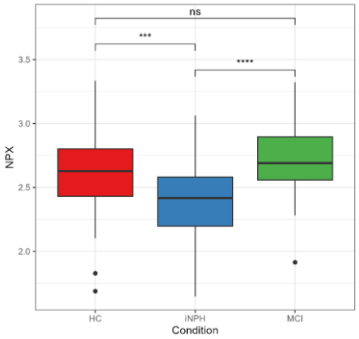


555) Gene Symbol: TOMM20

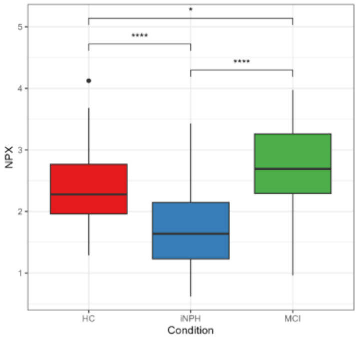


556) Gene Symbol: TPD52L2

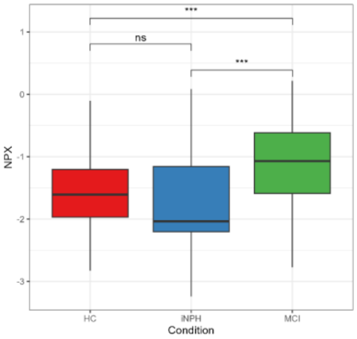


557) Gene Symbol: TPK1

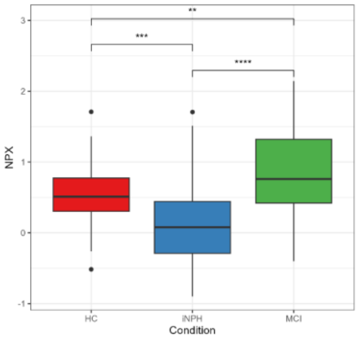


558) Gene Symbol: TPSG1

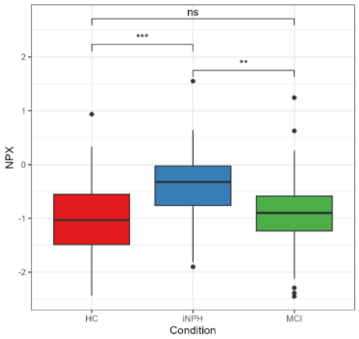


559) Gene Symbol: TREM2

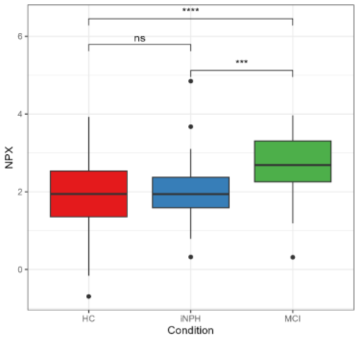


560) Gene Symbol: TSPAN1

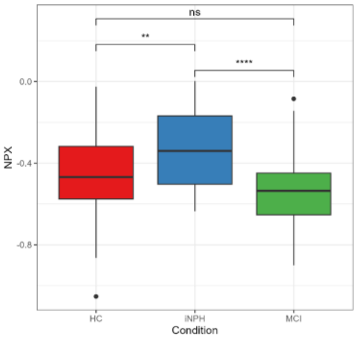

561) Gene Symbol: TTN

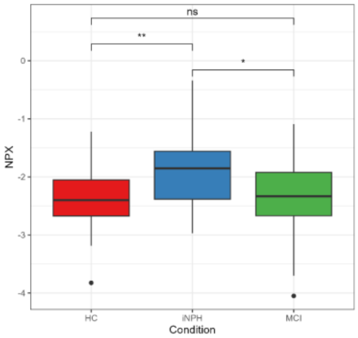


562) Gene Symbol: TWF2

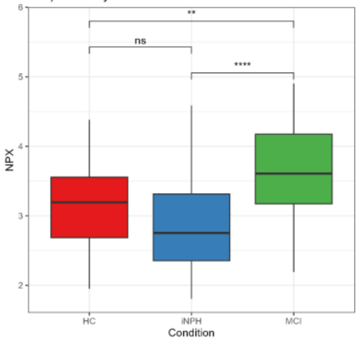


563) Gene Symbol: TXLNA

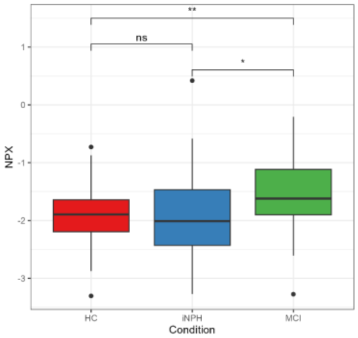


564) Gene Symbol: TXN

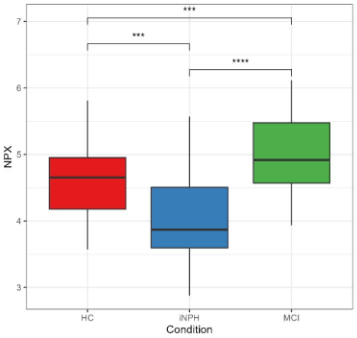


565) Gene Symbol: TXNDC15

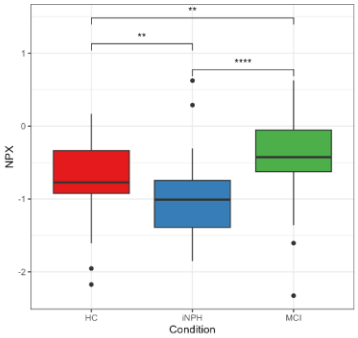


566) Gene Symbol: TXNDC5

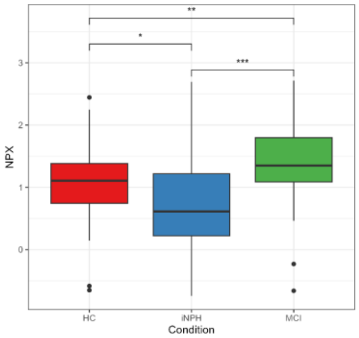


567) Gene Symbol: TXNRD1

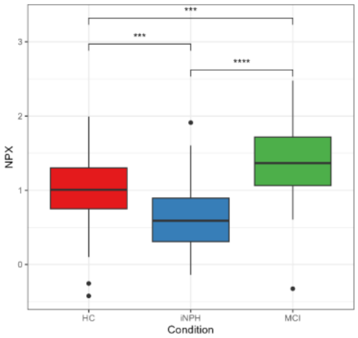


568) Gene Symbol: UBAC1

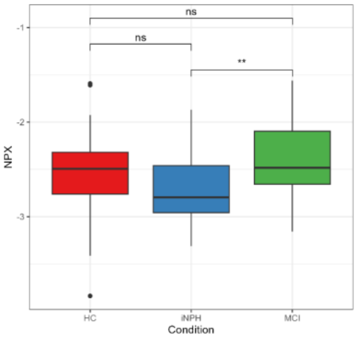


569) Gene Symbol: UBE2Z

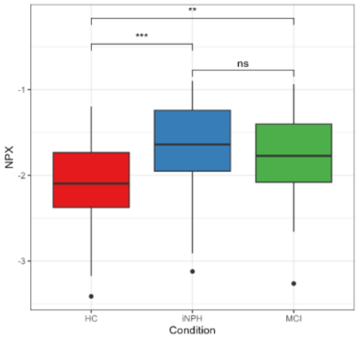


570) Gene Symbol: UBXN1

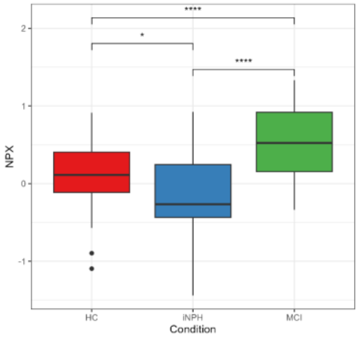


571) Gene Symbol: ULBP2

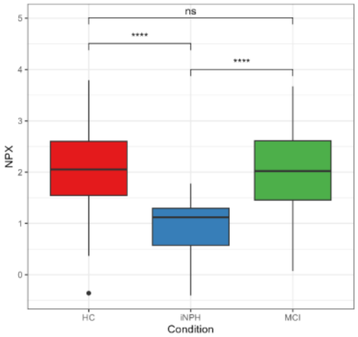


572) Gene Symbol: UROD

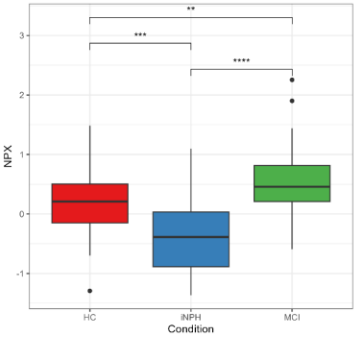


573) Gene Symbol: VASN

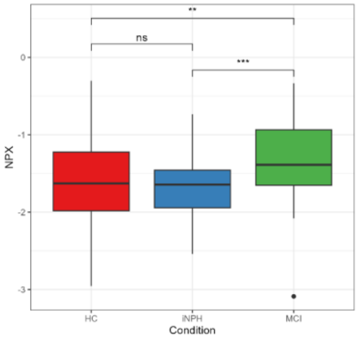


574) Gene Symbol: VAV3

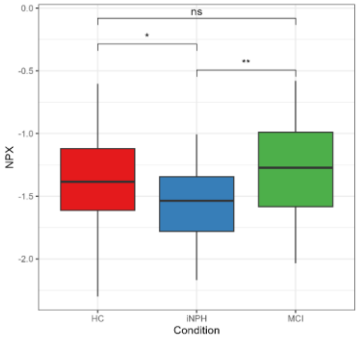


575) Gene Symbol: VEGFA

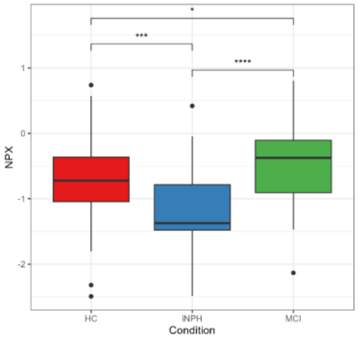


576) Gene Symbol: VIPR1

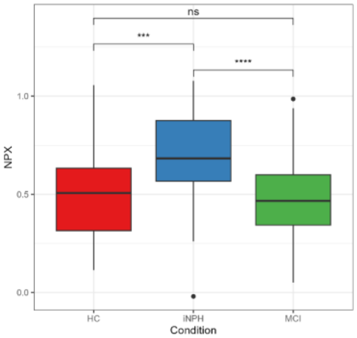


577) Gene Symbol: VNN2

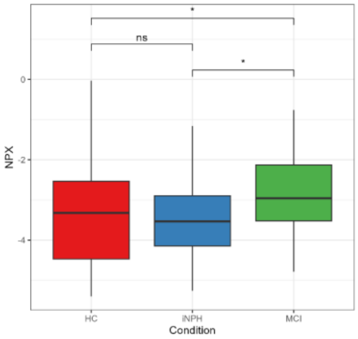


578) Gene Symbol: VPS28

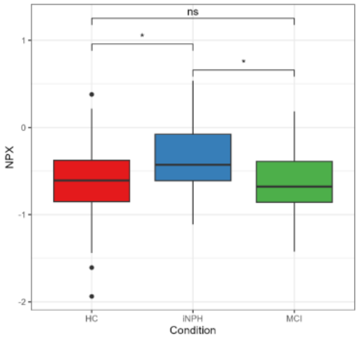


579) Gene Symbol: VPS53

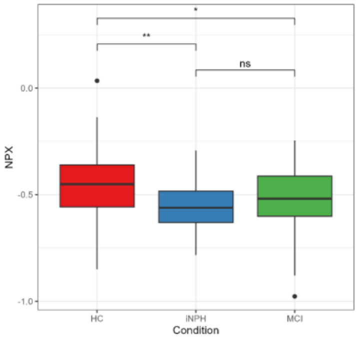


580) Gene Symbol: VSIG4

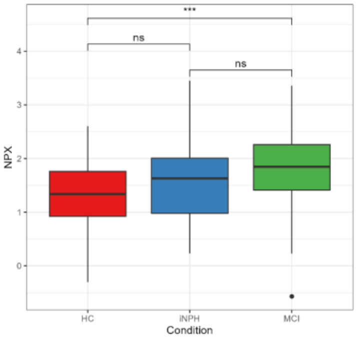

**Figure s3. Boxplots of Kruskal-Wallis Analysis of Normalized Protein Expression (NPX) Across Different Conditions.** This figure illustrates boxplots for the comparative analysis of NPX levels between conditions such as idiopathic Normal Pressure Hydrocephalus (iNPH), Mild Cognitive Impairment (MCI), and Healthy Control (HC). Each boxplot represents a unique protein, denoted by its gene symbol, and identified through the Kruskal-Wallis test for differences across conditions. Post-hoc pairwise Wilcoxon tests with Benjamini-Hochberg (BH) adjustments were performed, with significance levels indicated on the plots as follows: $p < 0.0001$ = '****', $p < 0.001$= '***', $p < 0.01$ = '**', $p < 0.05$ = '*', and 'ns' for non-significant differences.

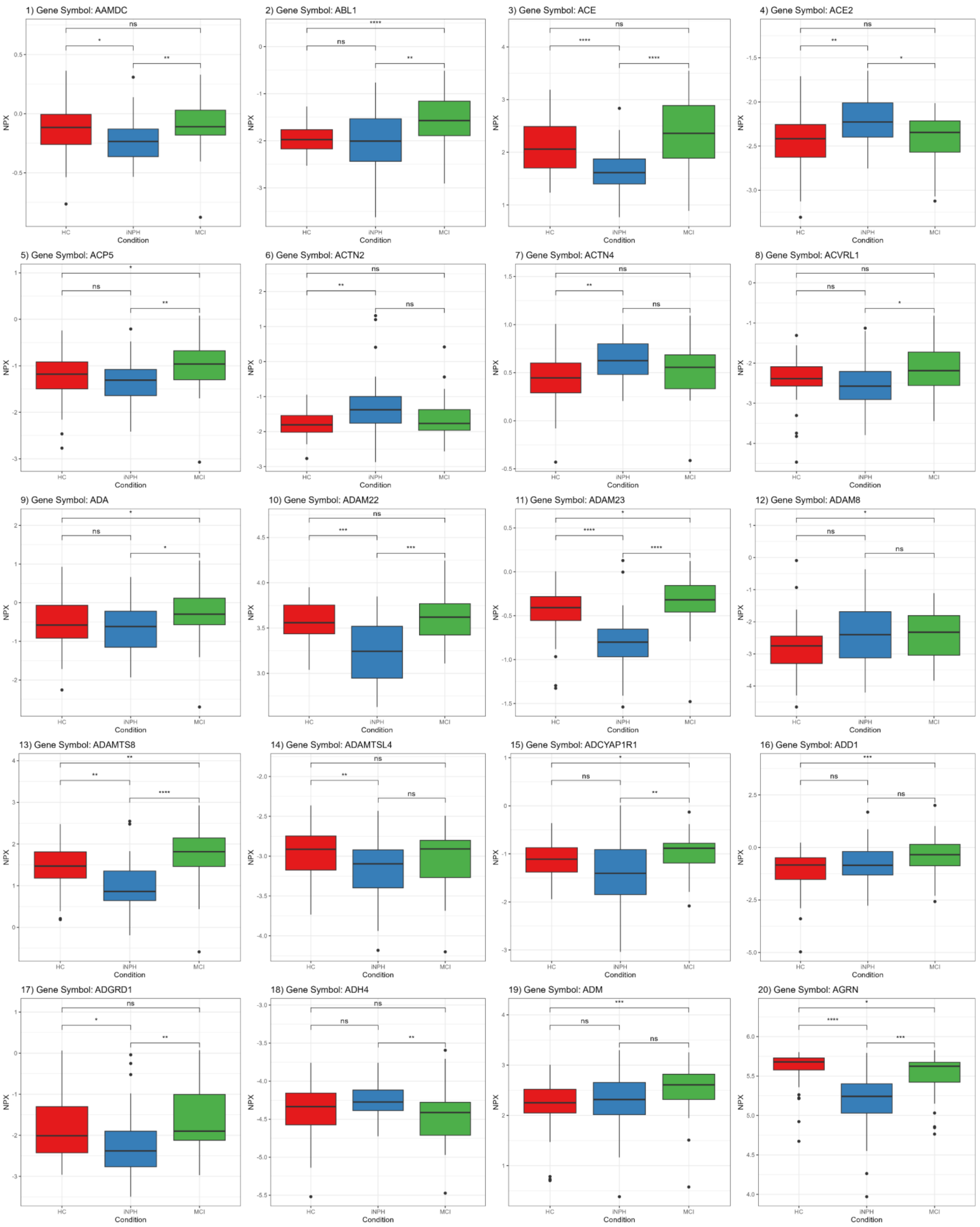
1) Gene Symbol: AAMDC
2) Gene Symbol: ABL1
3) Gene Symbol: ACE
4) Gene Symbol: ACE2
5) Gene Symbol: ACP5
6) Gene Symbol: ACTN2
7) Gene Symbol: ACTN4
8) Gene Symbol: ACVRL1
9) Gene Symbol: ADA
10) Gene Symbol: ADAM22
11) Gene Symbol: ADAM23
12) Gene Symbol: ADAM8
13) Gene Symbol: ADAMTS8
14) Gene Symbol: ADAMTSL4
15) Gene Symbol: ADCYAP1R1
16) Gene Symbol: ADD1
17) Gene Symbol: ADGRD1
18) Gene Symbol: ADH4
19) Gene Symbol: ADM
20) Gene Symbol: AGRN
NPX
HC
iNPH
MCI
Condition

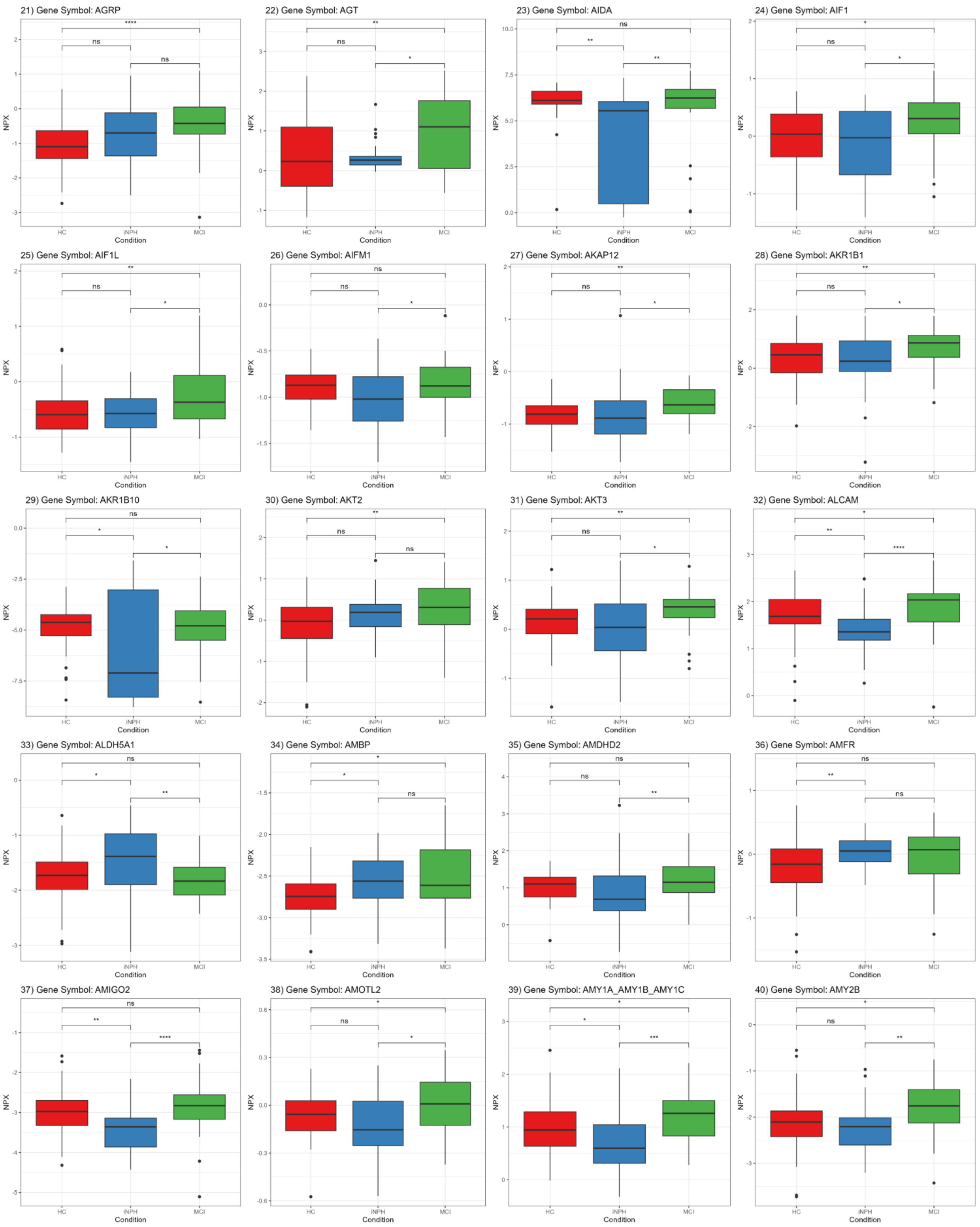
21) Gene Symbol: AGRP
22) Gene Symbol: AGT
23) Gene Symbol: AIDA
24) Gene Symbol: AIF1
25) Gene Symbol: AIF1L
26) Gene Symbol: AIFM1
27) Gene Symbol: AKAP12
28) Gene Symbol: AKR1B1
29) Gene Symbol: AKR1B10
30) Gene Symbol: AKT2
31) Gene Symbol: AKT3
32) Gene Symbol: ALCAM
33) Gene Symbol: ALDH5A1
34) Gene Symbol: AMBP
35) Gene Symbol: AMDHD2
36) Gene Symbol: AMFR
37) Gene Symbol: AMIGO2
38) Gene Symbol: AMOTL2
39) Gene Symbol: AMY1A_AMY1B_AMY1C
40) Gene Symbol: AMY2B
NPX
Condition
HC
iNPH
MCI

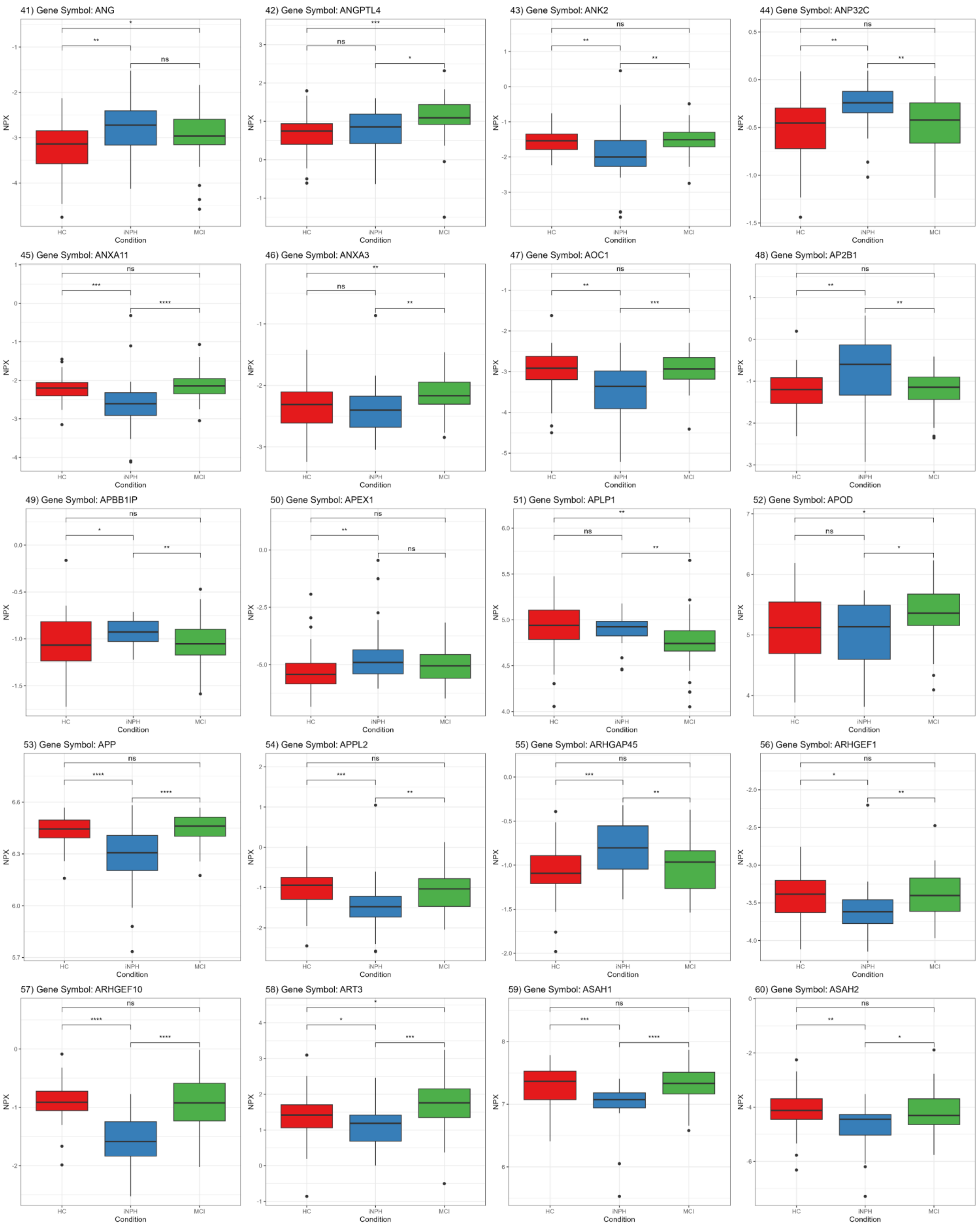
41) Gene Symbol: ANG
42) Gene Symbol: ANGPTL4
43) Gene Symbol: ANK2
44) Gene Symbol: ANP32C
45) Gene Symbol: ANXA11
46) Gene Symbol: ANXA3
47) Gene Symbol: AOC1
48) Gene Symbol: AP2B1
49) Gene Symbol: APBB1IP
50) Gene Symbol: APEX1
51) Gene Symbol: APLP1
52) Gene Symbol: APOD
53) Gene Symbol: APP
54) Gene Symbol: APPL2
55) Gene Symbol: ARHGAP45
56) Gene Symbol: ARHGEF1
57) Gene Symbol: ARHGEF10
58) Gene Symbol: ART3
59) Gene Symbol: ASAH1
60) Gene Symbol: ASAH2
NPX
Condition
HC
iNPH
MCI

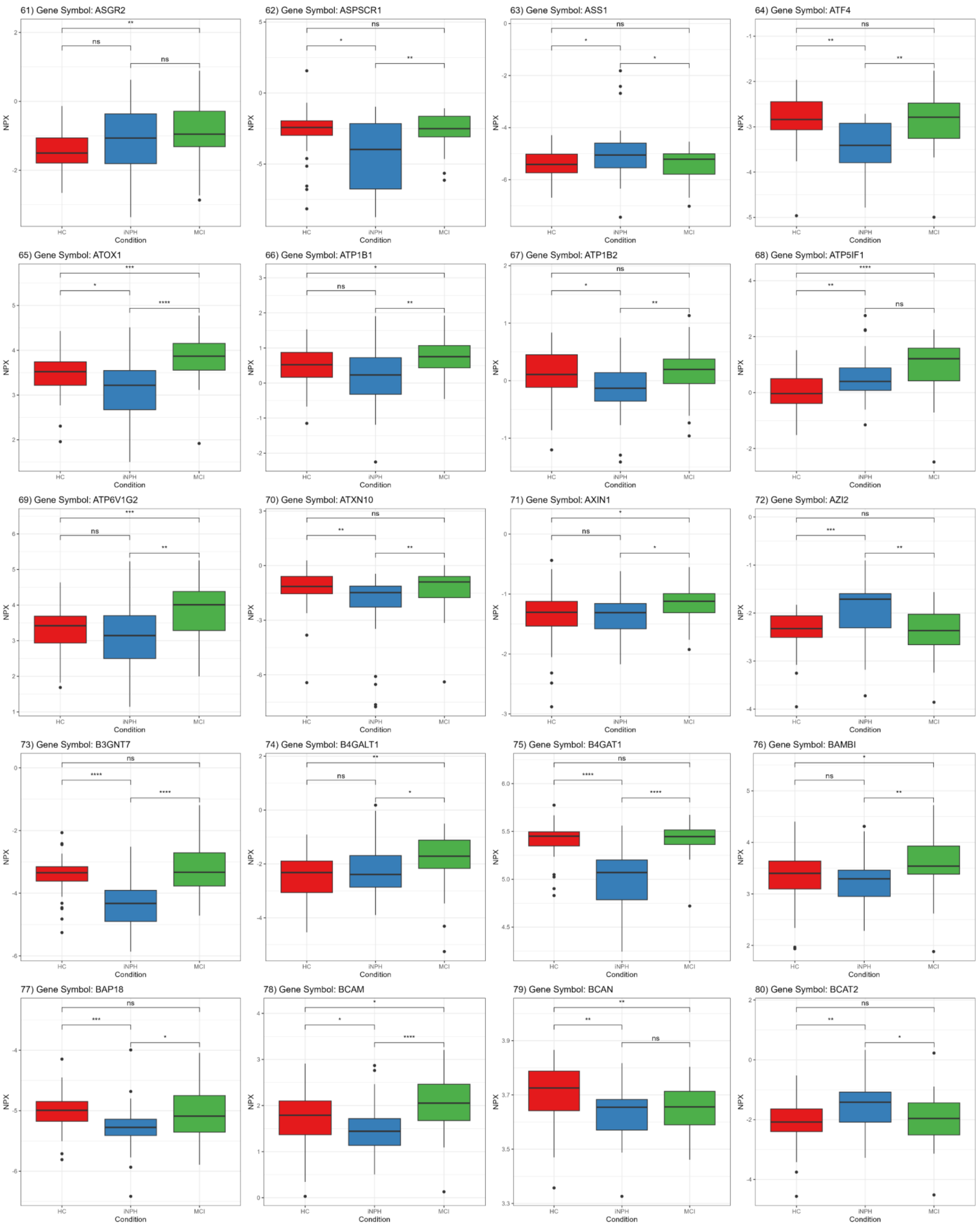
61) Gene Symbol: ASGR2
62) Gene Symbol: ASPSCR1
63) Gene Symbol: ASS1
64) Gene Symbol: ATF4
65) Gene Symbol: ATOX1
66) Gene Symbol: ATP1B1
67) Gene Symbol: ATP1B2
68) Gene Symbol: ATP5IF1
69) Gene Symbol: ATP6V1G2
70) Gene Symbol: ATXN10
71) Gene Symbol: AXIN1
72) Gene Symbol: AZI2
73) Gene Symbol: B3GNT7
74) Gene Symbol: B4GALT1
75) Gene Symbol: B4GAT1
76) Gene Symbol: BAMBI
77) Gene Symbol: BAP18
78) Gene Symbol: BCAM
79) Gene Symbol: BCAN
80) Gene Symbol: BCAT2
NPX
Condition
HC
iNPH
MCI

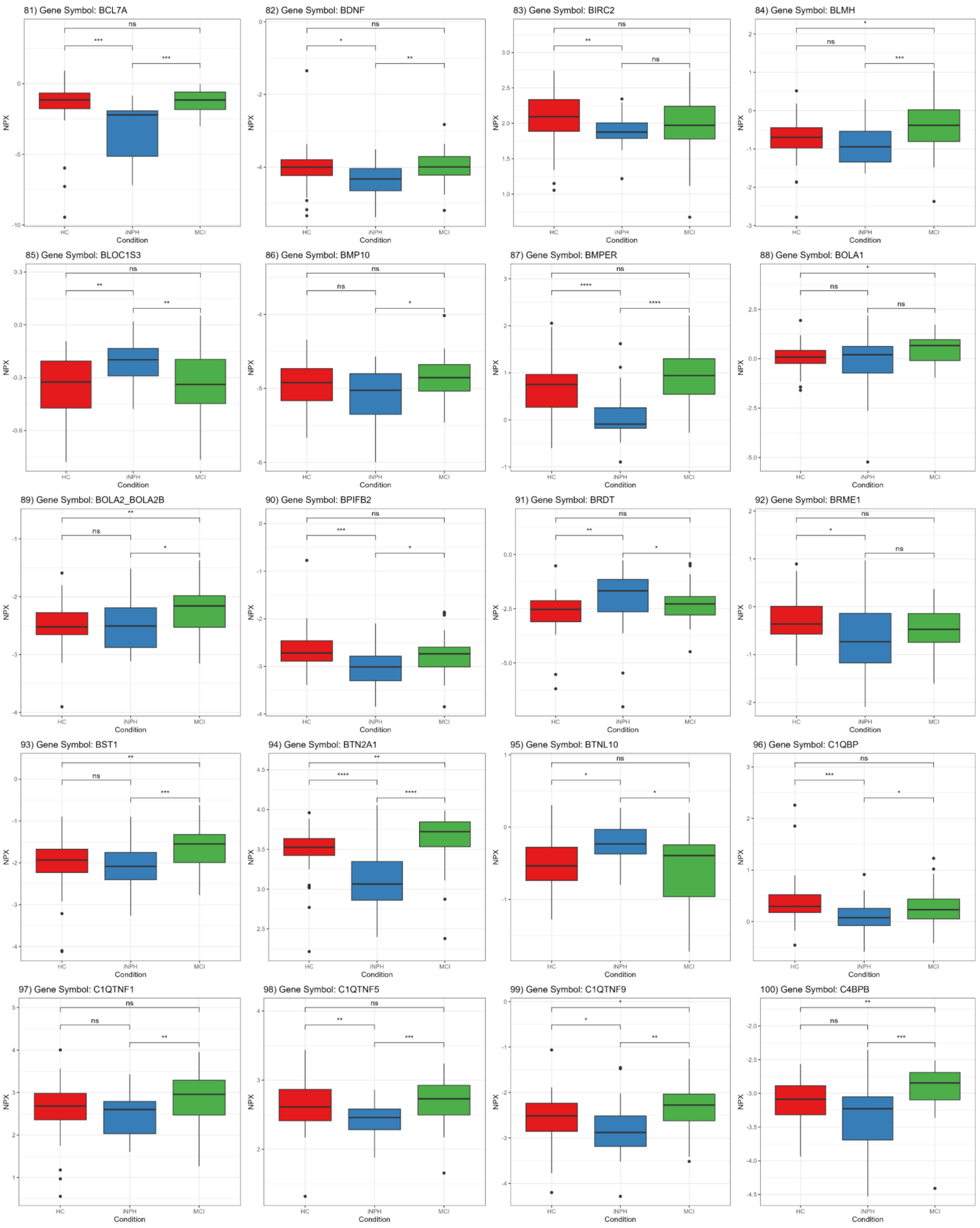
81) Gene Symbol: BCL7A
82) Gene Symbol: BDNF
83) Gene Symbol: BIRC2
84) Gene Symbol: BLMH
85) Gene Symbol: BLOC1S3
86) Gene Symbol: BMP10
87) Gene Symbol: BMPER
88) Gene Symbol: BOLA1
89) Gene Symbol: BOLA2_BOLA2B
90) Gene Symbol: BPIFB2
91) Gene Symbol: BRDT
92) Gene Symbol: BRME1
93) Gene Symbol: BST1
94) Gene Symbol: BTN2A1
95) Gene Symbol: BTNL10
96) Gene Symbol: C1QBP
97) Gene Symbol: C1QTNF1
98) Gene Symbol: C1QTNF5
99) Gene Symbol: C1QTNF9
100) Gene Symbol: C4BPB
NPX
Condition
HC
iNPH
MCI

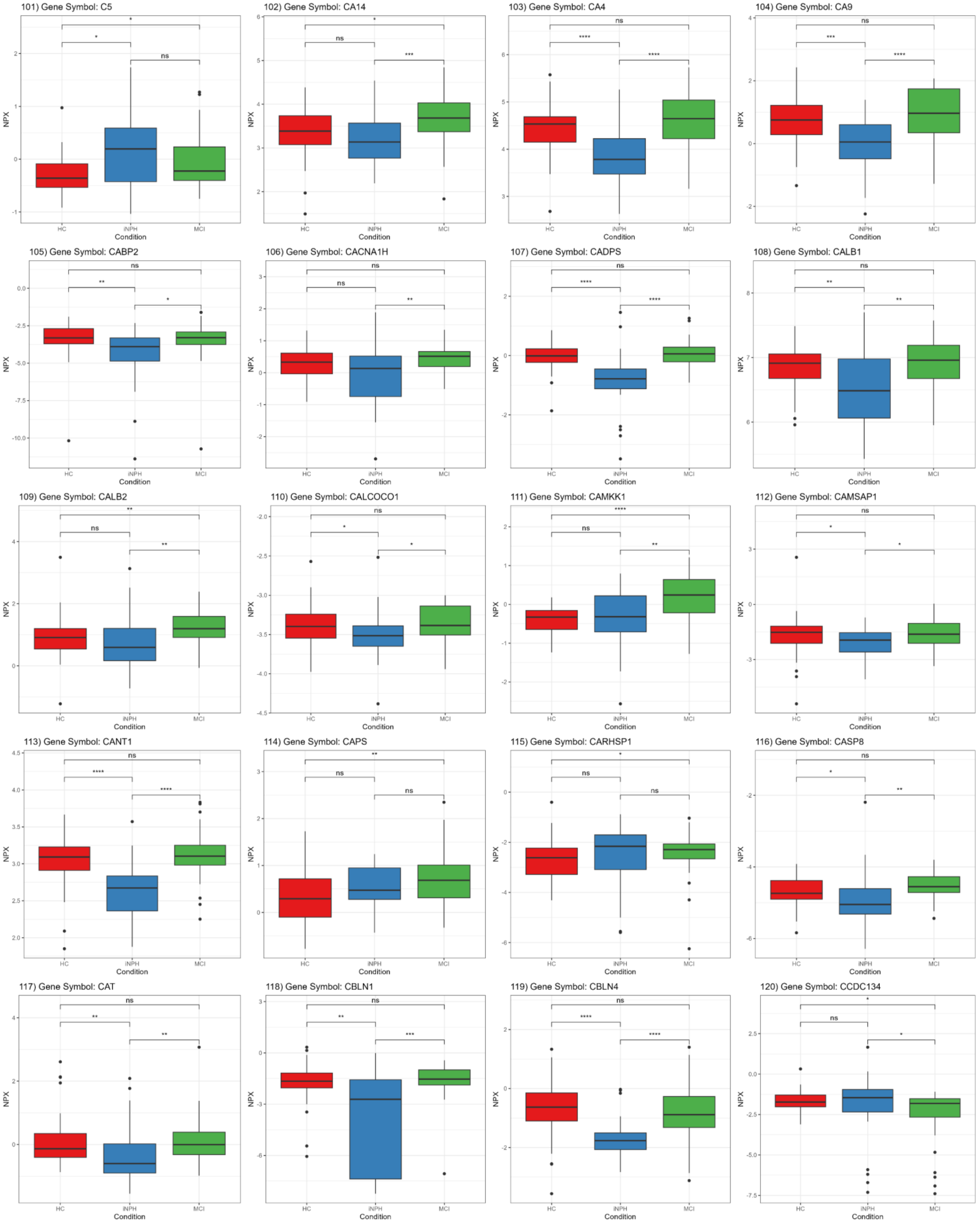
101) Gene Symbol: C5
102) Gene Symbol: CA14
103) Gene Symbol: CA4
104) Gene Symbol: CA9
105) Gene Symbol: CABP2
106) Gene Symbol: CACNA1H
107) Gene Symbol: CADPS
108) Gene Symbol: CALB1
109) Gene Symbol: CALB2
110) Gene Symbol: CALCOCO1
111) Gene Symbol: CAMKK1
112) Gene Symbol: CAMSAP1
113) Gene Symbol: CANT1
114) Gene Symbol: CAPS
115) Gene Symbol: CARHSP1
116) Gene Symbol: CASP8
117) Gene Symbol: CAT
118) Gene Symbol: CBLN1
119) Gene Symbol: CBLN4
120) Gene Symbol: CCDC134
NPX
Condition
HC
iNPH
MCI

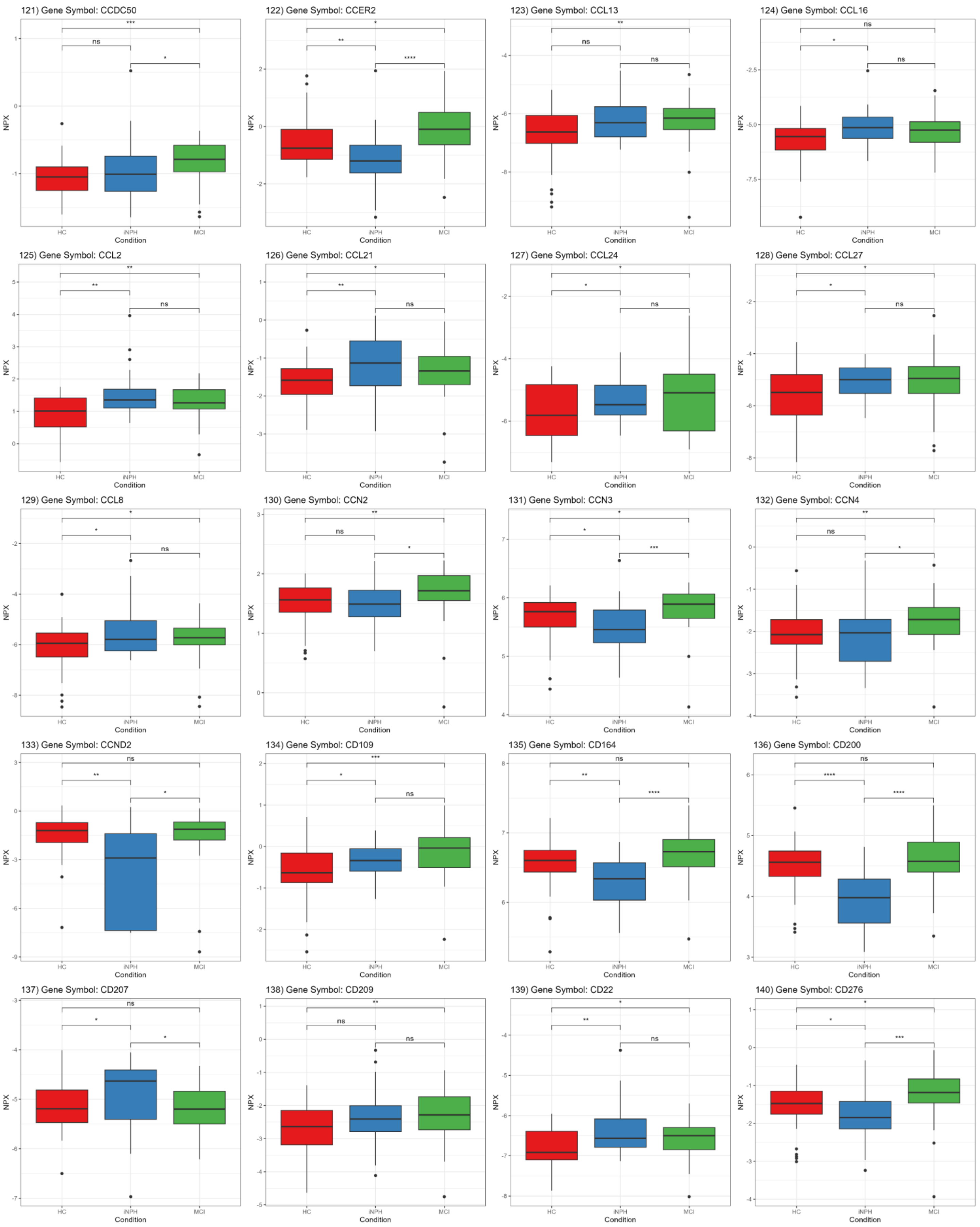
121) Gene Symbol: CCDC50
122) Gene Symbol: CCER2
123) Gene Symbol: CCL13
124) Gene Symbol: CCL16
125) Gene Symbol: CCL2
126) Gene Symbol: CCL21
127) Gene Symbol: CCL24
128) Gene Symbol: CCL27
129) Gene Symbol: CCL8
130) Gene Symbol: CCN2
131) Gene Symbol: CCN3
132) Gene Symbol: CCN4
133) Gene Symbol: CCND2
134) Gene Symbol: CD109
135) Gene Symbol: CD164
136) Gene Symbol: CD200
137) Gene Symbol: CD207
138) Gene Symbol: CD209
139) Gene Symbol: CD22
140) Gene Symbol: CD276
NPX
Condition
HC
iNPH
MCI

141) Gene Symbol: CD300LF

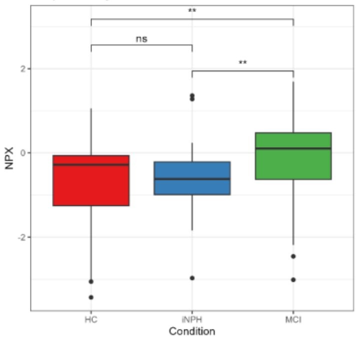


142) Gene Symbol: CD34

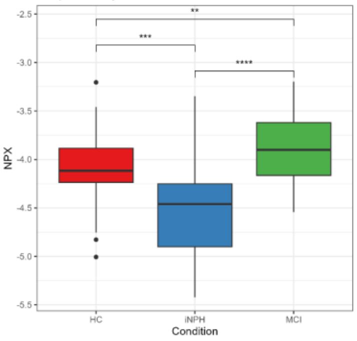


143) Gene Symbol: CD36

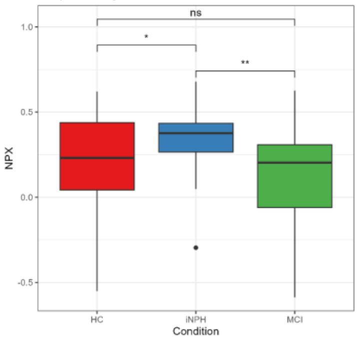


144) Gene Symbol: CD4

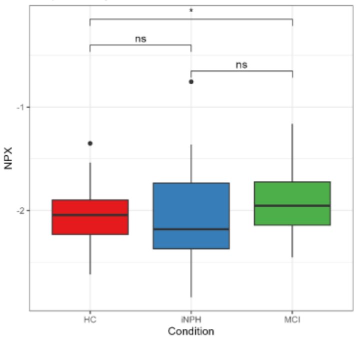


145) Gene Symbol: CD40

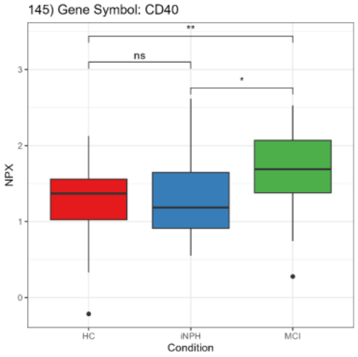


146) Gene Symbol: CD46

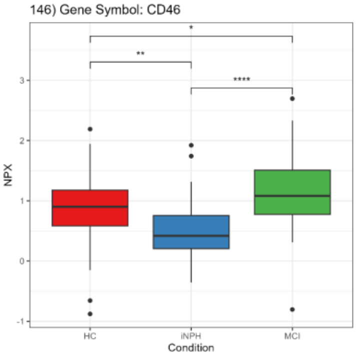


147) Gene Symbol: CD55

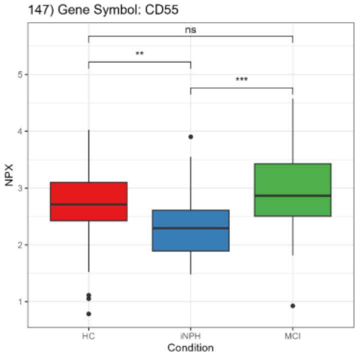


148) Gene Symbol: CD58

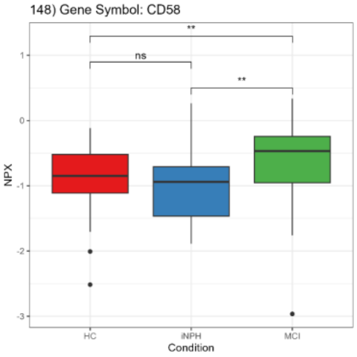


149) Gene Symbol: CD59

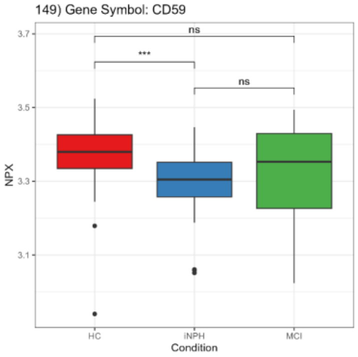


150) Gene Symbol: CD5L

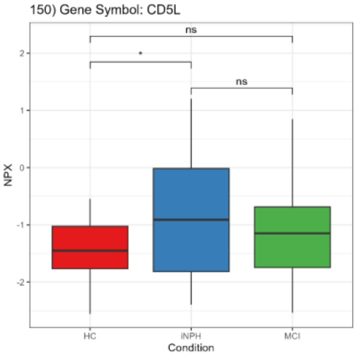


151) Gene Symbol: CD74

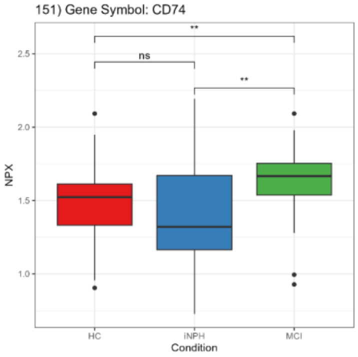


152) Gene Symbol: CD80

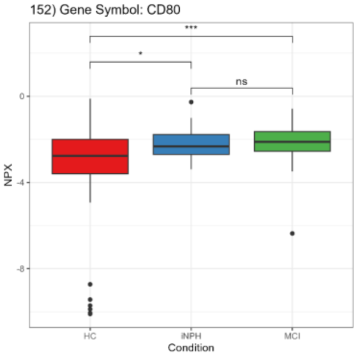


153) Gene Symbol: CD82

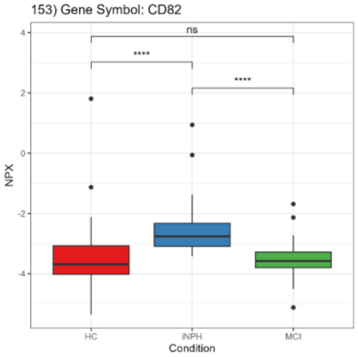


154) Gene Symbol: CD83

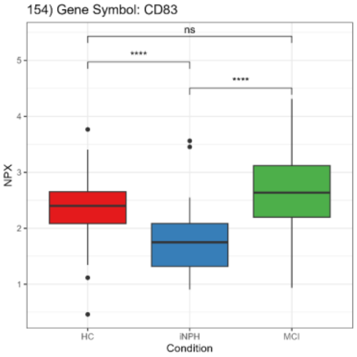


155) Gene Symbol: CD99L2

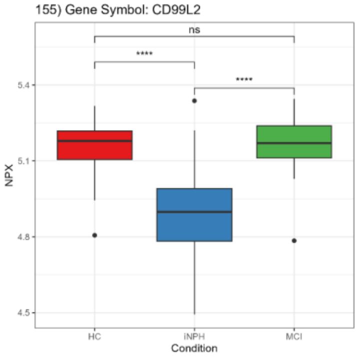


156) Gene Symbol: CDC42BPB

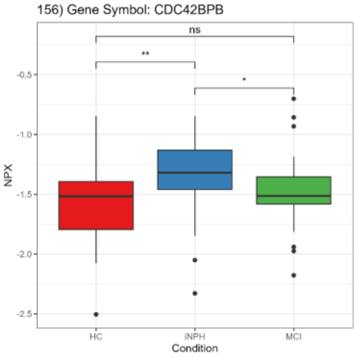


157) Gene Symbol: CDH15

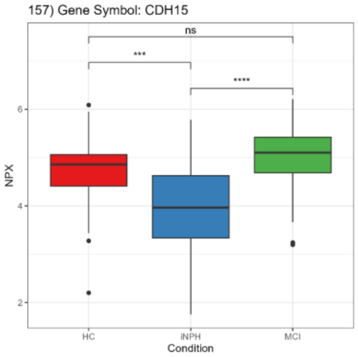


158) Gene Symbol: CDH2

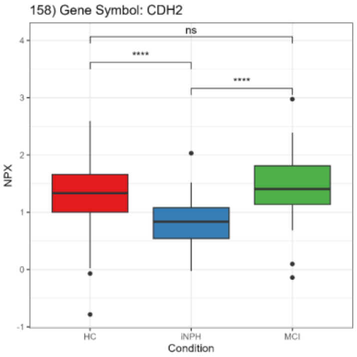


159) Gene Symbol: CDH3

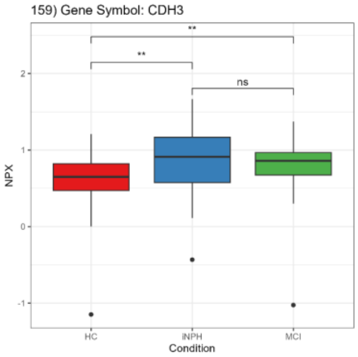


160) Gene Symbol: CDH6

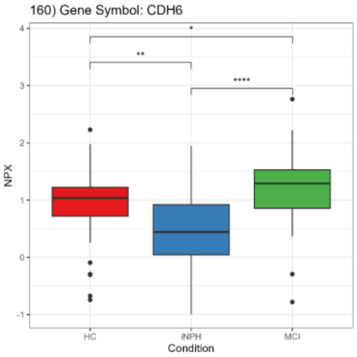

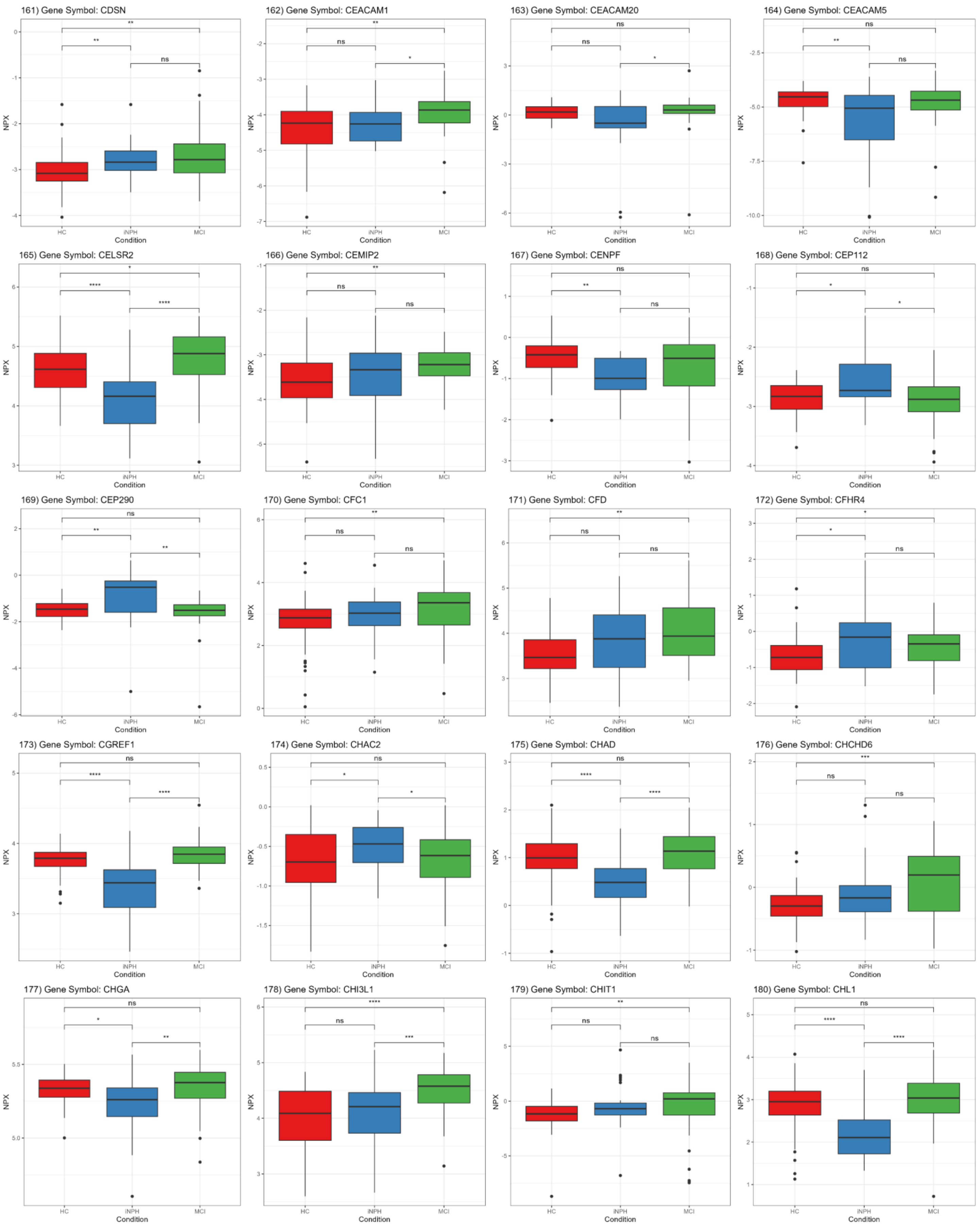
161) Gene Symbol: CDSN
162) Gene Symbol: CEACAM1
163) Gene Symbol: CEACAM20
164) Gene Symbol: CEACAM5
165) Gene Symbol: CELSR2
166) Gene Symbol: CEMIP2
167) Gene Symbol: CENPF
168) Gene Symbol: CEP112
169) Gene Symbol: CEP290
170) Gene Symbol: CFC1
171) Gene Symbol: CFD
172) Gene Symbol: CFHR4
173) Gene Symbol: CGREF1
174) Gene Symbol: CHAC2
175) Gene Symbol: CHAD
176) Gene Symbol: CHCHD6
177) Gene Symbol: CHGA
178) Gene Symbol: CHI3L1
179) Gene Symbol: CHIT1
180) Gene Symbol: CHL1
NPX
Condition
HC
iNPH
MCI

181) Gene Symbol: CHM

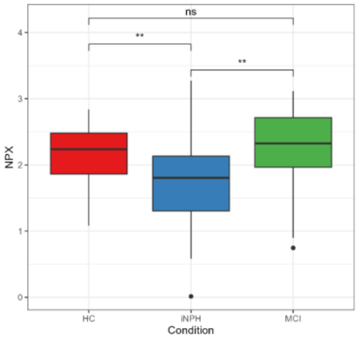


182) Gene Symbol: CHRDL1

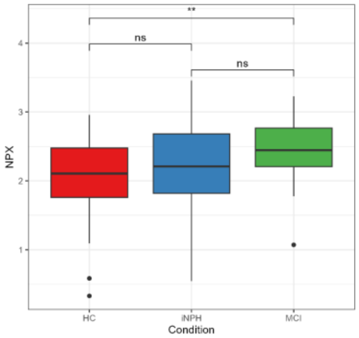


183) Gene Symbol: CIAPIN1

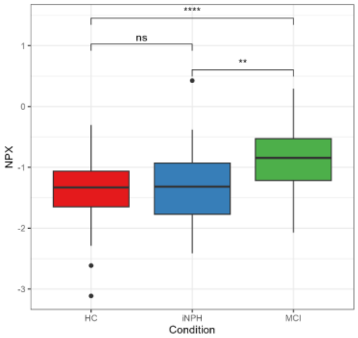


184) Gene Symbol: CKB

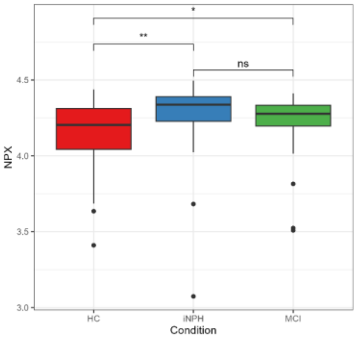


185) Gene Symbol: CKMT1A_CKMT1B

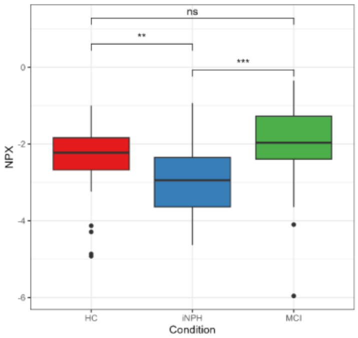


186) Gene Symbol: CLC

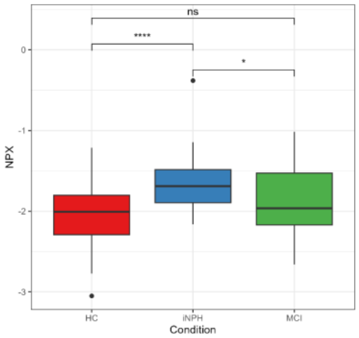


187) Gene Symbol: CLEC11A

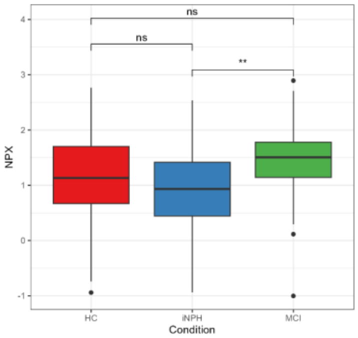


188) Gene Symbol: CLEC14A

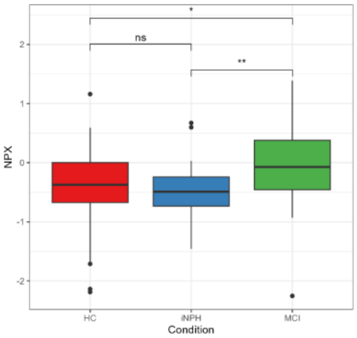


189) Gene Symbol: CLEC2L

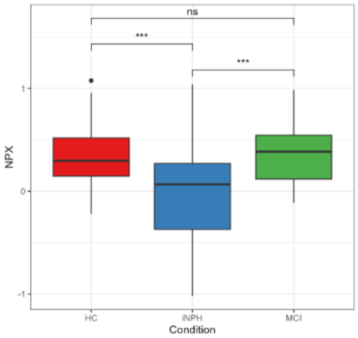


190) Gene Symbol: CLGN

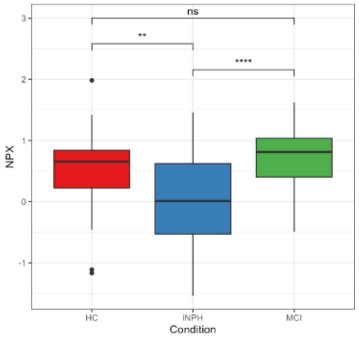


191) Gene Symbol: CLMP

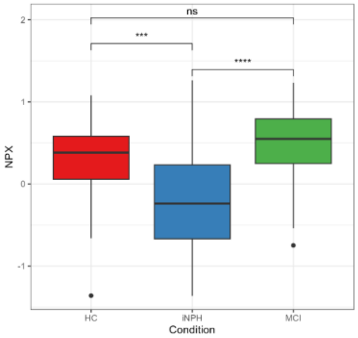


192) Gene Symbol: CLSTN2

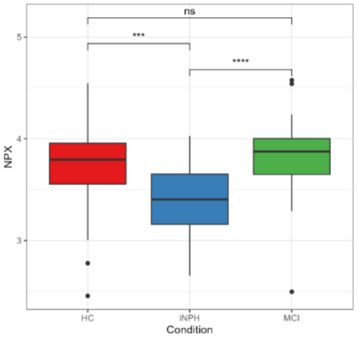


193) Gene Symbol: CLUL1

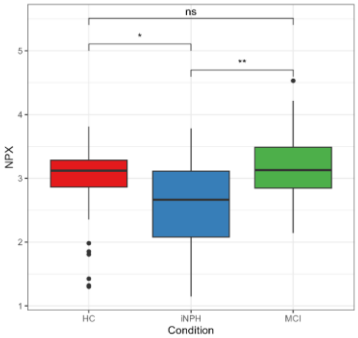


194) Gene Symbol: CNP

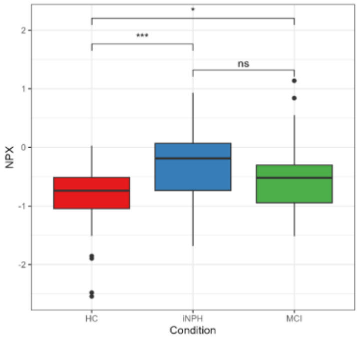


195) Gene Symbol: CNPY4

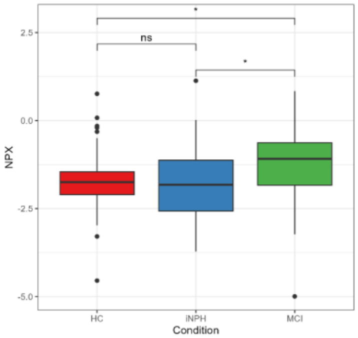


196) Gene Symbol: CNTN1

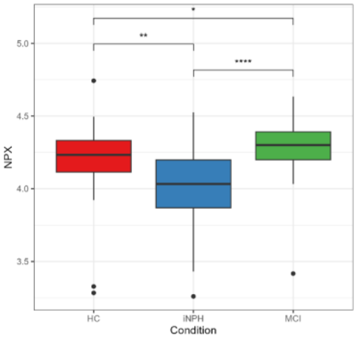


197) Gene Symbol: CNTN3

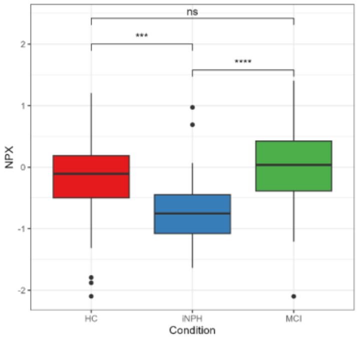


198) Gene Symbol: CNTN4

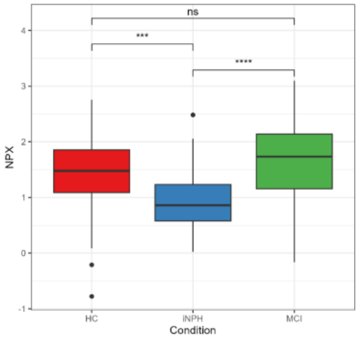


199) Gene Symbol: CNTN5

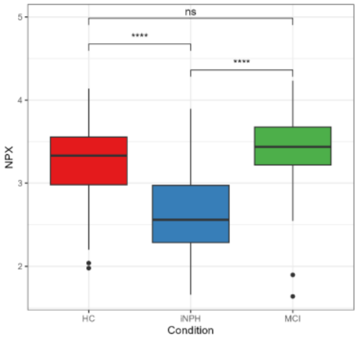


200) Gene Symbol: CNTNAP2

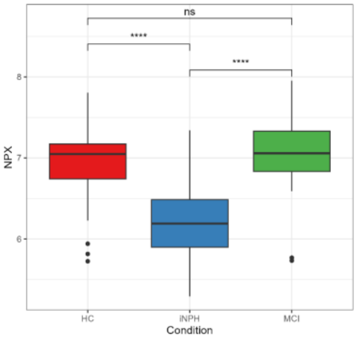

201) Gene Symbol: COCH

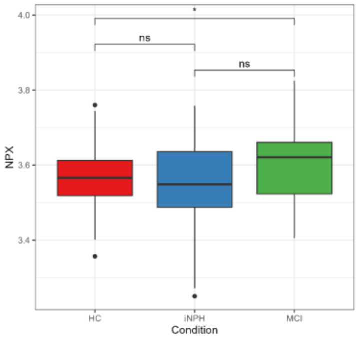


202) Gene Symbol: COL18A1

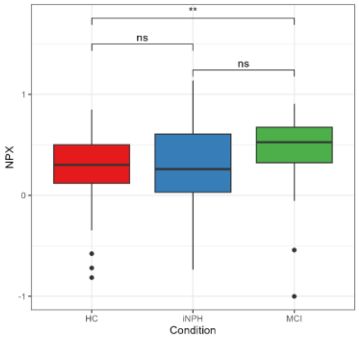


203) Gene Symbol: COL3A1

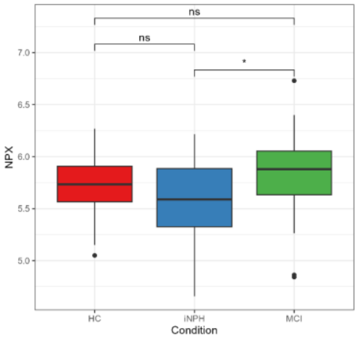


204) Gene Symbol: COL6A3

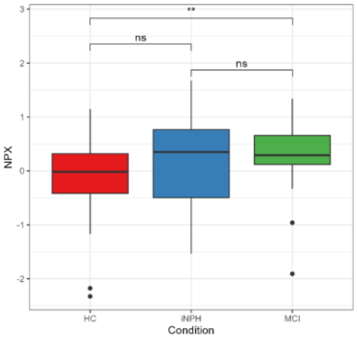


205) Gene Symbol: COL9A2

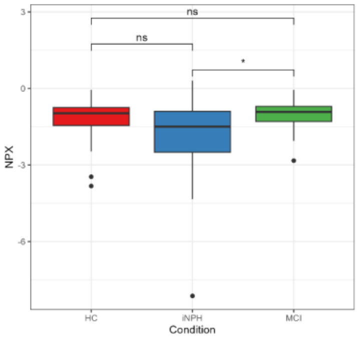


206) Gene Symbol: COLEC12

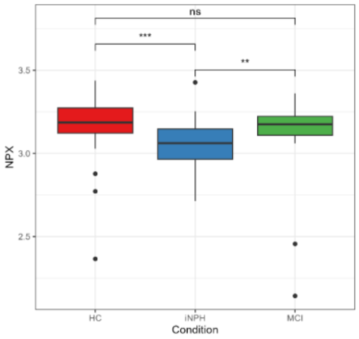


207) Gene Symbol: COMMD1

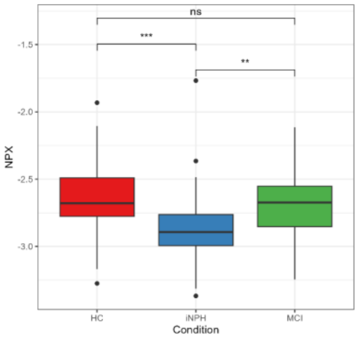


208) Gene Symbol: COPB2

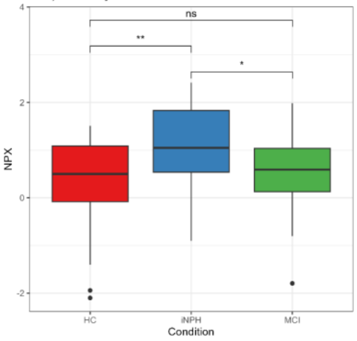


209) Gene Symbol: COX6B1

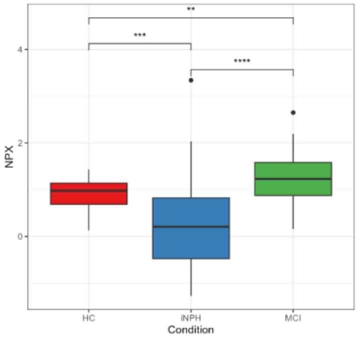


210) Gene Symbol: CPA1

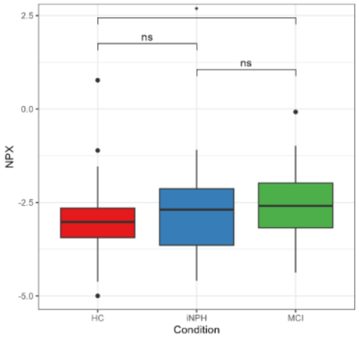


211) Gene Symbol: CPB1

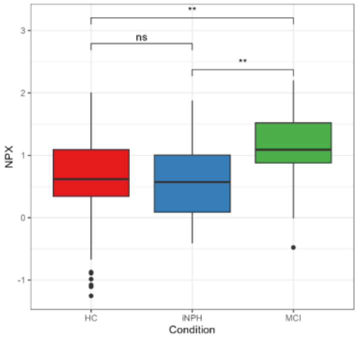


212) Gene Symbol: CPE

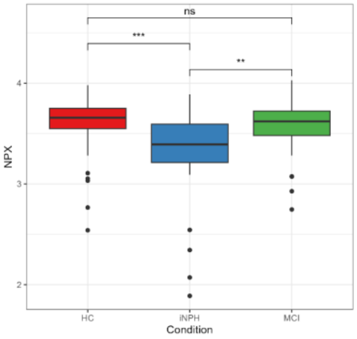


213) Gene Symbol: CPM

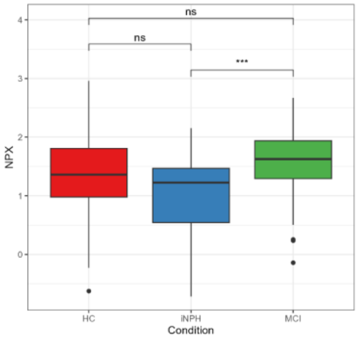


214) Gene Symbol: CPPED1

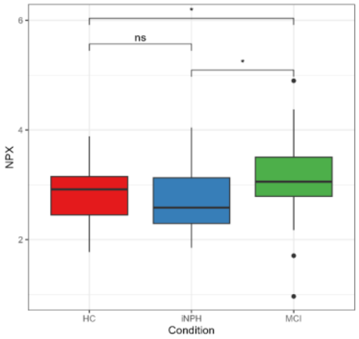


215) Gene Symbol: CPQ

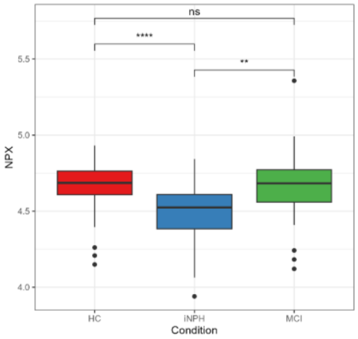


216) Gene Symbol: CPXM1

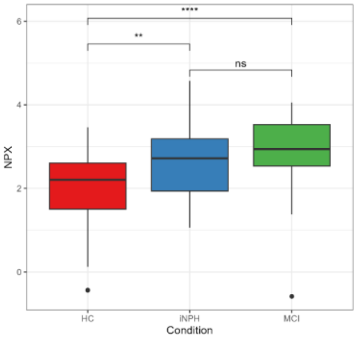


217) Gene Symbol: CRADD

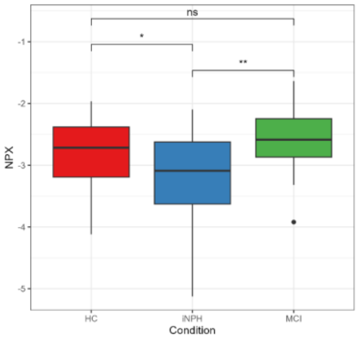


218) Gene Symbol: CREG1

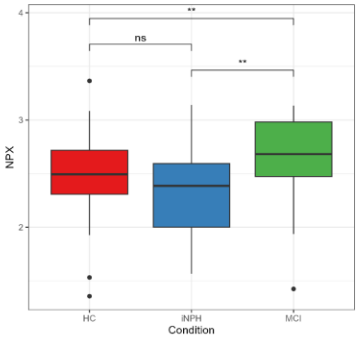


219) Gene Symbol: CRELD2

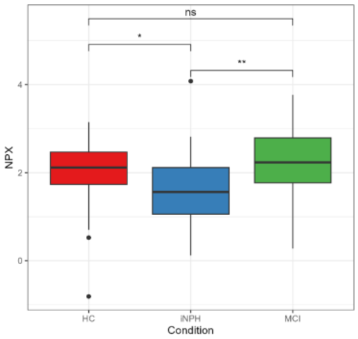


220) Gene Symbol: CRH

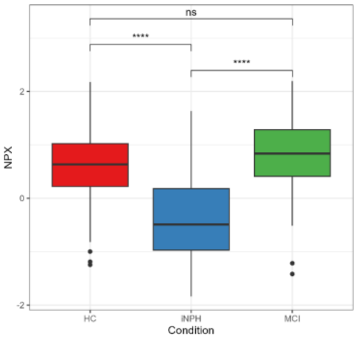

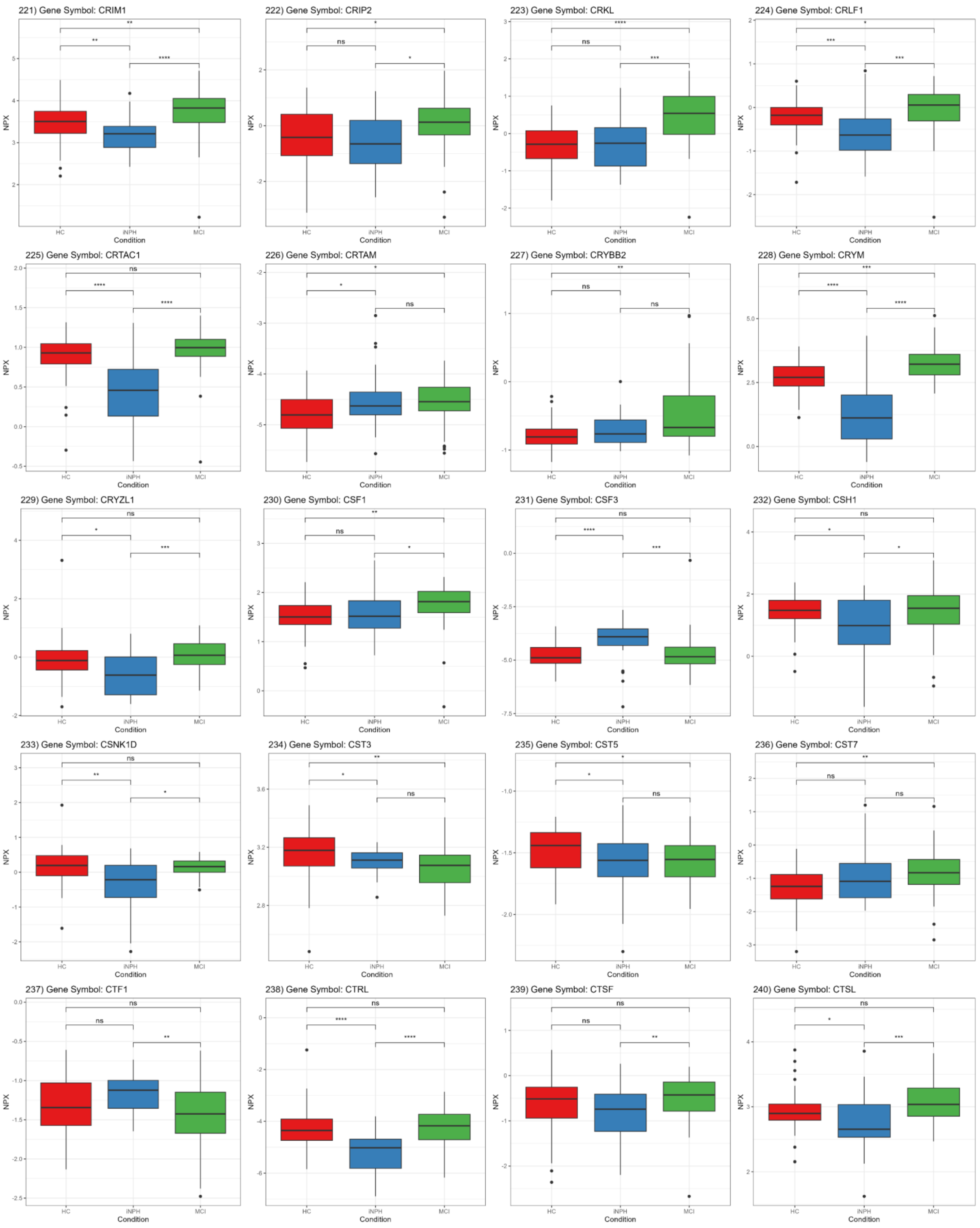
221) Gene Symbol: CRIM1
222) Gene Symbol: CRIP2
223) Gene Symbol: CRKL
224) Gene Symbol: CRLF1
225) Gene Symbol: CRTAC1
226) Gene Symbol: CRTAM
227) Gene Symbol: CRYBB2
228) Gene Symbol: CRYM
229) Gene Symbol: CRYZL1
230) Gene Symbol: CSF1
231) Gene Symbol: CSF3
232) Gene Symbol: CSH1
233) Gene Symbol: CSNK1D
234) Gene Symbol: CST3
235) Gene Symbol: CST5
236) Gene Symbol: CST7
237) Gene Symbol: CTF1
238) Gene Symbol: CTRL
239) Gene Symbol: CTSF
240) Gene Symbol: CTSL
NPX
Condition
HC
iNPH
MCI

241) Gene Symbol: CTSO

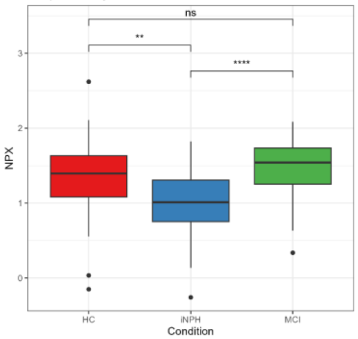


242) Gene Symbol: CXCL1

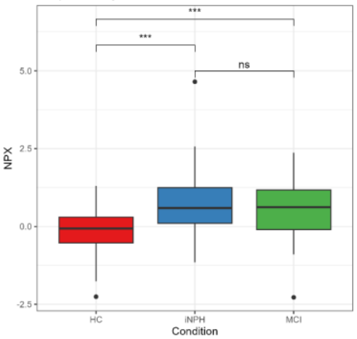


243) Gene Symbol: CXCL11

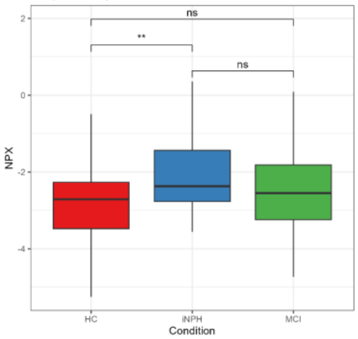


244) Gene Symbol: CXCL13

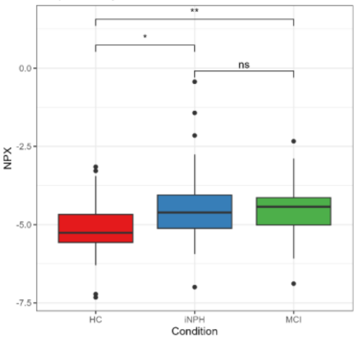


245) Gene Symbol: CXCL14

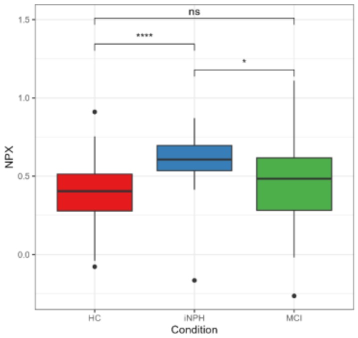


246) Gene Symbol: CXCL16

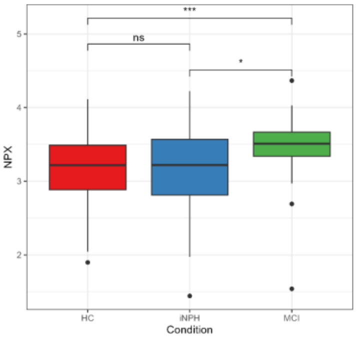


247) Gene Symbol: CXCL3

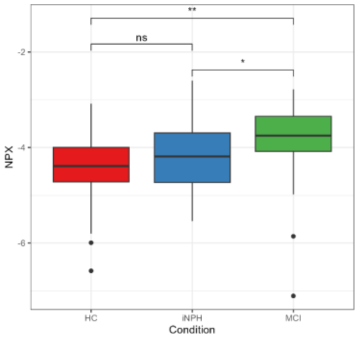


248) Gene Symbol: CXCL5

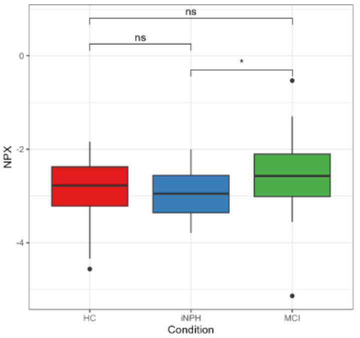


249) Gene Symbol: CXCL8

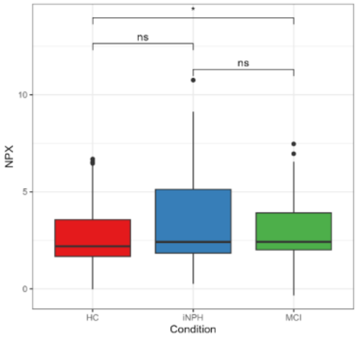


250) Gene Symbol: CXCL9

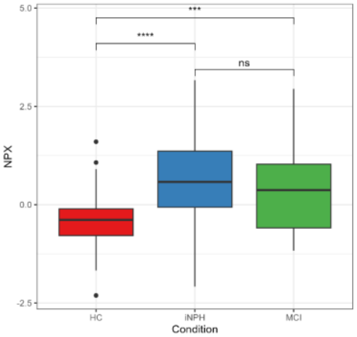


251) Gene Symbol: CYP24A1

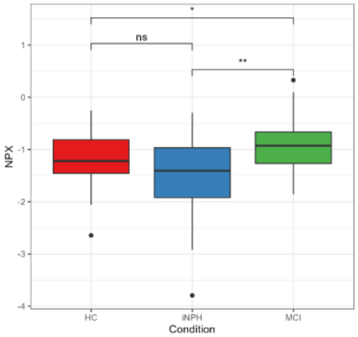


252) Gene Symbol: CYTL1

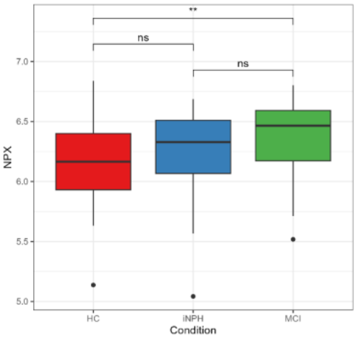


253) Gene Symbol: DAAM1

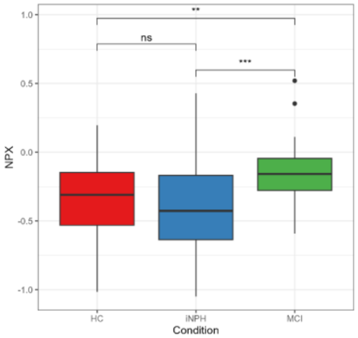


254) Gene Symbol: DAG1

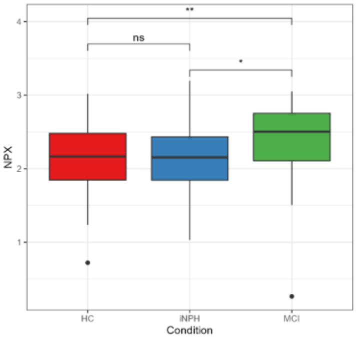


255) Gene Symbol: DAPK2

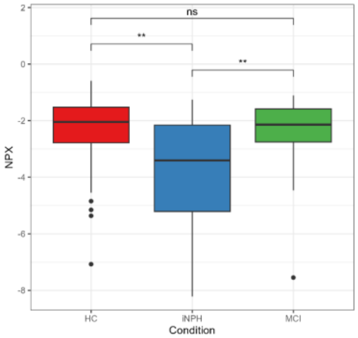


256) Gene Symbol: DAPP1

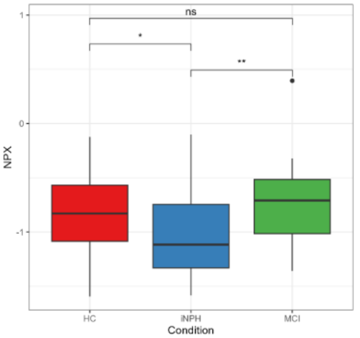


257) Gene Symbol: DBI

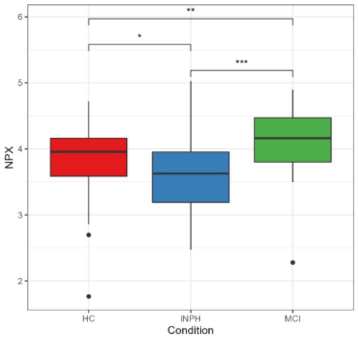


258) Gene Symbol: DBNL

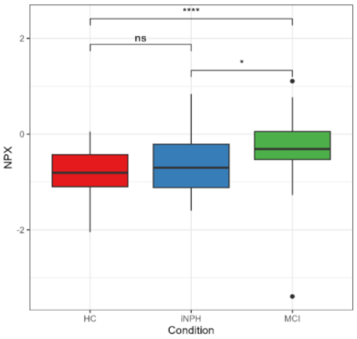


259) Gene Symbol: DCBLD2

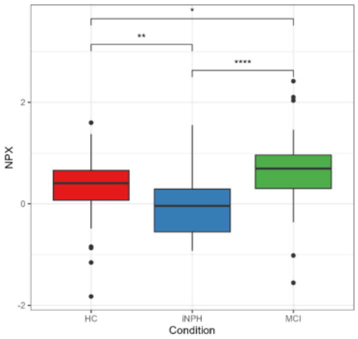


260) Gene Symbol: DCC

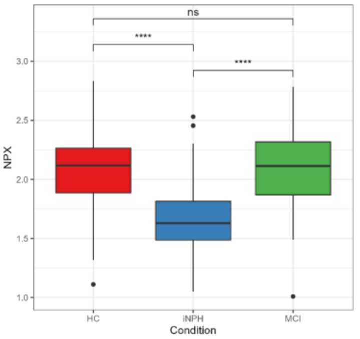

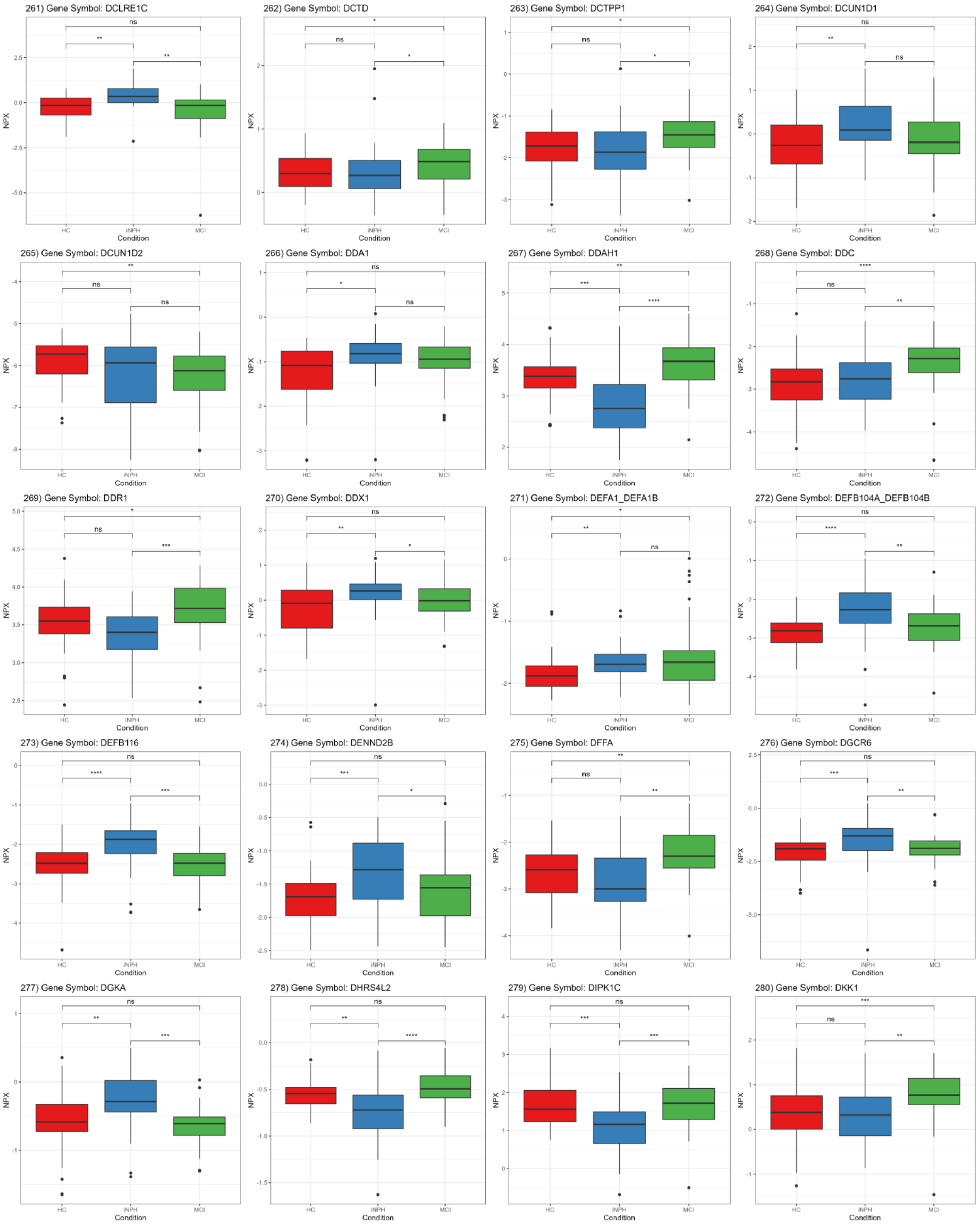
261) Gene Symbol: DCLRE1C
262) Gene Symbol: DCTD
263) Gene Symbol: DCTPP1
264) Gene Symbol: DCUN1D1
265) Gene Symbol: DCUN1D2
266) Gene Symbol: DDA1
267) Gene Symbol: DDAH1
268) Gene Symbol: DDC
269) Gene Symbol: DDR1
270) Gene Symbol: DDX1
271) Gene Symbol: DEFA1_DEFA1B
272) Gene Symbol: DEFB104A_DEFB104B
273) Gene Symbol: DEFB116
274) Gene Symbol: DENND2B
275) Gene Symbol: DFFA
276) Gene Symbol: DGCR6
277) Gene Symbol: DGKA
278) Gene Symbol: DHRS4L2
279) Gene Symbol: DIPK1C
280) Gene Symbol: DKK1
NPX
Condition
HC
iNPH
MCI

281) Gene Symbol: DKK3

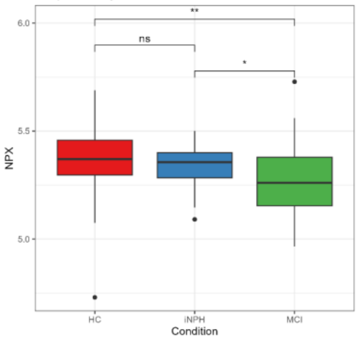


282) Gene Symbol: DLG4

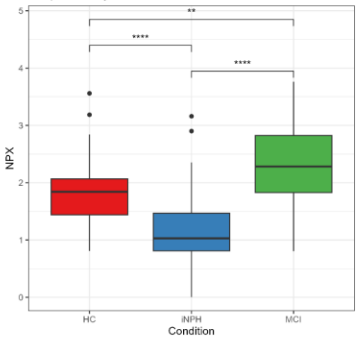


283) Gene Symbol: DLK1

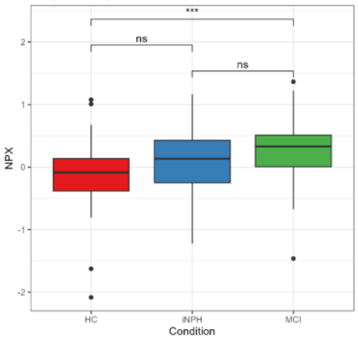


284) Gene Symbol: DLL1

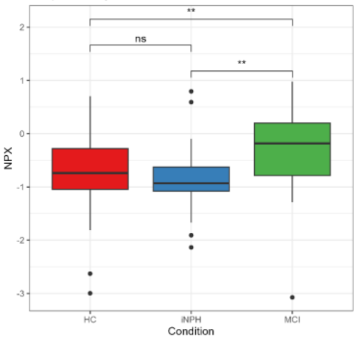


285) Gene Symbol: DMD

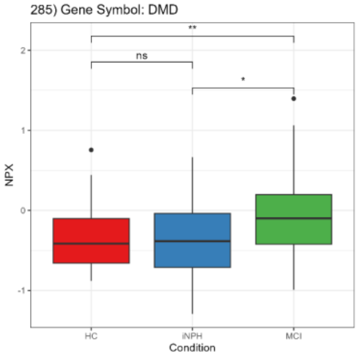


286) Gene Symbol: DMP1

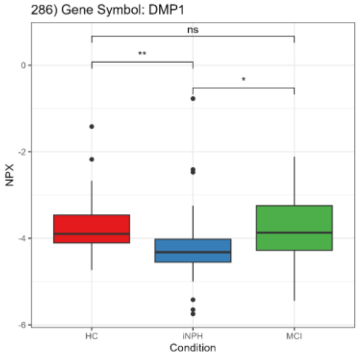


287) Gene Symbol: DNAJB8

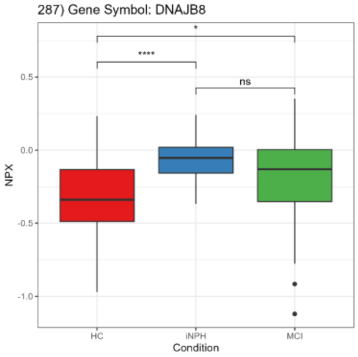


288) Gene Symbol: DNER

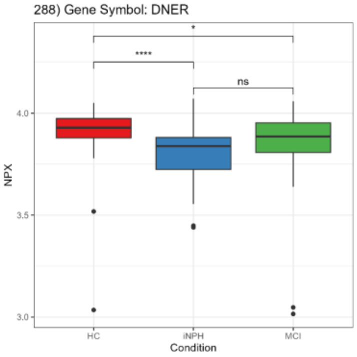


289) Gene Symbol: DNM1

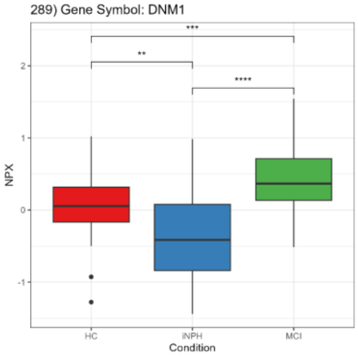


290) Gene Symbol: DNM3

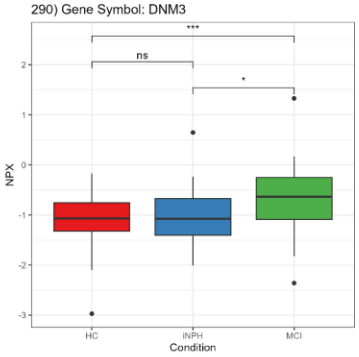


291) Gene Symbol: DNPH1

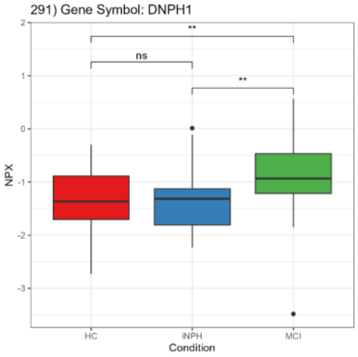


292) Gene Symbol: DOC2B

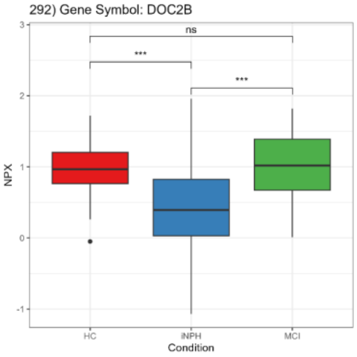


293) Gene Symbol: DOK1

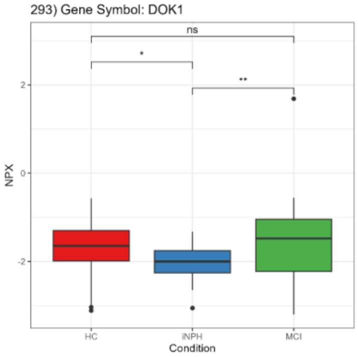


294) Gene Symbol: DPP6

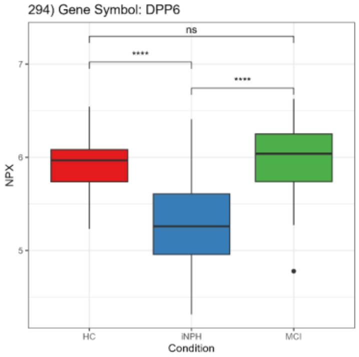


295) Gene Symbol: DPP7

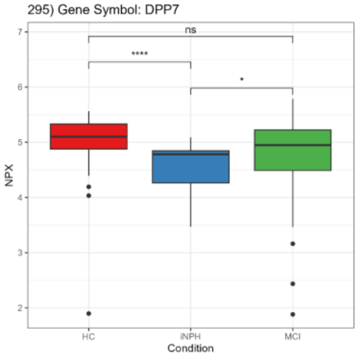


296) Gene Symbol: DRAXIN

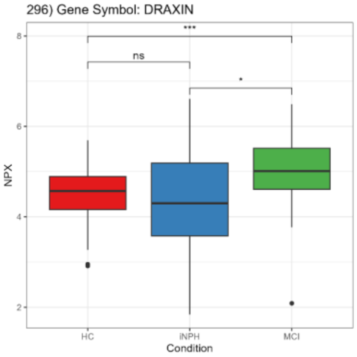


297) Gene Symbol: DSC2

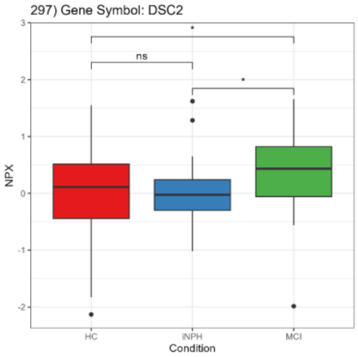


298) Gene Symbol: DSG2

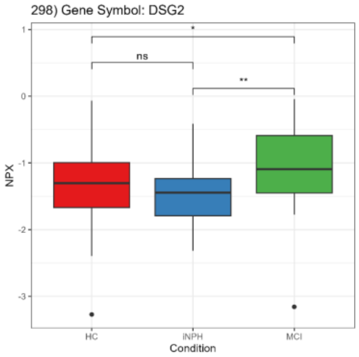


299) Gene Symbol: DUSP13

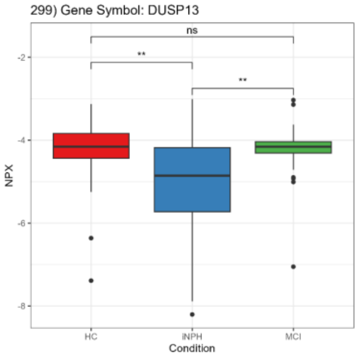


300) Gene Symbol: DUT

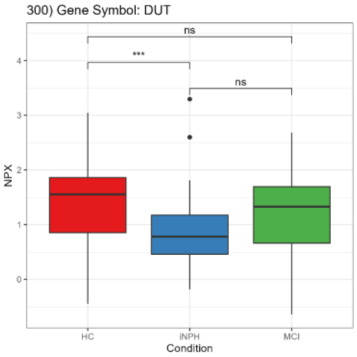

301) Gene Symbol: DXO

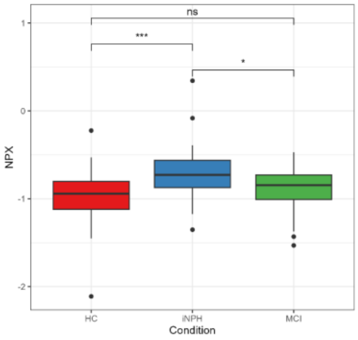


302) Gene Symbol: EDA2R

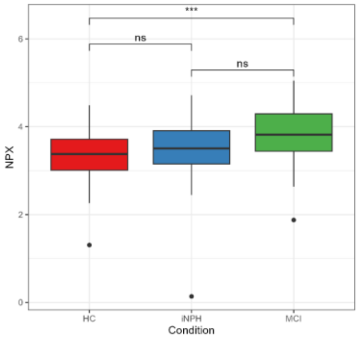


303) Gene Symbol: EFEMP1

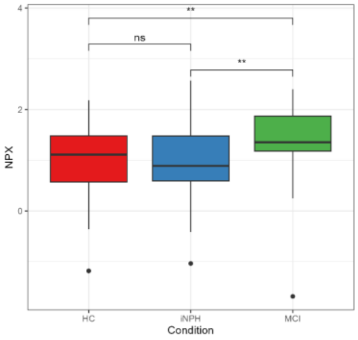


304) Gene Symbol: EFHD1

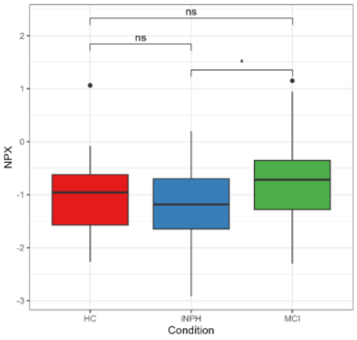


305) Gene Symbol: EFNB2

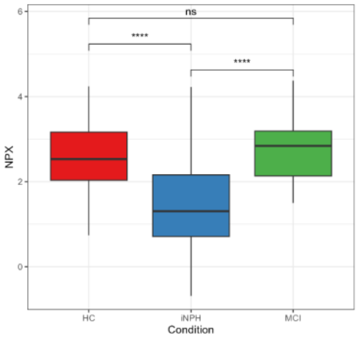


306) Gene Symbol: EGFL7

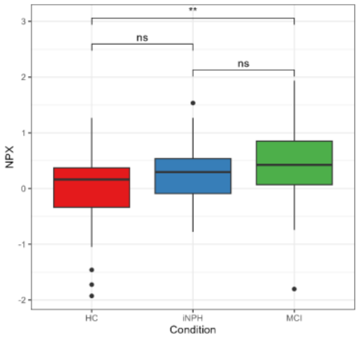


307) Gene Symbol: EGFR

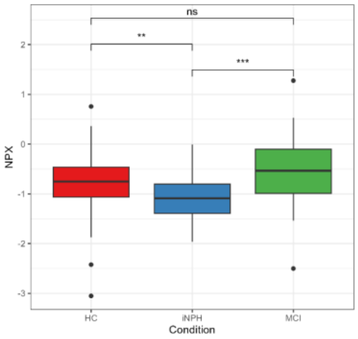


308) Gene Symbol: EHBP1

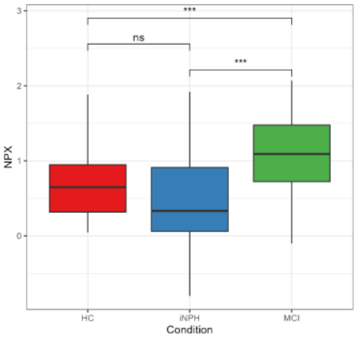


309) Gene Symbol: EIF2AK3

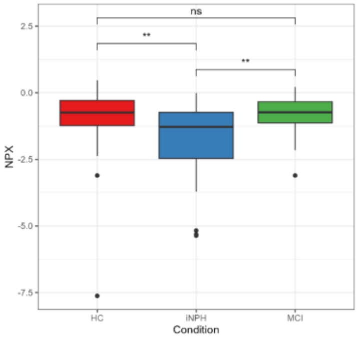


310) Gene Symbol: EIF4E

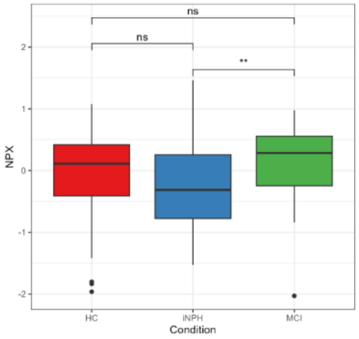


311) Gene Symbol: EIF4EBP1

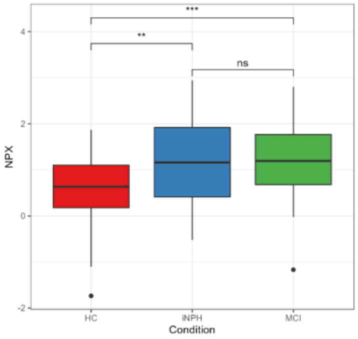


312) Gene Symbol: EIF4G1

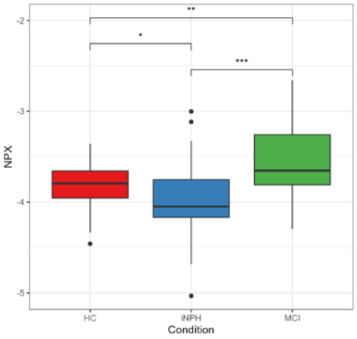


313) Gene Symbol: EIF5

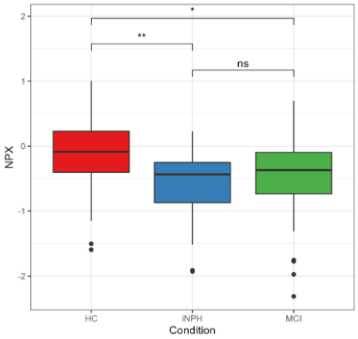


314) Gene Symbol: ELAC1

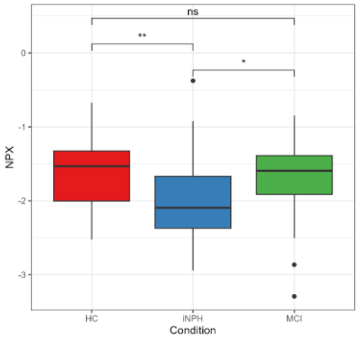


315) Gene Symbol: ELOA

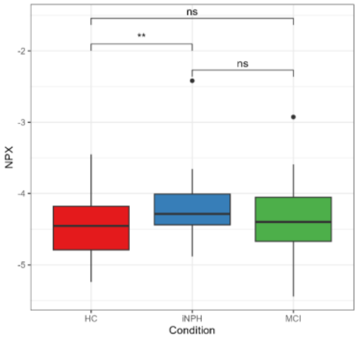


316) Gene Symbol: ENDOU

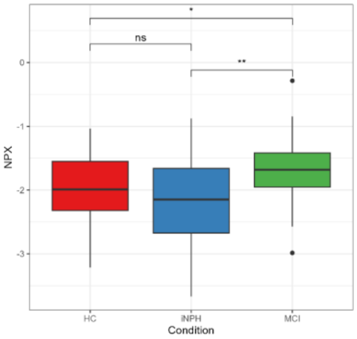


317) Gene Symbol: ENO1

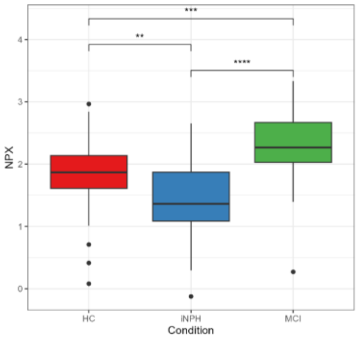


318) Gene Symbol: ENO2

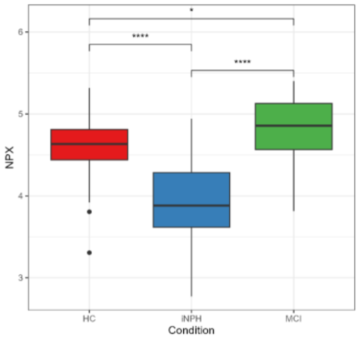


319) Gene Symbol: ENO3

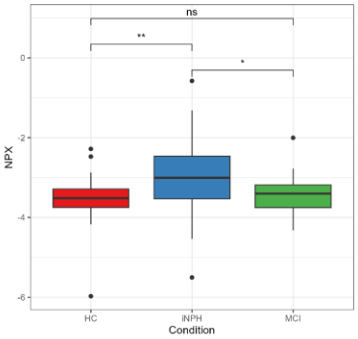


320) Gene Symbol: ENPP2

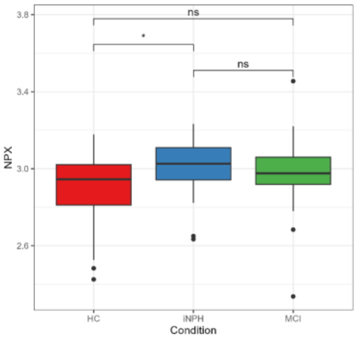

321) Gene Symbol: ENPP5

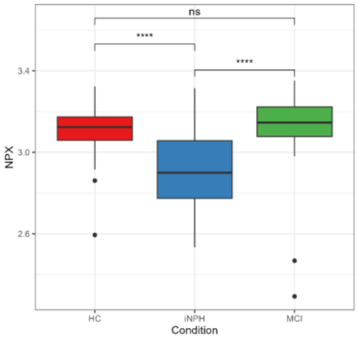


322) Gene Symbol: ENTPD6

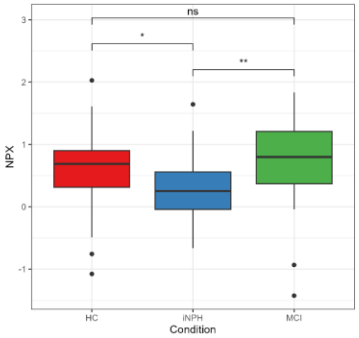


323) Gene Symbol: EPHA2

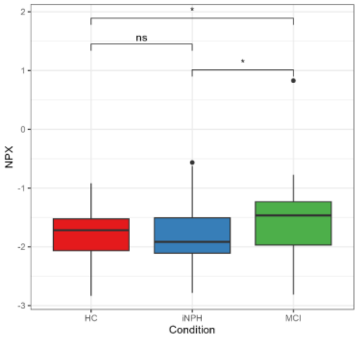


324) Gene Symbol: EPHA4

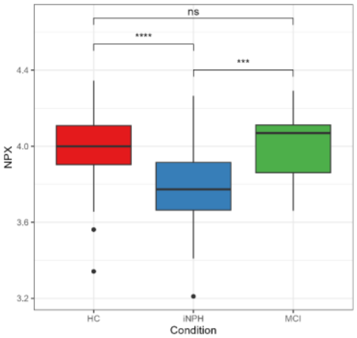


325) Gene Symbol: ERBB3

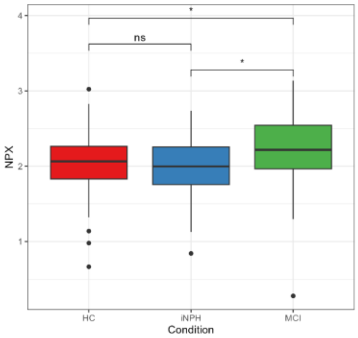


326) Gene Symbol: ERC2

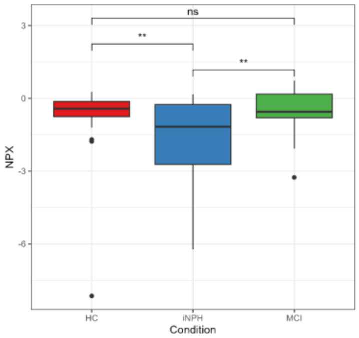


327) Gene Symbol: ERN1

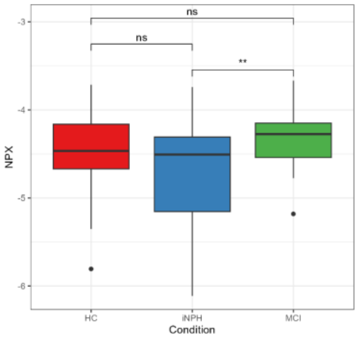


328) Gene Symbol: ESAM

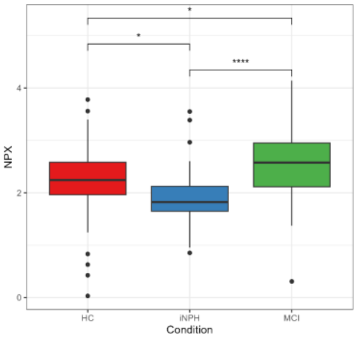


329) Gene Symbol: ESM1

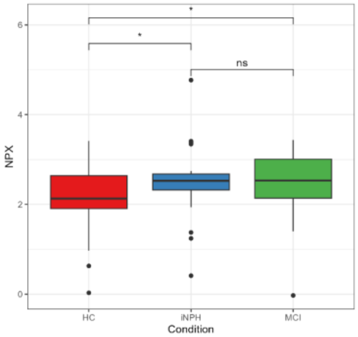


330) Gene Symbol: ESR1

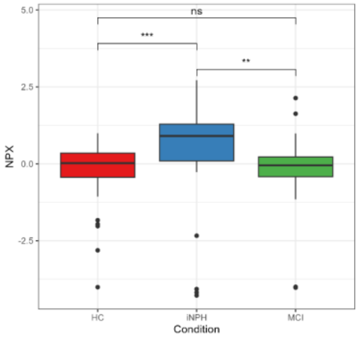


331) Gene Symbol: ESYT2

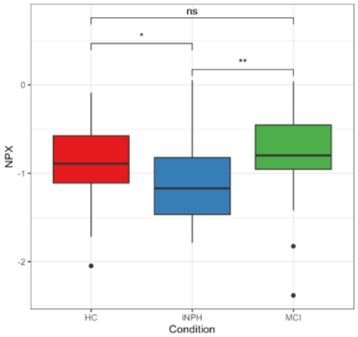


332) Gene Symbol: EVI5

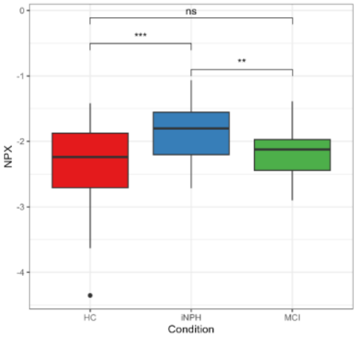


333) Gene Symbol: EXTL1

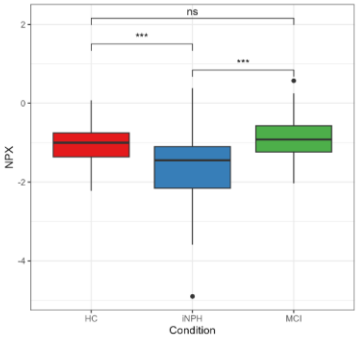


334) Gene Symbol: F11R

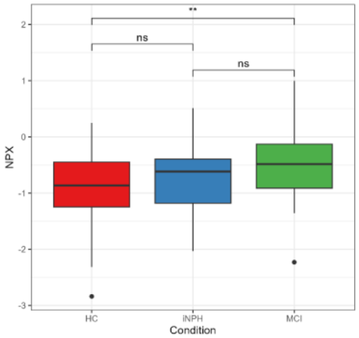


335) Gene Symbol: F2R

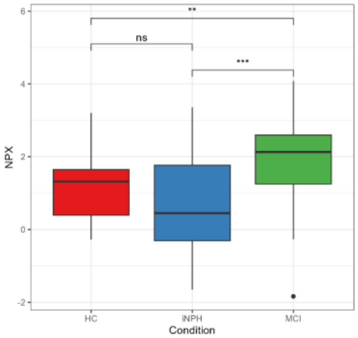


336) Gene Symbol: F3

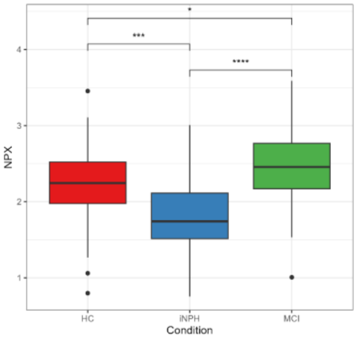


337) Gene Symbol: FABP1

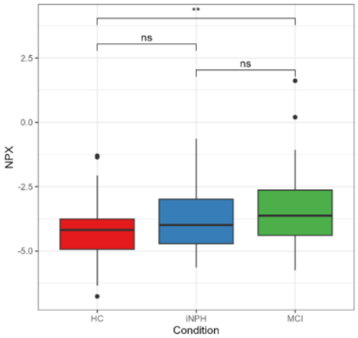


338) Gene Symbol: FABP2

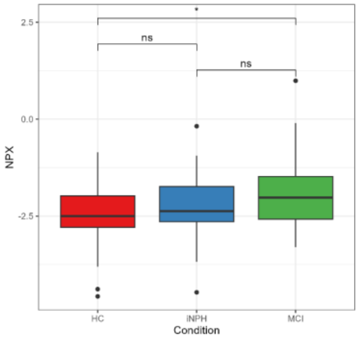


339) Gene Symbol: FABP5

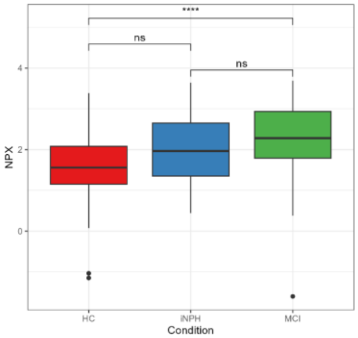


340) Gene Symbol: FAM13A

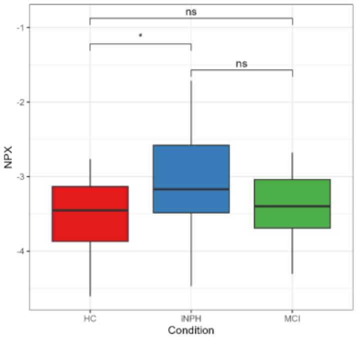

341) Gene Symbol: FAM172A

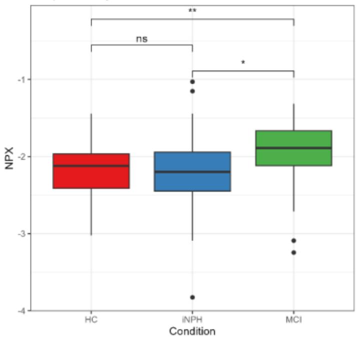


342) Gene Symbol: FAM3C

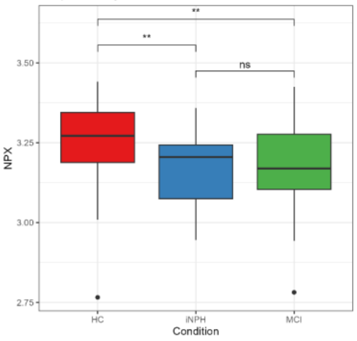


343) Gene Symbol: FCAMR

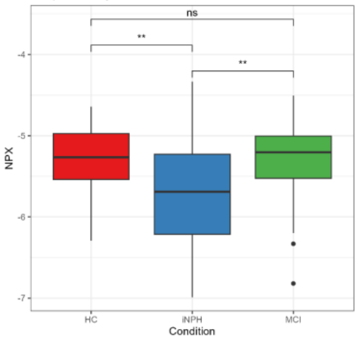


344) Gene Symbol: FCER1A

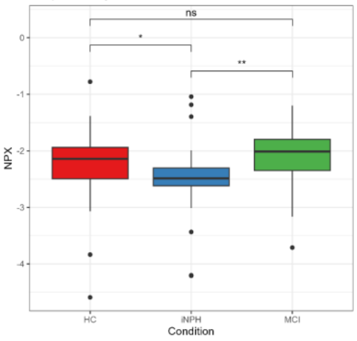


345) Gene Symbol: FCGR3B

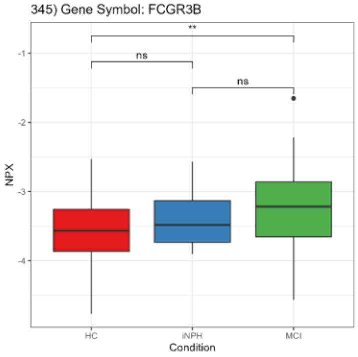


346) Gene Symbol: FCRL6

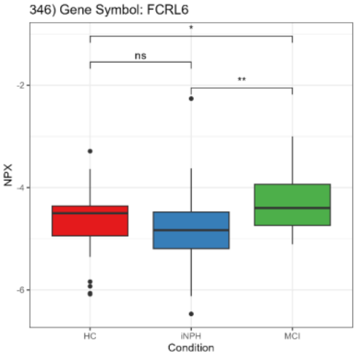


347) Gene Symbol: FDX2

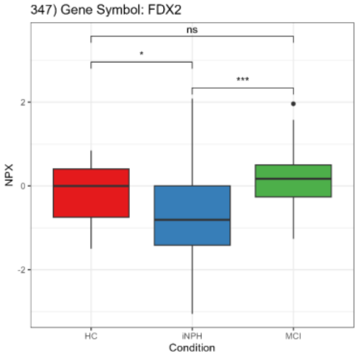


348) Gene Symbol: FGF19

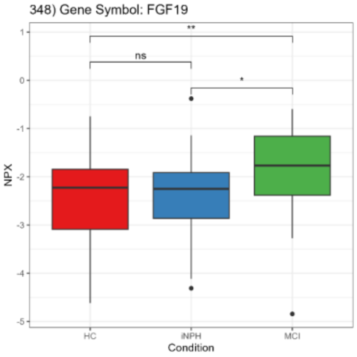


349) Gene Symbol: FGF21

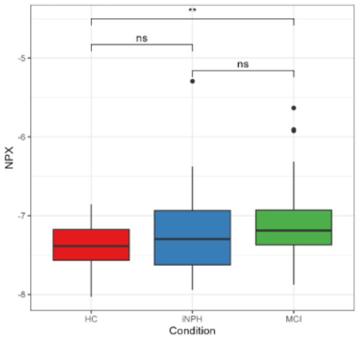


350) Gene Symbol: FGF5

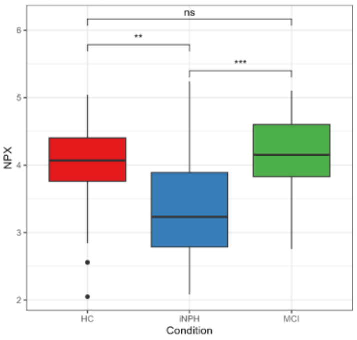


351) Gene Symbol: FGFBP1

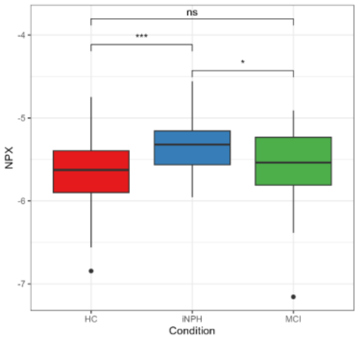


352) Gene Symbol: FGFR2

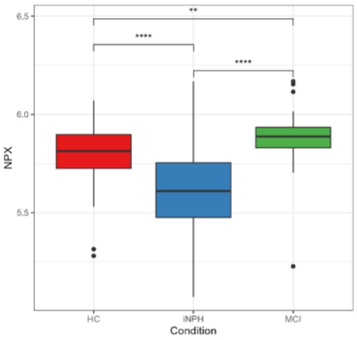


353) Gene Symbol: FHIP2A

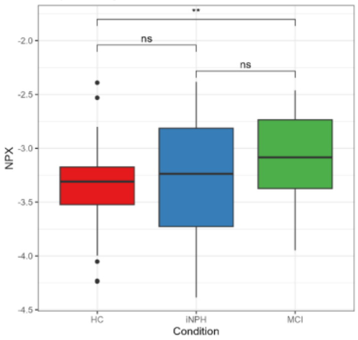


354) Gene Symbol: FHIT

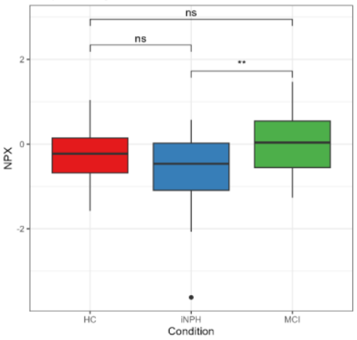


355) Gene Symbol: FKBP14

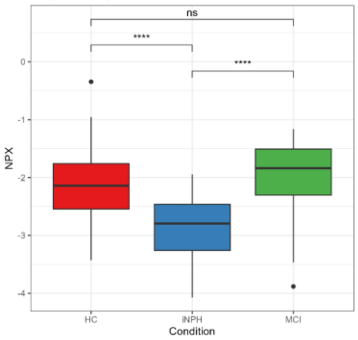


356) Gene Symbol: FKBP7

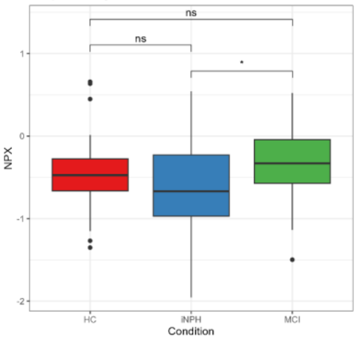


357) Gene Symbol: FLRT2

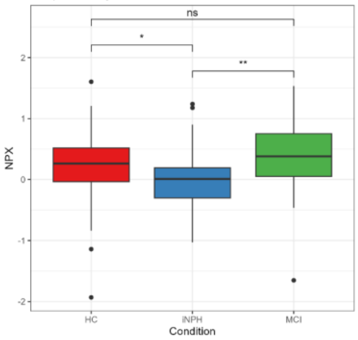


358) Gene Symbol: FLT4

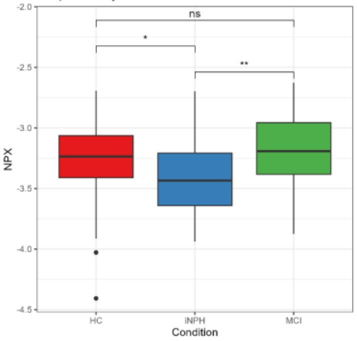


359) Gene Symbol: FMR1

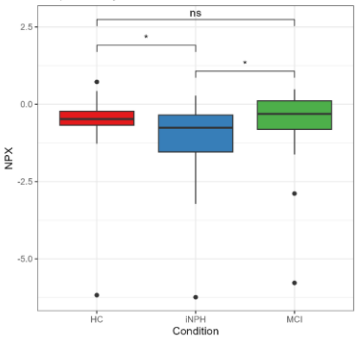


360) Gene Symbol: FOLR2

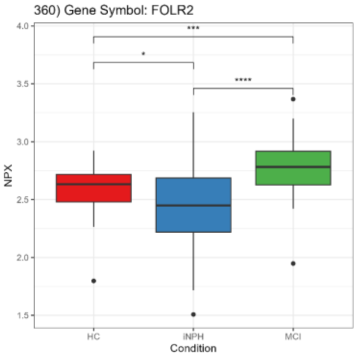

361) Gene Symbol: FOLR3

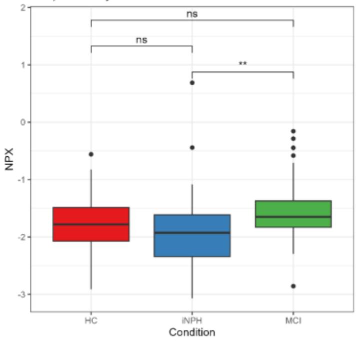


362) Gene Symbol: FSHB

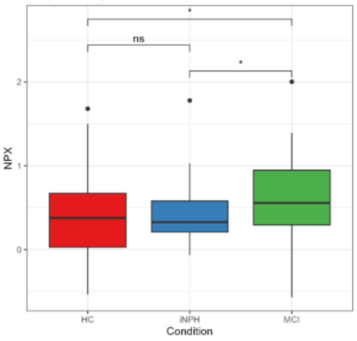


363) Gene Symbol: FST

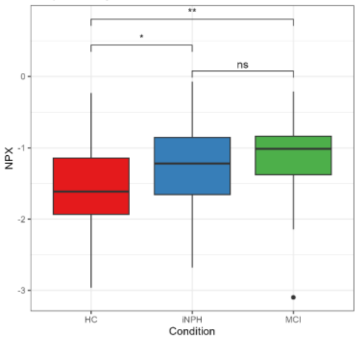


364) Gene Symbol: FSTL3

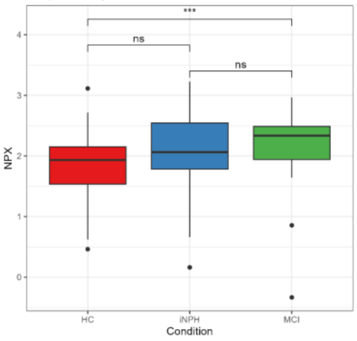


365) Gene Symbol: FUCA1

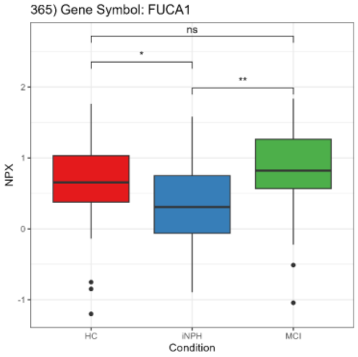


366) Gene Symbol: FURIN

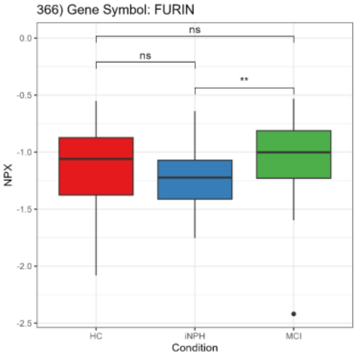


367) Gene Symbol: FUT3_FUT5

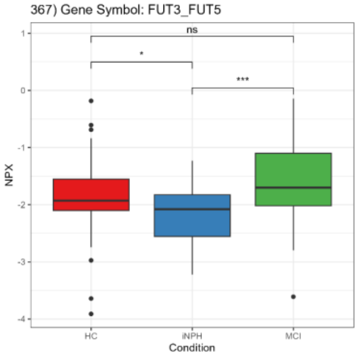


368) Gene Symbol: FXN

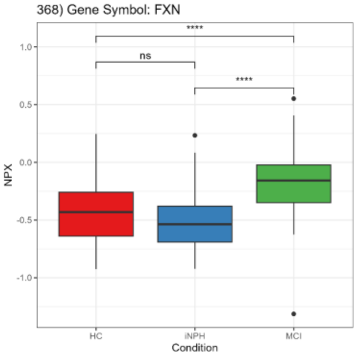


369) Gene Symbol: GAL

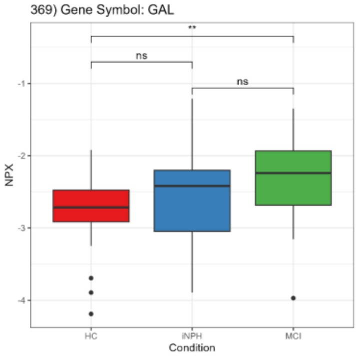


370) Gene Symbol: GALNT10

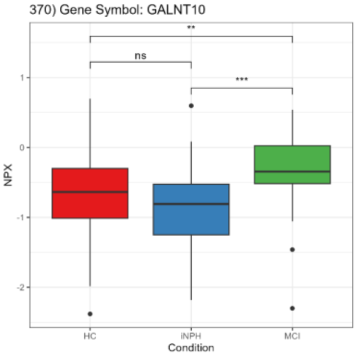


371) Gene Symbol: GALNT2

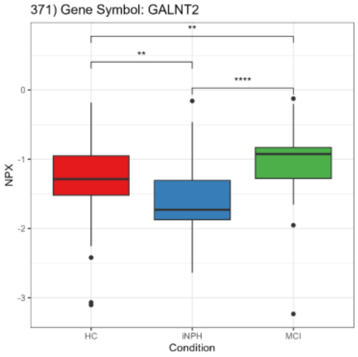


372) Gene Symbol: GALNT3

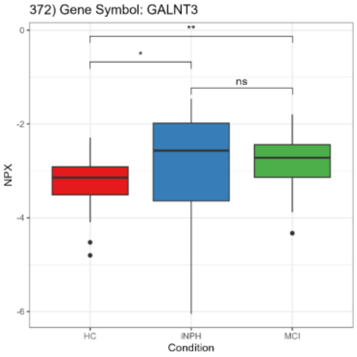


373) Gene Symbol: GALNT7

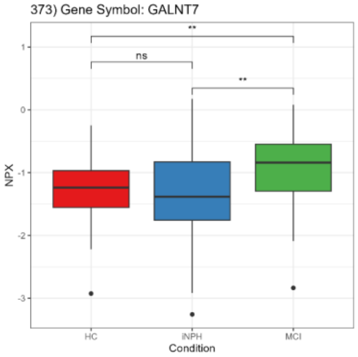


374) Gene Symbol: GAMT

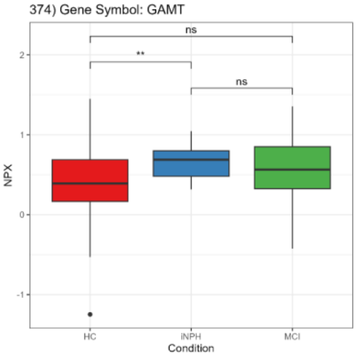


375) Gene Symbol: GAPDH

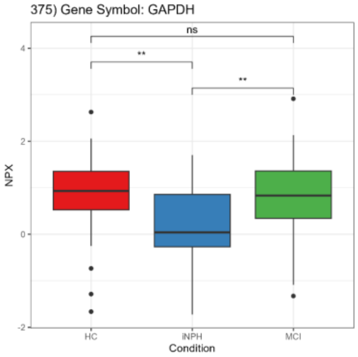


376) Gene Symbol: GAST

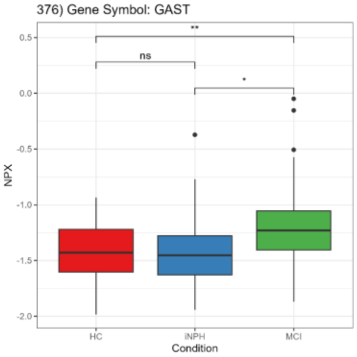


377) Gene Symbol: GATD3

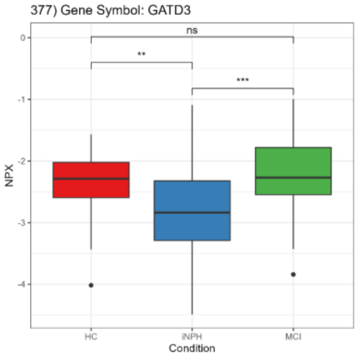


378) Gene Symbol: GBP2

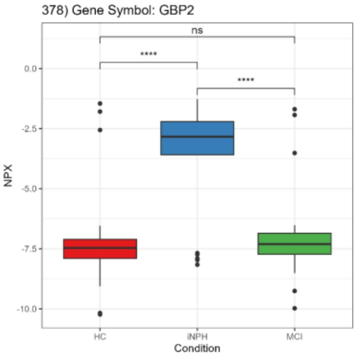


379) Gene Symbol: GCG

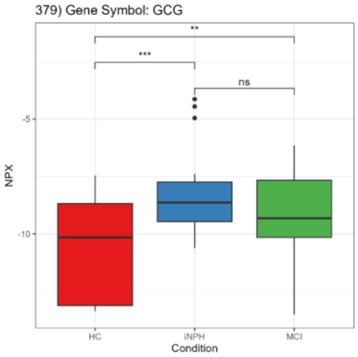


380) Gene Symbol: GCHFR

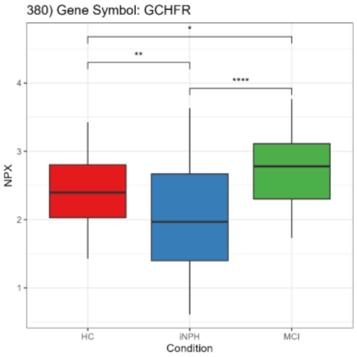

381) Gene Symbol: GCLM

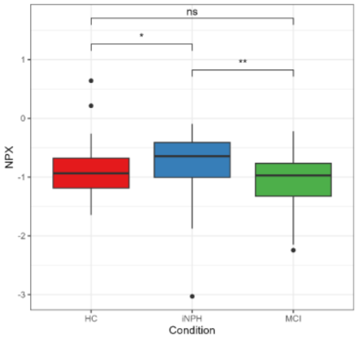


382) Gene Symbol: GCNT1

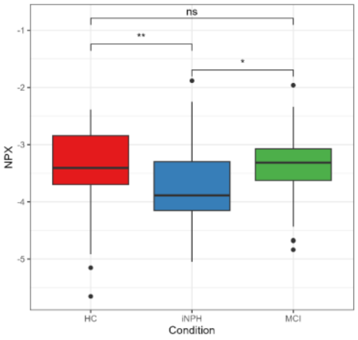


383) Gene Symbol: GFAP

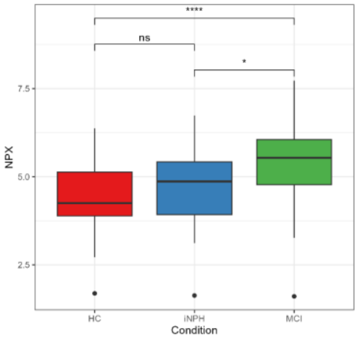


384) Gene Symbol: GFRA1

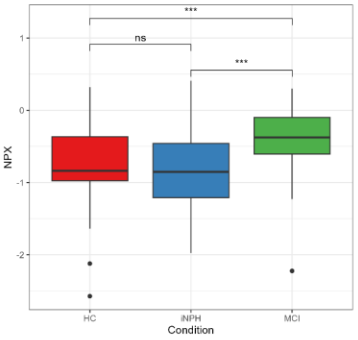


385) Gene Symbol: GFRA2

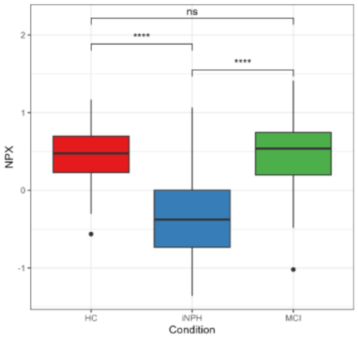


386) Gene Symbol: GFRAL

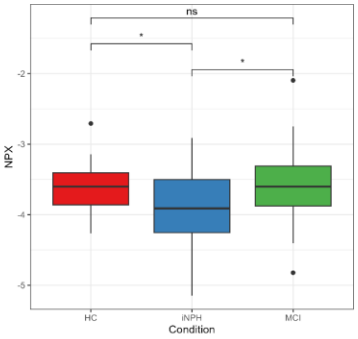


387) Gene Symbol: GGACT

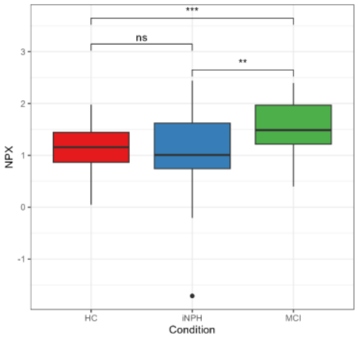


388) Gene Symbol: GGCT

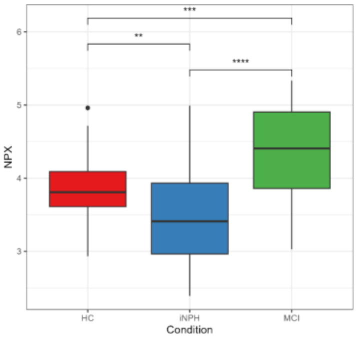


389) Gene Symbol: GHRL

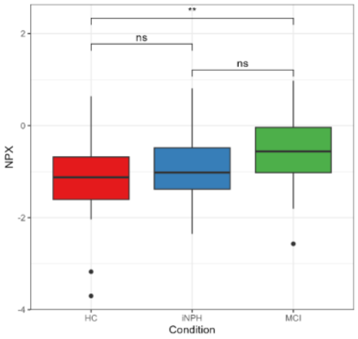


390) Gene Symbol: GIMAP7

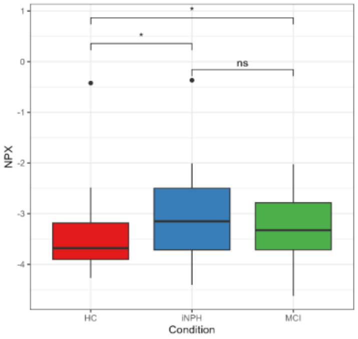


391) Gene Symbol: GIPC2

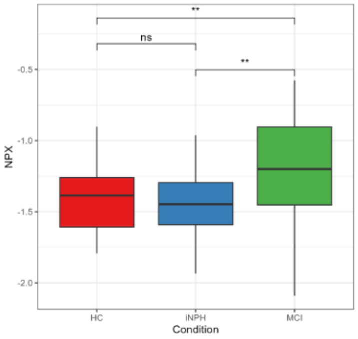


392) Gene Symbol: GIPC3

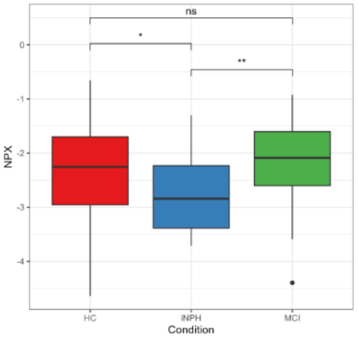


393) Gene Symbol: GKN1

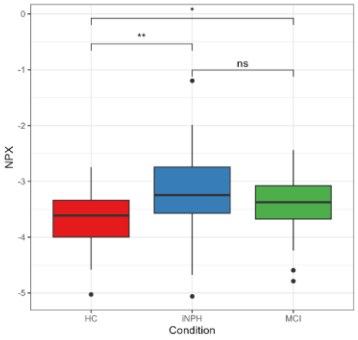


394) Gene Symbol: GLA

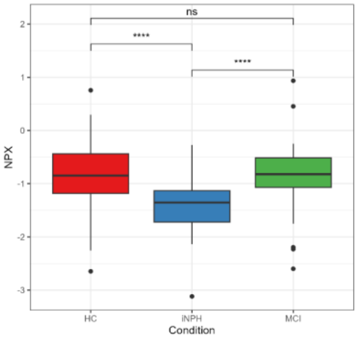


395) Gene Symbol: GLOD4

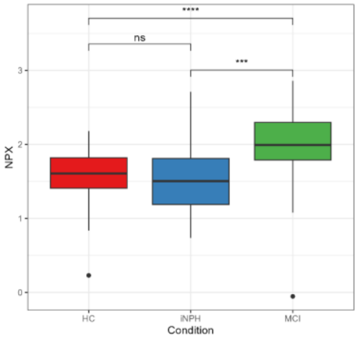


396) Gene Symbol: GLRX

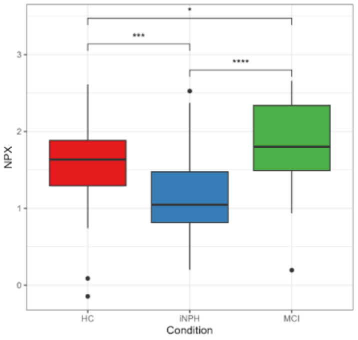


397) Gene Symbol: GM2A

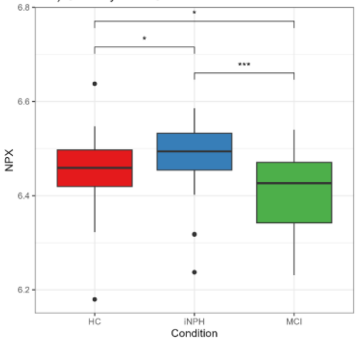


398) Gene Symbol: GMPR2

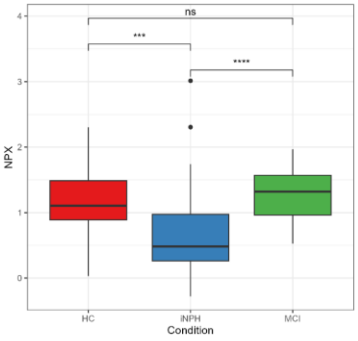


399) Gene Symbol: GOLM2

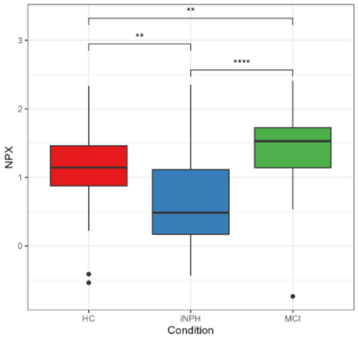


400) Gene Symbol: GOPC

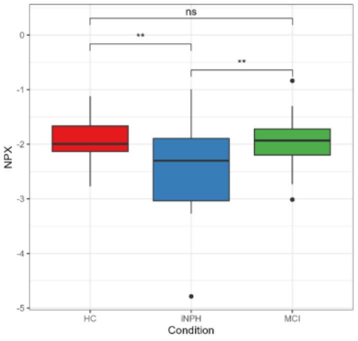

401) Gene Symbol: GOT1

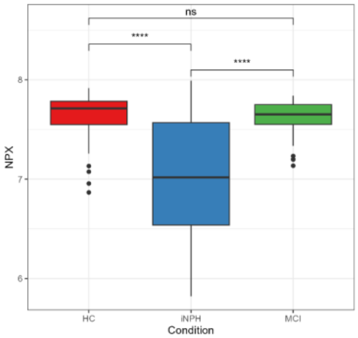


402) Gene Symbol: GP1BB

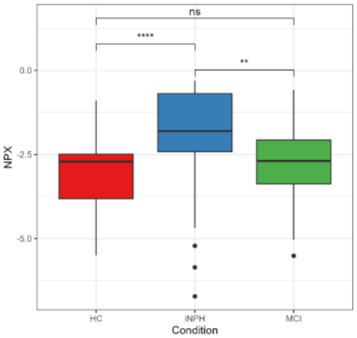


403) Gene Symbol: GPC5

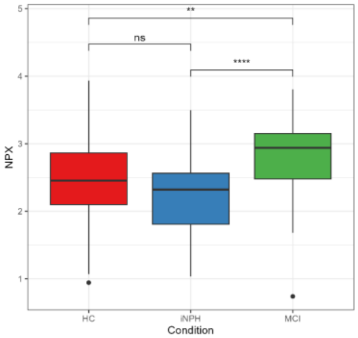


404) Gene Symbol: GPD1

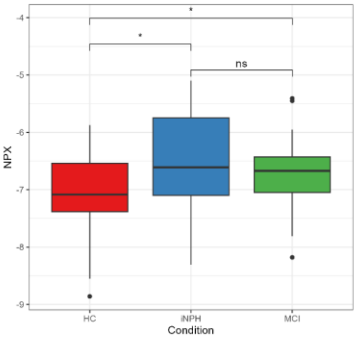


405) Gene Symbol: GPHA2

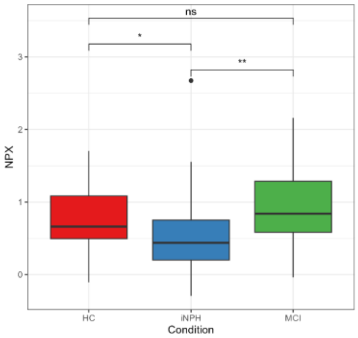


406) Gene Symbol: GPR158

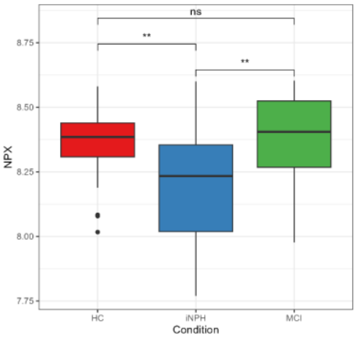


407) Gene Symbol: GPR37

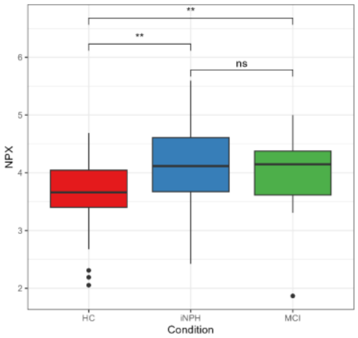


408) Gene Symbol: GRN

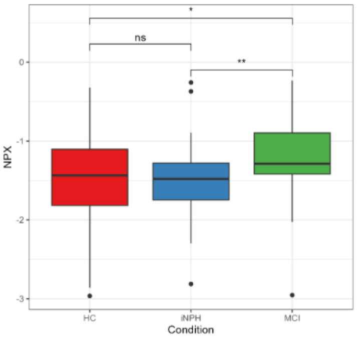


409) Gene Symbol: GRP

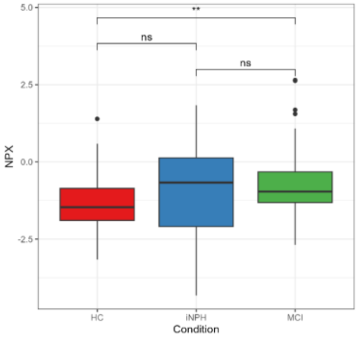


410) Gene Symbol: GSAP

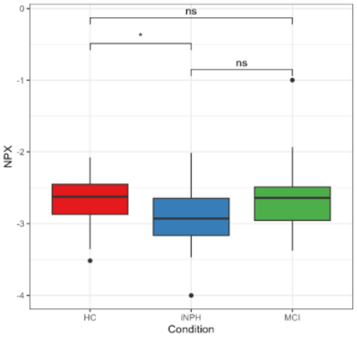


411) Gene Symbol: GSR

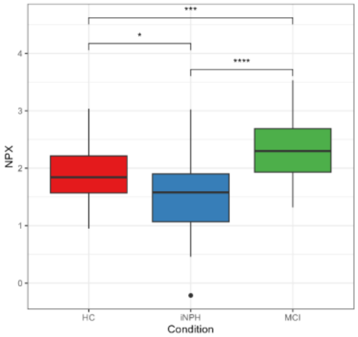


412) Gene Symbol: GZMA

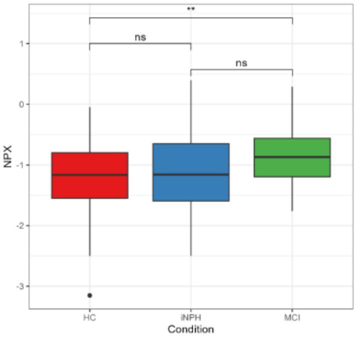


413) Gene Symbol: HAGH

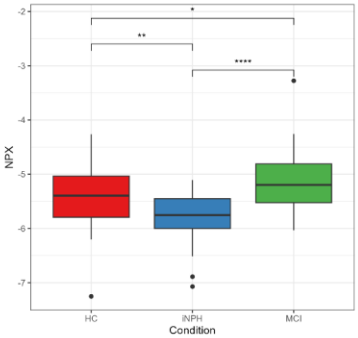


414) Gene Symbol: HARS1

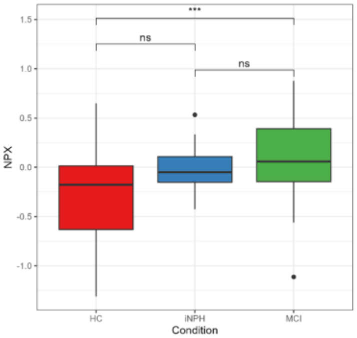


415) Gene Symbol: HAVCR2

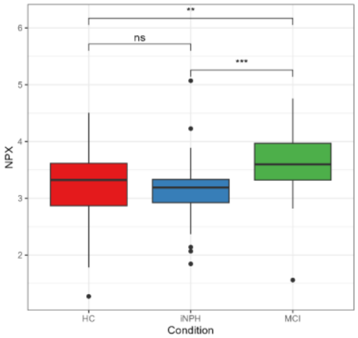


416) Gene Symbol: HBEGF

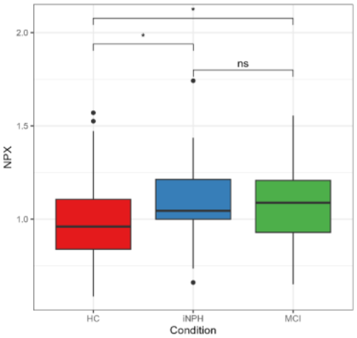


417) Gene Symbol: HDAC8

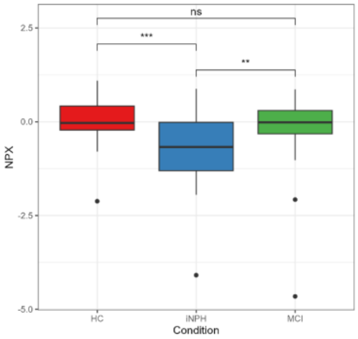


418) Gene Symbol: HGF

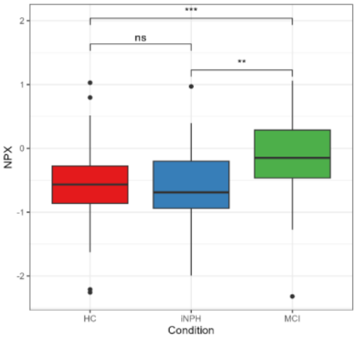


419) Gene Symbol: HHEX

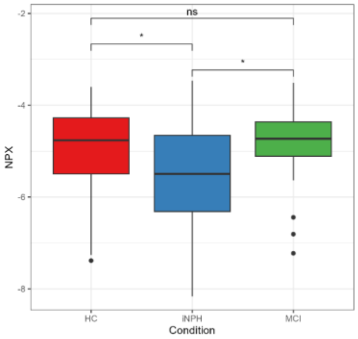


420) Gene Symbol: HK2

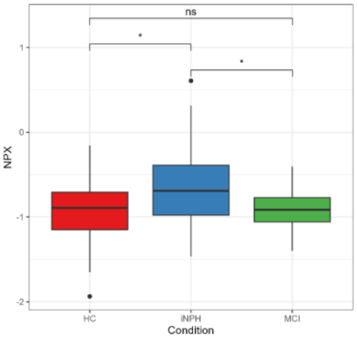

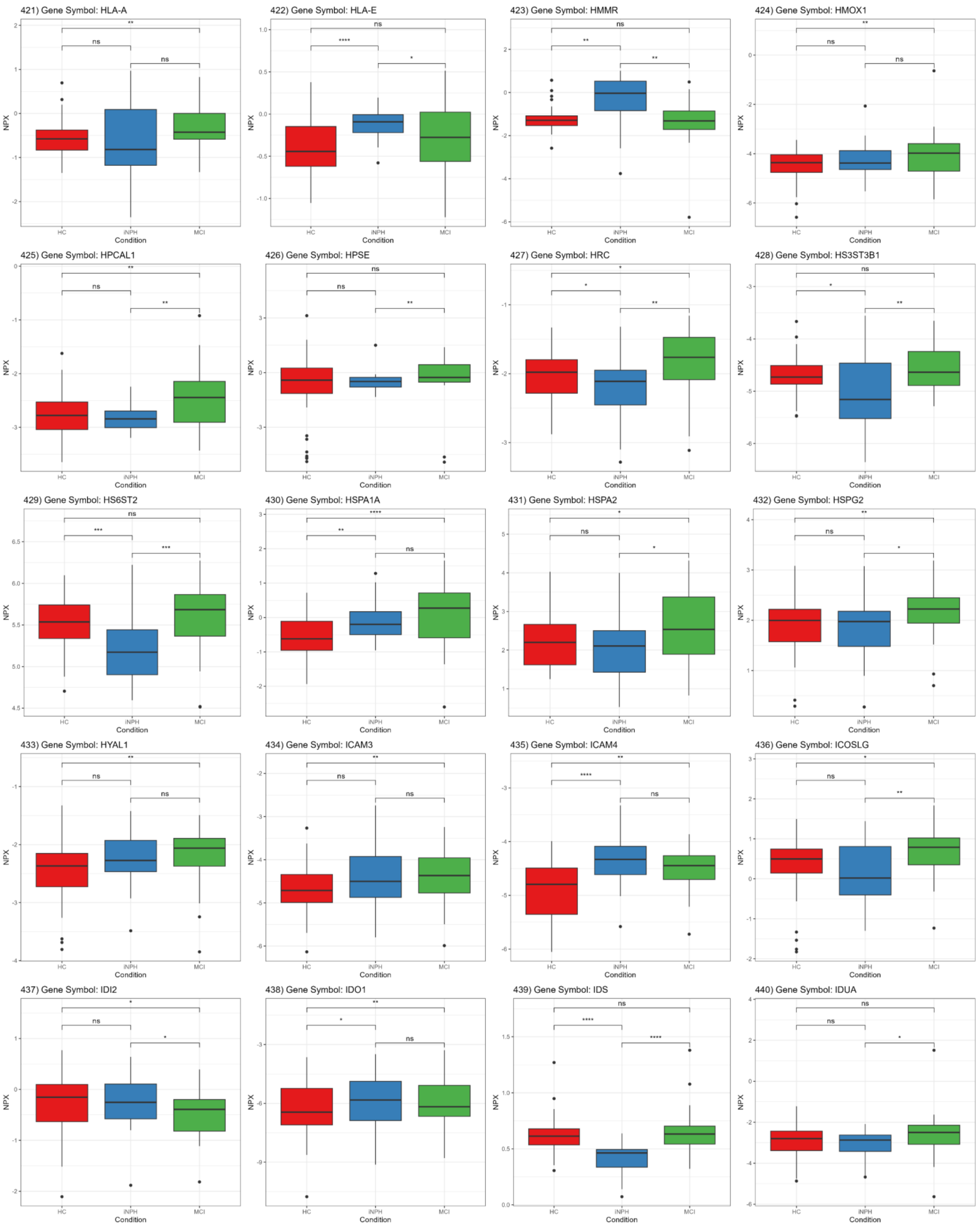
421) Gene Symbol: HLA-A
422) Gene Symbol: HLA-E
423) Gene Symbol: HMMR
424) Gene Symbol: HMOX1
425) Gene Symbol: HPCAL1
426) Gene Symbol: HPSE
427) Gene Symbol: HRC
428) Gene Symbol: HS3ST3B1
429) Gene Symbol: HS6ST2
430) Gene Symbol: HSPA1A
431) Gene Symbol: HSPA2
432) Gene Symbol: HSPG2
433) Gene Symbol: HYAL1
434) Gene Symbol: ICAM3
435) Gene Symbol: ICAM4
436) Gene Symbol: ICOSLG
437) Gene Symbol: IDI2
438) Gene Symbol: IDO1
439) Gene Symbol: IDS
440) Gene Symbol: IDUA
NPX
Condition
HC
iNPH
MCI

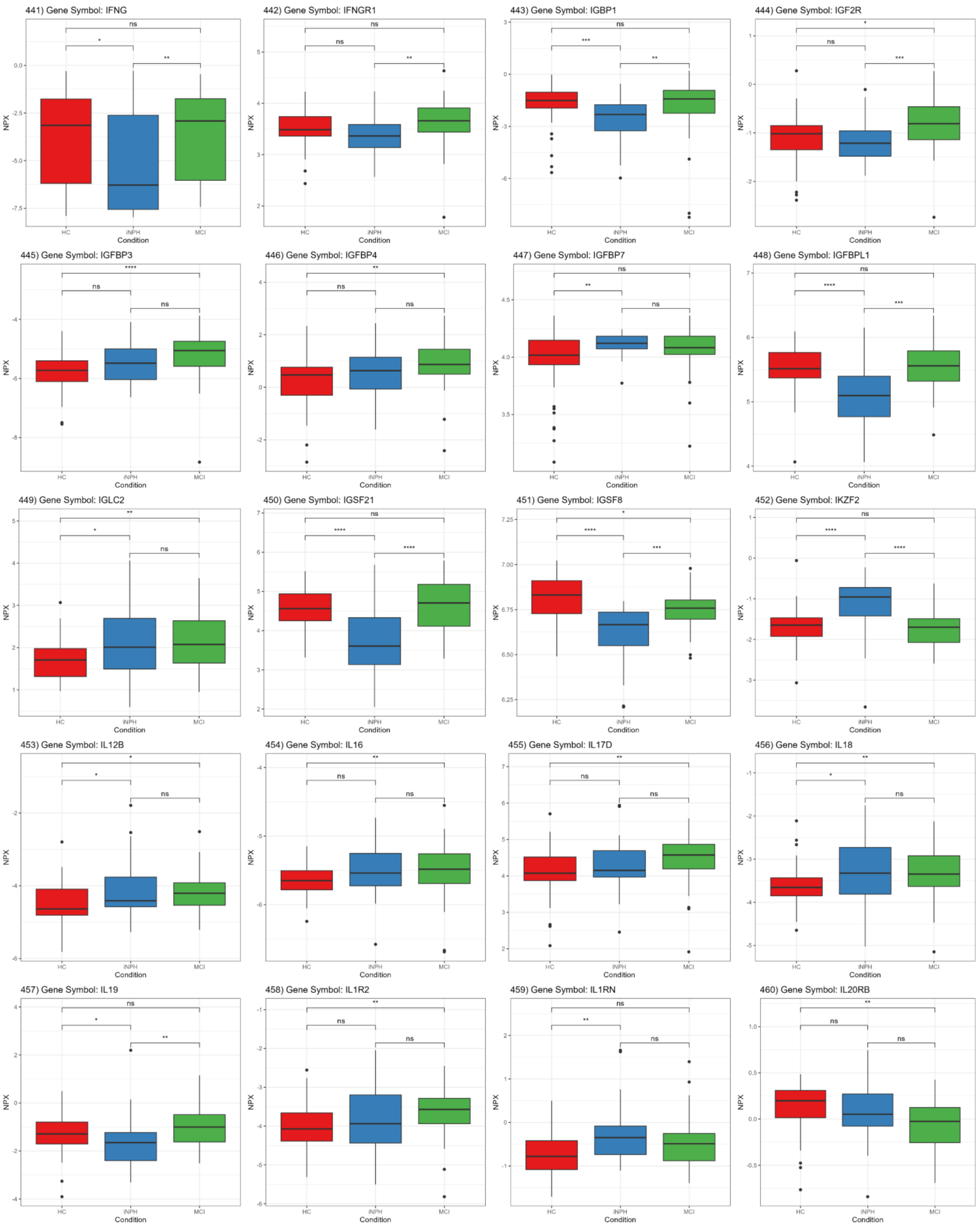

441) Gene Symbol: IFNG
442) Gene Symbol: IFNGR1
443) Gene Symbol: IGBP1
444) Gene Symbol: IGF2R
445) Gene Symbol: IGFBP3
446) Gene Symbol: IGFBP4
447) Gene Symbol: IGFBP7
448) Gene Symbol: IGFBPL1
449) Gene Symbol: IGLC2
450) Gene Symbol: IGSF21
451) Gene Symbol: IGSF8
452) Gene Symbol: IKZF2
453) Gene Symbol: IL12B
454) Gene Symbol: IL16
455) Gene Symbol: IL17D
456) Gene Symbol: IL18
457) Gene Symbol: IL19
458) Gene Symbol: IL1R2
459) Gene Symbol: IL1RN
460) Gene Symbol: IL20RB
NPX
Condition
HC
iNPH
MCI

461) Gene Symbol: IL22RA1

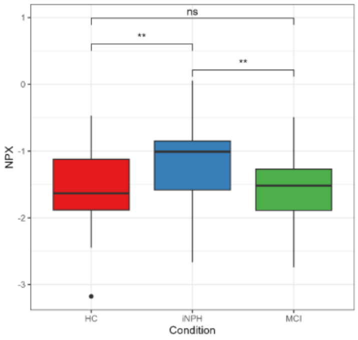


462) Gene Symbol: IL36A

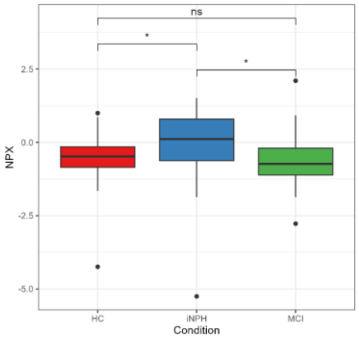


463) Gene Symbol: IL4

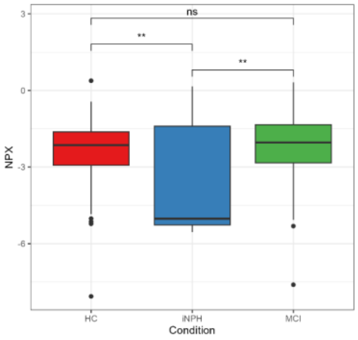


464) Gene Symbol: IL4R

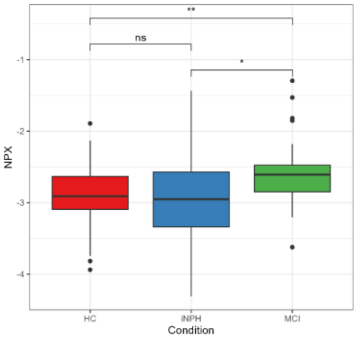


465) Gene Symbol: IL5RA

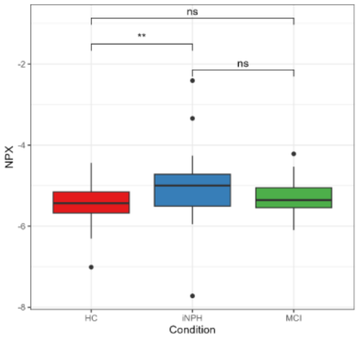


466) Gene Symbol: IL6

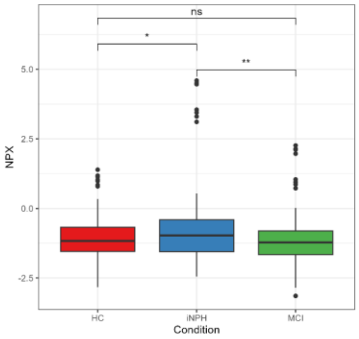


467) Gene Symbol: IL6ST

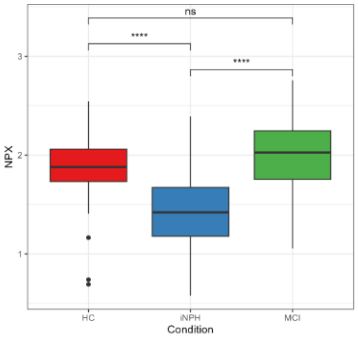


468) Gene Symbol: IMPA1

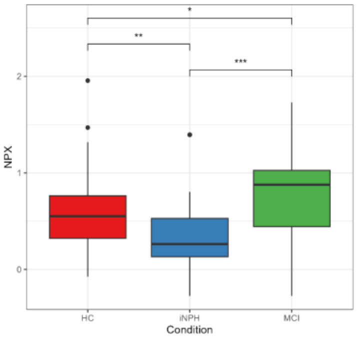


469) Gene Symbol: IMPG1

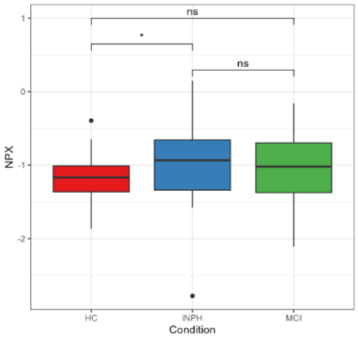


470) Gene Symbol: ING1

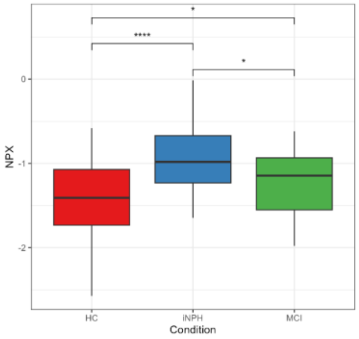


471) Gene Symbol: INHBB

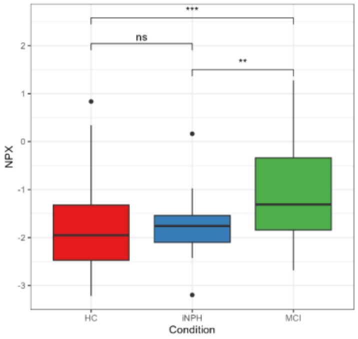


472) Gene Symbol: INPP1

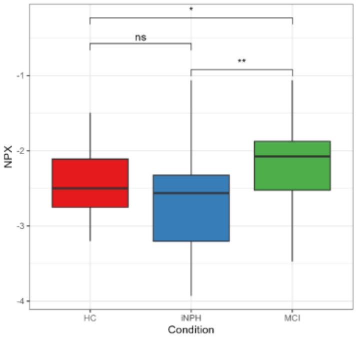


473) Gene Symbol: INPP5D

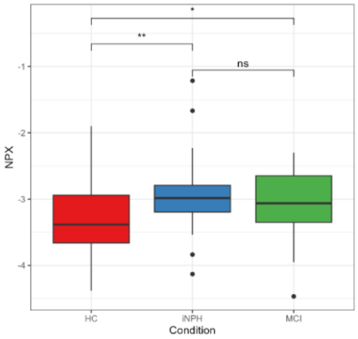


474) Gene Symbol: INSR

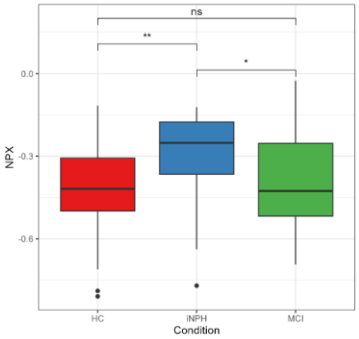


475) Gene Symbol: IRAK4

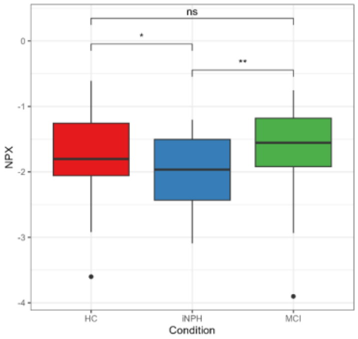


476) Gene Symbol: ISLR2

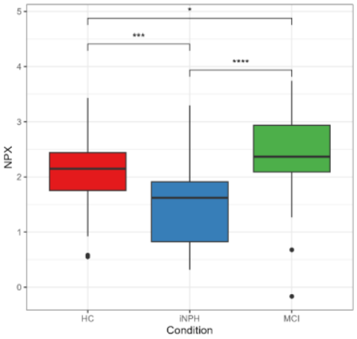


477) Gene Symbol: ITGAV

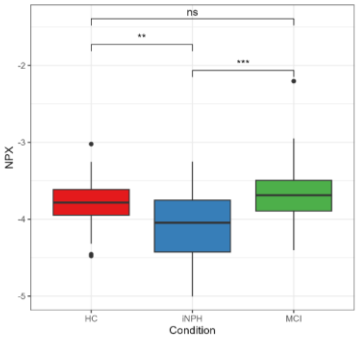


478) Gene Symbol: ITGB5

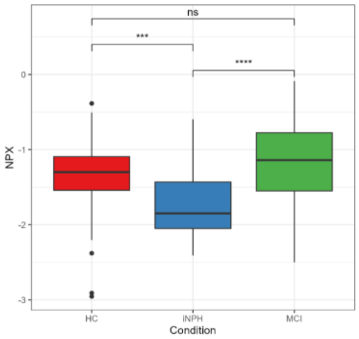


479) Gene Symbol: ITPR1

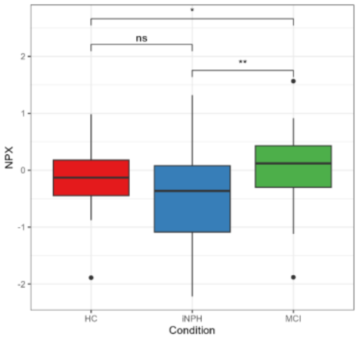


480) Gene Symbol: IVD

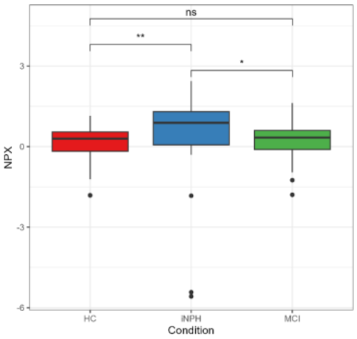

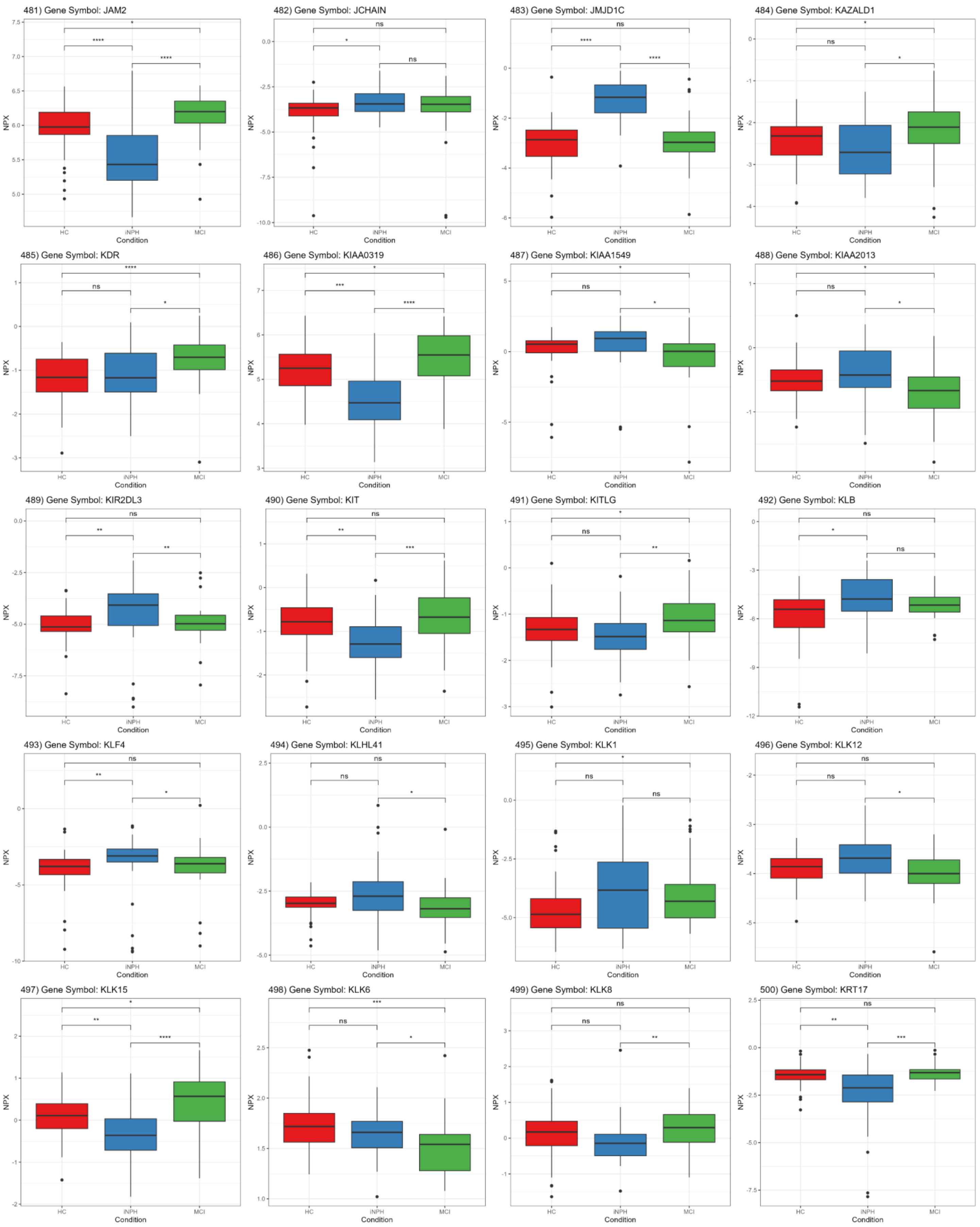
481) Gene Symbol: JAM2
482) Gene Symbol: JCHAIN
483) Gene Symbol: JMJD1C
484) Gene Symbol: KAZALD1
485) Gene Symbol: KDR
486) Gene Symbol: KIAA0319
487) Gene Symbol: KIAA1549
488) Gene Symbol: KIAA2013
489) Gene Symbol: KIR2DL3
490) Gene Symbol: KIT
491) Gene Symbol: KITLG
492) Gene Symbol: KLB
493) Gene Symbol: KLF4
494) Gene Symbol: KLHL41
495) Gene Symbol: KLK1
496) Gene Symbol: KLK12
497) Gene Symbol: KLK15
498) Gene Symbol: KLK6
499) Gene Symbol: KLK8
500) Gene Symbol: KRT17
NPX
Condition
HC
iNPH
MCI

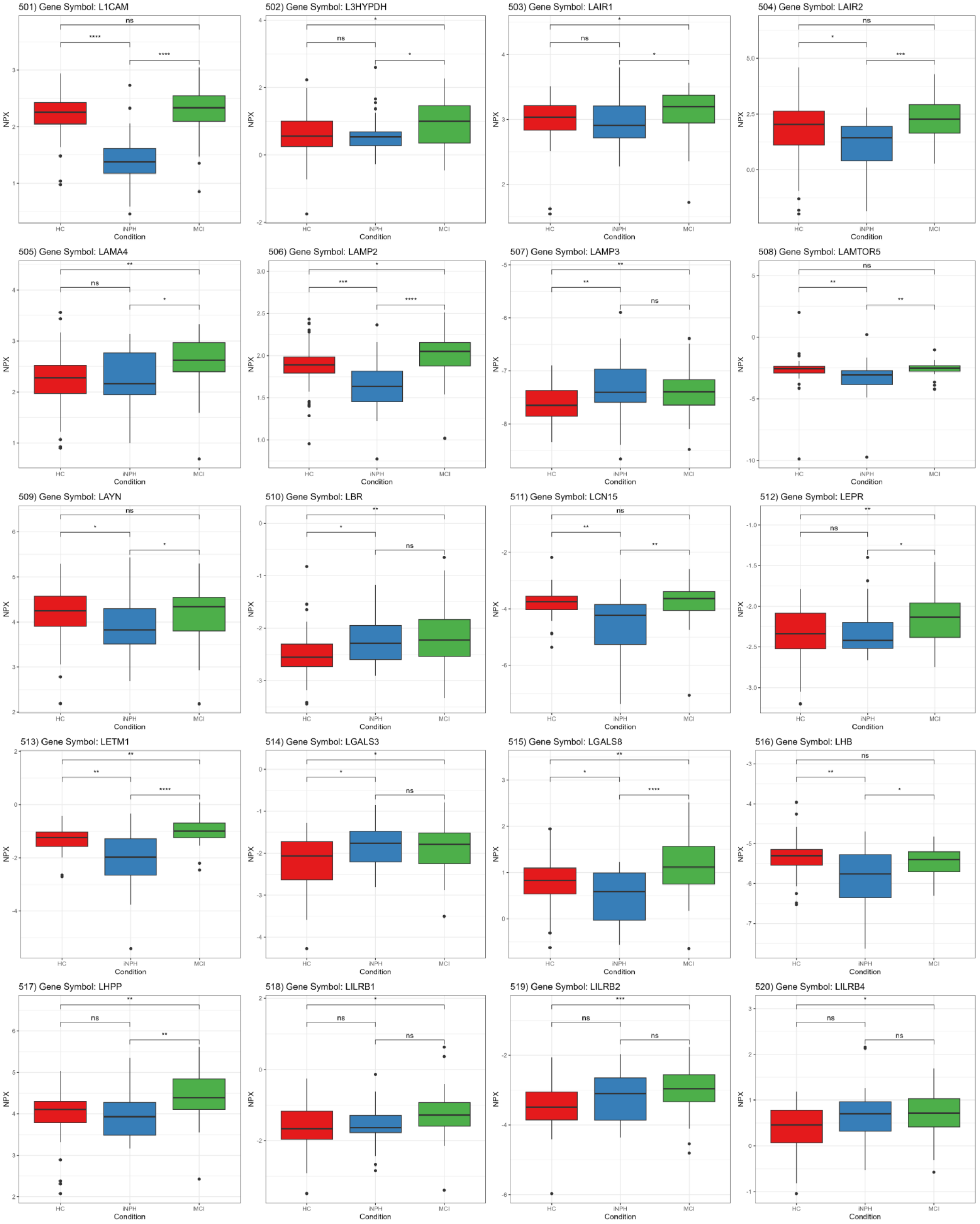
501) Gene Symbol: L1CAM
502) Gene Symbol: L3HYPDH
503) Gene Symbol: LAIR1
504) Gene Symbol: LAIR2
505) Gene Symbol: LAMA4
506) Gene Symbol: LAMP2
507) Gene Symbol: LAMP3
508) Gene Symbol: LAMTOR5
509) Gene Symbol: LAYN
510) Gene Symbol: LBR
511) Gene Symbol: LCN15
512) Gene Symbol: LEPR
513) Gene Symbol: LETM1
514) Gene Symbol: LGALS3
515) Gene Symbol: LGALS8
516) Gene Symbol: LHB
517) Gene Symbol: LHPP
518) Gene Symbol: LILRB1
519) Gene Symbol: LILRB2
520) Gene Symbol: LILRB4
NPX
Condition
HC
iNPH
MCI

521) Gene Symbol: LMOD1

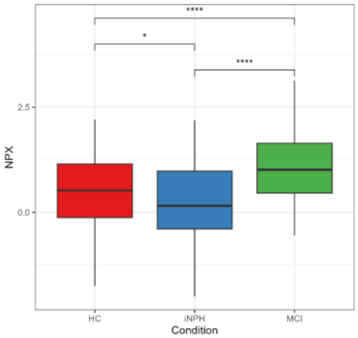


522) Gene Symbol: LMOD2

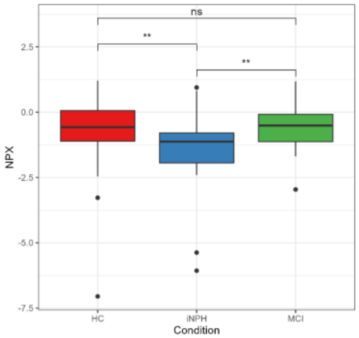


523) Gene Symbol: LPP

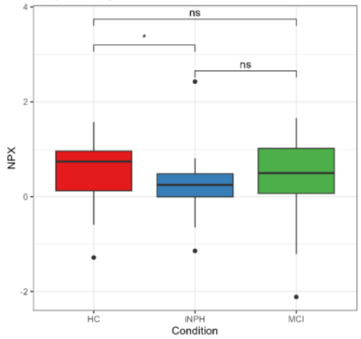


524) Gene Symbol: LRFN2

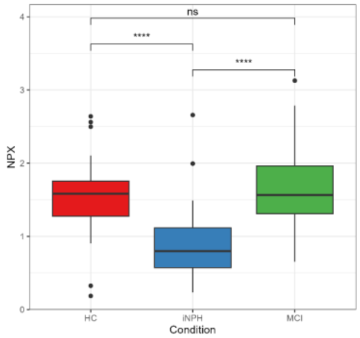


525) Gene Symbol: LRIG1

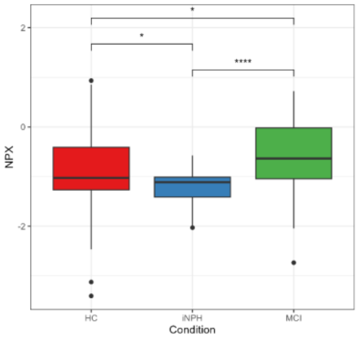


526) Gene Symbol: LRP11

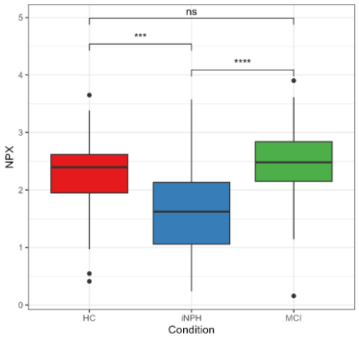


527) Gene Symbol: LRP2BP

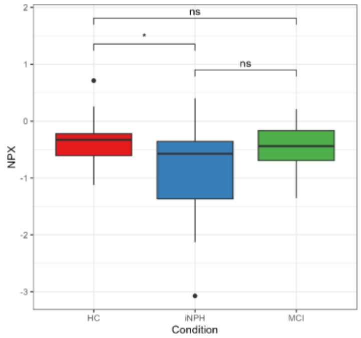


528) Gene Symbol: LRPAP1

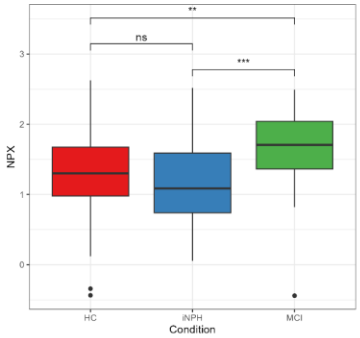


529) Gene Symbol: LRRC25

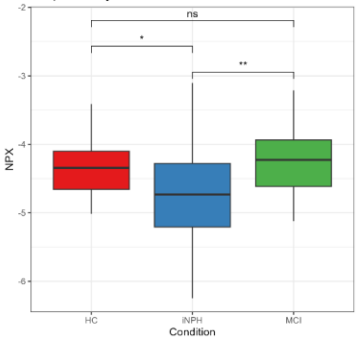


530) Gene Symbol: LRRN1

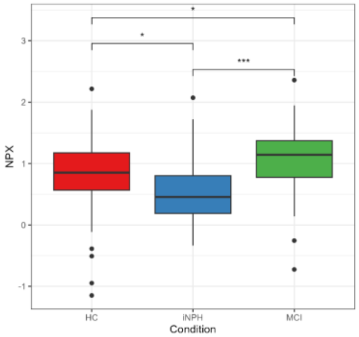


531) Gene Symbol: LSP1

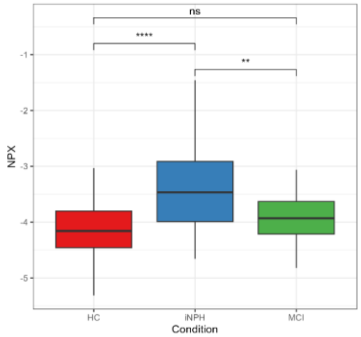


532) Gene Symbol: LTA4H

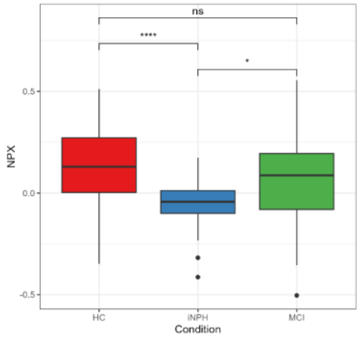


533) Gene Symbol: LTBP2

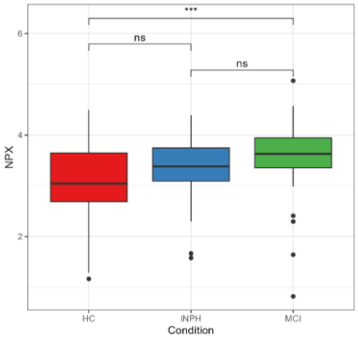


534) Gene Symbol: LTBR

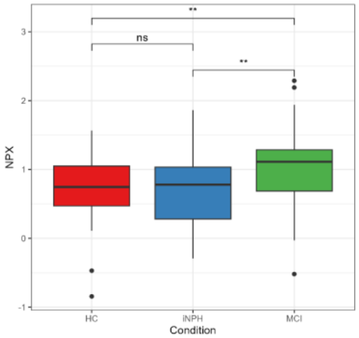


535) Gene Symbol: LY75

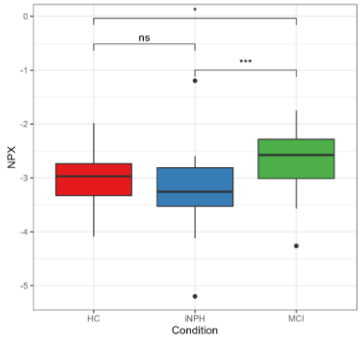


536) Gene Symbol: LYN

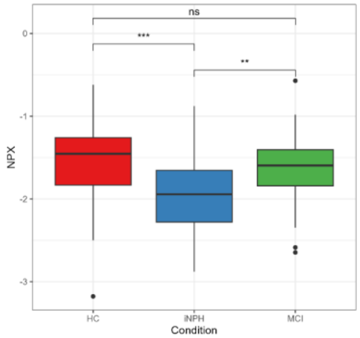


537) Gene Symbol: LYPD3

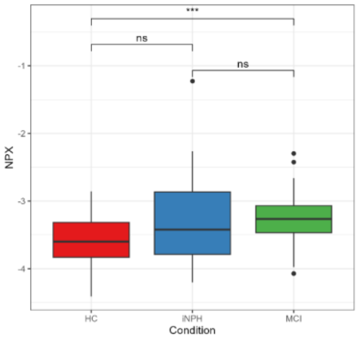


538) Gene Symbol: M6PR

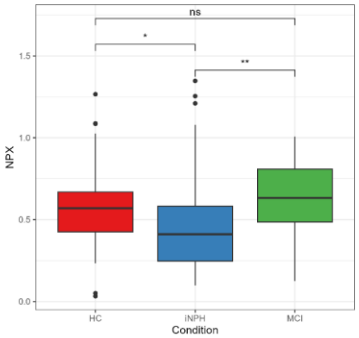


539) Gene Symbol: MAD1L1

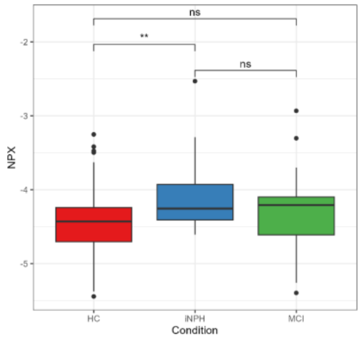


540) Gene Symbol: MAEA

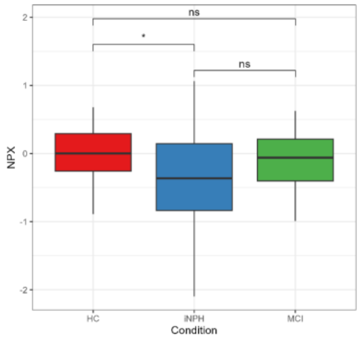

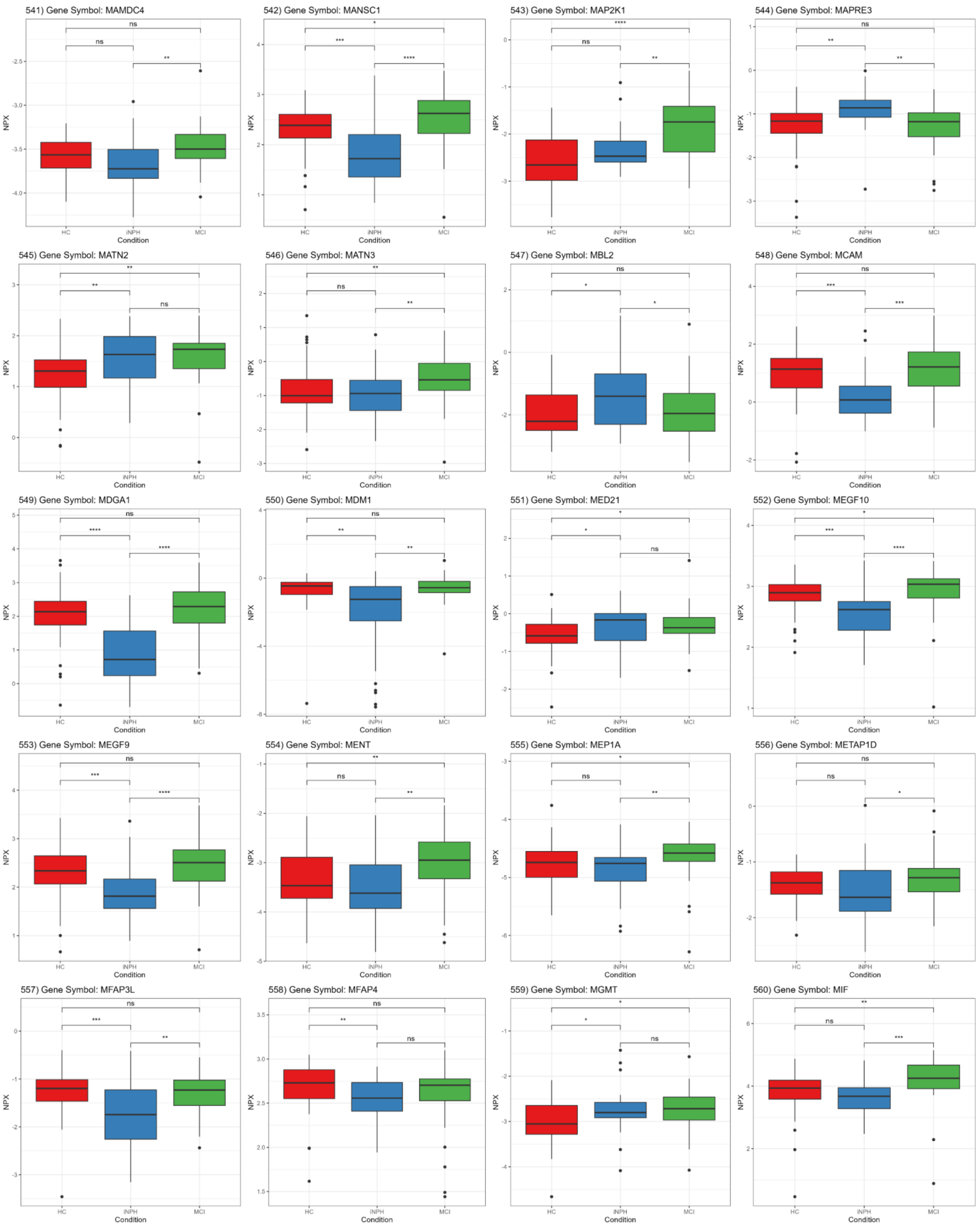
541) Gene Symbol: MAMDC4
542) Gene Symbol: MANSC1
543) Gene Symbol: MAP2K1
544) Gene Symbol: MAPRE3
545) Gene Symbol: MATN2
546) Gene Symbol: MATN3
547) Gene Symbol: MBL2
548) Gene Symbol: MCAM
549) Gene Symbol: MDGA1
550) Gene Symbol: MDM1
551) Gene Symbol: MED21
552) Gene Symbol: MEGF10
553) Gene Symbol: MEGF9
554) Gene Symbol: MENT
555) Gene Symbol: MEP1A
556) Gene Symbol: METAP1D
557) Gene Symbol: MFAP3L
558) Gene Symbol: MFAP4
559) Gene Symbol: MGMT
560) Gene Symbol: MIF
NPX
Condition
HC
iNPH
MCI

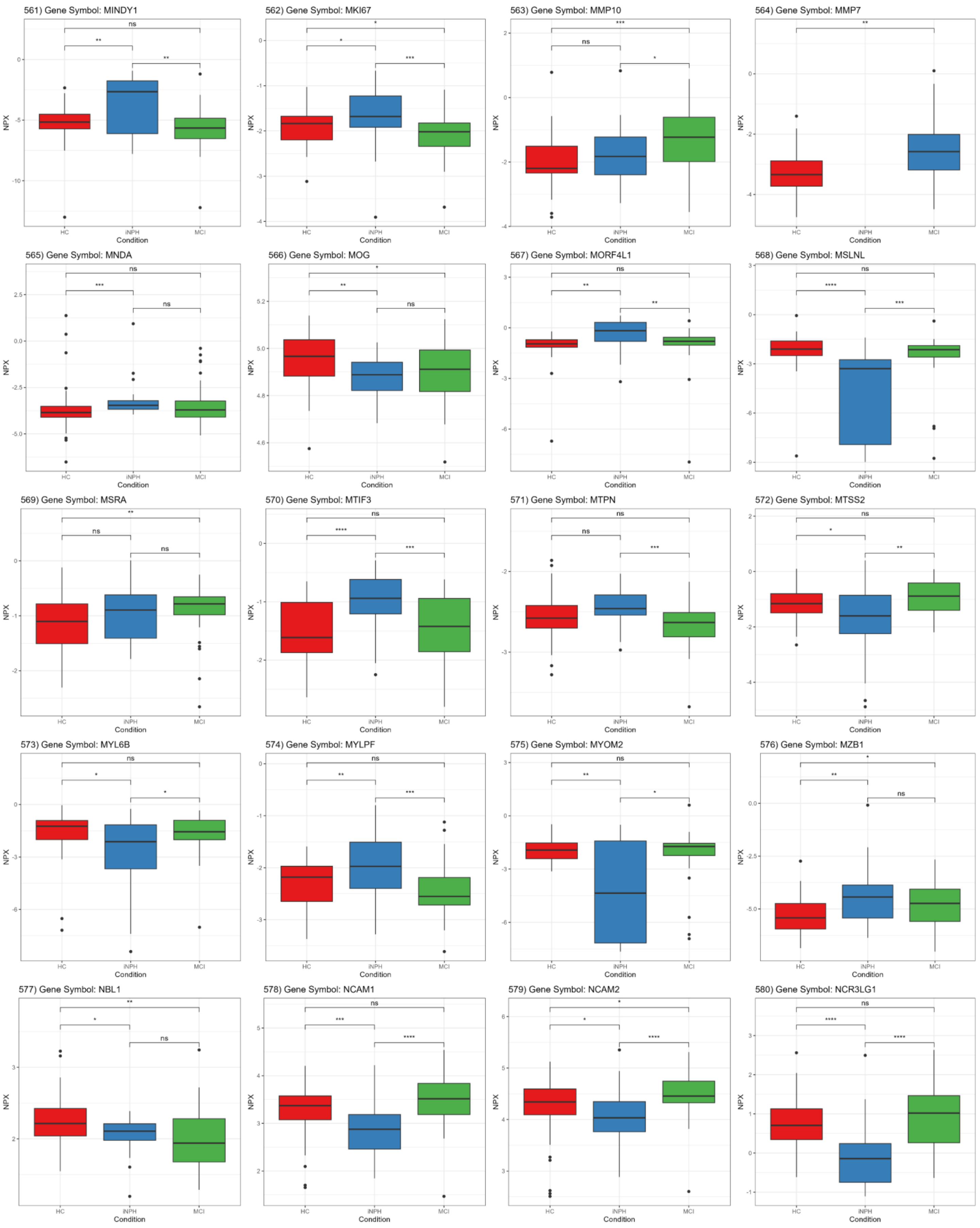
561) Gene Symbol: MINDY1
562) Gene Symbol: MKI67
563) Gene Symbol: MMP10
564) Gene Symbol: MMP7
565) Gene Symbol: MNDA
566) Gene Symbol: MOG
567) Gene Symbol: MORF4L1
568) Gene Symbol: MSLNL
569) Gene Symbol: MSRA
570) Gene Symbol: MTIF3
571) Gene Symbol: MTPN
572) Gene Symbol: MTSS2
573) Gene Symbol: MYL6B
574) Gene Symbol: MYLPF
575) Gene Symbol: MYOM2
576) Gene Symbol: MZB1
577) Gene Symbol: NBL1
578) Gene Symbol: NCAM1
579) Gene Symbol: NCAM2
580) Gene Symbol: NCR3LG1
NPX
Condition
HC
iNPH
MCI

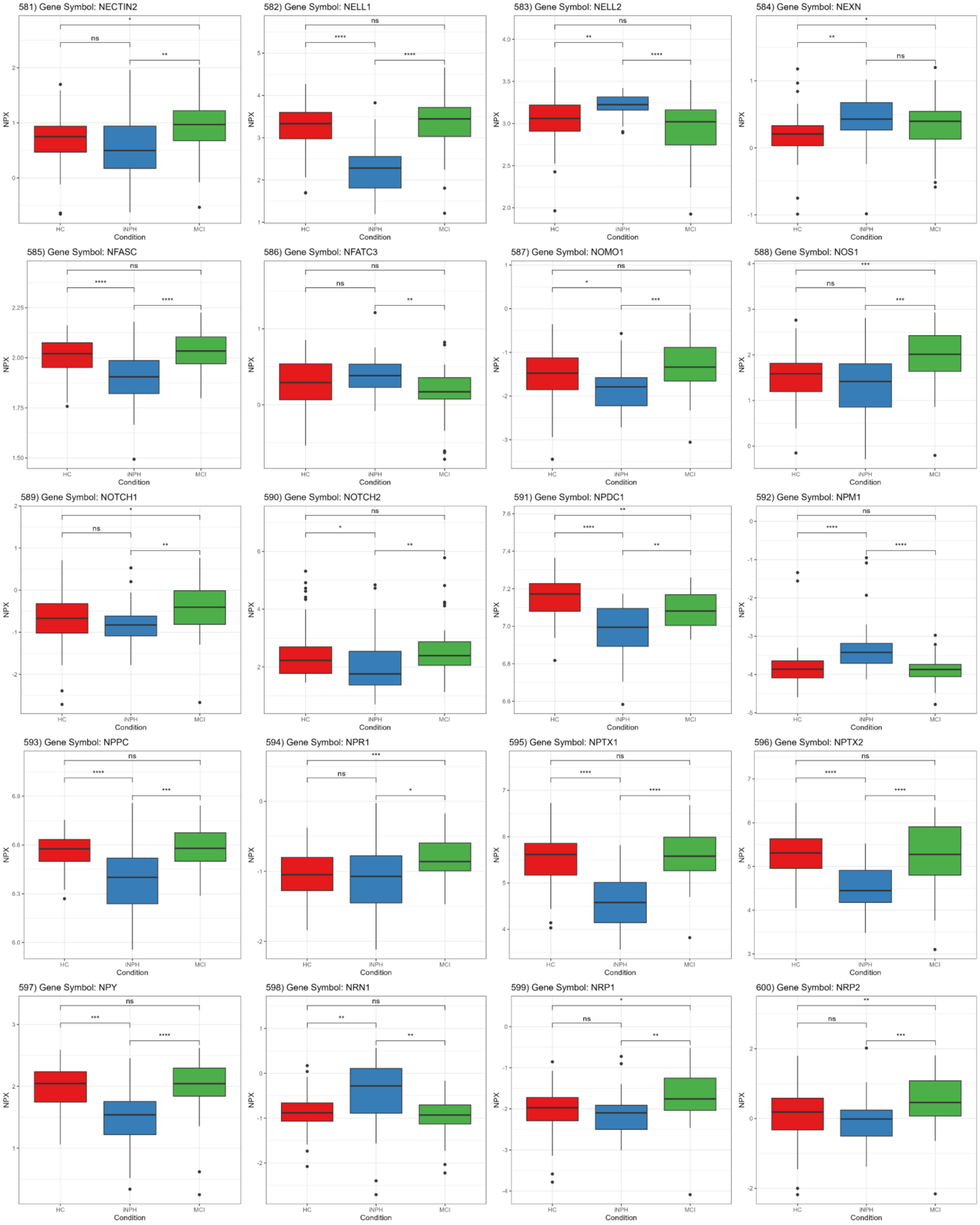
581) Gene Symbol: NECTIN2
582) Gene Symbol: NELL1
583) Gene Symbol: NELL2
584) Gene Symbol: NEXN
585) Gene Symbol: NFASC
586) Gene Symbol: NFATC3
587) Gene Symbol: NOMO1
588) Gene Symbol: NOS1
589) Gene Symbol: NOTCH1
590) Gene Symbol: NOTCH2
591) Gene Symbol: NPDC1
592) Gene Symbol: NPM1
593) Gene Symbol: NPPC
594) Gene Symbol: NPR1
595) Gene Symbol: NPTX1
596) Gene Symbol: NPTX2
597) Gene Symbol: NPY
598) Gene Symbol: NRN1
599) Gene Symbol: NRP1
600) Gene Symbol: NRP2
NPX
Condition
HC
iNPH
MCI

601) Gene Symbol: NRXN3

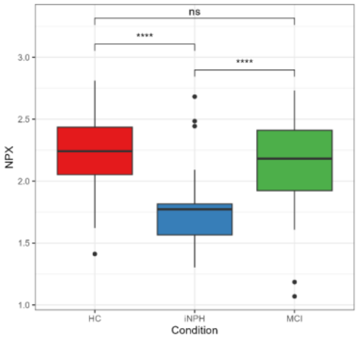


602) Gene Symbol: NSFL1C

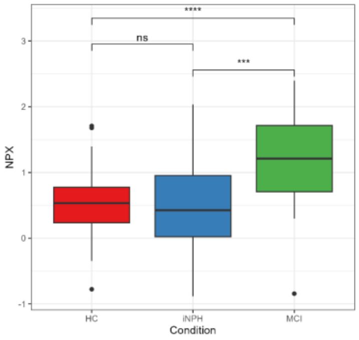


603) Gene Symbol: NT-proBNP

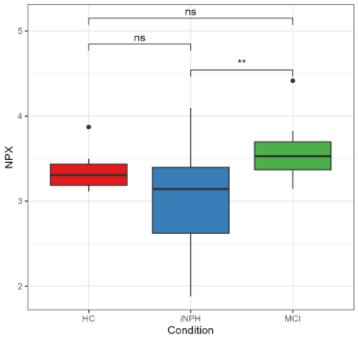


604) Gene Symbol: NT5C1A

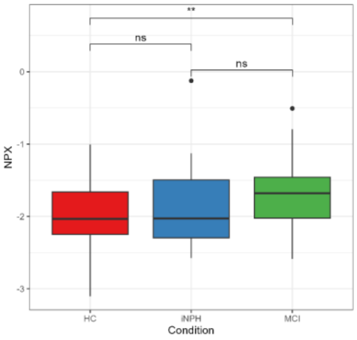


605) Gene Symbol: NTproBNP

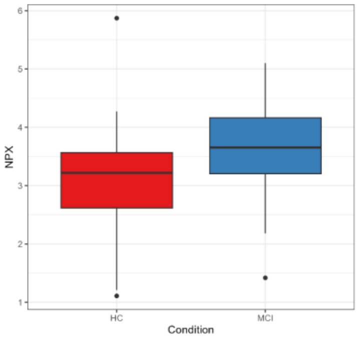


606) Gene Symbol: NTRK2

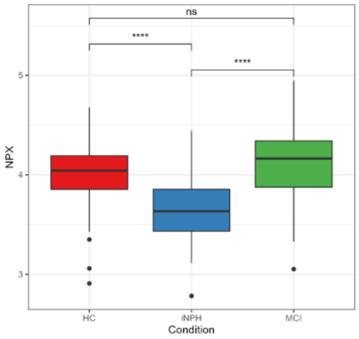


607) Gene Symbol: NTRK3

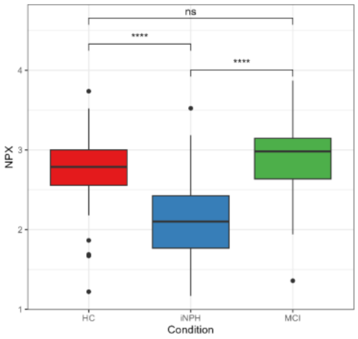


608) Gene Symbol: NUBP1

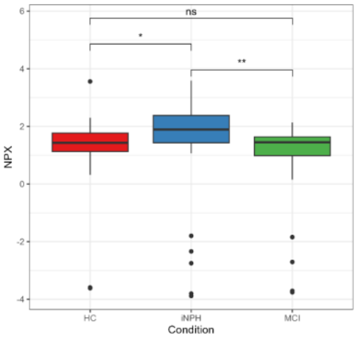


609) Gene Symbol: NUCB2

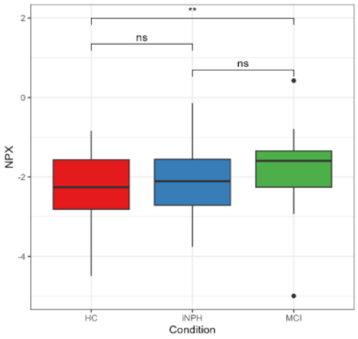


610) Gene Symbol: NUDT16

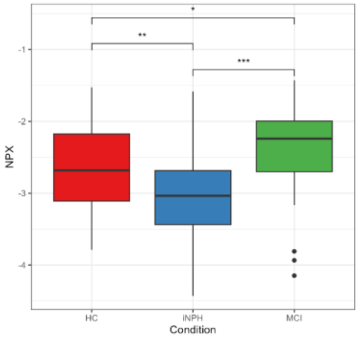


611) Gene Symbol: NUDT5

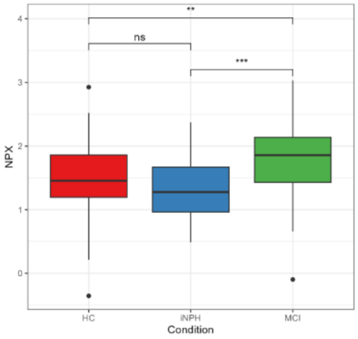


612) Gene Symbol: NXPH1

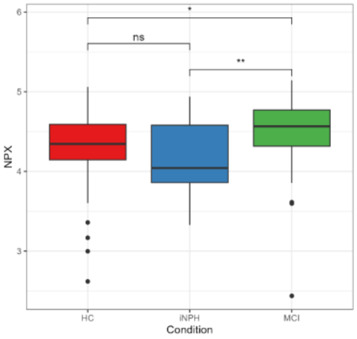


613) Gene Symbol: OGN

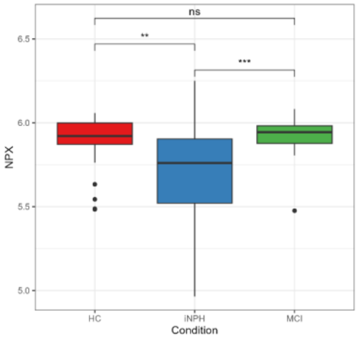


614) Gene Symbol: OLFM4

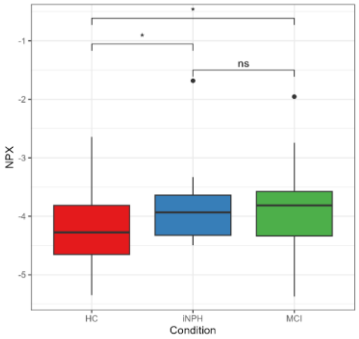


615) Gene Symbol: OLR1

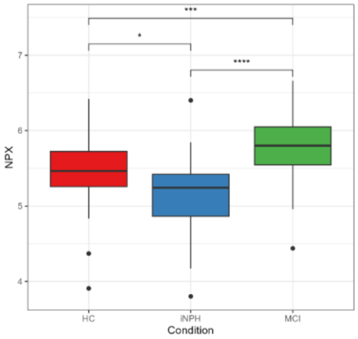


616) Gene Symbol: OMD

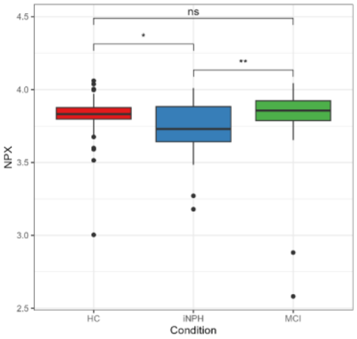


617) Gene Symbol: OMG

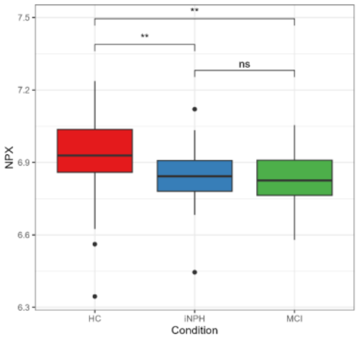


618) Gene Symbol: OMP

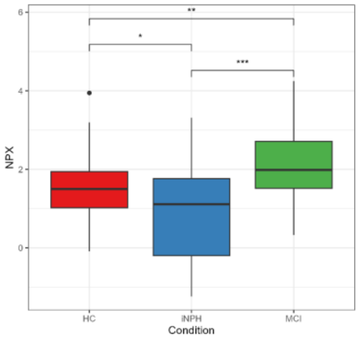


619) Gene Symbol: OPHN1

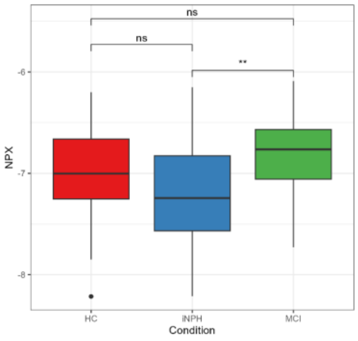


620) Gene Symbol: OPTC

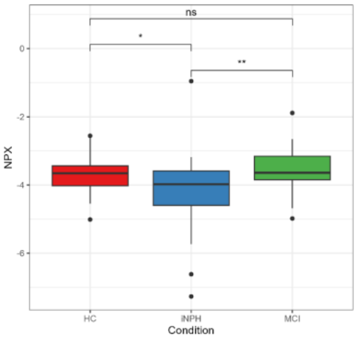

621) Gene Symbol: OSCAR

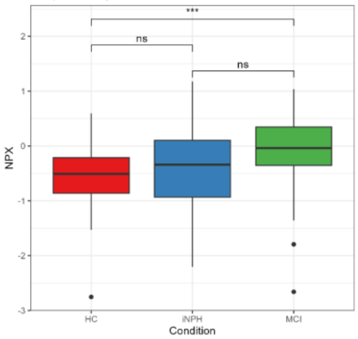


622) Gene Symbol: OSM

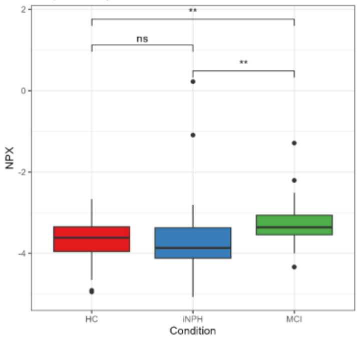


623) Gene Symbol: OSTN

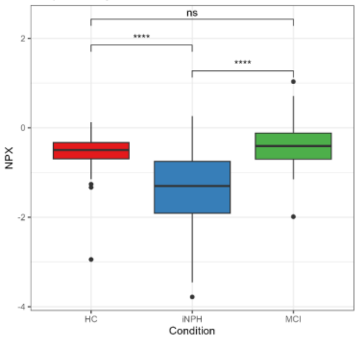


624) Gene Symbol: OTOA

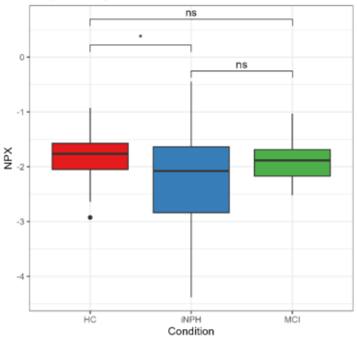


625) Gene Symbol: OXCT1

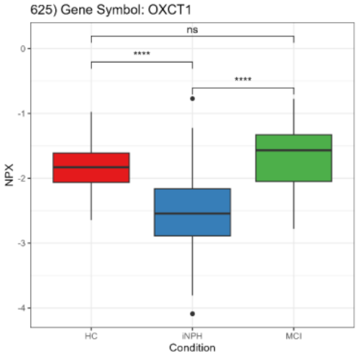


626) Gene Symbol: OXT

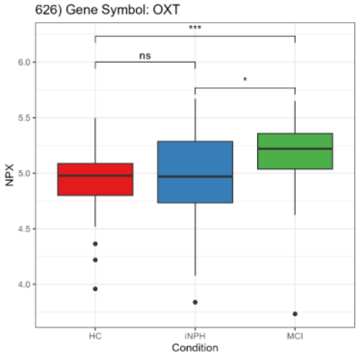


627) Gene Symbol: P4HB

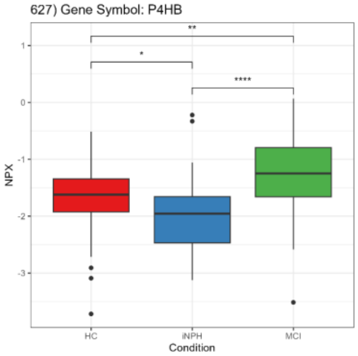


628) Gene Symbol: PACS2

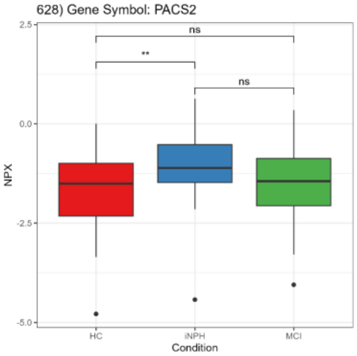


629) Gene Symbol: PADI4

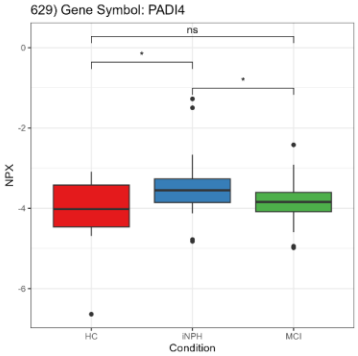


630) Gene Symbol: PAG1

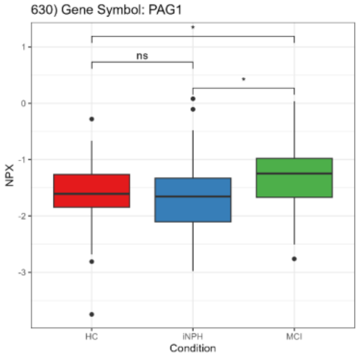


631) Gene Symbol: PALM

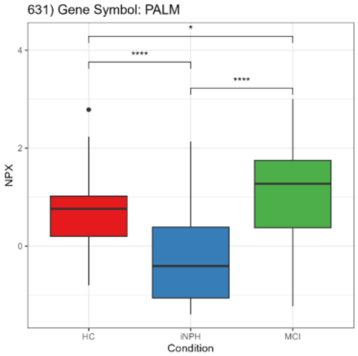


632) Gene Symbol: PAM

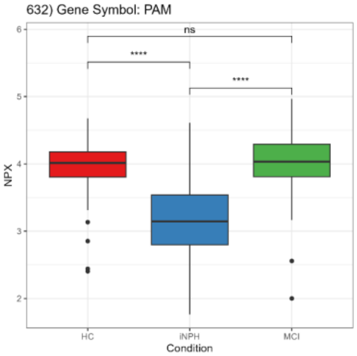


633) Gene Symbol: PAMR1

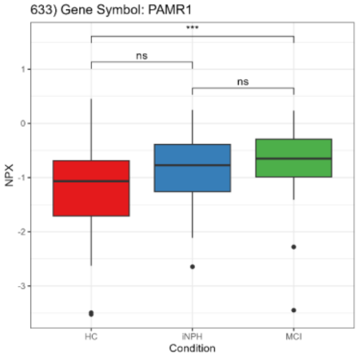


634) Gene Symbol: PCARE

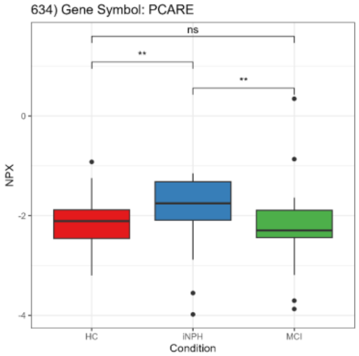


635) Gene Symbol: PCDH1

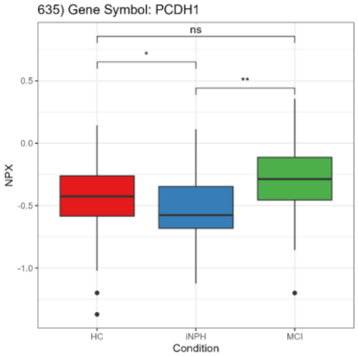


636) Gene Symbol: PCDH17

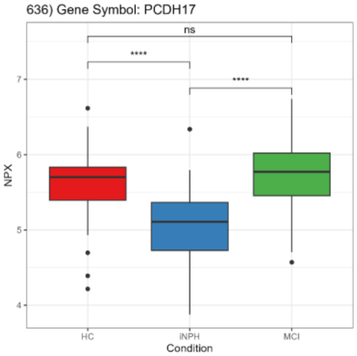


637) Gene Symbol: PCOLCE

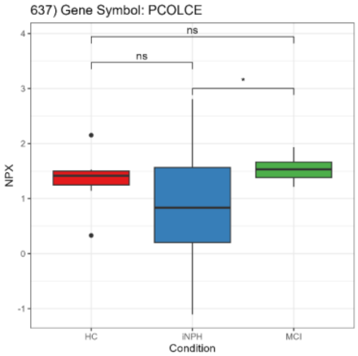


638) Gene Symbol: PCSK7

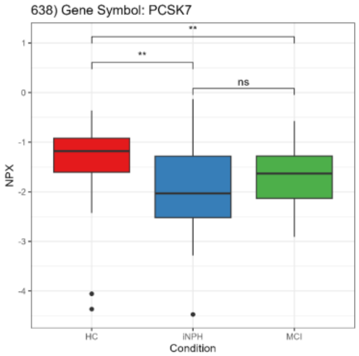


639) Gene Symbol: PDAP1

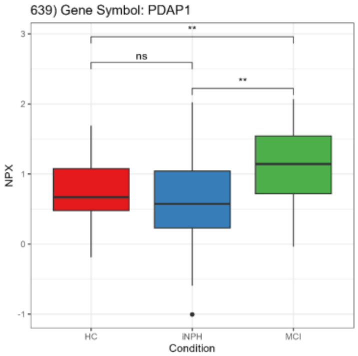


640) Gene Symbol: PDCD1

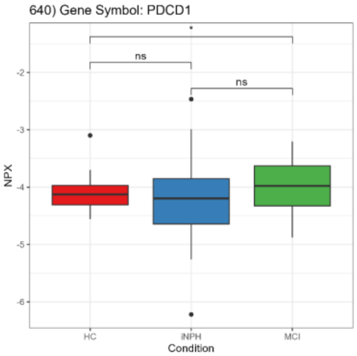

641) Gene Symbol: PDCD5

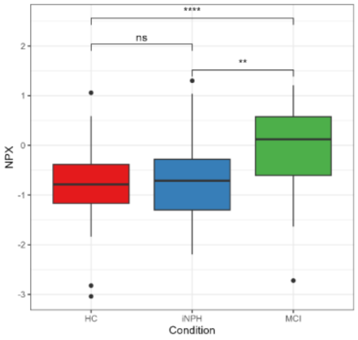


642) Gene Symbol: PDLIM5

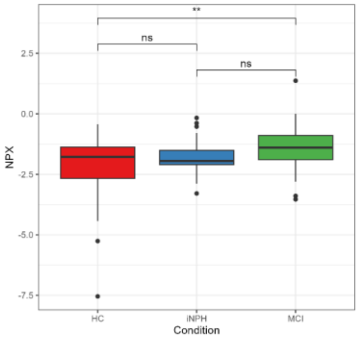


643) Gene Symbol: PEBP1

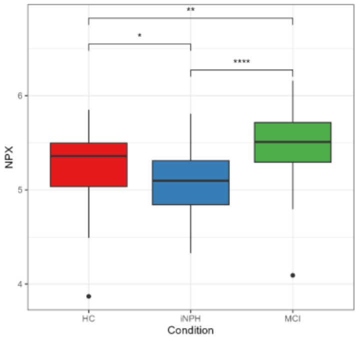


644) Gene Symbol: PECR

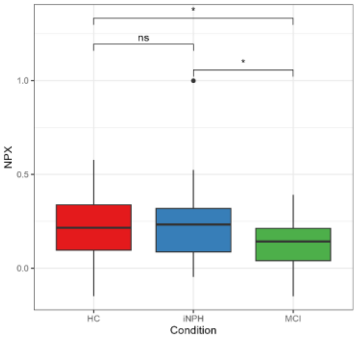


645) Gene Symbol: PER3

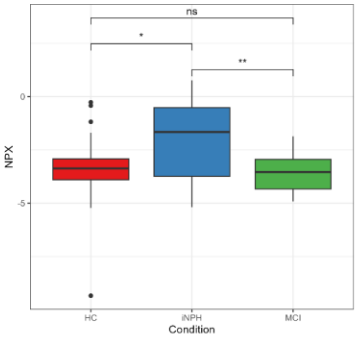


646) Gene Symbol: PFDN2

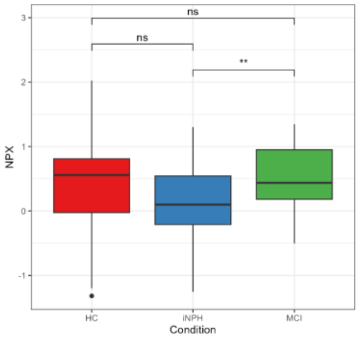


647) Gene Symbol: PFDN6

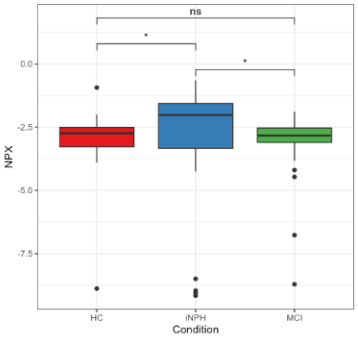


648) Gene Symbol: PGD

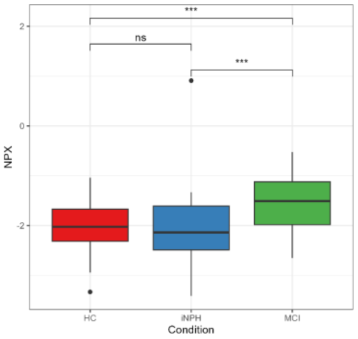


649) Gene Symbol: PGF

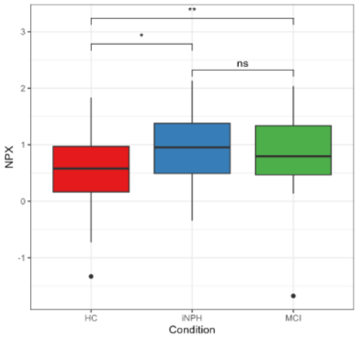


650) Gene Symbol: PHLDB1

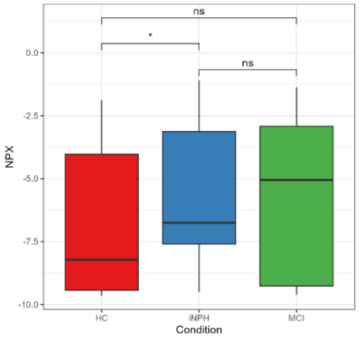


651) Gene Symbol: PHLDB2

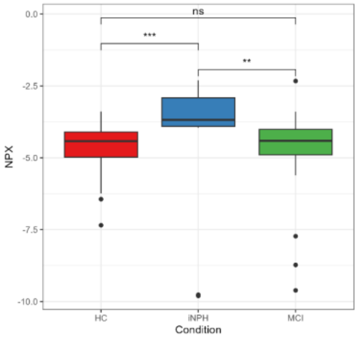


652) Gene Symbol: PIK3IP1

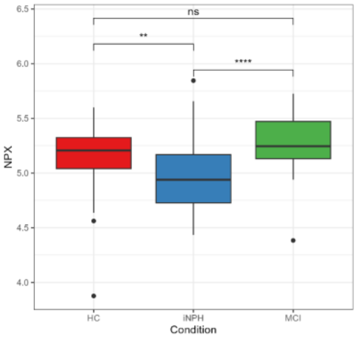


653) Gene Symbol: PILRA

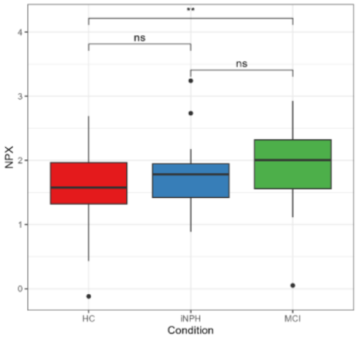


654) Gene Symbol: PILRB

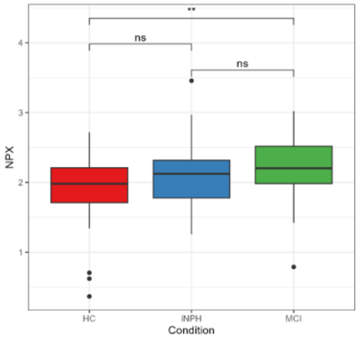


655) Gene Symbol: PINLYP

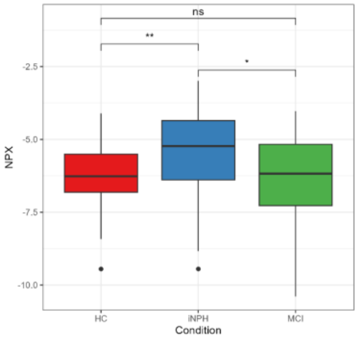


656) Gene Symbol: PITHD1

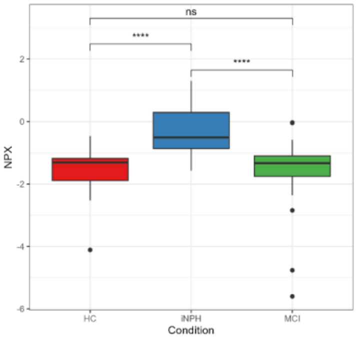


657) Gene Symbol: PLA2G10

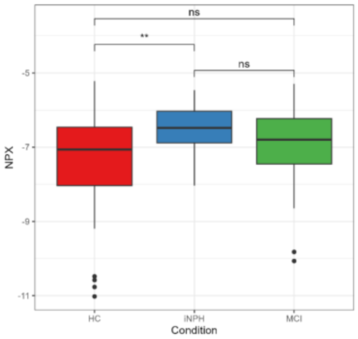


658) Gene Symbol: PLAT

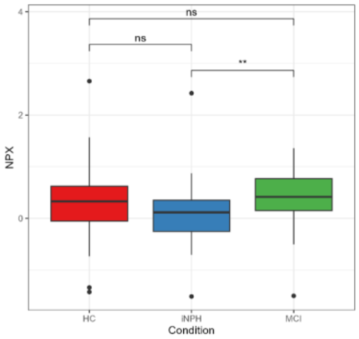


659) Gene Symbol: PLAU

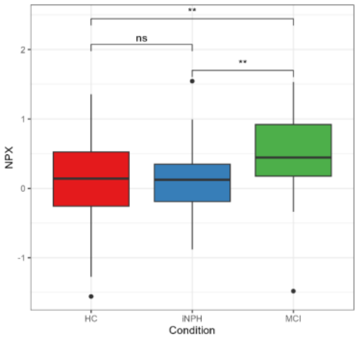


660) Gene Symbol: PLAUR

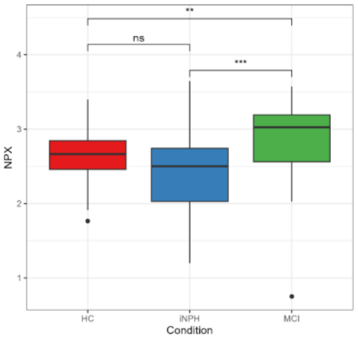

661) Gene Symbol: PLCB2

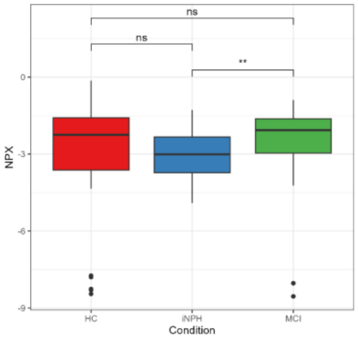


662) Gene Symbol: PLIN1

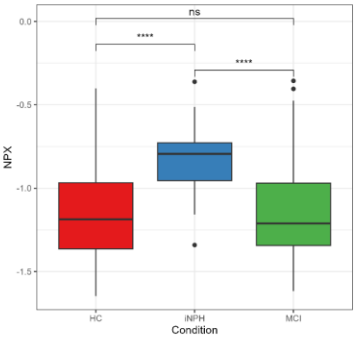


663) Gene Symbol: PLPBP

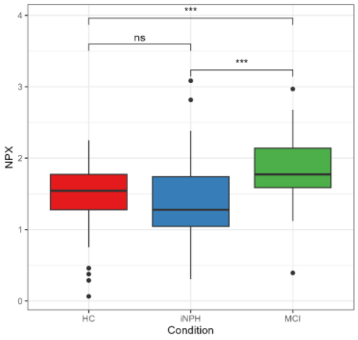


664) Gene Symbol: PLTP

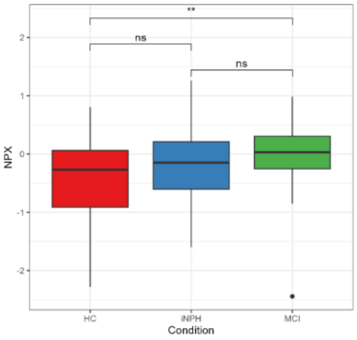


665) Gene Symbol: PLXDC2

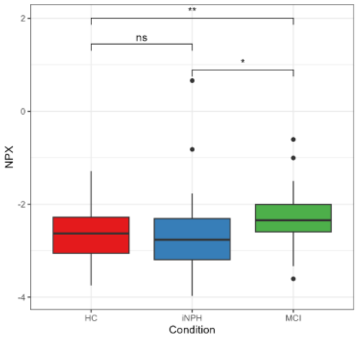


666) Gene Symbol: PLXNB2

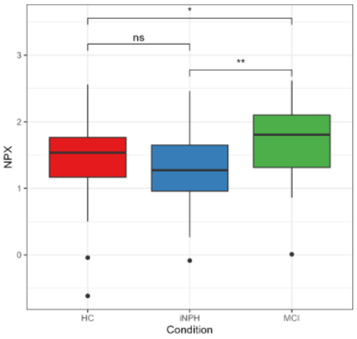


667) Gene Symbol: PLXNB3

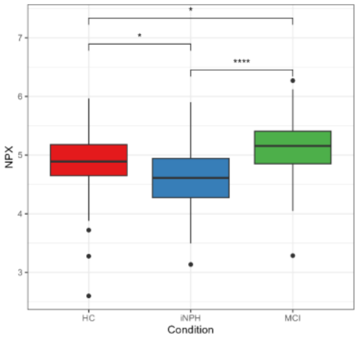


668) Gene Symbol: PMM2

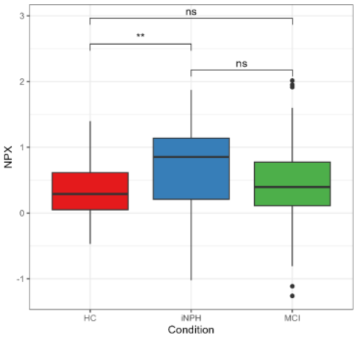


669) Gene Symbol: PMS1

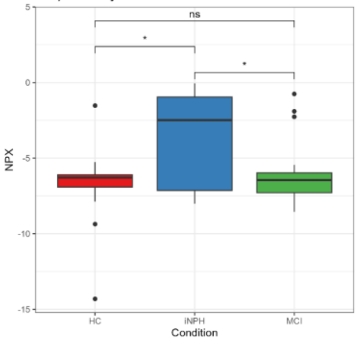


670) Gene Symbol: PODXL2

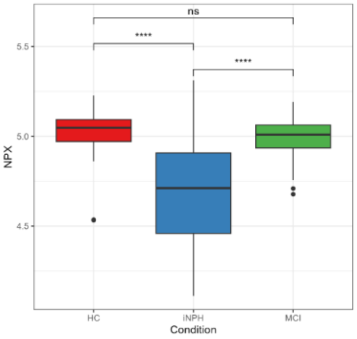


671) Gene Symbol: PPBP

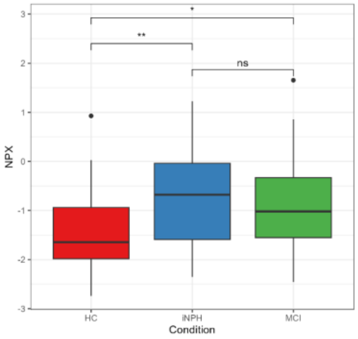


672) Gene Symbol: PPCDC

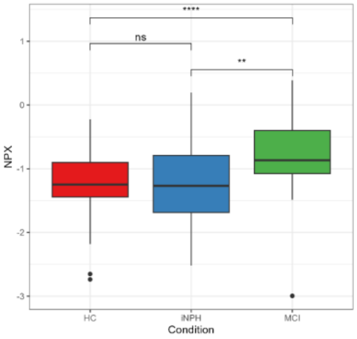


673) Gene Symbol: PPM1B

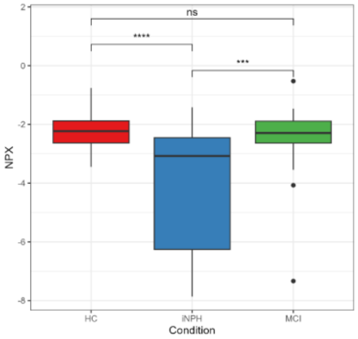


674) Gene Symbol: PPP1R14A

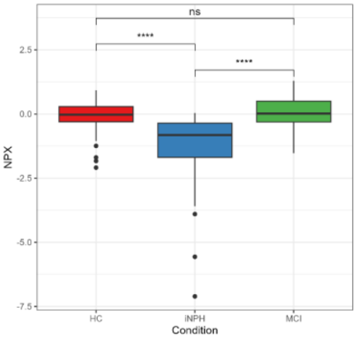


675) Gene Symbol: PPP1R9B

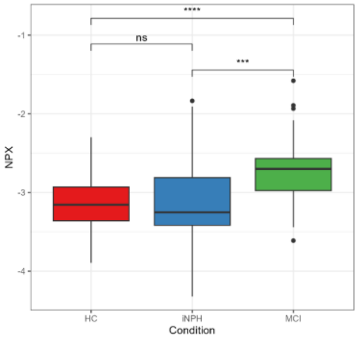


676) Gene Symbol: PPP2R5A

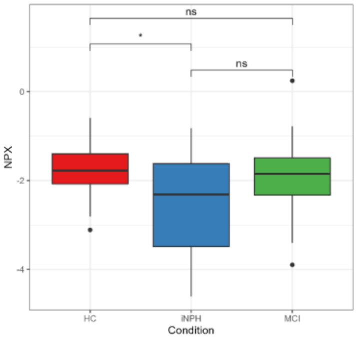


677) Gene Symbol: PPP3R1

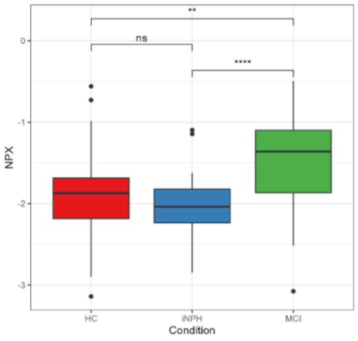


678) Gene Symbol: PPY

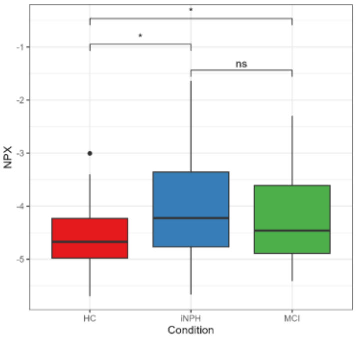


679) Gene Symbol: PRAP1

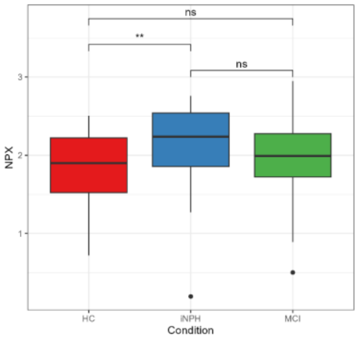


680) Gene Symbol: PRC1

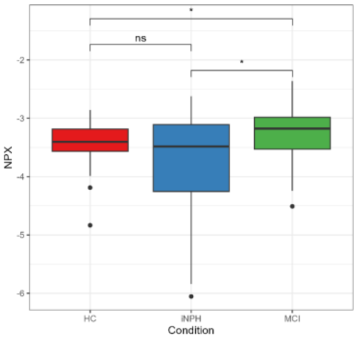

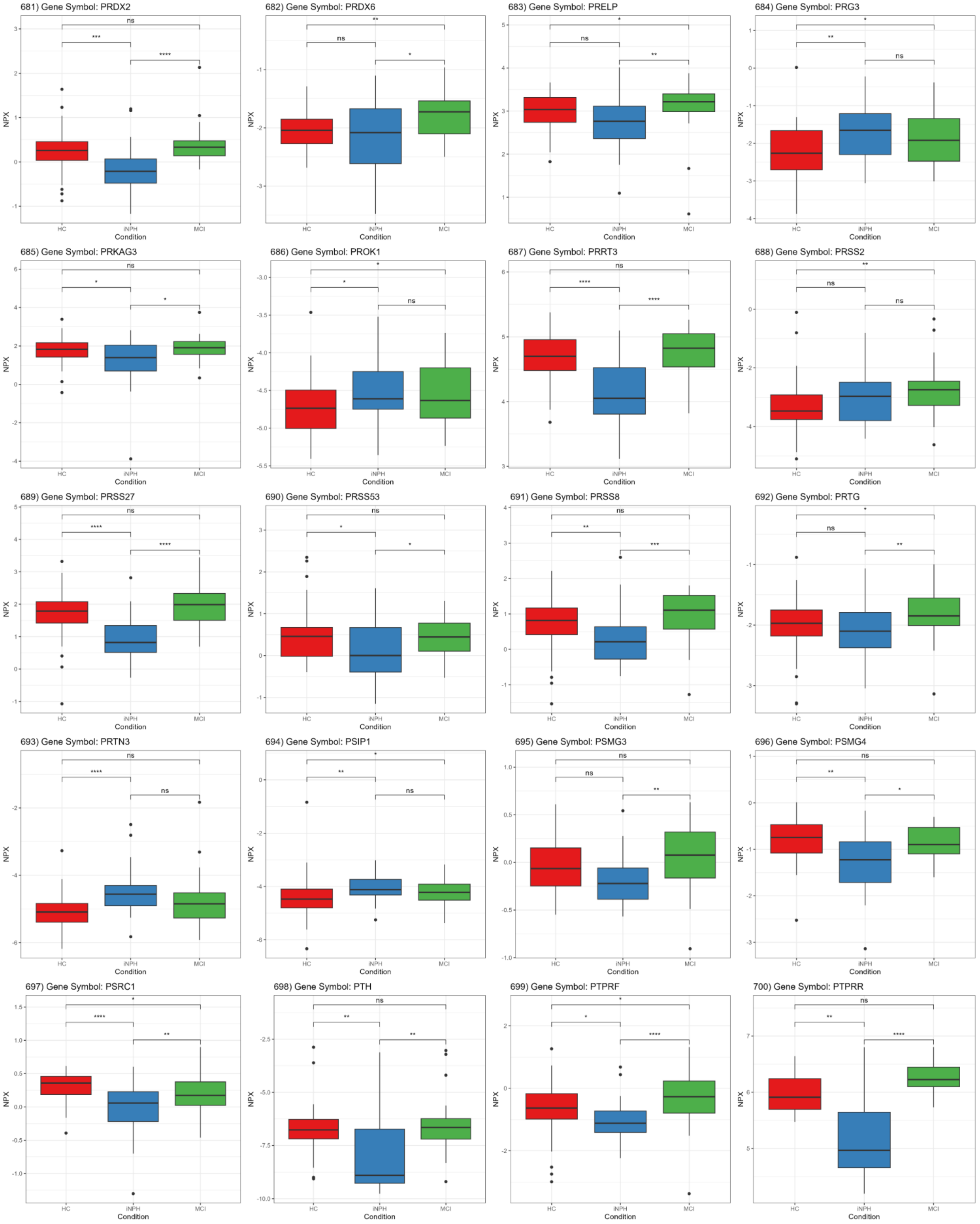
681) Gene Symbol: PRDX2
ns
***
****
682) Gene Symbol: PRDX6
**
ns
*
683) Gene Symbol: PRELP
*
ns
**
684) Gene Symbol: PRG3
*
**
ns
685) Gene Symbol: PRKAG3
ns
*
*
686) Gene Symbol: PROK1
*
*
ns
687) Gene Symbol: PRRT3
ns
****
****
688) Gene Symbol: PRSS2
**
ns
ns
689) Gene Symbol: PRSS27
ns
****
****
690) Gene Symbol: PRSS53
ns
*
*
691) Gene Symbol: PRSS8
ns
**
***
692) Gene Symbol: PRTG
*
ns
**
693) Gene Symbol: PRTN3
ns
****
ns
694) Gene Symbol: PSIP1
*
**
ns
695) Gene Symbol: PSMG3
ns
ns
**
696) Gene Symbol: PSMG4
ns
**
*
697) Gene Symbol: PSRC1
*
****
**
698) Gene Symbol: PTH
ns
**
**
699) Gene Symbol: PTPRF
*
*
****
700) Gene Symbol: PTPRR
ns
**
****
NPX
HC
iNPH
MCI
Condition

701) Gene Symbol: PTPRS

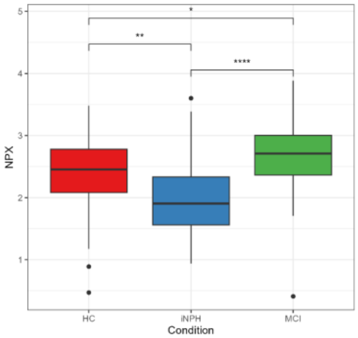


702) Gene Symbol: PTPRZ1

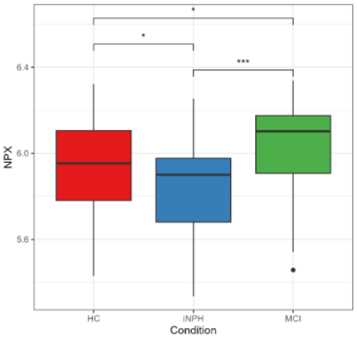


703) Gene Symbol: PVALB

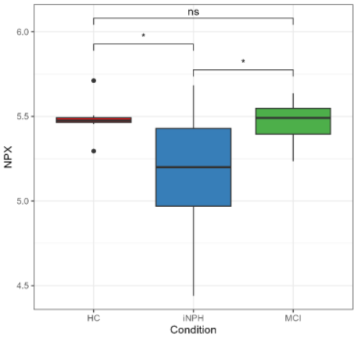


704) Gene Symbol: PVR

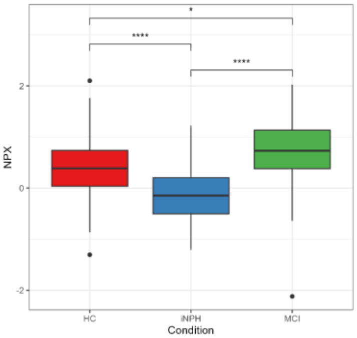


705) Gene Symbol: PXDNL

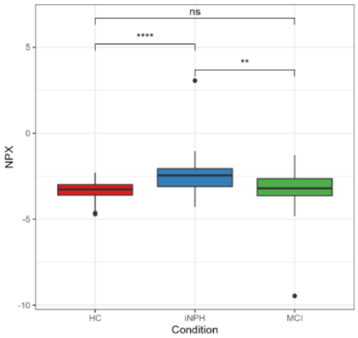


706) Gene Symbol: PYDC1

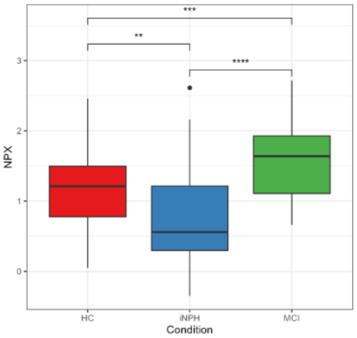


707) Gene Symbol: QDPR

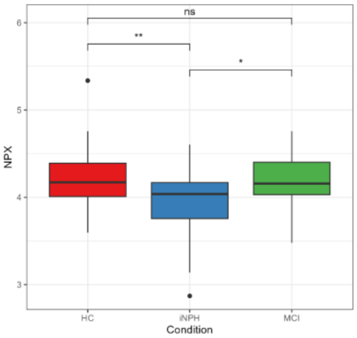


708) Gene Symbol: QPCT

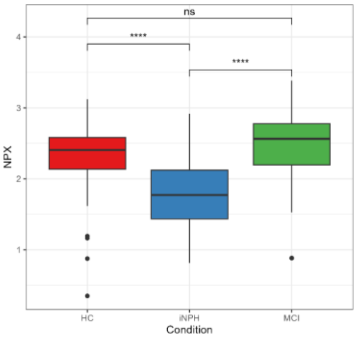


709) Gene Symbol: RAB33A

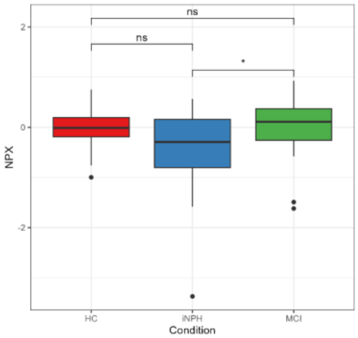


710) Gene Symbol: RAB39B

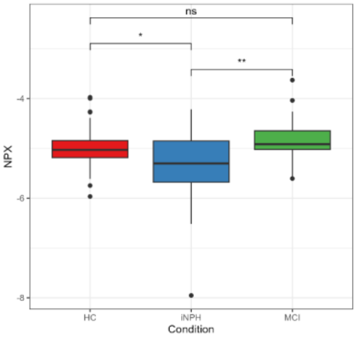


711) Gene Symbol: RAB3GAP1

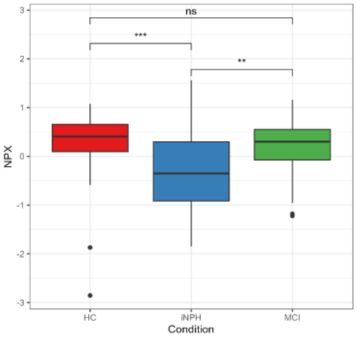


712) Gene Symbol: RAC3

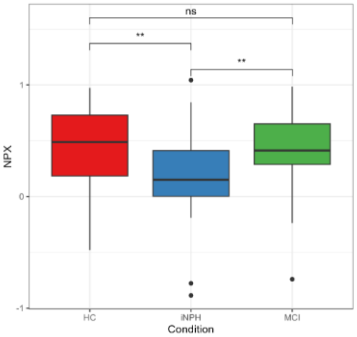


713) Gene Symbol: RAD23B

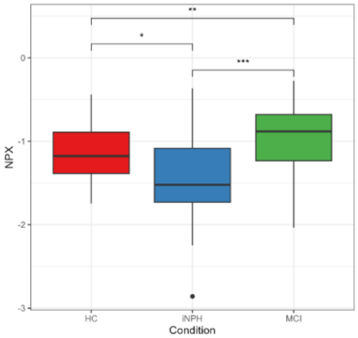


714) Gene Symbol: RAD51

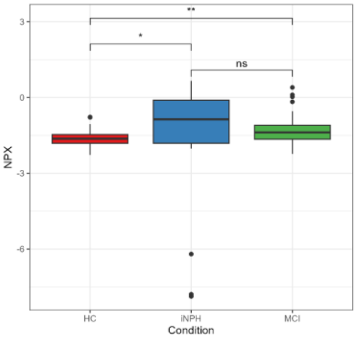


715) Gene Symbol: RALY

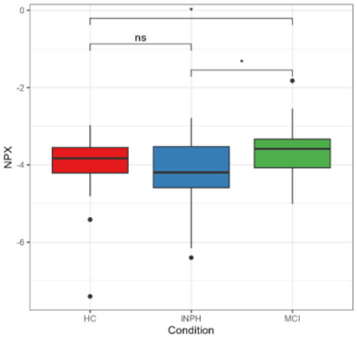


716) Gene Symbol: RARRES2

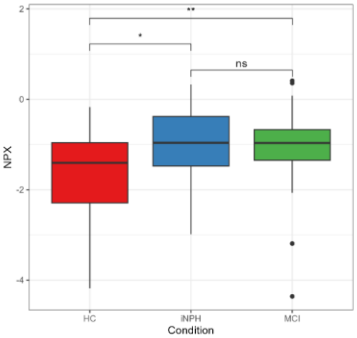


717) Gene Symbol: RBKS

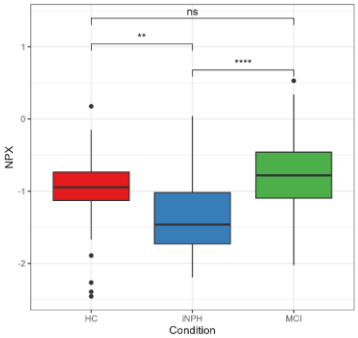


718) Gene Symbol: RBM19

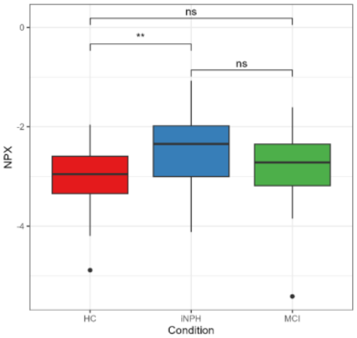


719) Gene Symbol: RBM25

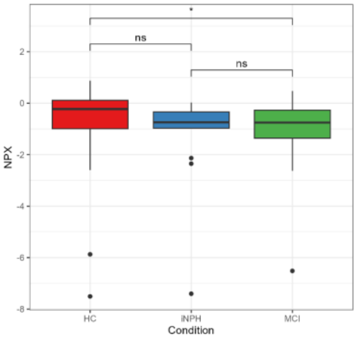


720) Gene Symbol: RBP2

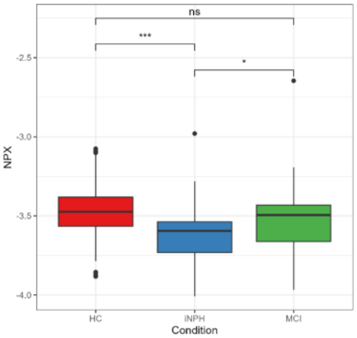

721) Gene Symbol: REG1B

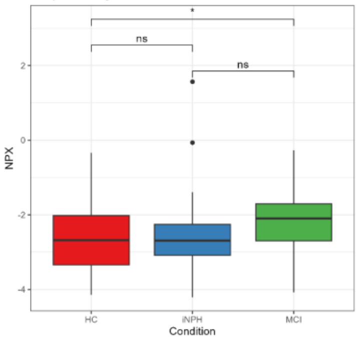


722) Gene Symbol: REG3A

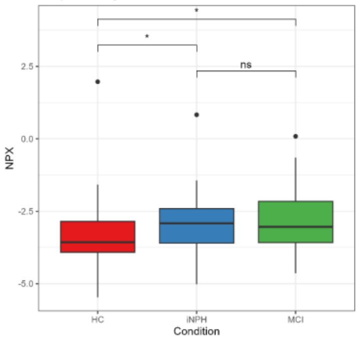


723) Gene Symbol: REG3G

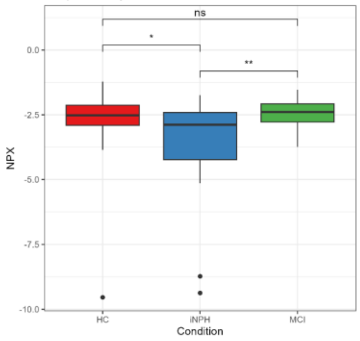


724) Gene Symbol: RELT

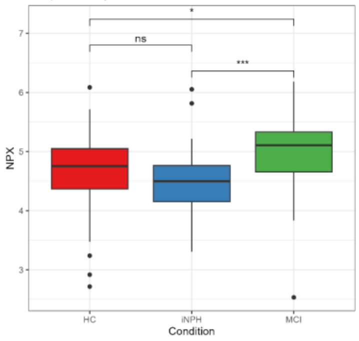


725) Gene Symbol: REXO2

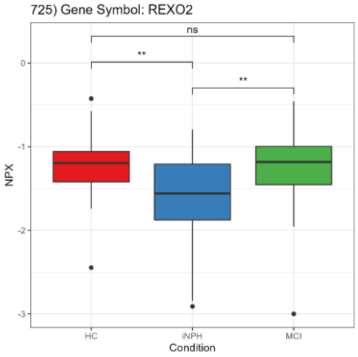


726) Gene Symbol: RGMA

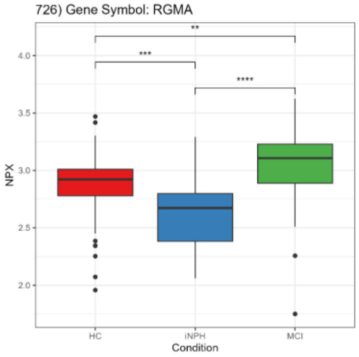


727) Gene Symbol: RGMB

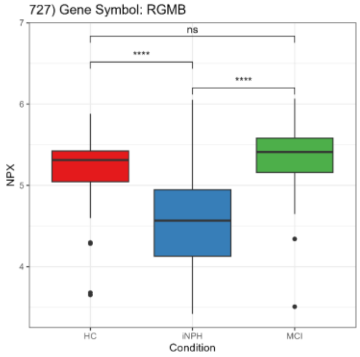


728) Gene Symbol: RHOC

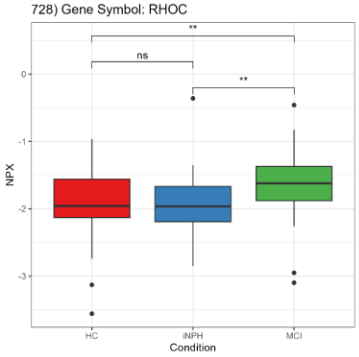


729) Gene Symbol: RILPL2

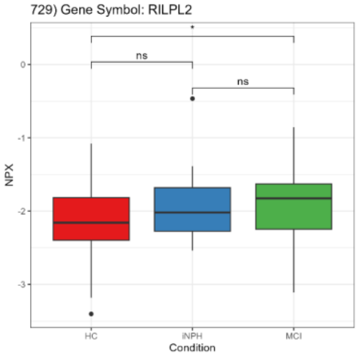


730) Gene Symbol: RNASE3

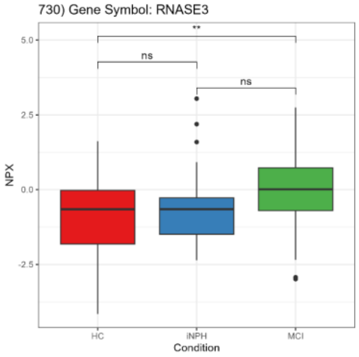


731) Gene Symbol: RNASE4

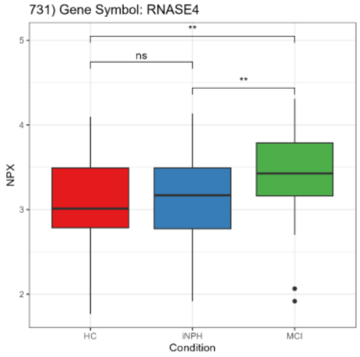


732) Gene Symbol: RNASET2

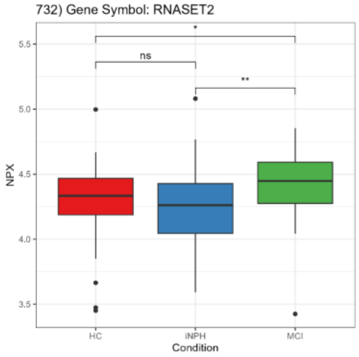


733) Gene Symbol: RNF43

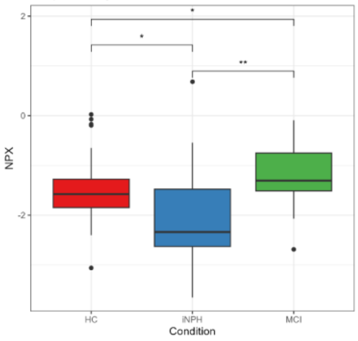


734) Gene Symbol: ROBO1

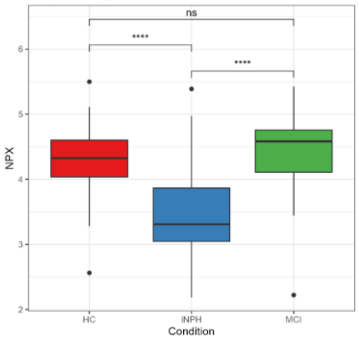


735) Gene Symbol: ROBO2

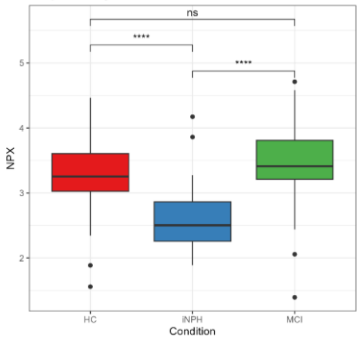


736) Gene Symbol: ROR1

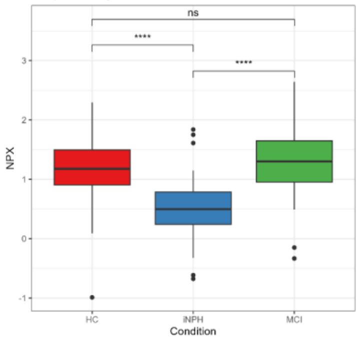


737) Gene Symbol: RPGR

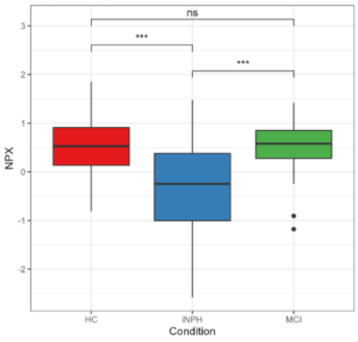


738) Gene Symbol: RSPO1

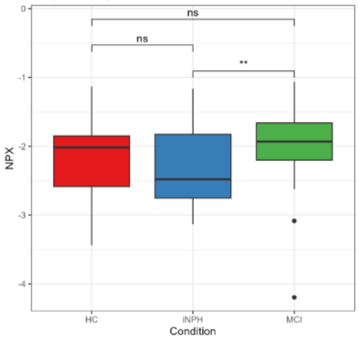


739) Gene Symbol: RTBDN

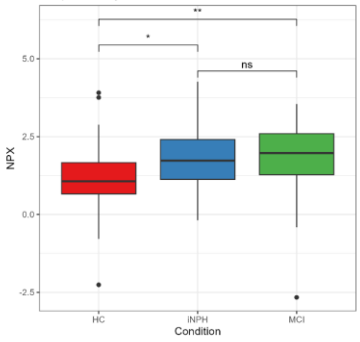


740) Gene Symbol: S100A12

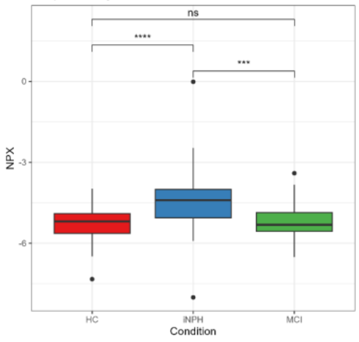

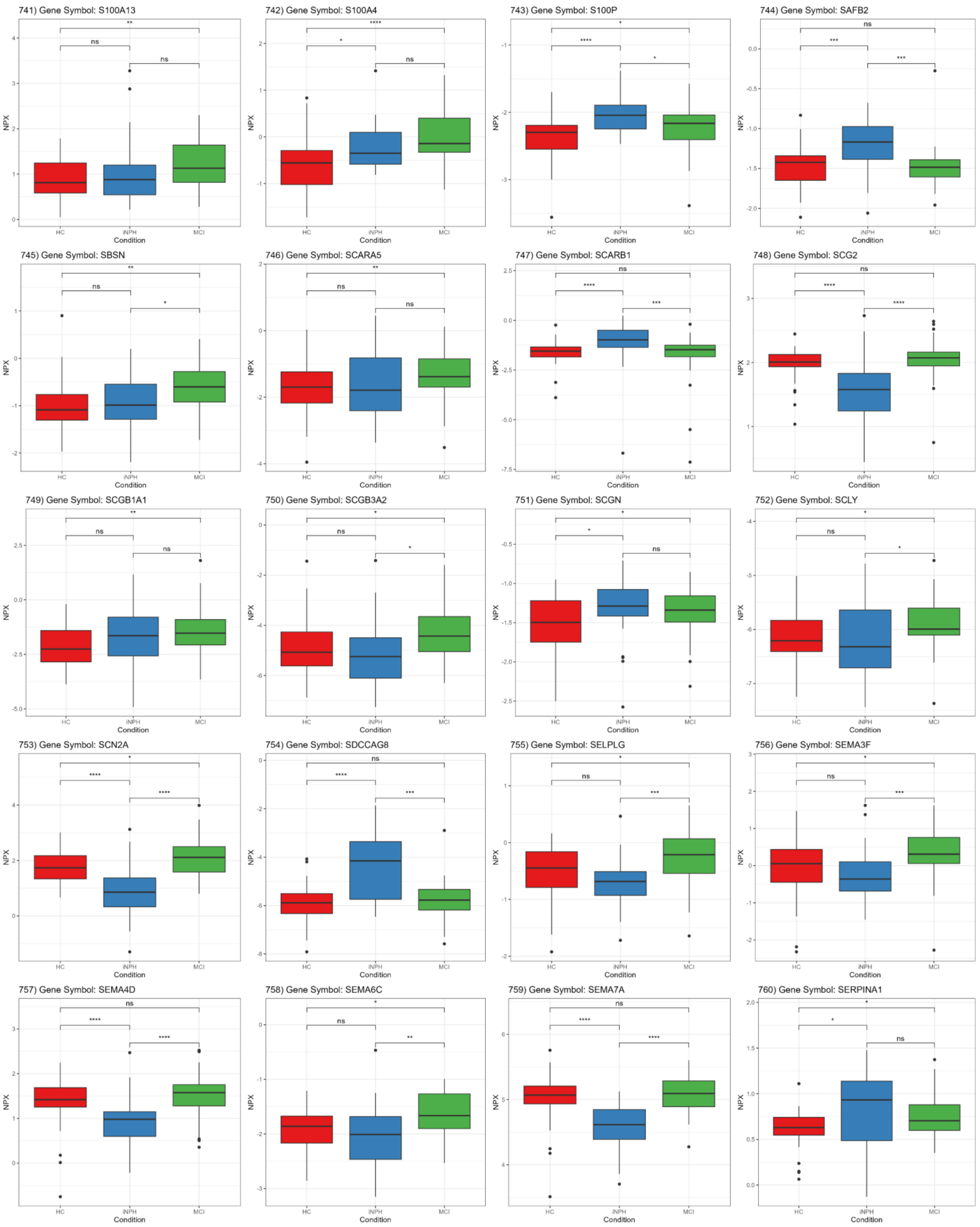
741) Gene Symbol: S100A13
742) Gene Symbol: S100A4
743) Gene Symbol: S100P
744) Gene Symbol: SAFB2
745) Gene Symbol: SBSN
746) Gene Symbol: SCARA5
747) Gene Symbol: SCARB1
748) Gene Symbol: SCG2
749) Gene Symbol: SCGB1A1
750) Gene Symbol: SCGB3A2
751) Gene Symbol: SCGN
752) Gene Symbol: SCLY
753) Gene Symbol: SCN2A
754) Gene Symbol: SDCCAG8
755) Gene Symbol: SELPLG
756) Gene Symbol: SEMA3F
757) Gene Symbol: SEMA4D
758) Gene Symbol: SEMA6C
759) Gene Symbol: SEMA7A
760) Gene Symbol: SERPINA1
NPX
Condition
HC
iNPH
MCI

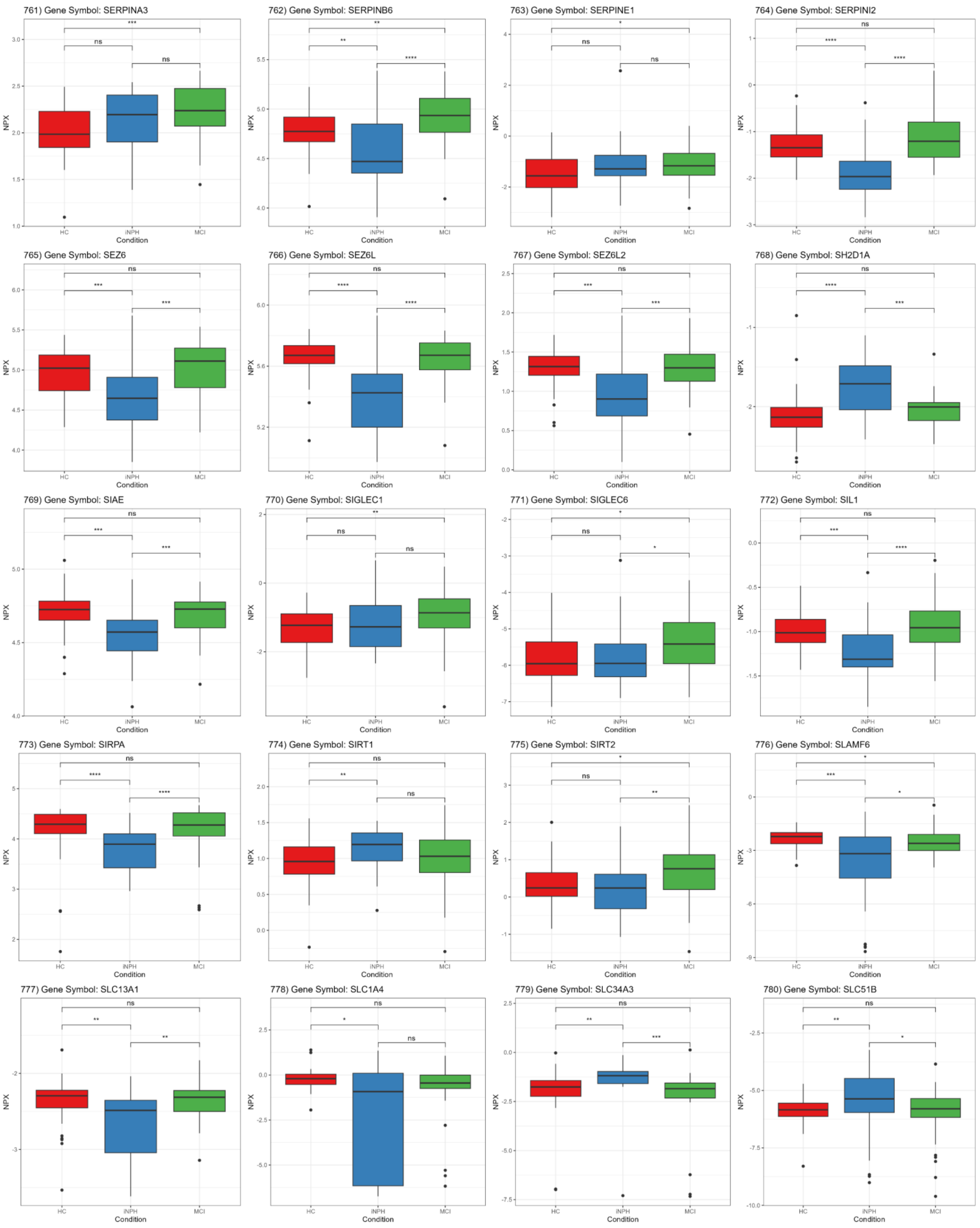
761) Gene Symbol: SERPINA3
762) Gene Symbol: SERPINB6
763) Gene Symbol: SERPINE1
764) Gene Symbol: SERPINI2
765) Gene Symbol: SEZ6
766) Gene Symbol: SEZ6L
767) Gene Symbol: SEZ6L2
768) Gene Symbol: SH2D1A
769) Gene Symbol: SIAE
770) Gene Symbol: SIGLEC1
771) Gene Symbol: SIGLEC6
772) Gene Symbol: SIL1
773) Gene Symbol: SIRPA
774) Gene Symbol: SIRT1
775) Gene Symbol: SIRT2
776) Gene Symbol: SLAMF6
777) Gene Symbol: SLC13A1
778) Gene Symbol: SLC1A4
779) Gene Symbol: SLC34A3
780) Gene Symbol: SLC51B
NPX
Condition
HC
iNPH
MCI

781) Gene Symbol: SLIRP

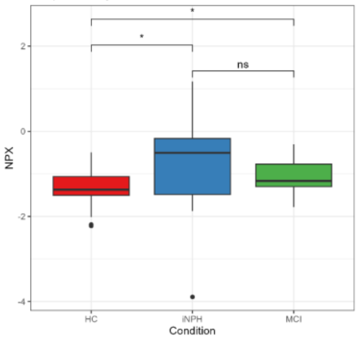


782) Gene Symbol: SLIT2

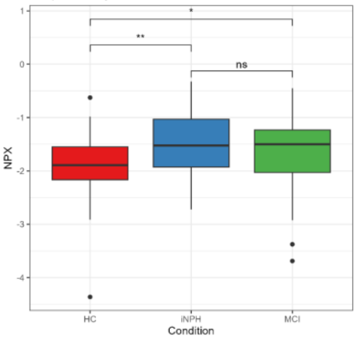


783) Gene Symbol: SLITRK1

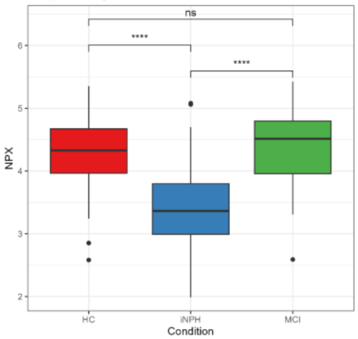


784) Gene Symbol: SLITRK2

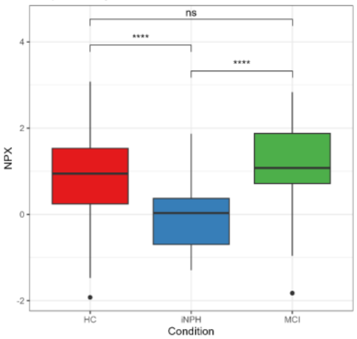


785) Gene Symbol: SMOC1

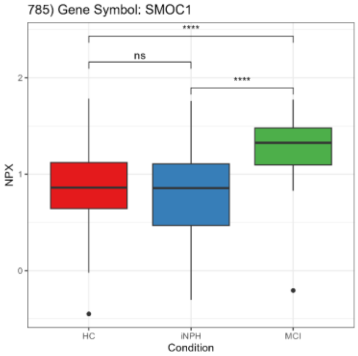


786) Gene Symbol: SMOC2

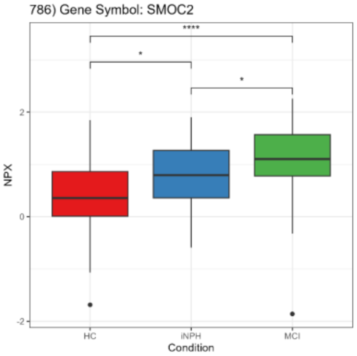


787) Gene Symbol: SMPD3

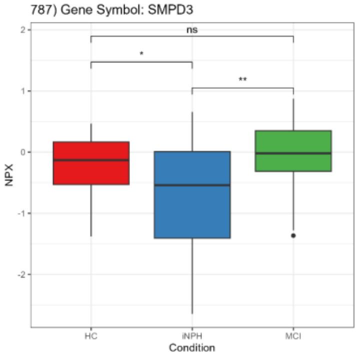


788) Gene Symbol: SMPDL3B

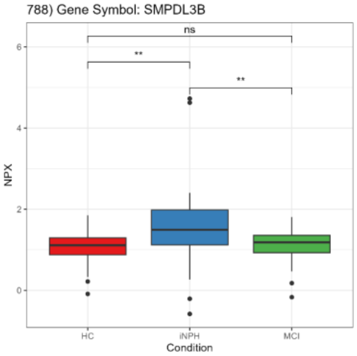


789) Gene Symbol: SMS

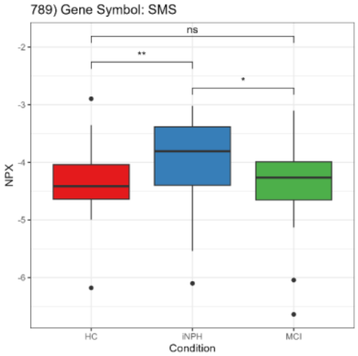


790) Gene Symbol: SNAP25

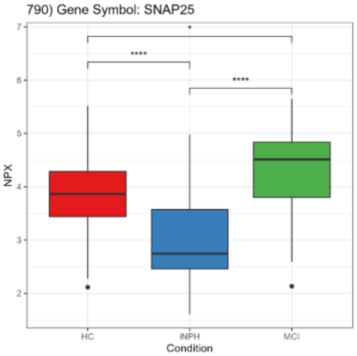


791) Gene Symbol: SNAP29

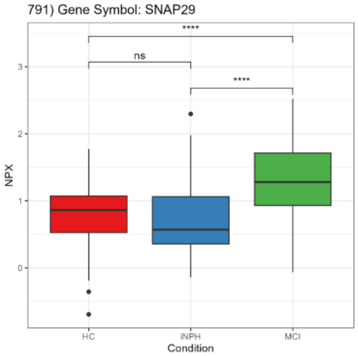


792) Gene Symbol: SNAPIN

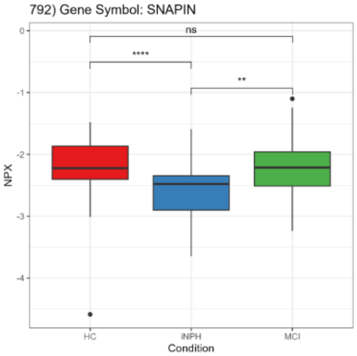


793) Gene Symbol: SNCA

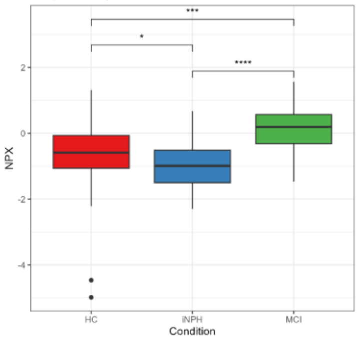


794) Gene Symbol: SNCG

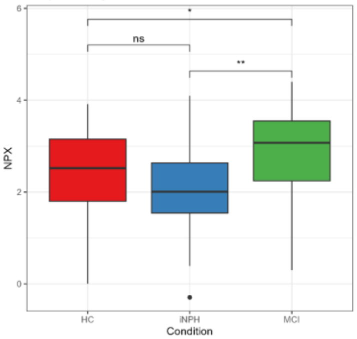


795) Gene Symbol: SNED1

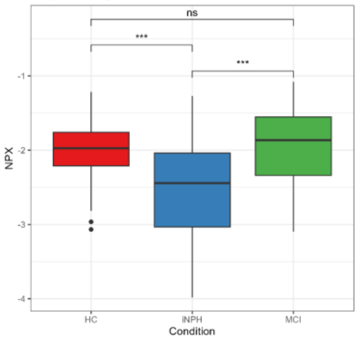


796) Gene Symbol: SNX15

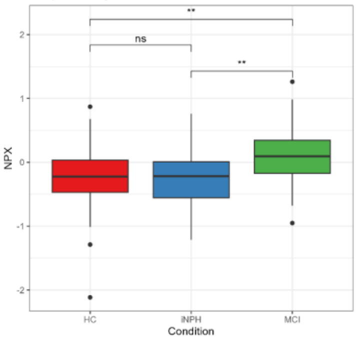


797) Gene Symbol: SNX18

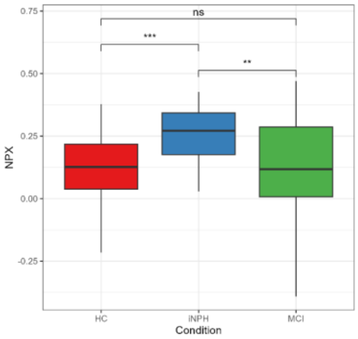


798) Gene Symbol: SOD1

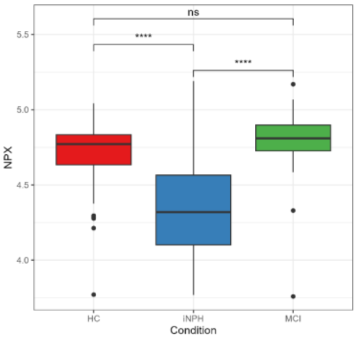


799) Gene Symbol: SOD2

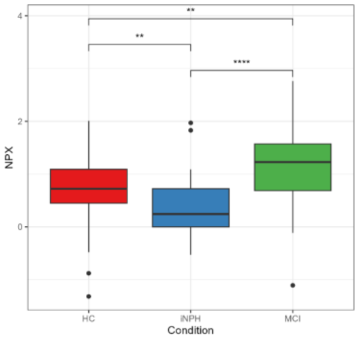


800) Gene Symbol: SOD3

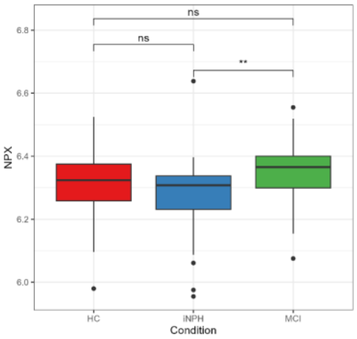

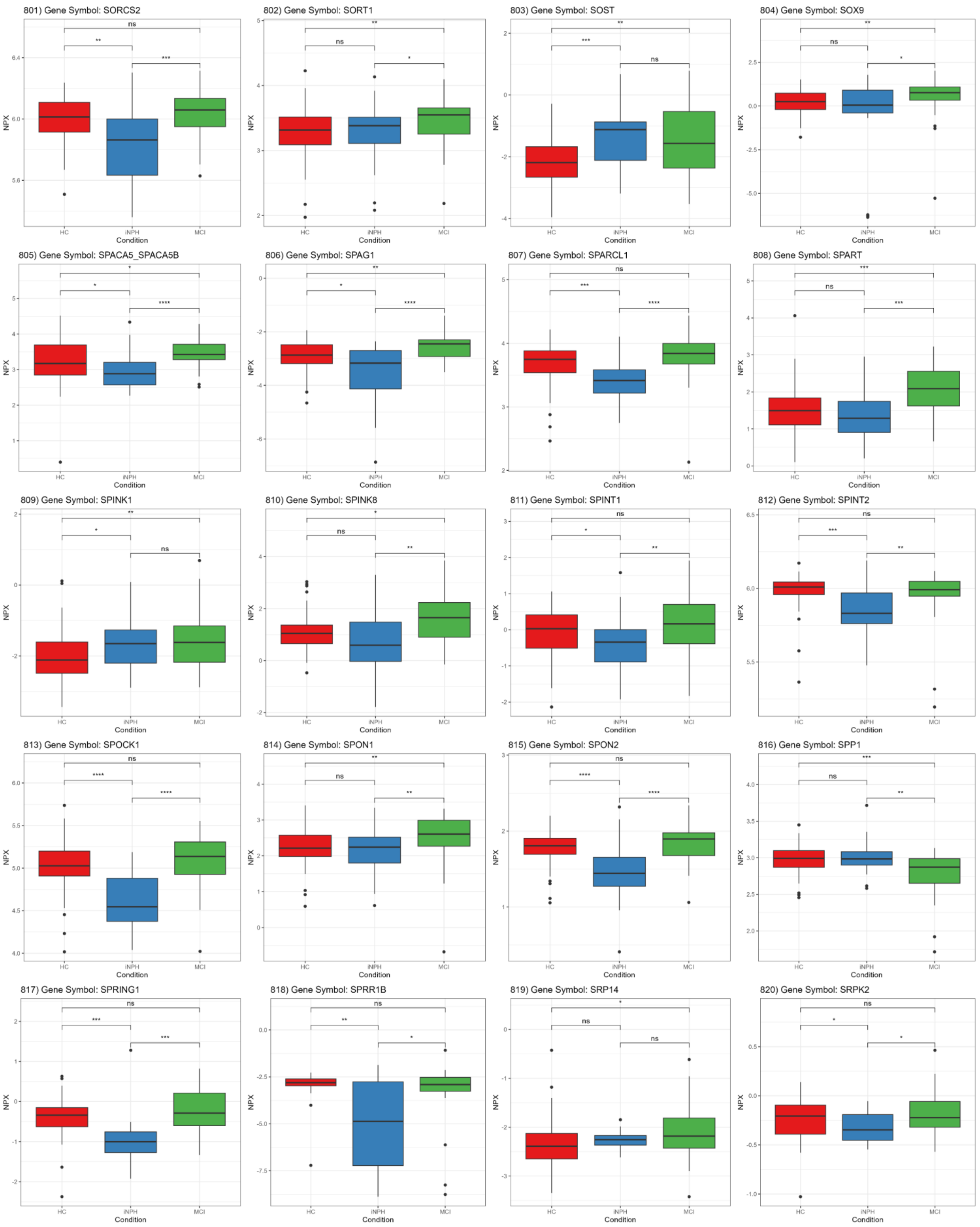
801) Gene Symbol: SORCS2
802) Gene Symbol: SORT1
803) Gene Symbol: SOST
804) Gene Symbol: SOX9
805) Gene Symbol: SPACA5_SPACA5B
806) Gene Symbol: SPAG1
807) Gene Symbol: SPARCL1
808) Gene Symbol: SPART
809) Gene Symbol: SPINK1
810) Gene Symbol: SPINK8
811) Gene Symbol: SPINT1
812) Gene Symbol: SPINT2
813) Gene Symbol: SPOCK1
814) Gene Symbol: SPON1
815) Gene Symbol: SPON2
816) Gene Symbol: SPP1
817) Gene Symbol: SPRING1
818) Gene Symbol: SPRR1B
819) Gene Symbol: SRP14
820) Gene Symbol: SRPK2
NPX
Condition
HC
iNPH
MCI

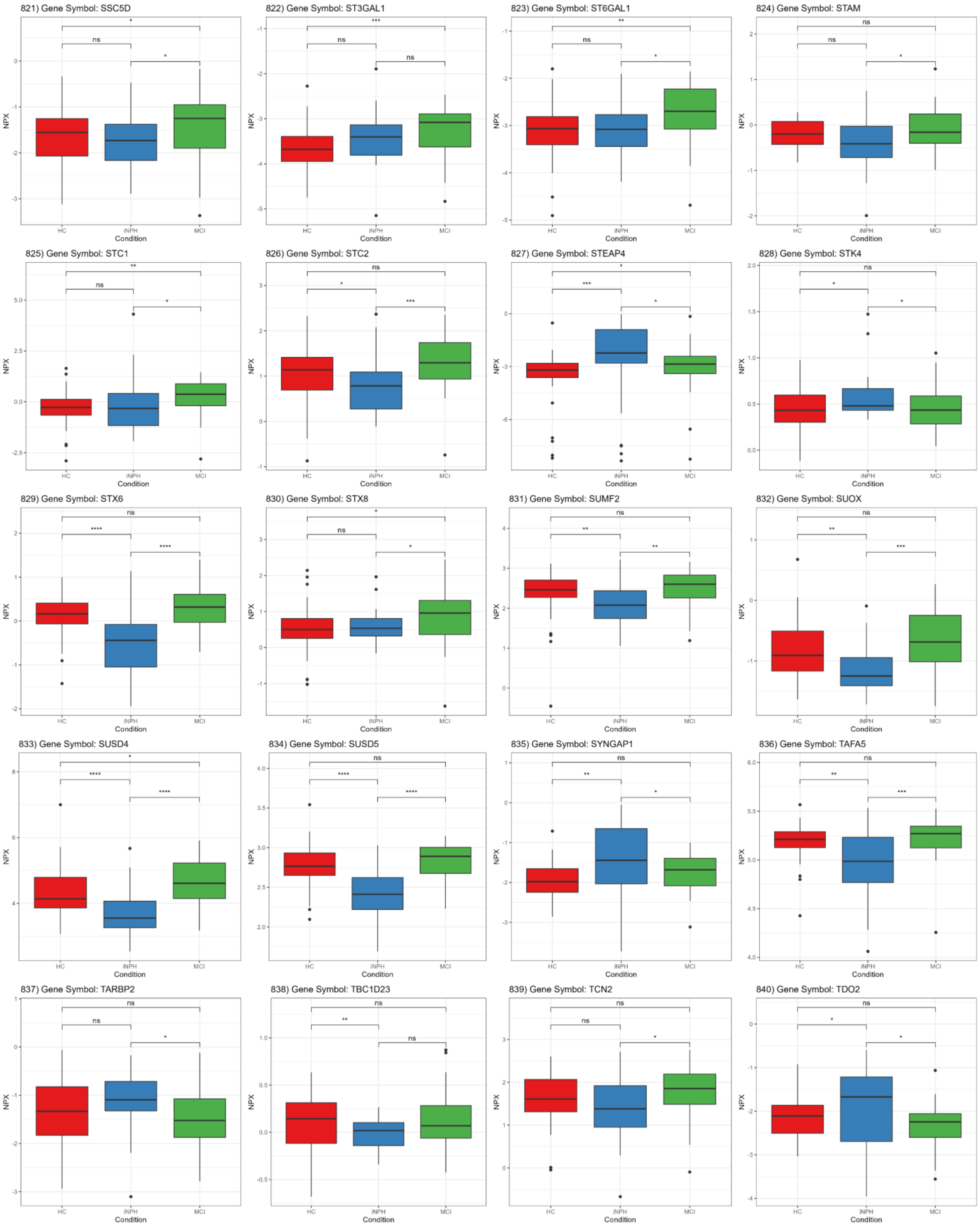
821) Gene Symbol: SSC5D
822) Gene Symbol: ST3GAL1
823) Gene Symbol: ST6GAL1
824) Gene Symbol: STAM
825) Gene Symbol: STC1
826) Gene Symbol: STC2
827) Gene Symbol: STEAP4
828) Gene Symbol: STK4
829) Gene Symbol: STX6
830) Gene Symbol: STX8
831) Gene Symbol: SUMF2
832) Gene Symbol: SUOX
833) Gene Symbol: SUSD4
834) Gene Symbol: SUSD5
835) Gene Symbol: SYNGAP1
836) Gene Symbol: TAFA5
837) Gene Symbol: TARBP2
838) Gene Symbol: TBC1D23
839) Gene Symbol: TCN2
840) Gene Symbol: TDO2
NPX
Condition
HC
iNPH
MCI

841) Gene Symbol: TEX101

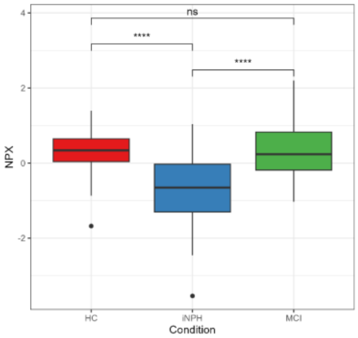


842) Gene Symbol: TFF1

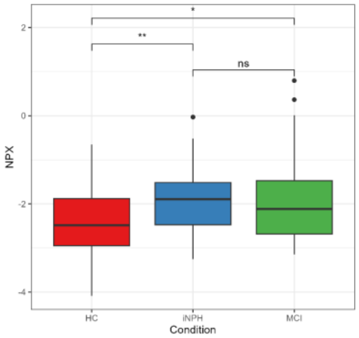


843) Gene Symbol: TFPI

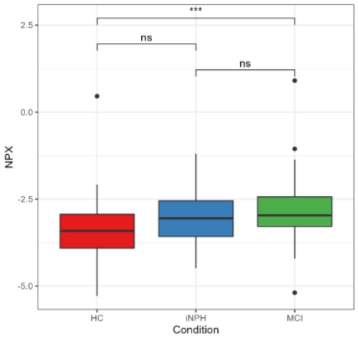


844) Gene Symbol: TFPI2

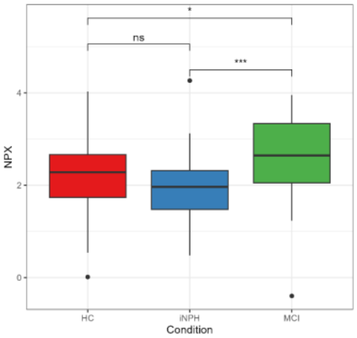


845) Gene Symbol: TGFBI

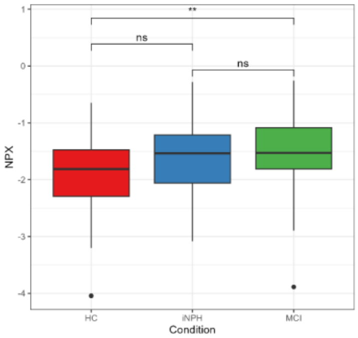


846) Gene Symbol: TGFBR1

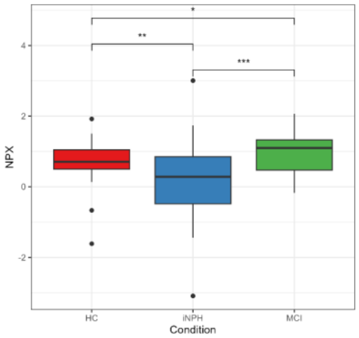


847) Gene Symbol: TGFBR2

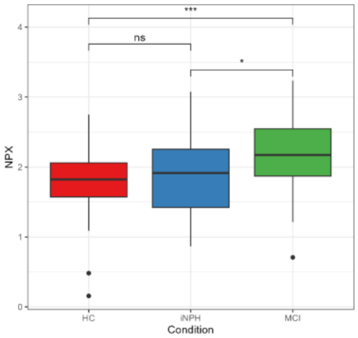


848) Gene Symbol: TGFBR3

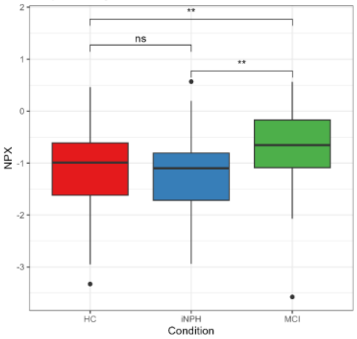


849) Gene Symbol: TGOLN2

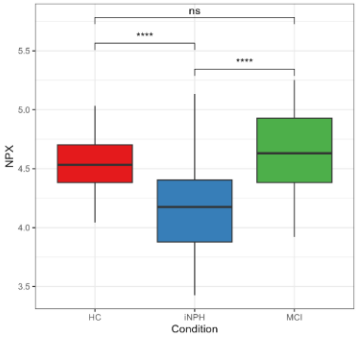


850) Gene Symbol: THAP12

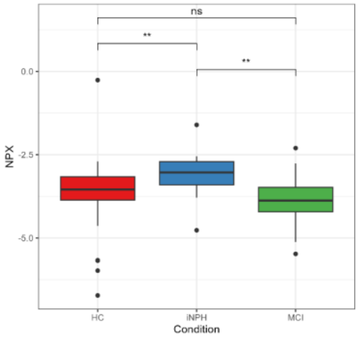


851) Gene Symbol: THBS4

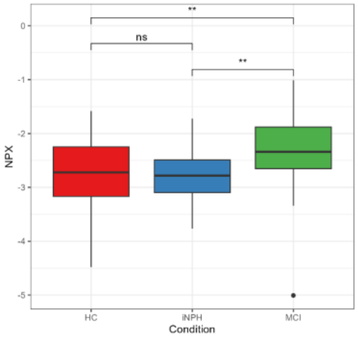


852) Gene Symbol: THPO

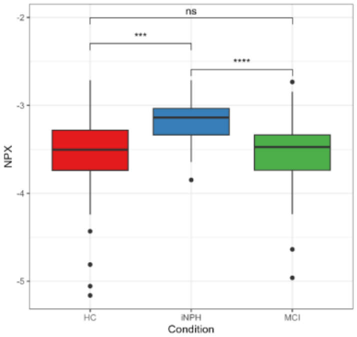


853) Gene Symbol: THSD1

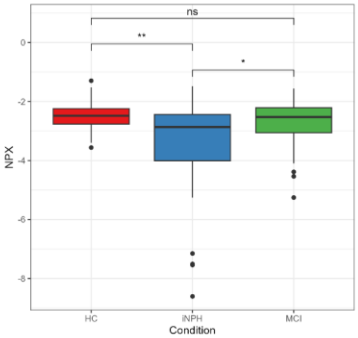


854) Gene Symbol: THTPA

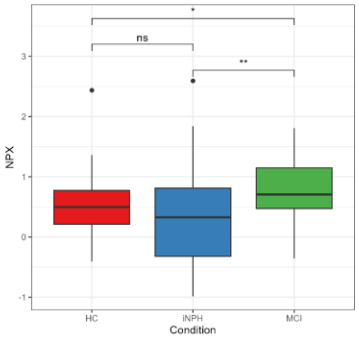


855) Gene Symbol: TIE1

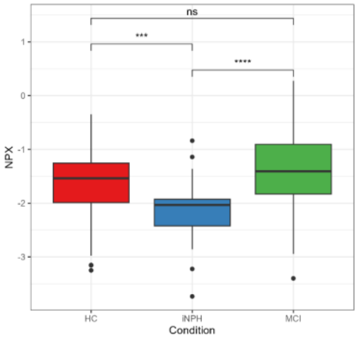


856) Gene Symbol: TIGAR

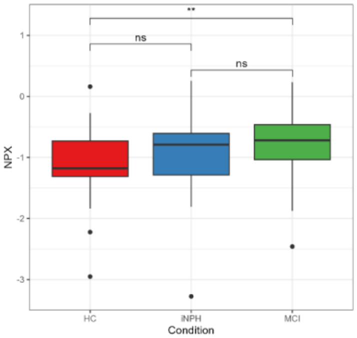


857) Gene Symbol: TIMM10

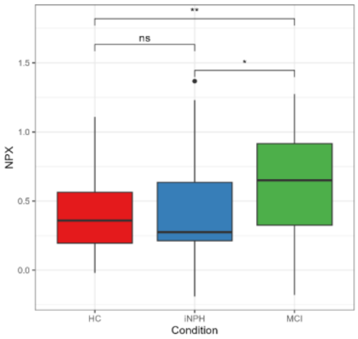


858) Gene Symbol: TIMP1

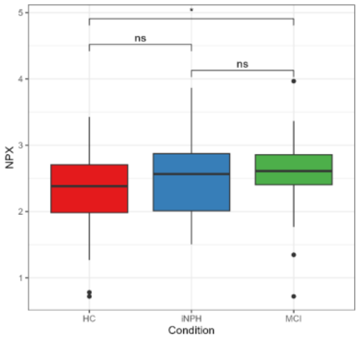


859) Gene Symbol: TIMP3

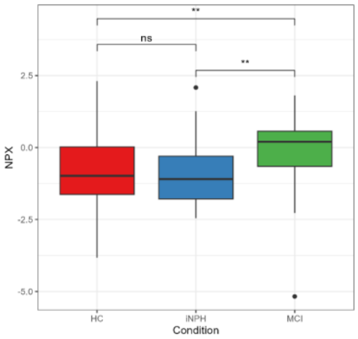


860) Gene Symbol: TIMP4

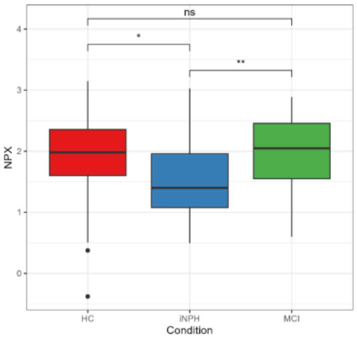

861) Gene Symbol: TINAGL1

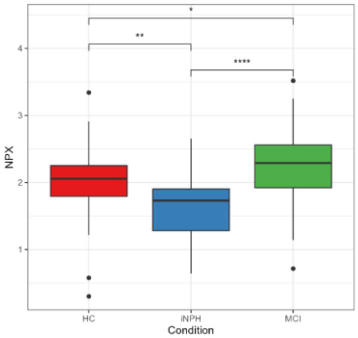


862) Gene Symbol: TK1

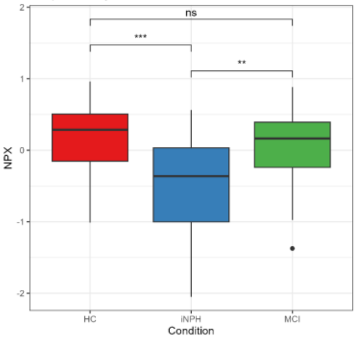


863) Gene Symbol: TMED4

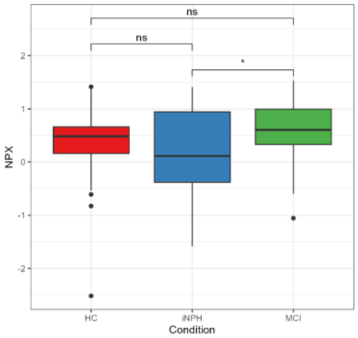


864) Gene Symbol: TMEM132A

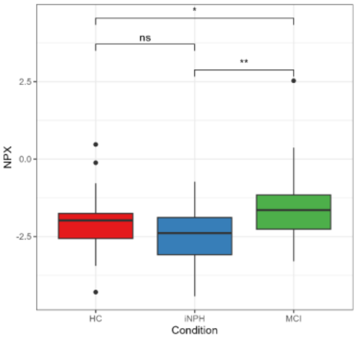


865) Gene Symbol: TMSB10

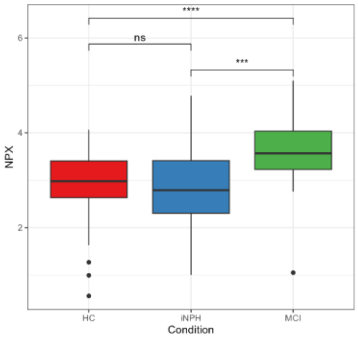


866) Gene Symbol: TNC

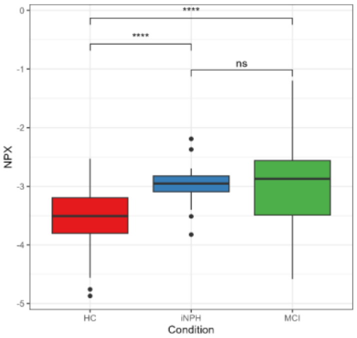


867) Gene Symbol: TNFAIP8

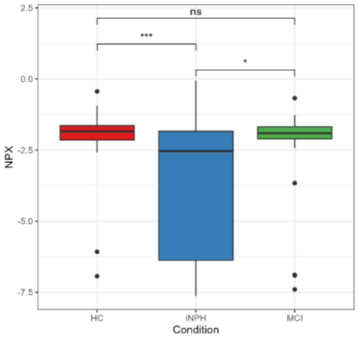


868) Gene Symbol: TNFRSF10B

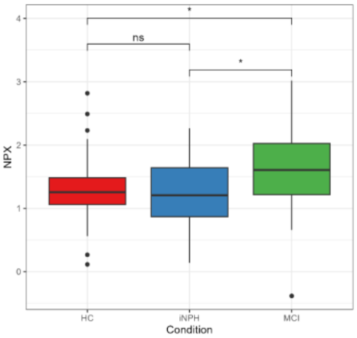


869) Gene Symbol: TNFRSF12A

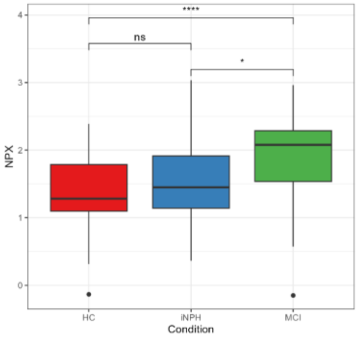


870) Gene Symbol: TNFRSF13B

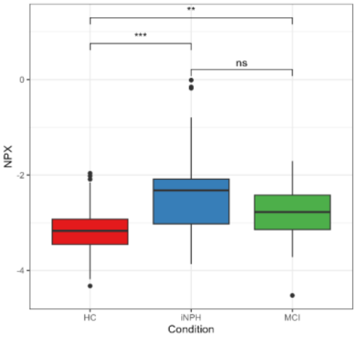


871) Gene Symbol: TNFRSF13C

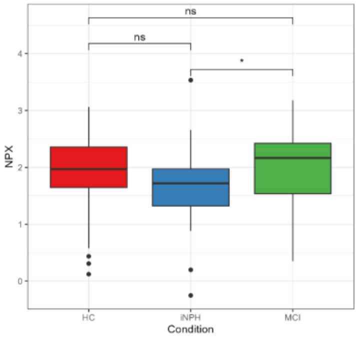


872) Gene Symbol: TNFRSF17

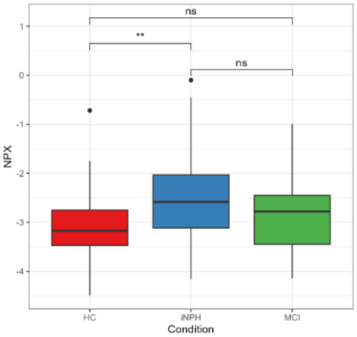


873) Gene Symbol: TNFRSF19

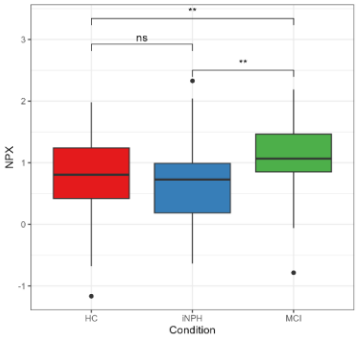


874) Gene Symbol: TNFRSF1A

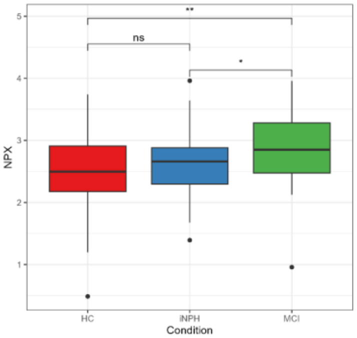


875) Gene Symbol: TNFRSF21

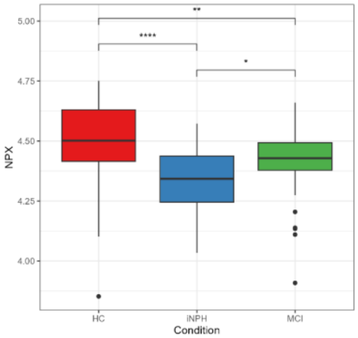


876) Gene Symbol: TNFRSF4

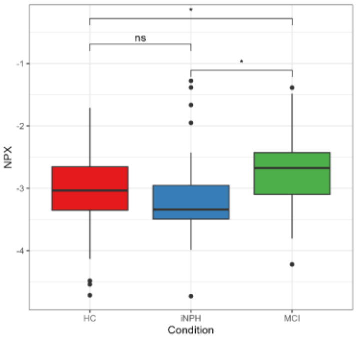


877) Gene Symbol: TNFRSF8

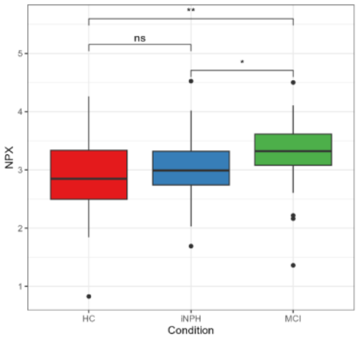


878) Gene Symbol: TNFSF11

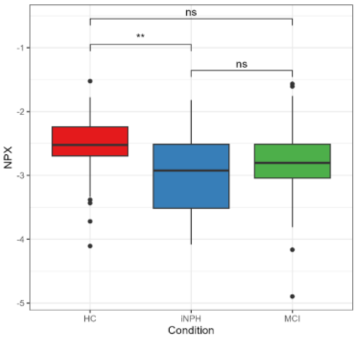


879) Gene Symbol: TNFSF13

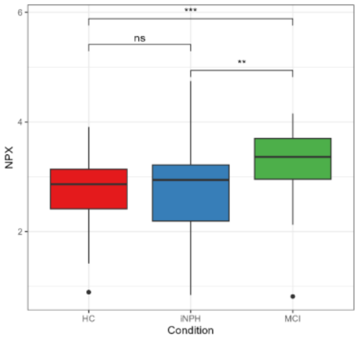


880) Gene Symbol: TNN

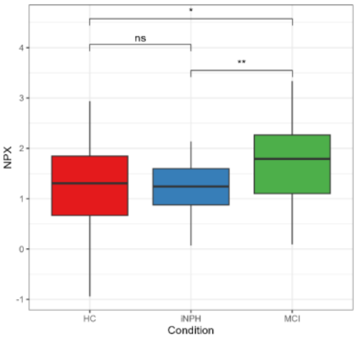

881) Gene Symbol: TNNI3

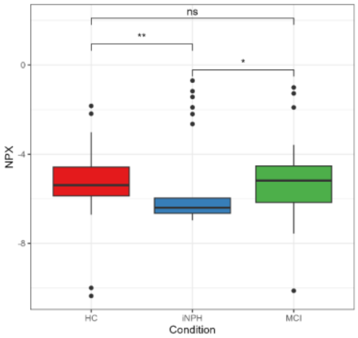


882) Gene Symbol: TP53BP1

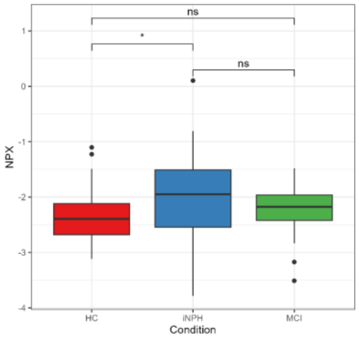


883) Gene Symbol: TP53I3

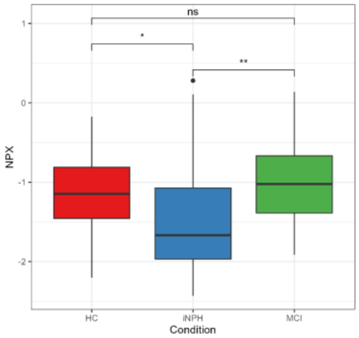


884) Gene Symbol: TPM3

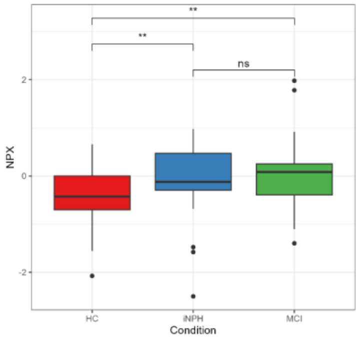


885) Gene Symbol: TPP1

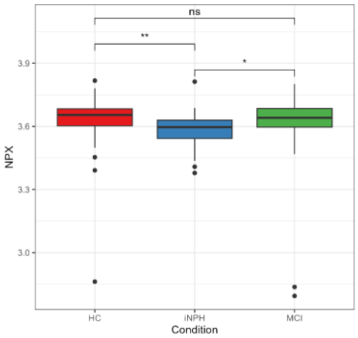


886) Gene Symbol: TPT1

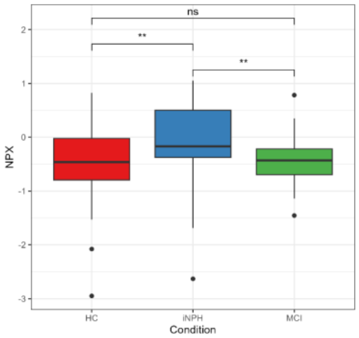


887) Gene Symbol: TREML2

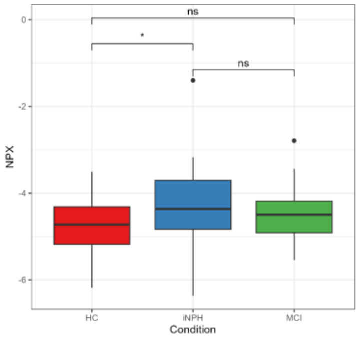


888) Gene Symbol: TRIAP1

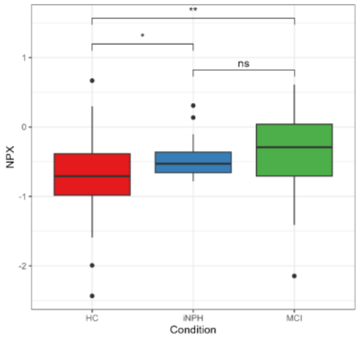


889) Gene Symbol: TRIM24

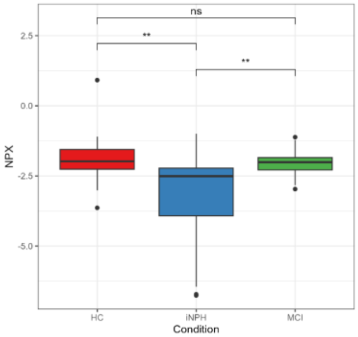


890) Gene Symbol: TRIM5

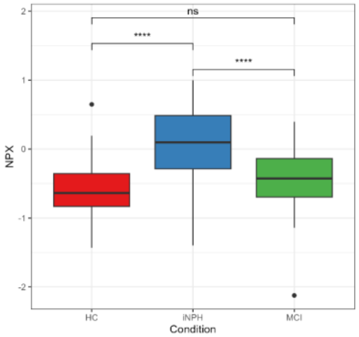


891) Gene Symbol: TSPAN7

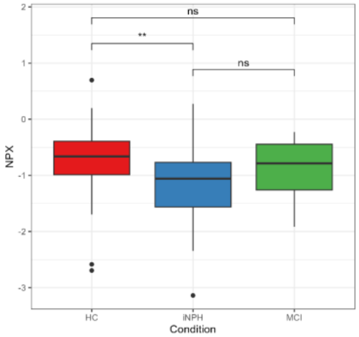


892) Gene Symbol: TTR

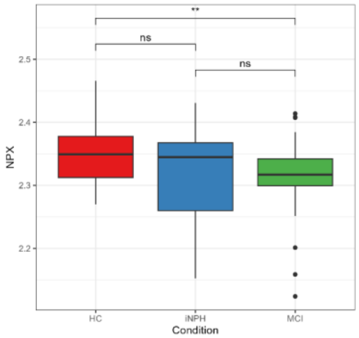


893) Gene Symbol: TYRO3

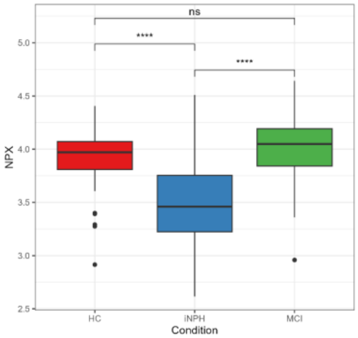


894) Gene Symbol: UBE2B

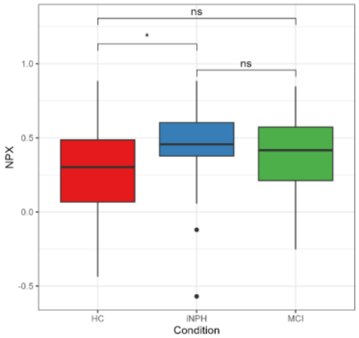


895) Gene Symbol: UBE2L6

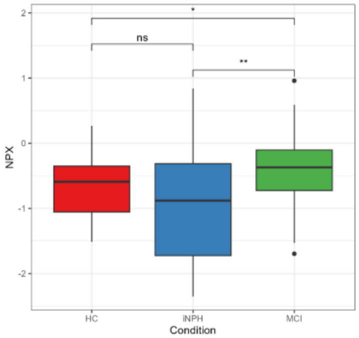


896) Gene Symbol: UFD1

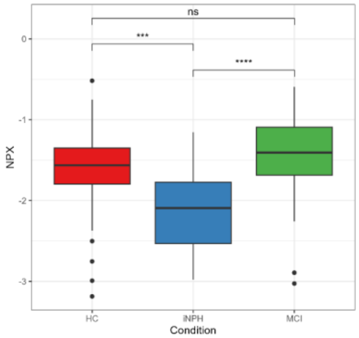


897) Gene Symbol: UGDH

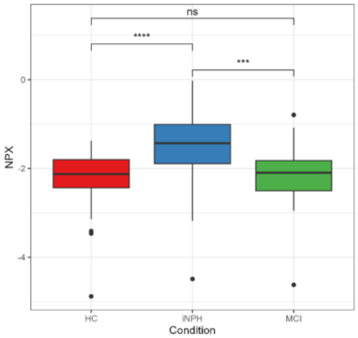


898) Gene Symbol: UNC5D

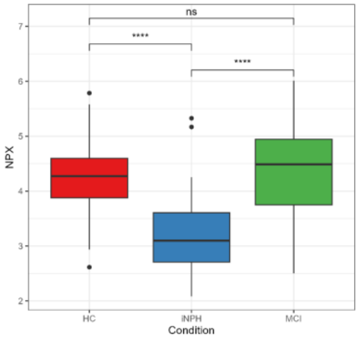


899) Gene Symbol: UNC79

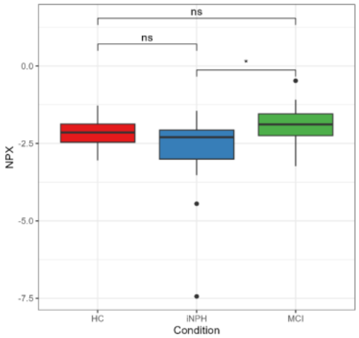


900) Gene Symbol: UNG

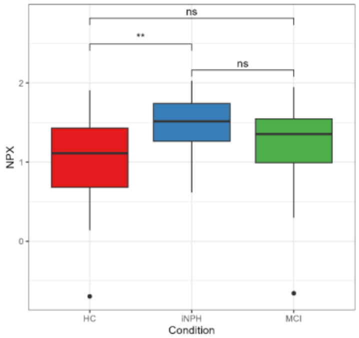

901) Gene Symbol: UPB1

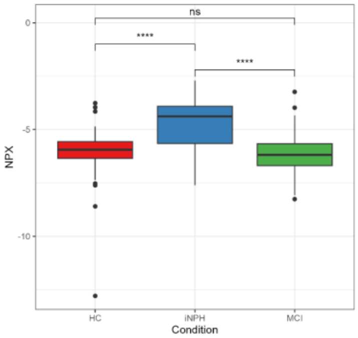


902) Gene Symbol: USP25

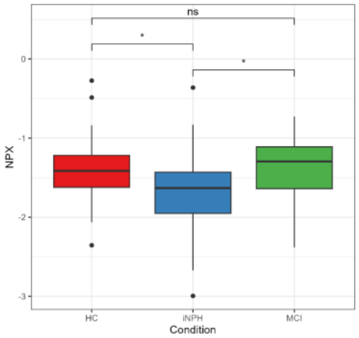


903) Gene Symbol: USP28

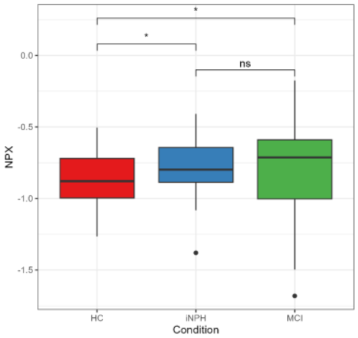


904) Gene Symbol: VASP

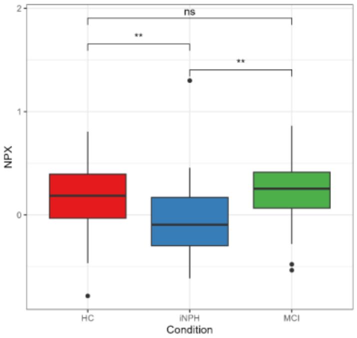


905) Gene Symbol: VCAM1

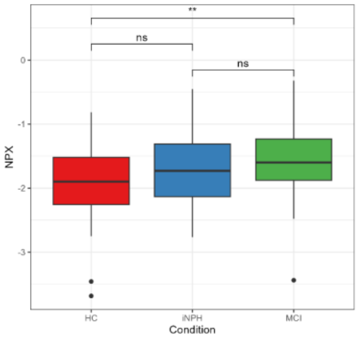


906) Gene Symbol: VCAN

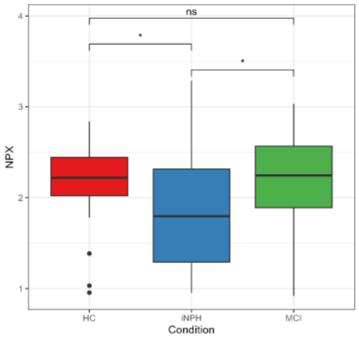


907) Gene Symbol: VEGFC

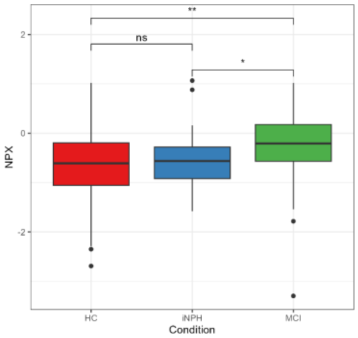


908) Gene Symbol: VGF

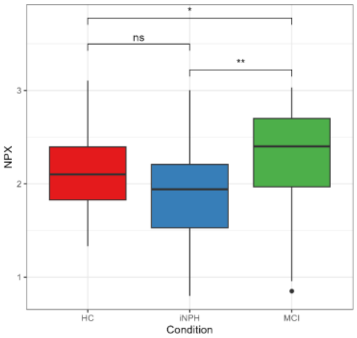


909) Gene Symbol: VIM

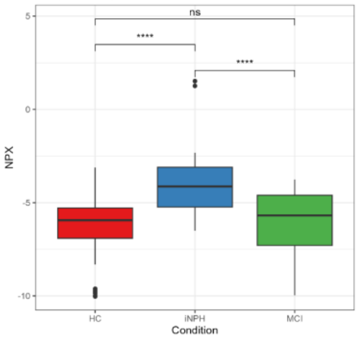


910) Gene Symbol: VSIG10

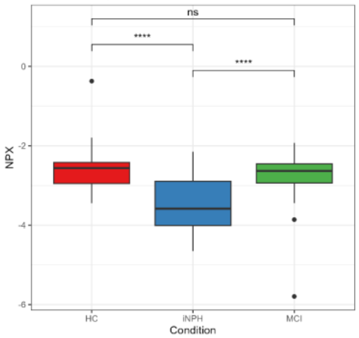


911) Gene Symbol: VSIR

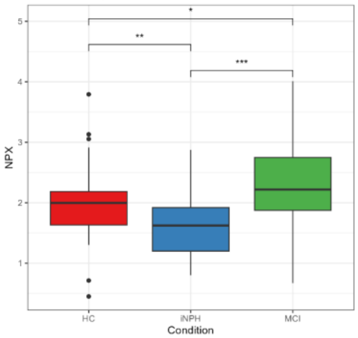


912) Gene Symbol: VSNL1

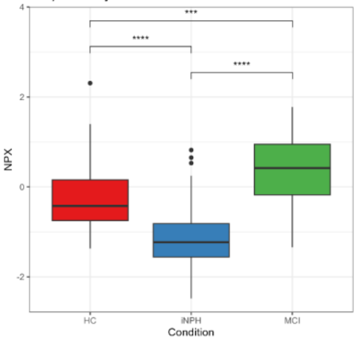


913) Gene Symbol: VTCN1

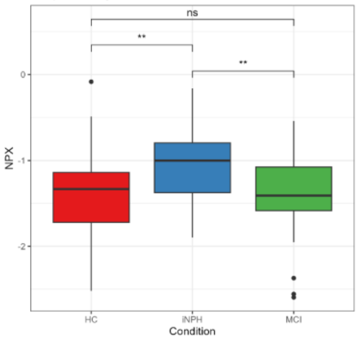


914) Gene Symbol: VWA5A

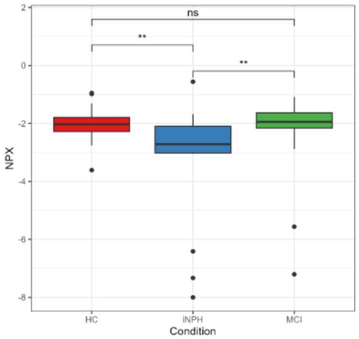


915) Gene Symbol: VWC2

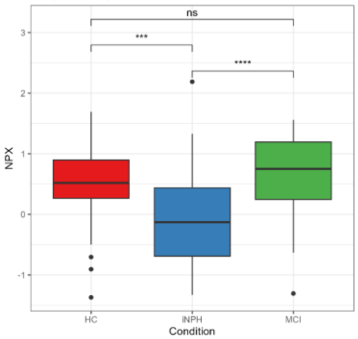


916) Gene Symbol: VWF

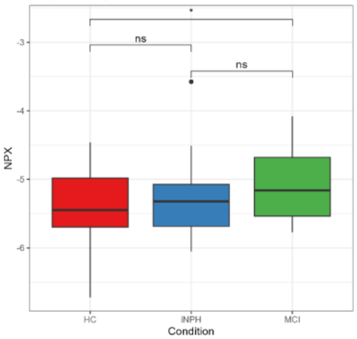


917) Gene Symbol: WASF3

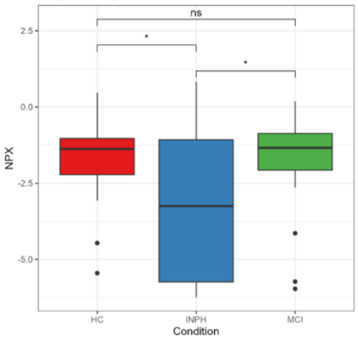


918) Gene Symbol: WASL

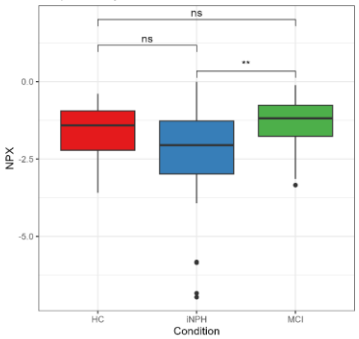


919) Gene Symbol: WDR46

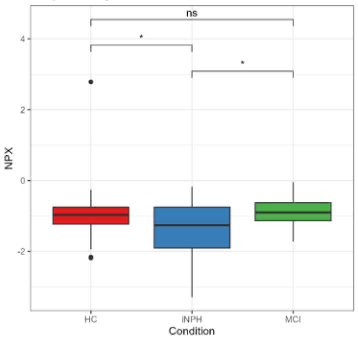


920) Gene Symbol: WFDC1

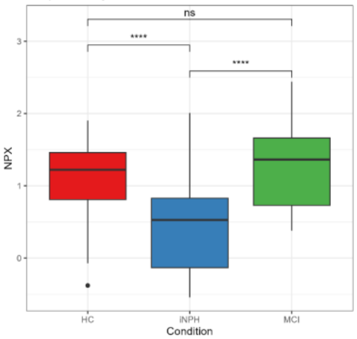

**Figure s4. Over-representation analysis (ORA) results Across Idiopathic Normal Pressure Hydrocephalus (iNPH), Mild Cognitive Impairment (MCI), and Healthy Control (HC) Conditions.** This figure represents the gene ontological (GO) terms of biological processes (BP) between iNPH and HC (a,b) and iNPH and MCI (c, d) that were significant (False discovery rate (FDR) < 0.05).

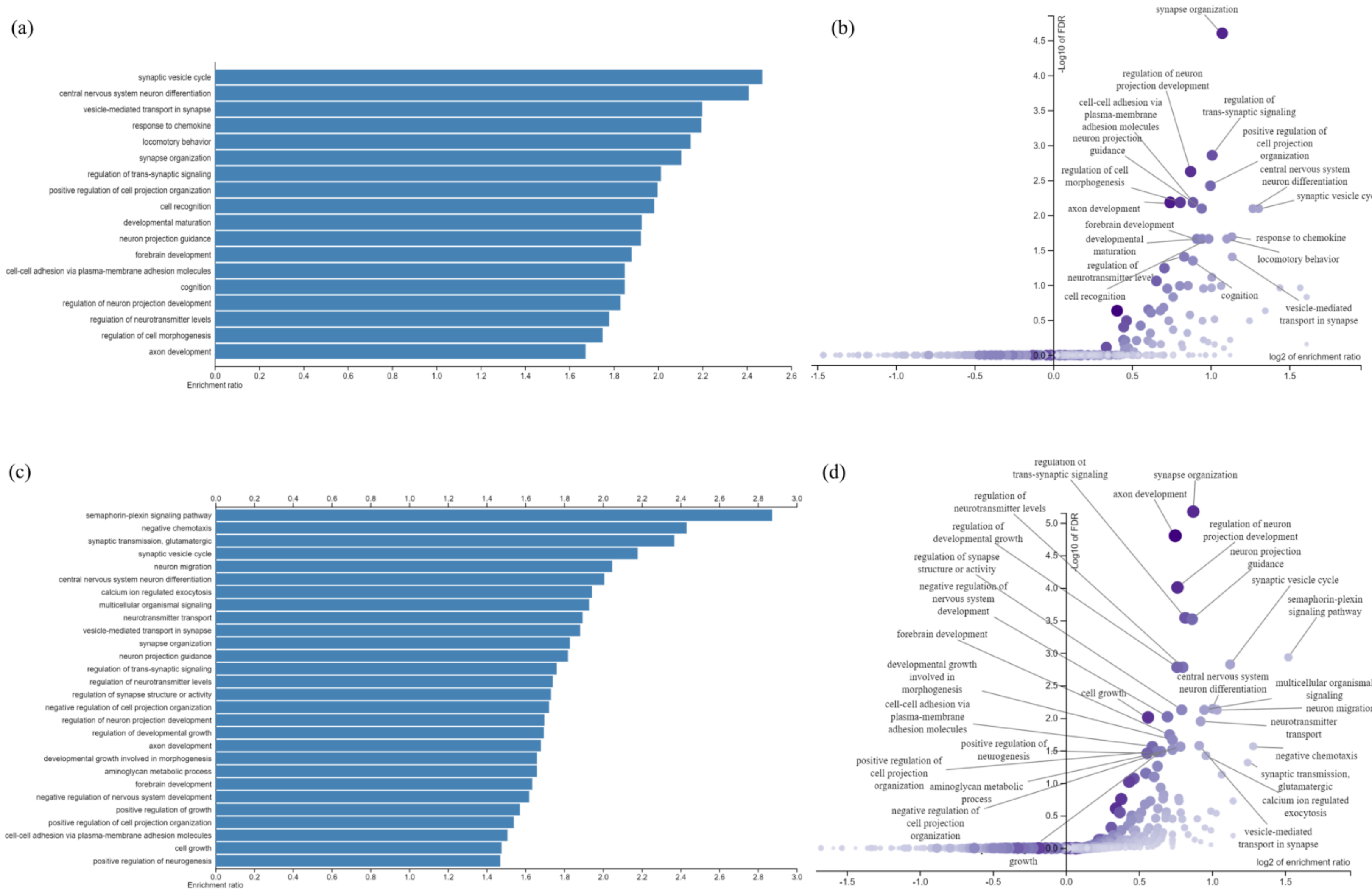

**Figure s5. Principal component analysis (PCA) plot of the of Normalized Protein Expression (NPX) Across Idiopathic Normal Pressure Hydrocephalus (iNPH), Mild Cognitive Impairment (MCI), and Healthy Control (HC) Conditions.** This figure presents a PCA plot based on the NPX data of all proteins, providing a visual representation of the variance and clustering of protein expression profiles across the three studied conditions: iNPH, MCI, and HC. Each data point corresponds to a sample, colored and labeled according to its respective condition.

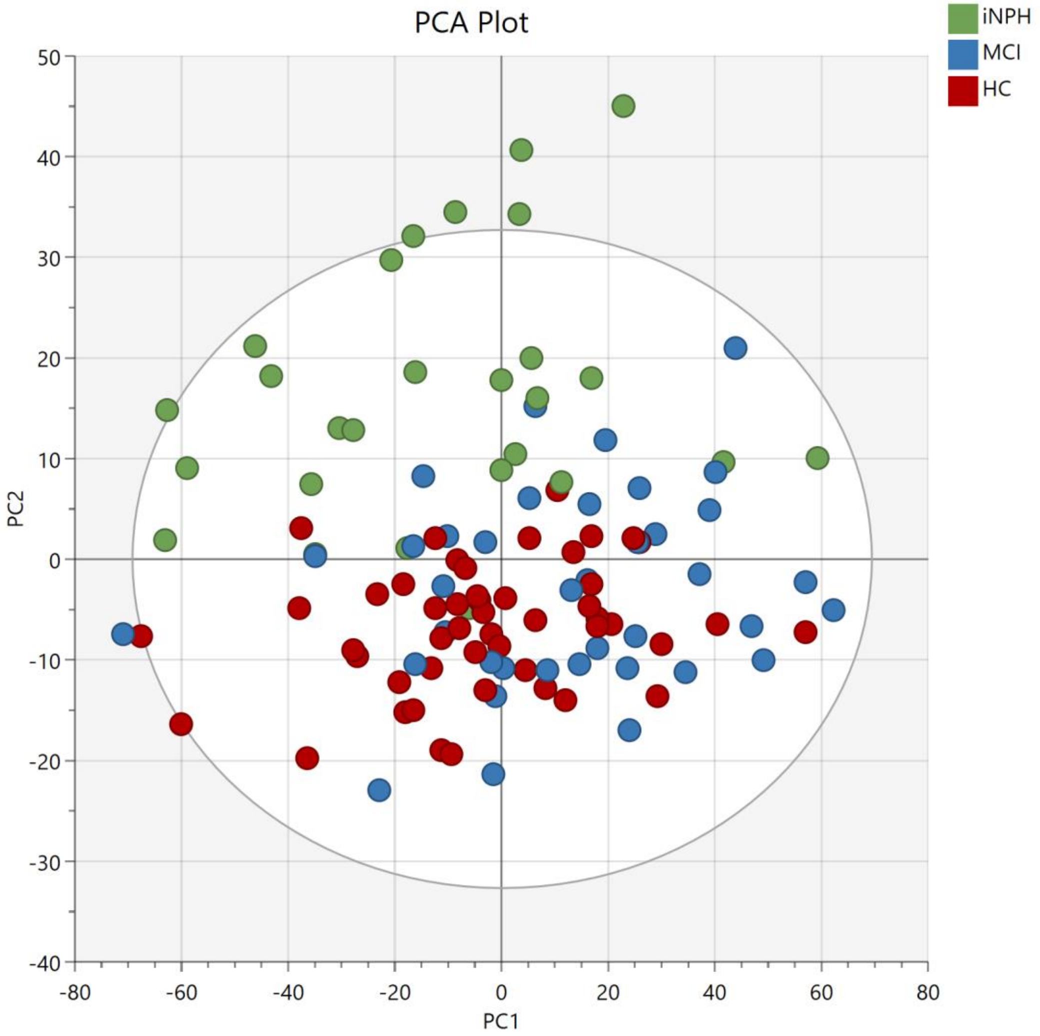
